# Brain charts for neuroanatomical sex differences across the human lifespan

**DOI:** 10.64898/2026.05.09.724029

**Authors:** Margaret Gardner, Lena Dorfschmidt, Dabriel Zimmerman, Sheila Shanmugan, Jakob Seidlitz, Richard A.I. Bethlehem, Laura Pritschet, Andrew A Chen, Lifespan Brain Chart Consortium, Armin Raznahan, Russell T Shinohara, Aaron Alexander-Bloch, the Alzheimer’s Disease Neuroimaging Initiative

## Abstract

Population-level sex differences in human neuroanatomy have been highly debated, with prior literature limited by narrow age windows and assumptions about regions’ scaling with total brain size. Building on recent work charting brain structures across the lifespan, we use magnetic resonance imaging data from over 100,000 individuals (51.7% F) to model normative reference trajectories of 241 structural features capturing age-varying sex effects. Results show replicable age-varying sex differences in nearly all features, and sex’s effects survive nonlinear correction for total brain size, with sex biases tending to increase across the lifespan. Notably, models are sensitive to atypicalities in six neuropsychiatric disorders. This work establishes and quantifies population-level sex biases in the largest-to-date study of sex differences in the human brain.

## Introduction

The availability of massive neuroimaging samples has led to recent advances in “brain charts”, population-scale normative reference models of imaging-derived phenotypes’ (IDPs’) trajectories across the human lifespan(*1–7*). These resources have provided unprecedented insight into population trends in brain structure, as well as benchmarks for quantifying individual variation used to identify patterns of deviation across neuropsychiatric disorders(*1*, *8*).

Typically, brain charts have been modeled with a nonlinear effect of age, controlling for sex and technical covariates(*1*, *5*). However, this approach does not capture age-varying sex effects, which have been described in a range of structural IDPs and developmental epochs(*9–18*). Failure to capture even subtle population-level differences between males and females – i.e. “biases” towards being larger or more variable in one sex – limits models’ accuracy in both sexes and may impair the ability to uncover disease biomarkers, particularly for the numerous disorders with sex biases in prevalence, course, and/or presentation(*19–21*). Moreover, directly quantifying sex differences in IDPs reveals the cumulative impact of sex- and gender-related phenomena on brain structures (**Box 1**), enabling targeted assessments of regions or ages with prominent sex biases and aiding interpretation of findings in mixed-sex samples.

However, the presence of sex biases in neuroanatomy remains widely debated(*22–25*). Inconsistent results may be attributable to small, underpowered samples; limited age ranges; and variable methods to account for sex differences in overall brain size (see DeCasien et al(*21*) for review). These limitations are addressable using large datasets and nonlinear corrections for allometry, or how regions scale with total brain size(*15*, *21*, *26*), yet, such methods remain only sparsely applied.

Here, we aim to overcome these common limitations and leverage the power of normative modeling in the largest-to-date study of sex differences in the human brain. We fit brain charts incorporating age-varying sex effects in the Lifespan Brain Chart Consortium, a global sample of anatomical scans of over 100,000 individuals from the second trimester of gestation to 100 years of age(*1*, *7*). While some prior brain charts have been fitted separately in each sex to allow independent age trajectories(*2–4*), we fit “sex-moderated” brain charts with a nonlinear moderator to directly quantify sex effects’ significance and increase models’ power. Next, we re-fit brain charts using a novel semiparametric method of controlling for total brain size, which we used to quantify sex biases in median and variability across the lifespan. We also demonstrate the utility of individual reference scores derived from sex-moderated brain charts to identify deviations in six neuropsychiatric disorders. We hypothesized that sex-moderated brain charts would better explain variability in IDPs than traditional models without age-varying sex effects and that, for most IDPs, sex effects would remain significant even when controlling for overall brain size. Moreover, we hypothesized that reference scores derived from sex-moderated brain charts would be more sensitive to disease effects than traditional models. Sex-moderated and total-size-corrected brain charts for each IDP are available at www.brainchart.io (upon publication) to derive reference scores for new neuroimaging samples.

### Box 1

**Sex and Gender**

Here we investigate biases in neuroanatomy by sex, a biological phenomenon based on anatomical and physiological traits usually associated with sex chromosomes(*27*). Studies in humans and animal models have demonstrated that sex directly impacts neuroanatomy via gene expression and sex hormone concentrations(*28–32*). The social phenomenon of gender – which encompasses gender identity, expression, and related environmental factors(*27*) – is also critically important to neuroscience and psychology(*33*, *34*). Here we focus on sex differences, as sex was most frequently recorded in consortium studies. However, we acknowledge that the sex differences reported may result from any combination of sex- or gender-related differences in genetic, hormonal, environmental, or social factors(*20*, *21*, *33*, *35*). More systematic assessments of sex- and gender-related covariates can help disambiguate the causes and potential functional or clinical consequences of these neuroanatomical biases.

## Results

### Sex effects vary significantly with age in nearly all brain structures

We assessed the significance of age-by-sex interactions on brain structure using split-half cross-validation. Briefly, we used a global consortium of over 100,000 structural MRI scans spanning the human lifespan to derive neuroanatomical measures of: 7 global phenotypes; 30 subcortical volumes; and thickness, surface area, and volume measures for 68 cortical regions. After quality control, we fit a generalized additive model for location, scale and shape (GAMLSS) for each IDP using penalized splines to capture nonlinear age and age-by-sex interaction effects (**Fig 1A**, Extended Figures). Model selection was performed for each IDP in one half of the data (training) to establish additional covariates and penalties on nonlinear terms. The best-fit model was then fitted in the second half of the dataset (testing) to assess age-by-sex interaction’s significance via likelihood ratio tests. This procedure was done twice per IDP, swapping the train and test sets. As results were highly consistent across split-halves, we visualize test sample B with full results in Supplement Section A.

**Figure 1.**
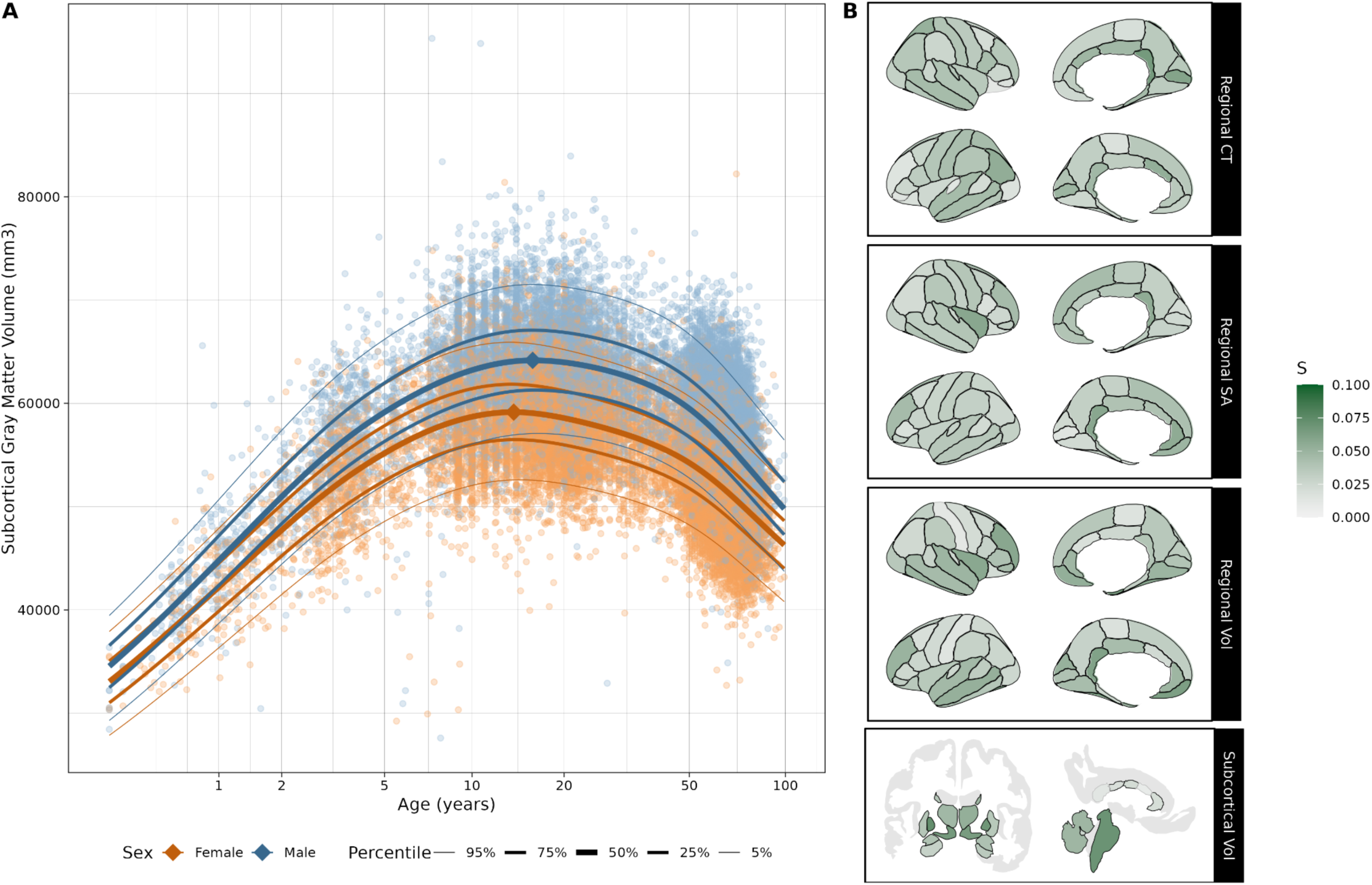
Nearly all neuroanatomical features exhibit sex biases that vary across the lifespan. **(A)** An exemplar brain chart quantifying the distribution of subcortical gray matter volume in females (orange) and males (blue) across the lifespan. Line thicknesses correspond to the population percentile, while diamonds indicate peak volumes for each sex. **(B)** Magnitude of the age-varying component of sex (age-by-sex term) on IDPs in test sample B. Effect sizes (S) are reported using the robust effect size index (RESI). Black outlines indicate significant effects at p <0.05, using false discovery rate (FDR) correction for multiple comparisons. Abbrv: CT, cortical thickness; SA, surface area; Vol, volume.

In nearly all IDPs, significant age-varying sex effects replicated across both split halves (Split Half A, 233 of 240 converged features, mean effect size S(*36*)=0.037; Split Half B, 232 of 241 converged features, mean S=0.036; **Figs 1B** & S5, Supplemental Data S1).

We also performed two sets of sensitivity analyses. First, in addition to our quality control procedures, we assessed sensitivity to scans’ segmentation quality by refitting training and testing models, weighting each scan inversely to its normed surface hole count(*37*). Second, given documented challenges of accurately quantifying cortical thickness in infants(*38–40*), we refit models of thickness IDPs excluding individuals younger than two years old. Results were consistent with our main analyses and are presented in Supplement Section C.

### Sex biases are not explained by total brain size

We next tested whether sex effects survived when controlling for total brain size, as sex differences have been suggested to primarily reflect males’ larger average brain volume(*21*, *24*). We conducted the same model selection procedure described above, expanding training models’ search space to include a nonlinear effect of total size in up to three statistical moments. For volumetric and other global IDPs (i.e. mean cortical thickness and total surface area), total size was operationalized as total brain volume; for regional cortical thickness models, total size was operationalized as mean cortical thickness; and for regional surface area models, it was operationalized as total surface area. Following model selection, we fit models in the held-out test split to chart normative trajectories (**Fig 2A**), including differences in each sex’s median and variability (**Figs 2B-C**). We again used likelihood ratio tests to probe the significance of all sex effects, as well as sex’s age-varying component specifically. Even when controlling for nonlinear scaling effects, sex was robustly and replicably significant in nearly every IDP (A, 238 of 239 converged features, mean S=0.070; B, 239 of 239 features, mean S=0.070; **Figs 2D** & S6). The age-varying component of sex also remained significant in most IDPs (A, 161 of 239 features, mean S = 0.034; B, 169 of 239 features, mean S=0.032; Fig S7).

**Figure 2.**
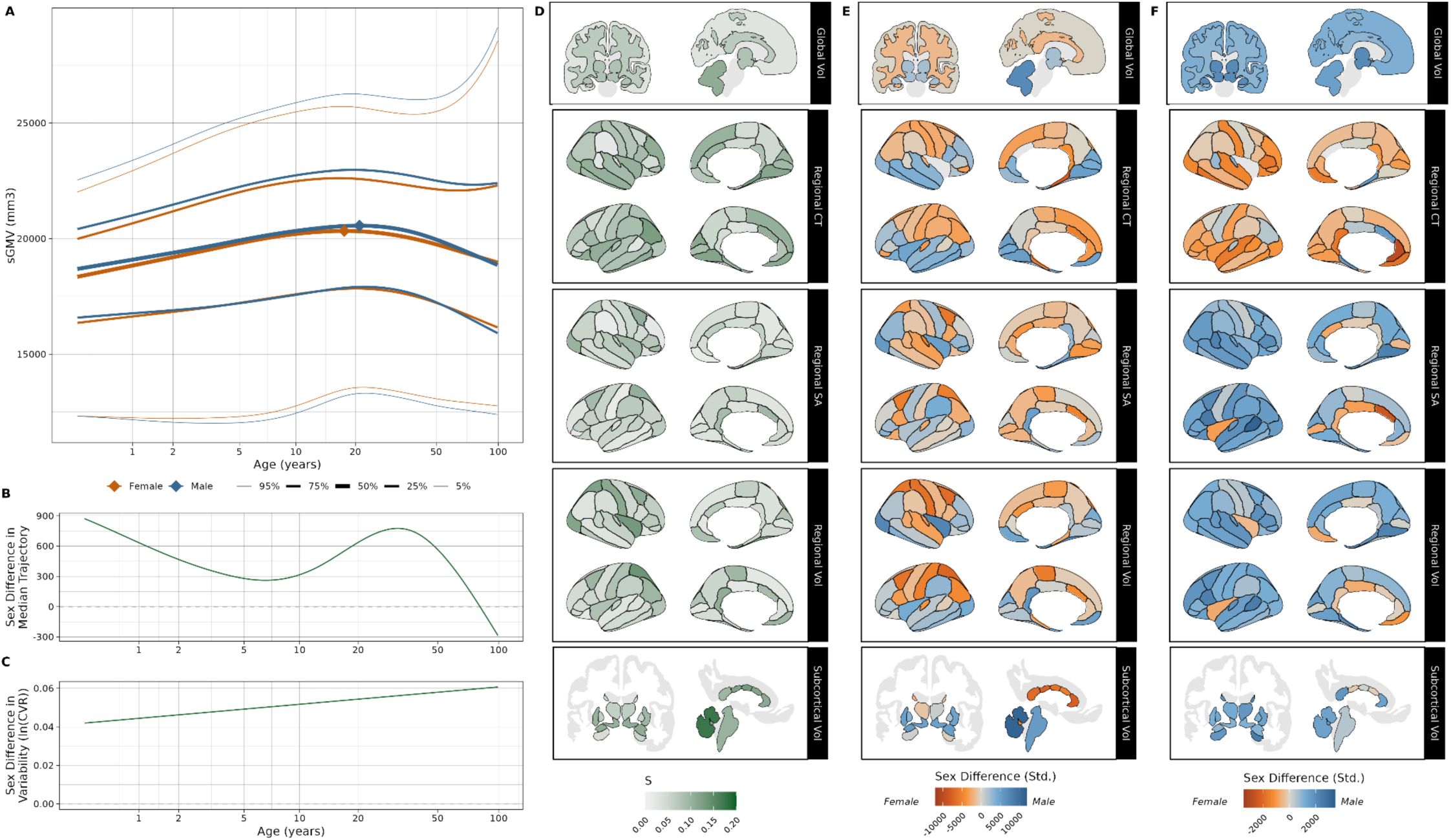
Sex biases in neuroanatomy persist when accounting for scaling effects. **(A)** An exemplar brain chart of subcortical gray matter volume (sGMV), residualizing nonlinear total brain volume effects. Line thicknesses correspond to the population percentile, while diamonds indicate peak volumes for each sex. **(B)** Sex difference in the median trajectory of subcortical gray matter volume over the lifespan, with positive values indicating male bias. **(C)** Sex differences in variability quantified as the log coefficient of variation ratio (CVR) across the lifespan. Positive values indicate greater variability in males. **(D)** Effect size (S) using the robust effect size index (RESI) of sex (both age-by-sex and sex intercept terms) after controlling for total size, in test sample B. **(E - F)** Sex bias in IDP median (E) and variability (F) across the lifespan controlling for total size, standardized by scaling relative to each IDP’s standard deviation. Black outlines indicate significant effects at p <0.05, FDR-corrected. Abbrv: CT, cortical thickness; SA, surface area; Vol, volume; Std., standardized

To probe the directionality of sex biases and their dynamics across the lifespan, we charted trajectories of sex biases in each IDP. Specifically, we plotted standardized differences between male and female median trajectories (i.e. 50th centile line; **Fig 2B**), as well as the ratio of male and female coefficients of variation (**Fig 2C**). We then summarized each IDP’s tendency to be male or female-biased by integrating these difference trajectories, with positive values indicating that males tend to be larger or more variable than females. We found that IDPs are almost equally likely to be male or female biased in their median values (A, 51.0% male biased; B, 53.1% male biased; **Fig 2E** & S8), while most are more variable in males (A, 71.1% male biased; B, 67.8% male biased; **Fig 2F** & S9).

### Sex biases tend to increase across the lifespan

Next, we queried how sex biases change across the lifespan by grouping IDPs by these total-size-corrected sex difference trajectories. We first assessed how long each IDP was male- or female-biased, revealing that for both median and variability, most were consistently biased towards a single sex for over 95% of the lifespan (Figs S10 & S11). Beyond a consistent male or female bias, IDPs with significant age-by-sex effects were subcategorized as “Diverging” or “Converging”, respectively, if sex differences’ magnitude increased or decreased with age. IDPs with nonsignificant age-by-sex effects when controlling for total brain size were noted as “Stable”. Finally, the remaining IDPs – which had significant age-moderated sex effects but were not consistently biased for more than 95% of the lifespan – were categorized as “Switch” if they were biased towards different sexes at the beginning and end of the lifespan. A small minority of IDPs with more complex trajectories were labeled as “Complex”. We used these groupings (median, **Fig 3A**, Fig S12; variability, Fig S13) to quantify common sex-bias trajectories (median sex-biases, **Fig 3B**, Fig S12; variability sex-biases, Fig S13) and summarized trends within each group (median, **Fig 3C**, Fig S12; variability, Fig S13).

**Figure 3.**
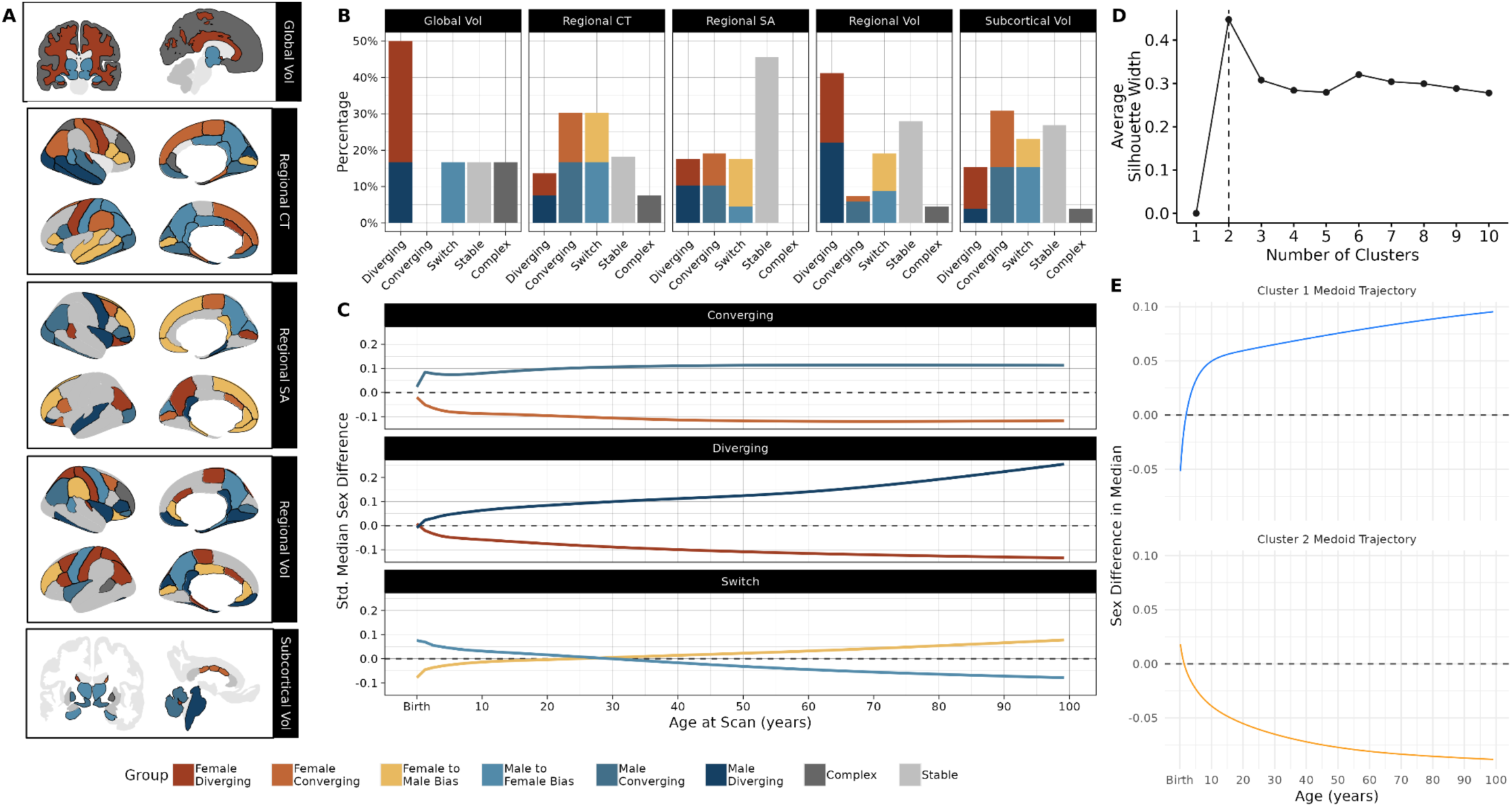
Grouping on total-size-corrected sex bias trajectories reveals that male and female medians tend to diverge across the lifespan. **(A)** Imaging-derived phenotypes (IDPs) colored by groups indicating the trajectory of sex biases controlling for total size: female or male diverging, where features that are consistently biased towards one sex become more biased with age; female or male converging, where bias decreases over time; female bias to male bias or male to female bias, where bias changes from one sex to another; stable, where sex biases do not significantly vary with age; complex, for significantly age-varying sex biases that do not fit any previously defined groups. **(B)** Bar plots of IDPs in each group **(C)** Sex bias trajectories summarized within each group. Summaries were created by standardizing sex-difference trajectories within each IDP before fitting a simple generalized additive model to constituent IDPs’ sex bias estimates at each age. **(D)** Plot of average silhouette width by number of k-medoid clusters, indicating a two-cluster solution. **(E)** Scaled sex-bias trajectories of cluster medoids for 2-cluster solution: left parahippocampal volume for the diverging female cluster (lower) and right pars triangularis volume for the diverging male cluster (upper). Abbrv: CT, cortical thickness; SA, surface area; Vol, volume; Std., standardized

We found that among IDPs with age-varying sex effects, sex difference tends to increase in magnitude (A, Diverging = 33.1%; B, Diverging = 25.5%; **Fig 3B**, Fig S12). Grouping by sex differences in variability produced similar results, with roughly half of all features either becoming increasingly male-biased (A, 21.3%, B, 17.6%) or remaining stable (A, 32.6%, B, 29.3%; Fig S13).

We complemented this *a priori* grouping with data-driven k-medoids clustering, which minimizes within-cluster distance between regions’ sex difference trajectories. In each split-half, we identified two clusters as the optimal solution for both median and variability sex differences, one diverging male and one diverging female (median, **Fig 3D-E**, Fig S14 & S15; variability, Fig S16 & S17). In total, these results suggest that population-level differences in females and males tend to increase across the lifespan.

### Neuroanatomical sex biases in context

We identify points of interest in global phenotypes’ sex bias trajectories, controlling for total brain size, alongside critical reproductive, neurodevelopmental, and psychiatric milestones (**Fig 4**). Interestingly, subcortical gray matter volumes and growth rates peak earlier in females than in males, mirroring females’ earlier average pubertal onset(*41*). Given high subcortical concentrations of sex hormone receptors(*41*), we speculate that these sex biases may be driven by pubertal processes. Meanwhile, white matter sex biases in middle- and late-life align with recently-described sex differences in structural connectivity(*42*). Finally, we observe a distinct change in cortical gray matter volume’s sex bias trajectory around midlife, when it transitions from increasingly female-biased to become male-biased, possibly driven by volume reductions reported during menopause(*43*).

**Figure 4.**
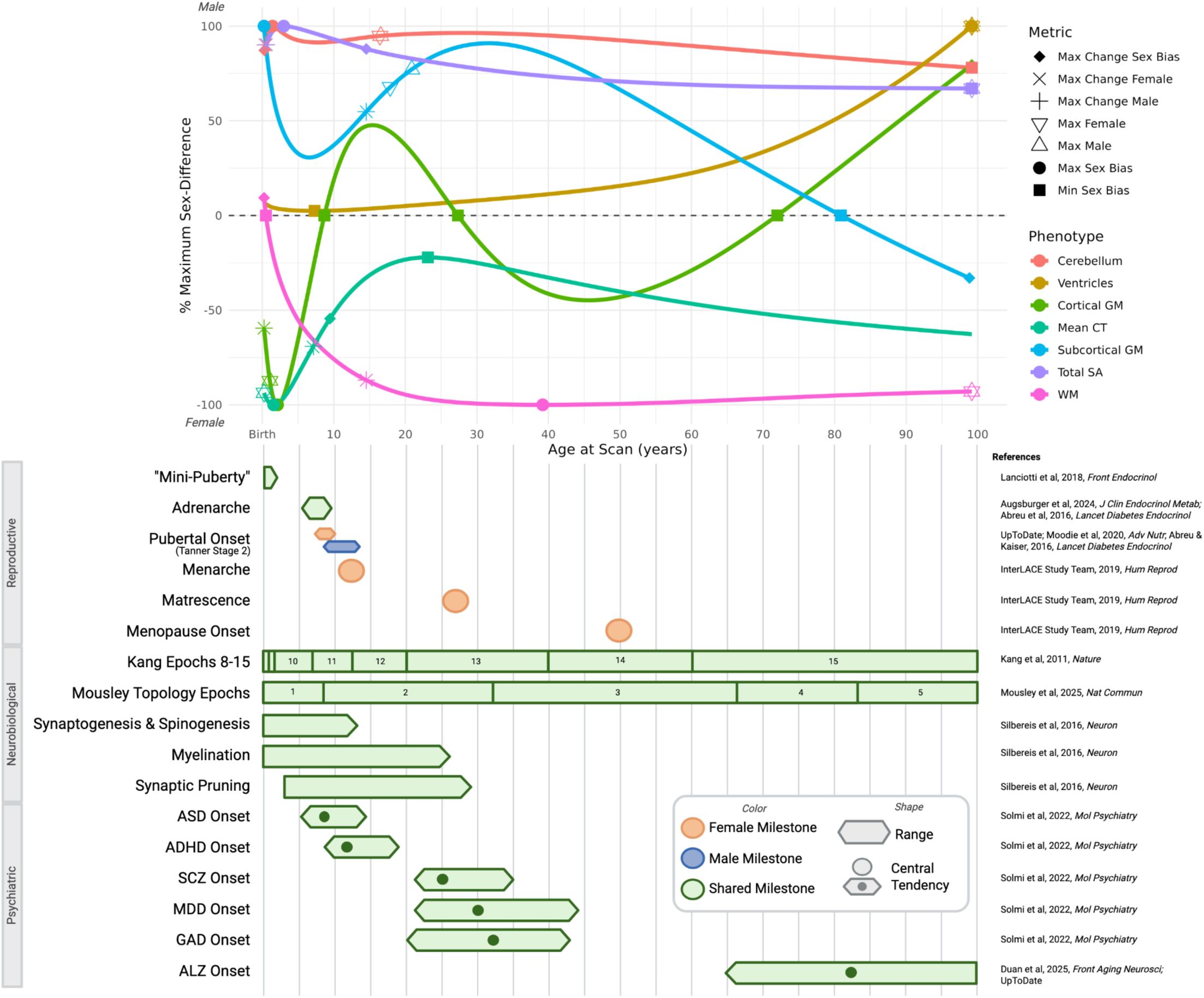
Sex bias trajectories of total-size-corrected global phenotypes in the context of reproductive, neurobiological, and psychiatric milestones. Top, a graphical summary of sex differences in the median trajectories of global neuroanatomical phenotypes across the lifespan. Symbols denote the age of key neuroanatomical peaks. Trajectories are visualized by averaging across two split-halves and scaling relative to their maximum sex bias. Bottom, a graphical summary of reproductive, neurobiological, and psychiatric milestones derived from the literature. Bars represent age ranges while circles indicate mean or median reported age; see Supplement for further discussion. Abbrv: ADHD, attention-deficit/hyperactivity disorder; ALZ, Alzheimer’s disease; ASD, autism spectrum disorder; GAD, generalized anxiety disorder; MDD, major depressive disorder; SCZ, schizophrenia; CT, cortical thickness; SA, surface area; Vol, volume. Figure created with BioRender.

### Brain charts with age-varying sex biases distinguish structural deviations in clinical samples

We assessed whether reference scores benchmarked against sex-moderated brain charts were sensitive to neuroanatomical deviations in six neuropsychiatric disorders. Here, we used large (N > 1,000) samples of individuals diagnosed with schizophrenia, Alzheimer’s disease, autism spectrum disorder, major depressive disorder, generalized anxiety disorder, or attention-deficit/hyperactivity disorder(Fig S3, Table S4).

In each diagnosis, t-tests comparing centile z-scores revealed case-control differences in neuroanatomical deviations (**Fig 5A & B**; Supplemental Data S2). These both replicated known clinical differences, such as extensive IDP reductions among those with Alzheimer’s and schizophrenia, and revealed subtler effects, including increased surface area and decreased thickness in frontal and temporal regions in major depressive disorder, as well as increased volume and surface area in temporal and parietal regions in generalized anxiety disorder.

**Figure 5.**
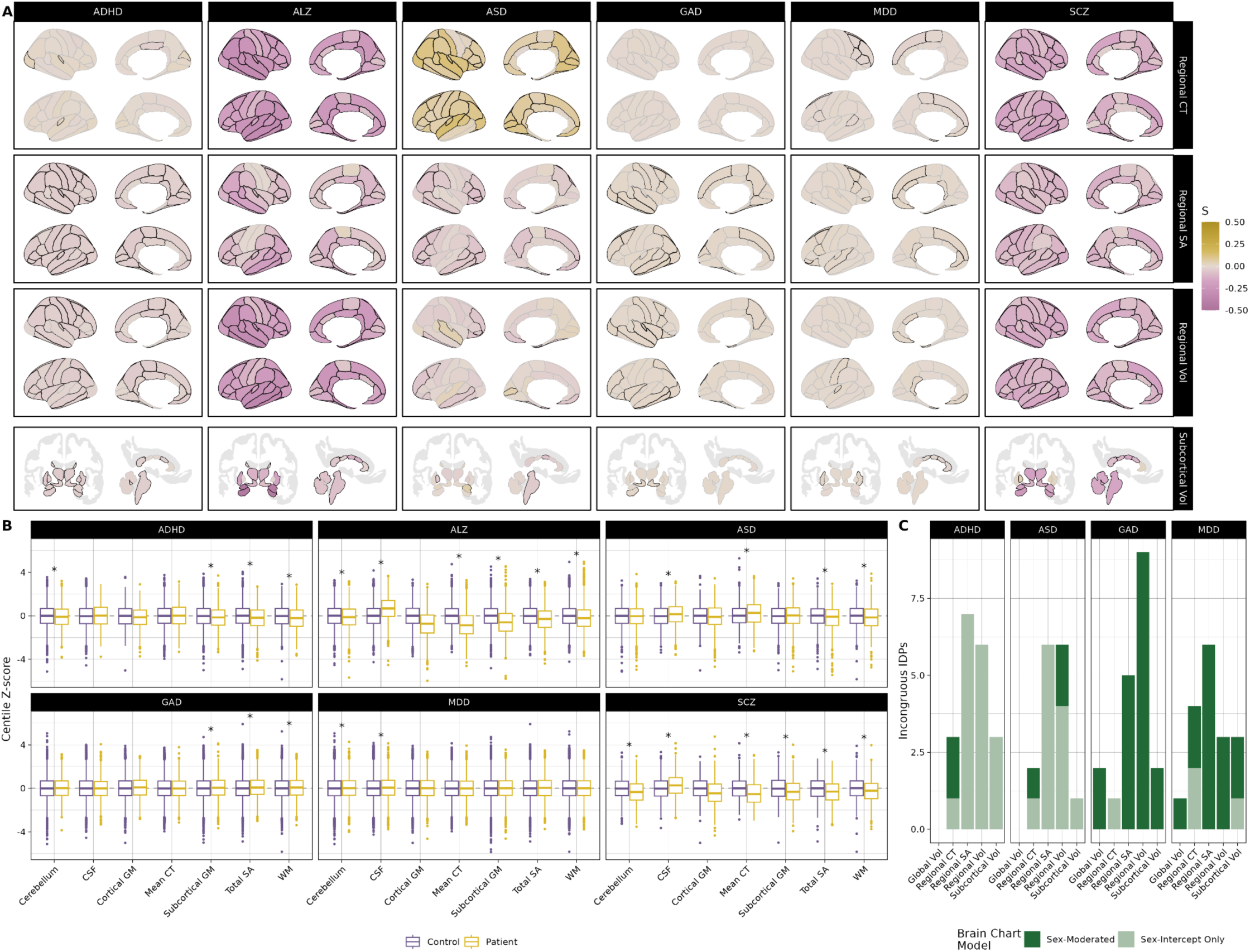
Brain charts reveal structural deviations in clinical populations, with notable differences when accounting for age-varying sex biases. **(A)** Case-control differences in centile z-score. Effect size (S) using the robust effect size index (RESI), with positive values indicating cases tend to be larger than controls and negative values indicating they tend to be smaller. Black outlines indicate significant effects at p <0.05, FDR-corrected. **(B)** Case-control differences in centile z-scores of global imaging-derived phenotypes (IDPs). Asterisks indicate significant effects at p < 0.05, FDR-corrected. **(C)** Bar plots showing discrepant case-control results across sex-moderated and sex-intercept models. Y-axis quantifies IDPs with significant diagnostic effects only when centile z-scores are derived from brain charts with age-varying sex effects (dark green) or from traditional sex-intercept brain charts (light green). Abbrv: ADHD, attention-deficit/hyperactivity disorder; ALZ, Alzheimer’s disease; ASD, autism spectrum disorder; GAD, generalized anxiety disorder; MDD, major depressive disorder; SCZ, schizophrenia; CT, cortical thickness; SA, surface area; Vol, volume.

We also assessed whether individuals with each diagnosis were over- or under-represented among extreme percentiles, i.e., more or less likely than controls to be below the 5th percentile or above the 95th percentile for a given IDP, which produced similar results (Fig S18). We further quantified regional disease-associated atypicalities by testing case-control differences in reference scores derived from total-size-corrected models (Figs S31-S32).

Finally, we queried whether benchmarking against sex-moderated brain charts impacted case-control analyses relative to traditional sex-intercept models, which hold sex-effects constant across the lifespan. While results were largely consistent (Figs S19-S30), sex-intercept models overestimated regions with case-control differences in attention-deficit/hyperactivity and autism spectrum disorders, while failing to identify deviations in generalized anxiety and major depressive disorders (**Fig 5C**). This suggests that failure to account for age-varying sex differences in neuroanatomy may reduce the specificity of biomarkers of male-biased disorders in younger samples and sensitivity to atypicalities in female-biased disorders with predominantly adult samples.

## Discussion

We use normative reference models trained on a global neuroimaging sample and split-half cross-validation to establish and quantify sex biases in neuroanatomical imaging-derived phenotypes (IDPs) across the human lifespan. First, we show that sex differences vary across the lifespan in nearly all IDPs. Next, we demonstrate that sex differences remain even when robustly controlling for total brain size, such that neuroanatomical sex biases are not reducible to differences in males’ and females’ average brain size. Finally, we show that models accounting for age-moderated sex bias are sensitive to disease effects and may yield improvements over reference scores derived from sex-intercept models. Together, this work resolves several outstanding questions in the existing literature on sex differences in human neuroanatomy and, to our knowledge, is the largest study of sex differences in human neuroanatomy to date.

We emphasize that population-level sex biases in structure do not necessarily imply differences in function(*44*), nor do they indicate that brain regions are better or more evolved in one sex than another. Similarly, while brain charts can produce individualized reference scores, one’s score does not indicate a more masculine versus feminine brain. While our results clearly demonstrate the advantages of incorporating sex – including its age-varying component – in analyses of neuroanatomy, sex differences must be interpreted in the context of myriad factors known to influence brain structure(*21*).

Our results provide strong, replicable evidence for sex biases in population neuroanatomical trajectories and clearly establish that sex biases are not artifacts of total size differences. We also replicate reports that roughly equal numbers of regions are male- and female-biased after correcting for total brain size, with male-biases in subcortex and occipital lobes and female-biases in medial structures and parietal lobes(*15*, *21*, *45*), as well as greater variability among males(*15*, *26*). We further find that sex biases tend to increase across the lifespan.

We hypothesize that this may reflect an accumulation of sex-biased hormonal and environmental exposures, including societal factors and reproductive events(*33*, *34*, *41*, *46*)(**Fig 4**). Interestingly, we found age-by-sex effects on variability in roughly 70% of IDPs, while the largest prior study of sex differences in neuroanatomical variance reported that sex biases tended to remain stable or decrease with age(*26*). This difference could be due to our order-of-magnitude larger sample size(*47*), our sample’s expanded age range(26), and/or GAMLSS’ ability to directly model variability rather than relying on residuals(*1*, *48*). While the current study cannot parse the mechanisms shaping these sex biases across the lifespan, we hope they will help target future experimental work.

Finally, we show that failure to account for age-varying sex effects can alter case-control differences in reference scores. While our case-control tests of scores derived from sex-intercept versus sex-moderated models were largely consistent, we found that traditional methods might yield more false positives in younger samples of male-biased disorders and false negatives in older samples of female-biased disorders (**Fig 5C**). To facilitate incorporating these age-varying sex differences into future work, reference scoring for out-of-sample neuroimaging data using the present models is available at www.brainchart.io.

This work has several limitations. First, we categorized individuals based only on sex reported by primary studies and cannot determine to what extent sex biases reported here are due to sex, gender, or related genetic, hormonal, psychosocial, or environmental factors(*21*). Similarly, we do not include individuals with differences in sexual development (previously termed “intersex”) due to insufficient sample size even in our consortium dataset. Next, overrepresentation of Western, educated, affluent individuals in consortium data may limit the generalizability of findings(*6*, *49*, *50*). Finally, the relative scarcity of neuroimaging data in fetal and infant subjects limits our ability to accurately resolve sex biases very early in life.

Despite these limitations, the present study contributes substantially to our knowledge of the presence of sex differences in human neuroanatomy and their temporal dynamics. Leveraging massive consortium data, normative modeling, semiparametric allometric scaling, and cross-validation, we show that age-varying sex biases are present in nearly all brain structures and that sex effects are not reducible to size differences. We further use individualized deviation scores to map sex-specific disease effects across six psychiatric disorders. Sex-moderated and total-size-corrected brain charts for each IDP are available at www.brainchart.io (upon publication) to derive reference scores for new neuroimaging data.

## Acknowledgements

We would like to acknowledge the individuals and organizations that have made the data used for this research available, including the following consortia: Lifespan Brain Chart Consortium (LBCC); Aging Brain: Vasculature, Ischemia, and Behavior Study (ABVIB); Alzheimer’s Disease Neuroimaging Initiative (ADNI); Australian Imaging, Biomarkers and Lifestyle (AIBL); Alzheimer’s Disease Repository Without Borders (ARWiBo); Biomarkers of Cognitive Decline among Normal Individuals (BIOCARD); Centre for Attention Learning and Memory (CALM); Cambridge Center for Ageing and Neuroscience (Cam-CAN); Chinese Color Nest Project (CCNP); Centers of Biomedical Research Excellence (COBRE); Consortium on Vulnerability to Externalizing Disorders and Addictions (cVEDA); Harvard Aging Brain Study (HABS); International Consortium for Brain Mapping (ICBM); IMAGEN; Neuroscience in Psychiatry Network (NSPN); Province of Ontario Neurodevelopmental Disorders (POND) Network; Pre-symptomatic Evaluation of Experimental or Novel Treatments for Alzheimer Disease (PREVENT-AD); and Vietnam Era Twin Study of Aging (VETSA). Lists of members and their affiliations appear in the Supplement. The consortium investigators who designed and implemented these studies and/or provided data did not necessarily participate in the analysis or writing of this report. This manuscript reflects the views of the authors and may not reflect the opinions or views of the investigators or funders providing data, including those acknowledged below.

## Funding

National Institutes of Health grant R01 MH133843 (A.A.B)

National Institutes of Health grant F31 MH136691 (M.G.)

FOCUS on Health and Leadership for Women and Penn PROMOTES Research on Sex and Gender in Health grant RAising the Investment in Sex and gender Evidence (RAISE) Pilot Grant Award (A.A.B., M.G., R.T.S., S.S.)

Burroughs Wellcome Fund Career Award for Medical Scientists (S.S)

Foundation for Women’s Health Award (S.S)

National Institutes of Health grant DP5 OD036142 (S.S.)

## Author Contributions

Conceptualization: A.A.B., M.G.

Methodology: A.A.B., M.G., R.T.S

Software: A.A.C., M.G.

Formal Analysis: M.G.

Investigation: M.G.

Resources: LBCC

Data Curation: D.Z, J.S., L.D., R.A.I.B

Writing - Original Draft: M.G.

Writing - Review & Editing: A.A.B., A.A.C., A.R., D.Z., J.S., L.D., L.P., R.A.I.B, R.T.S., S.S.

Visualization: M.G.

Supervision: A.A.B.

Funding Acquisition: A.A.B., M.G.

## Competing Interests

M.G., D.Z., J.S., and A.A.B. have an inventorship interest in intellectual property licensed to Centile Bioscience by the Children’s Hospital of Philadelphia. J.S., R.A.I.B., and A.A.B. hold shares in and J.S. is director of Centile Bioscience. R.T.S. has received consulting income from Octave Bioscience, Sanofi, and compensation for scientific reviewing from the American Medical Association.

## Data, code, and materials availability

Data used in the preparation of this article were obtained from the Lifespan Brain Chart consortium (LBCC), which includes multiple publicly-available MRI datasets. These datasets are subject to their respective data access and sharing policies and can be accessed via the following websites: ABCD (https://abcdstudy.org/); the International Neuroimaging Data-Sharing Initiative (https://fcon_1000.projects.nitrc.org/); the Laboratory of NeuroImaging (https://loni.usc.edu/); the BarcelonaBeta Brain Research Center (https://www.barcelonabeta.org/en); the Alzheimer’s Disease Repository Without Borders (ARWiBo) (https://www.arwibo.it); CamCAN (http://www.mrc-cbu.cam.ac.uk/datasets/camcan/); the Open Science Framework (https://osf.io/); The Global Alzheimer’s Association Interactive Network (https://www.gaain.org/); Brain-CODE (https://www.braincode.ca/); Science Data Bank (https://www.scidb.cn/en); Longitudinal Online Research and Imaging System (https://www.loris.ca/); the Human Connectome Project (https://www.humanconnectome.org/); Information eXtraction from Images (IXI) (https://brain-development.org/ixi-dataset/); figshare (https://figshare.com/); Collaborative Informatics and Neuroimaging Suite (COINS) (https://coins.trendscenter.org/); National Alzheimer’s Coordinating Center (NACC) (https://naccdata.org/); NeuroImaging Tools and Resources Collaboratory (NITRC) (https://www.nitrc.org/); Neuroscience in Psychiatry Network (NSPN) (https://nspn.org.uk/); Open Access Series of Imaging Studies (OASIS) (https://sites.wustl.edu/oasisbrains/); OpenNeuro (https://openneuro.org/); BICR Resource (https://bicr-resource.atr.jp/); UK Biobank (UKB), a major biomedical database (https://www.ukbiobank.ac.uk/); YOUth Cohort Study (https://www.uu.nl/en/research/youth-cohort-study/data-access); and the OpenPain project (https://www.openpain.org). Data accessed via NIMH Data Archive (https://nda.nih.gov/) are available at doi:.10.15154/3er3-dc69. Details of each dataset contained within the LBCC, including references, are available in **Supplemental Demographics Table**, attached. For dataset-specific acknowledgements, please see Extended Acknowledgements in the Supplement. All code is available at https://github.com/BGDlab/sex_mod_braincharts (*51*), and versioned packages are stored in https://hub.docker.com/r/mgardner457/r_gamlss.

## Notes

### Summary of Updates

Adding Supplemental files; add author Andrew A. Chen

