## Supplemental Material for "Brain charts for neuroanatomical sex differences across the human lifespan"

### Supplementary Materials

**The PDF file includes:**


Materials and Methods

- Figs S1 to S4
- Table S1 to S5

Supplementary Text

A. Sex Effect Replication Results
  a. Figs S5 to S17
B. Additional Case-Control Extended Results
  a. Figs S18 to S32
C. Sensitivity Analysis Results
  - C1. Weighting by scan quality
    a. Figs S33 to S57
  - C2. Removing infant and neonatal data
    a. Figs S58 to S76
D. Consortium Information
E. Extended Acknowledgements
F. Captions for Other Supplementary Materials


**Other Supplementary Materials for this manuscript include the following:**
Supplemental Demographics Table
Supplemental Data S1-3

Extended figures, including centile fan plots of training and testing models and sex difference trajectories for each phenotype, are available at https://drive.google.com/drive/folders/1xRb6deCtG0RPyG3JHpAm-wOaxu4a1pM9?usp=sharing.

### Materials and Methods

**Sample**

The Lifespan Brain Chart Consortium (LBCC) is a collection of structural MRI scans representing the range of the human lifespan, aggregated from primary studies around the globe. Details of the dataset and primary studies, including processing pipelines, can be found elsewhere(*7*). Briefly, scans were compiled from primary studies and segmented to Freesurfer's default atlases(*52*, *53*). For the present study, we used data from 138,261 unique individuals who participated in one of 127 primary studies. As in prior work(*7*, *54*), we also included 7,201 T1w MPRAGE scans from 5,992 unique patients at the Children's Hospital of Philadelphia; scans with no or limited imaging pathology per their clinical radiology report were curated and segmented using FreeSurfer's specialized *recon-all-clinical* pipeline(*55*), as detailed elsewhere(*56*). Global composite phenotypes were operationalized as defined in **Table S1**.

**Table S1. Operationalizations of global imaging-derived phenotypes from each processing pipeline**

| Imaging-Derived Phenotype | FreeSurfer pipeline operationalization | *Recon-all-clinical* and related *synthseg*-based pipelines operationalizations |
|---|---|---|
| Gray Matter Volume (GMV) | Total cortical gray matter volume | left cerebral cortex + right cerebral cortex |

| | | |
|---|---|---|
| White Matter Volume (WMV) | Total cortical or cerebral white matter volume | left cerebral white matter + right cerebral white matter |
| Cerebrospinal Fluid/Ventricles (CSF) | BrainSegVol - BrainSegVolNotVent | “Ventricles” or left lateral ventricle + right lateral ventricle + left inferior lateral ventricle + right inferior lateral ventricle + third ventricle + fourth ventricle |
| Subcortical Gray Matter Volume (sGMV) | Subcortical gray matter volume (thalamus, caudate nucleus, putamen, pallidum, hippocampus, amygdala, and nucleus accumbens area) | left thalamus + left caudate + left putamen + left pallidum + left hippocampus + left amygdala + left accumbens area + right thalamus + right caudate + right putamen + right pallidum + right hippocampus + right amygdala + right accumbens area |

From this initial sample of 184,508 scans, we excluded subjects with sex other than male or female (n=6 scans). For scans processed using standard FreeSurfer, we assessed image quality using the Euler index (available for 94 studies), an automated measure of the reconstructions' surface continuity often used as a robust, quantitative assessment of scan quality(*37*). As in prior work(*1*), we applied an adaptive threshold of the Euler index, excluding scans with surface hole counts greater than four median absolute deviations above the primary-study median. For scans processed using *recon-all-clinical* and related *synthseg*-based pipelines (27 studies, Supplemental Demographics Table), we applied the recommended threshold by excluding scans with any quality control index less than 0.65 across all brain structures(*57*). Together, these quality thresholds removed a total of 5,656 scans. Next, we removed scans if the value of any IDP was 0 (n=24 scans) or from subjects greater than 100 years of age (n=6 scans). In addition, we removed scans from any individual who was not a healthy control (i.e. known neuropsychiatric diagnoses, which were subsequently included in case-control analyses after benchmarking to reference models, or missing diagnosis data, n=41,044 scans). We then randomly selected a single scan to retain from subjects with more than one time point, which removed 26,262 scans. Finally, sites with fewer than 10 subjects per sex were removed, resulting in a final sample of 108,335 scans (n= 56,065 female) from 234 sites across 108 studies (Supplemental Demographics Table, attached). The age of the final sample ranged from 146 days post-conception, corresponding roughly to the second trimester of gestation, to 99.2 years post-birth.

**Ethics**

The research was reviewed by the Cambridge Psychology Research Ethics Committee (PRE.2020.104) and The Children’s Hospital of Philadelphia’s Institutional Review Board (IRB 20-017874) and deemed not to require PRE or IRB oversight as it consists of secondary analysis of de-identified primary datasets. Informed consent of participants (or their guardians) in primary studies is available in the study-specific references provided in the Supplemental Demographics Table.

**Software & Code**

Analyses were conducted using R versions 4.4.0 and 4.5.0(*58*). All code is available at https://github.com/BGDlab/sex_mod_braincharts (*51*), and versioned packages are stored in https://hub.docker.com/r/mgardner457/r_gamlss. Generative AI (ChatGPT and Claude) were used to assist code troubleshooting and optimization, as noted in the GitHub repository.

**Split-Half Significance Testing**

*Subsetting*

Following data curation, the full dataset was split into two subsamples, stratified by sex and study site (**Table S2, Fig S1**). These subsets were well-matched on age and Euler index. For each phenotype, samples were truncated prior to model fitting to ensure that nonlinear terms (age and total size) had at least 5 datapoints in the first and last 0.05% of their distributions, corresponding to at least 5 data points per penalized spline basis function. Each subset was used alternately as the training and testing sample in a split-half cross-validation approach, described below.

**Table S2. Demographics of curated LBCC Consortium data by split-half.**

| | A | B |
|---|---|---|
| N | 54177 | 54161 |
| N Studies | 108 | 108 |
| Sex (Female), n (%) | 28034 (51.7%) | 28033 (51.8%) |
| Age (years), Mean (SD) | 46.09 (25.7) | 46.09 (25.72) |
| Surface Holes, Mean (SD) | 54.2 (57.1) | 54.3 (57.7) |

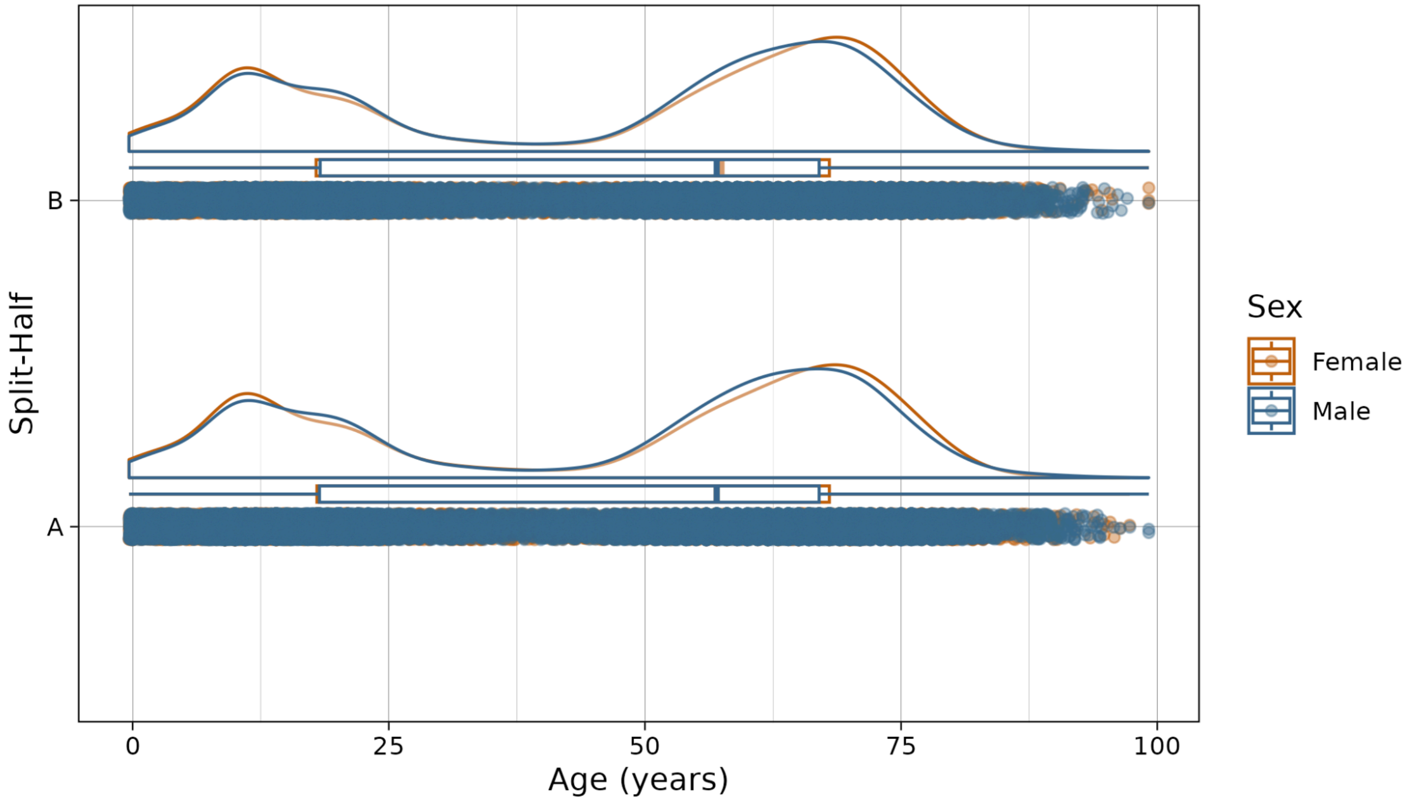


***Figure S1. Split-half samples of curated LBCC Consortium data.*** *For each split half, the figure shows a scatterplot of raw data (below), boxplot, and smoothed density plots of the distribution of male (blue) and female (orange) brain MRI scans included in the present sample.*

*Model Training & Selection*

We fit brain charts for each IDP using generalized additive models for location, scale and shape (GAMLSS)(*59*) using the *gamlss* package in R(*60*). We chose the Box-Cox t (BCTo) distribution(*48*) due to its flexibility across all four moments and log-link function in *mu*(*61*), which helped prevent convergence errors when IDP values were very small, particularly in early life. We performed model selection in the training set to determine covariate inclusion, as well as the penalties (*lambda*) of each penalized spline term. All models included, minimally, for study **i**, subject **j**, and IDP **k**:

$$f_k() = \text{BCTo}(\mu, \sigma, \nu, \tau) \text{ with}$$
$$log(\mu_{ijk}) = pb(\alpha_{\mu, age} age_{ij}) + \beta_{\mu, sex} sex_{ij} + pb(\gamma_{\mu, int} age_{ij} \times sex_{ij}) + random(\eta_{\mu, site} site_{ij}) + \zeta_{ijk}$$
$$log(\sigma_{ijk}) = pb(\alpha_{\sigma, age} age_{ij}) + \beta_{\sigma, sex} sex_{ij} + pb(\gamma_{\sigma, int} age_{ij} \times sex_{ij}) + random(\eta_{\sigma, site} site_{ij}) + \delta_{ijk}$$
$$\nu_{ijk} = 1$$
$$\tau_{ijk} = 1$$

Where *pb()* indicates penalized b-splines and *random()* indicates a random effect. The *age* term was operationalized as age in days post-conception, log-scaled to allow more wiggliness in early life as in prior work(*1*). We tested the inclusion of a FreeSurfer version term to control for variability in the versions used to process each scan in: *mu*; *mu* and *sigma*; *mu*, *sigma*, and *nu;* or no moments. We also tested the following possible formulas for *nu*:

Sex: $\nu_{ijk} = \beta_{\nu, sex} sex_{ij}$

Age (linear): $\nu_{ijk} = \alpha_{\nu, age} age_{ij}$

Site: $\nu_{ijk} = \eta_{\nu, site} site_{ij}$

Age (linear) and sex: $\nu_{ijk} = \alpha_{\nu, age} age_{ij} + \beta_{\nu, sex} sex_{ij}$

Age (linear) and site: $\nu_{ijk} = \alpha_{\nu, age} age_{ij} + \eta_{\nu, site} site_{ij}$

Sex and site: $\nu_{ijk} = \beta_{\nu, sex} sex_{ij} + \eta_{\nu, site} site_{ij}$

Age (linear), sex, and site: $\nu_{ijk} = \alpha_{\nu, age} age_{ij} + \beta_{\nu, sex} sex_{ij} + \eta_{\nu, site} site_{ij}$

Age (nonlinear): $\nu_{ijk} = pb(\alpha_{\nu, age} age_{ij})$

Age (nonlinear) and sex: $\nu_{ijk} = pb(\alpha_{\nu, age} age_{ij}) + \beta_{\nu, sex} sex_{ij}$

Age (nonlinear) and site: $\nu_{ijk} = pb(\alpha_{\nu, age} age_{ij}) + \eta_{\nu, site} site_{ij}$

Age (nonlinear), sex, and site: $\nu_{ijk} = pb(\alpha_{\nu, age} age_{ij}) + \beta_{\nu, sex} sex_{ij} + \eta_{\nu, site} site_{ij}$

*Tau* remained an intercept in all models. The penalty for each spline term, *lambda*, was determined automatically using the default GAIC method with k=log(n)(*61*). For all IDPs in each training sample, the best-fitting model was determined by the lowest BIC.

*Model Testing*

After model selection, the best-fitting model was then fit in the other split-half, which served as testing data. Test models preserved all covariates and penalties calculated in the training sample. To test the significance of age-varying sex effects (i.e. $age \times sex$) we fit a nested null model in the test sample that removed the age-by-sex moderators from both *mu* and *sigma*, then compared the full and null models using a likelihood ratio test. Effect sizes for age-varying sex effects were calculated as in prior work(*62*) using a generalized pseudo R-squared(*60*, *63*) to calculate Cohen's F-squared(*64*), which was then converted to the robust effect size index (RESI S)(*36*). For comparison, Cohen's *d=2S* for statistical models where Cohen's *d* is defined, but the RESI is generalizable to a much broader array of statistical models, including GAMLSS. Extended figures, including centile fan plots of training and testing models for each phenotype, are available at https://drive.google.com/drive/folders/1xRb6deCtG0RPyG3JHpAm-wOaxu4a1pM9?usp=sharing.

*Controlling for total brain size*

We tested whether sex effects remained significant after robustly controlling for total brain size. Given the diversity of IDPs modeled in this study, we operationalized the "total brain size" covariate as shown in Table S3 for each type of IDP:

**Table S3. "Total Brain Size" covariate corresponding to each IDP modeled**

| "Total Brain Size" Covariate | Calculation | IDPs Modeled with the Covariate |
|---|---|---|
| Total Brain Volume | Sum of cortical and subcortical gray matter, white matter, and cerebellar volumes | Volumes (global and regional), Total Surface Area, Mean Cortical Thickness |
| Mean Cortical Thickness | Mean thickness across 34 bilateral cortical regions | Regional Cortical Thickness |
| Total Surface Area | Sum of surface area across 34 bilateral cortical regions | Regional Surface Area |

To test sex effects when controlling for total brain size, we repeated the model selection procedure described above, extending the search space to include a nonlinear total size covariate in: *mu*; *mu* and *sigma*; or *mu*, *sigma*, and *nu*. We again used penalized b-splines to model the relationship between IDPs and total size, thus accounting for variable allometry across the brain. We chose not to fit models allowing sex-specific allometry (i.e., $sex \times total\ size$) based on prior literature suggesting that allometric scaling is highly conserved across sexes(*15*, *29*, *65*) and to facilitate clear interpretation of sex differences. All other model selection and testing procedures were identical to those described above, with the addition of a second null model, which removed both sex's intercept ($sex$) and age-moderated effects ($age \times sex$), to test for any effects of sex when accounting for total brain size.

*Sensitivity Analysis: Assessing the impact of scan quality*

We assessed whether image and segmentation quality impacted the significance of age-moderated sex effects – and sex's effects beyond total size – by re-fitting each IDP's training and testing models with each datapoint weighted by scan quality. Specifically, we used the large subset of scans with surface hole measures from the triangular mesh used by Freesurfer to reconstruct the cortical surface (**Table S4, Fig S2**). Notably, the minimum age at which surface hole measures were available in this sample is 1.5 years due to different processing pipelines used for younger participants.

**Table S4. Demographics of curated LBCC Consortium data with Surface Hole metric by split-half**

| | A | B |
|---|---|---|
| N | 42229 | 42185 |
| N Studies | 81 | 81 |
| Sex (Female), n (%) | 22171 (52.5%) | 22121 (52.4%) |
| Age (years), Mean (SD) | 52.73 (22.06) | 52.73 (22.07) |
| Surface Holes, Mean (SD) | 54.2 (57.1) | 54.3 (57.7) |

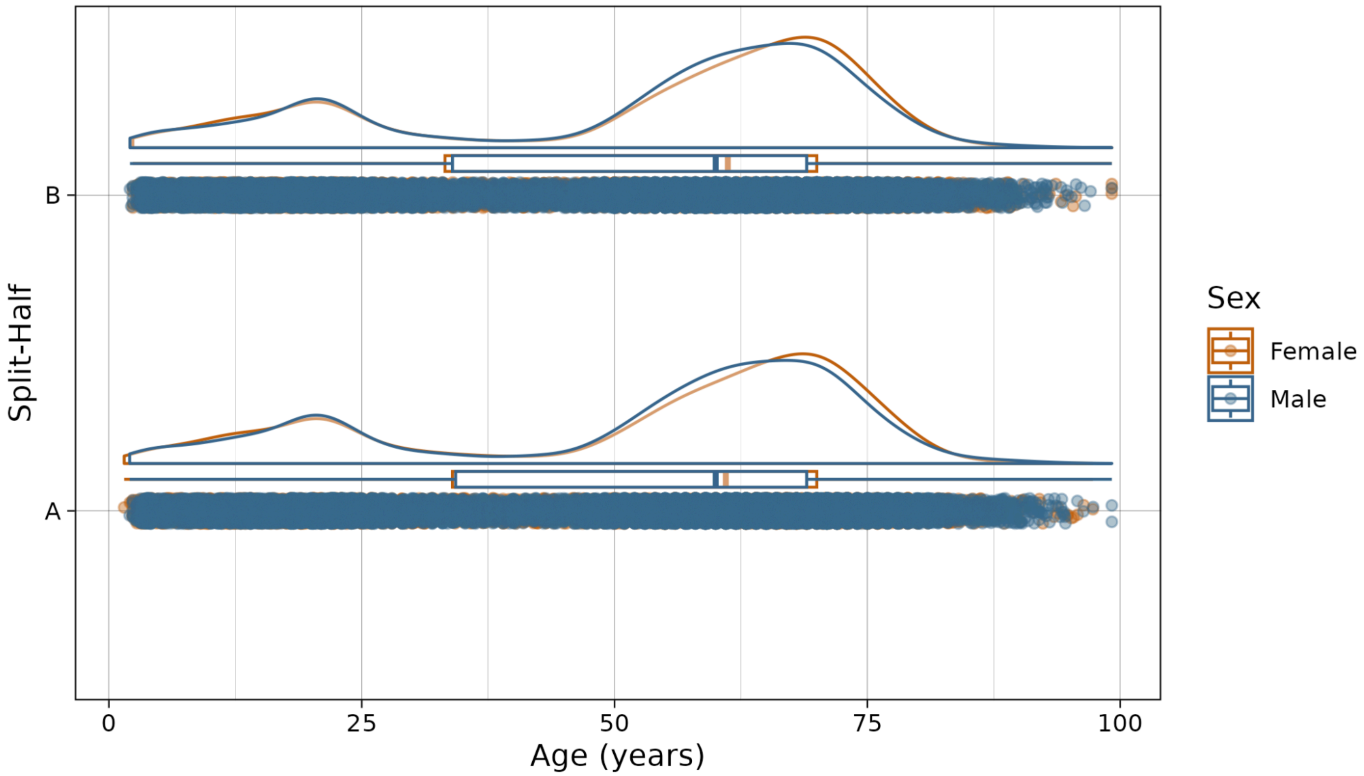


***Figure S2. Split-half samples of curated LBCC Consortium data with Surface Hole metric.*** *For each split half, the figure shows a scatterplot of raw data (below), boxplot, and smoothed density plots of the distribution of male (blue) and female (orange) brain MRI scans with accompanying data on surface holes included in the present sample.*

The number of surface holes, which is directly proportional to the Euler number of the mesh, has been validated against manual ratings as an automated measure of anatomical scan quality(*1*, *37*). To reduce the impact of poorer-quality scans while maintaining degrees of freedom and allowing study-specific Euler quantification, as recommended in previous literature(*37*), we standardized Surface Holes within each study using min-max normalization and calculated weights as follows:

$$weight = (1 - SurfaceHoles_{standardized}) \times (n/sum(1 - SurfaceHoles_{standardized}))$$

For each split-half and IDP, we refit the training model, passing these weight scores to gamlss' `weights` parameter. This weighted training model was refit in the test data, which were also weighted using this formula. Significance testing and effect size measurement were conducted as described above. The same procedure was followed to create and test sex effects in weighted models correcting for total brain size.

*Sensitivity Analysis: Removing fetal and infant cortical thickness data*

Given the difficulties in accurately quantifying cortical thickness in fetal and neonatal data, we re-ran analyses of cortical thickness excluding individuals less than 2 years old. Specifically, for each split-half and cortical thickness IDP, we refit the training and testing models in individuals aged 2 to 100 years, both with and without controlling for total size (here, mean cortical thickness).

**Quantifying Sex Biases**

To assess sex biases in IDP size and variability, we used the test models covarying for total size, which were fit as described in Table S3. All results were replicated across the two split-halves and analyses were repeated in models weighted by surface hole number (described above) to assess sensitivity to scan quality.

*Sex Difference Trajectories*

To quantify the trajectories of sex bias in each IDP across the lifespan, we applied `predict.gamlss()` to predict the *mu* and *sigma* parameters – corresponding to the 50th centile or population median and coefficient of variation (CV) trajectories, respectively(*48*, *66*) – for each sex at 500 points across the lifespan. To allow comparison across IDPs, predicted centile values were standardized relative to the male distribution of raw IDP values(*67*). Finally, at each point along the lifespan, sex biases in an IDP's population median were calculated by subtracting the scaled female 50th centile from the scaled male 50th centile, while sex biases in an IDP's variability were quantified as the log CV ratio(*26*, *68*), or the natural log of males' CV over females' CV. Thus, for an IDP at a given age, a positive sex difference in median or variability indicates males are larger or more variable than females, respectively. More specifically, if the median sex difference for an IDP is 1, for example, it indicates that males are one male-standard-deviation larger than females. Extended figures, including sex difference trajectories for each phenotype, are available at https://drive.google.com/drive/folders/1xRb6deCtG0RPyG3JHpAm-wOaxu4a1pM9?usp=sharing.

*Normalized Sex Differences*

While the likelihood ratio tests noted above quantified the overall magnitude of sex's impact on an IDP, we further quantified sex biases by taking the integral of the sex-difference trajectories described above. Specifically, because age was log-scaled in our models, we took the Riemann-Stieltjes integral of the sex difference vector over age. Since there was some variability in the age range each IDP was modeled on, we then scaled the resulting integral to reflect an age range of exactly 100 years to facilitate comparisons across IDPs. Thus, a positive normalized sex difference in median or variability indicates that, across the lifespan, the IDP tends to be overall larger or more variable in males compared to females.

In addition, we ran exploratory analyses to test for any spatial correlation between cortical IDPs' median sex bias and variability sex bias. Specifically, within cortical thickness, cortical surface area, and cortical volume IDPs, we calculated Spearman's correlation between the scaled integrals quantifying IDPs' sex differences in medians and the scaled integrals quantifying sex differences in variability. We used spatial null models to assess correlations' significance relative to 10,000 spatial-autocorrelation-preserving random samples(*69*–*71*).

*Grouping Sex Difference Trajectories*

To describe how sex biases change over the lifespan, we grouped IDPs by their sex difference trajectories, once by sex differences in the median and once by sex differences in the variability. First, any IDPs that did not have a significant age-moderated sex term when controlling for total brain size (see above) were categorized as "Stable". Next, we calculated the proportion of the lifespan modeled that an IDP's difference trajectory was positive (i.e., male-biased) or negative (i.e., female-biased) and assigned IDPs that were positive or negative for over 95% of the lifespan to either the "Male" or "Female", respectively. IDPs in these groups were further sub-categorized as "Diverging" or "Converging" based on whether their sex biases tended to increase or decrease across the lifespan. We assessed this by first squaring the sex difference trajectories such that a positive slope indicated an increase in sex difference magnitude, then taking the integral of the derivative of this trajectory, with a positive integral indicating males' and females' trajectories tend to diverge with age. Finally, the minority of IDPs that were not biased towards one sex for more than 95% of the lifespan were categorized as: "Male to Female Bias" if sex difference trajectories were positive (i.e. male biased) at the beginning of life and negative (i.e. female biased) at the end of life; "Female to Male Bias" if the sex differences were negative at the beginning of life and positive at the end of life; or "Complex" for more complex trajectories that did not fit these patterns. For visualization purposes, we summarized each group's trajectories by fitting a generalized additive model with a nonlinear age term to the standardized sex difference estimates of all IDPs in the group at each point along the lifespan.

*Quantifying the Age of Neuroanatomical Peaks*

We identified the age at which global IDPs reached several points of interest by extracting the ages corresponding to each of the following values. First, for each IDP, we averaged the 50th centile trajectories of each sex and sex-bias trajectory (i.e., male minus female 50th centile) across each split-half's test model. Next, for each sex, we defined the maximum value of the 50th centile trajectory ("Max Female" and "Max Male") and its derivative ("Max Change Female" and "Max Change Male"). The remaining points were defined using the trajectory of median sex bias itself, namely its maximum absolute value ("Max Sex Bias"), minimum absolute value ("Min Sex Bias"), and maximum absolute value of its derivative ("Max Change Sex Bias").

*Sensitivity Analysis: Data-driven trajectory clustering*

Given that the descriptive groupings outlined above were defined *a priori*, we also assessed a data-driven *k*-medoids clustering method to identify relevant groupings of regional sex differences. *k*-medoids facilitates interpretability by identifying one IDP's trajectory as the medoid, or center, of each cluster. Clustering was done twice, once on median sex bias and once on sex bias in variability, per split-half. IDPs were classified by the Euclidean distance between their sex bias trajectories, calculated via the `dist()` function, using the partitioning around the medoids algorithm implemented in the *cluster* package's `pam()`. The number of clusters was chosen to maximize average silhouette width, which indexes how close a datapoint is to members of its own cluster relative to other clusters, with the goal to minimize distance within clusters and maximize it between clusters(*72*).

**Case-control analyses**

*Clinical Samples*

The patient sample was compiled from the LBCC dataset. After filtering for quality control and randomly selecting one scan per subject, as described above, we used the diagnoses shared by each primary study in the dataset to identify individuals with primary diagnoses of: schizophrenia, Alzheimer's disease, autism spectrum disorder, major depressive disorder, generalized anxiety disorder, or attention-deficit/hyperactivity disorder. For all case-control analyses, clinical subjects were assessed relative to controls from the same study sites(**Table S5, Fig S3**). Given the small number of cases relative to controls, we did not include individuals from the UK Biobank sample with autism spectrum disorder or schizophrenia in these analyses.

**Table S5. Demographics of psychiatric samples and site-matched controls.**

| | Case | Control |
|---|---|---|
| **ADHD** | | |
| N | 1193 | 11090 |
| No. Studies | 6 | 6 |
| Female, n (%) | 334 (28%) | 5405 (48.7%) |
| Age (years), Mean (SD) | 11.74 (4.93) | 10.95 (2.99) |
| Surface Holes, Mean (SD) | 103.1 (74.3) | 100.3 (69.4) |
| **ALZ** | | |
| N | 2067 | 6020 |
| No. Studies | 9 | 9 |
| Female, n (%) | 1138 (55.1%) | 3884 (64.5%) |
| Age (years), Mean (SD) | 74.31 (8.9) | 65.93 (15.67) |
| Surface Holes, Mean (SD) | 126.2 (85.6) | 115.6 (86.6) |
| **ASD** | | |
| N | 1293 | 2087 |
| No. Studies | 10 | 10 |
| Female, n (%) | 295 (22.8%) | 937 (44.9%) |
| Age (years), Mean (SD) | 12.17 (8.92) | 13.12 (11.97) |
| Surface Holes, Mean (SD) | 126.6 (86.8) | 98.1 (71.3) |
| **GAD** | | |
| N | 2559 | 63419 |
| No. Studies | 4 | 4 |
| Female, n (%) | 1589 (62.1%) | 31448 (49.6%) |
| Age (years), Mean (SD) | 61 (15.57) | 56.21 (21.44) |
| Surface Holes, Mean (SD) | 32.7 (18.6) | 34 (20.1) |
| **MDD** | | |
| N | 7565 | 68419 |
| No. Studies | 14 | 14 |
| Female, n (%) | 4930 (65.2%) | 34484 (50.4%) |
| Age (years), Mean (SD) | 60.97 (12.27) | 57.53 (20.06) |
| Surface Holes, Mean (SD) | 34.1 (24.4) | 41 (35.9) |
| **SCZ** | | |
| N | 1027 | 1714 |
| No. Studies | 16 | 16 |
| Female, n (%) | 374 (36.4%) | 799 (46.6%) |
| Age (years), Mean (SD) | 32.63 (11.57) | 29.88 (12.19) |
| Surface Holes, Mean (SD) | 78.7 (48.5) | 71.5 (47.4) |

*Abbrv: ADHD, attention-deficit/hyperactivity disorder; ALZ, Alzheimer's disease; ASD, autism spectrum disorder; GAD, generalized anxiety disorder; MDD, major depressive disorder; SCZ, schizophrenia.*

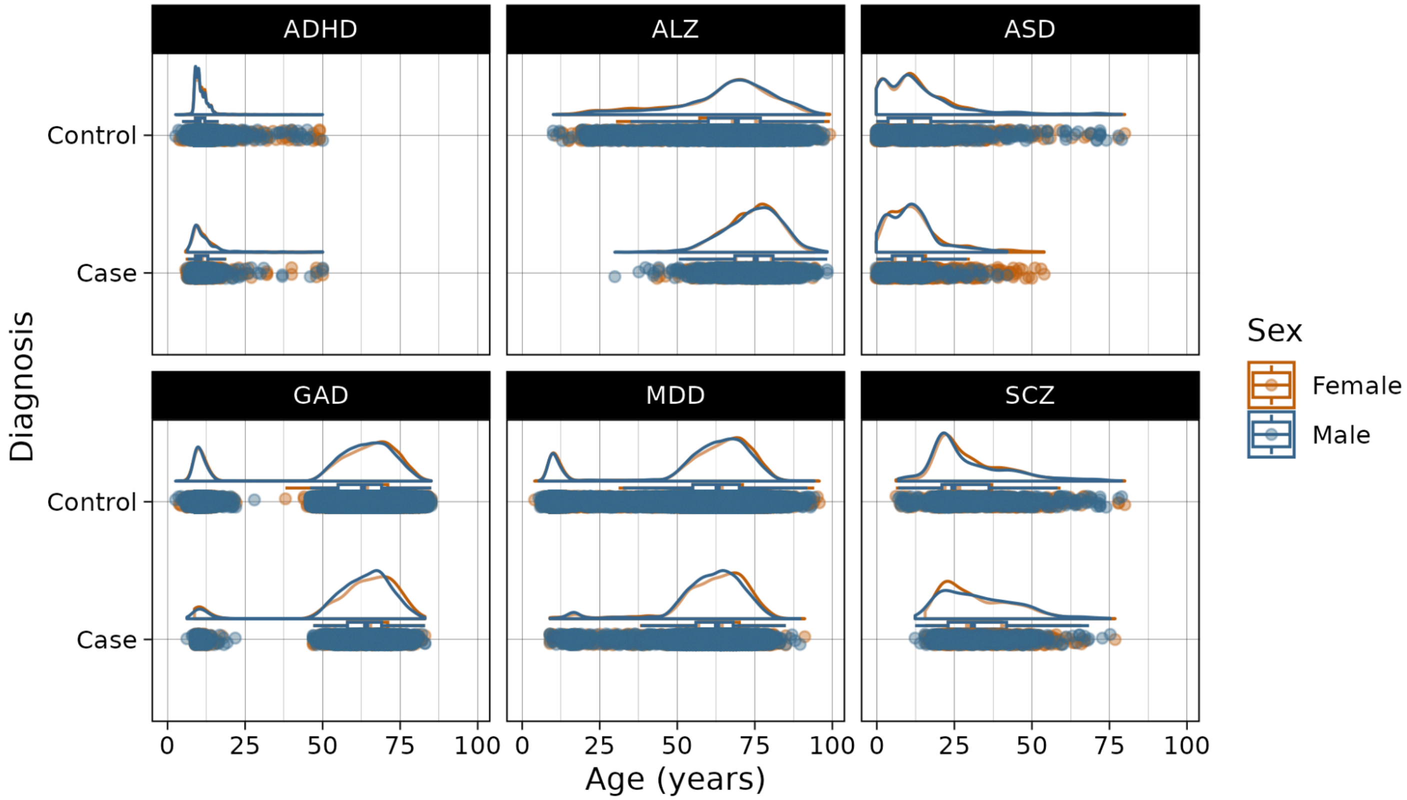


***Figure S3. Patient data in each diagnosis alongside site-matched controls.*** *Abbrv: ADHD, attention-deficit/hyperactivity disorder; ALZ, Alzheimer's disease; ASD, autism spectrum disorder; GAD, generalized anxiety disorder; MDD, major depressive disorder; SCZ, schizophrenia.*

*Reference Score Derivation*

For our primary analyses, we obtained reference scores for case and control subjects from models that did not control for total brain size, for interpretability of case-control differences. For each subject, we derived centile scores from test models using *gamlss*' `predictAll()` function. Centile z-scores were derived from centiles using R's `qnorm()` function; they are identical to traditional z-scores when the underlying distribution is normal. To prevent infinite centile z-scores, centiles of 0 and 1 were estimated as *1e-25* and 1–*1e-25,* respectively. For controls, all reference scores were calculated for the held-out sample only, i.e., from the test model that was trained on the other split half. For patients, centile z-scores were calculated on both test models for each IDP (one from each split half) and averaged, back-transforming to obtain mean centile scores as needed. For IDPs where the test models only converged in one split-half, centiles from the converged model were used without averaging.

To test how accounting for age-varying sex effects impacted reference scores, scores for each subject were derived twice for each IDP, once from the full sex-moderated model and once from the sex-intercept-only null model. These analyses were also repeated using reference scores derived from total-size-corrected models. Notably, reference scores of cortical thickness IDPs controlling for total brain size (i.e. regional thickness scores controlling for mean cortical thickness) were not well-correlated across our two split-half testing models (**Fig S4**). Thus, in our supplemental analyses of regional case-control differences in total-size-corrected reference scores, we used scores derived from our sensitivity models of cortical thickness IDPs fit on individuals ages 2 years or older (described above).

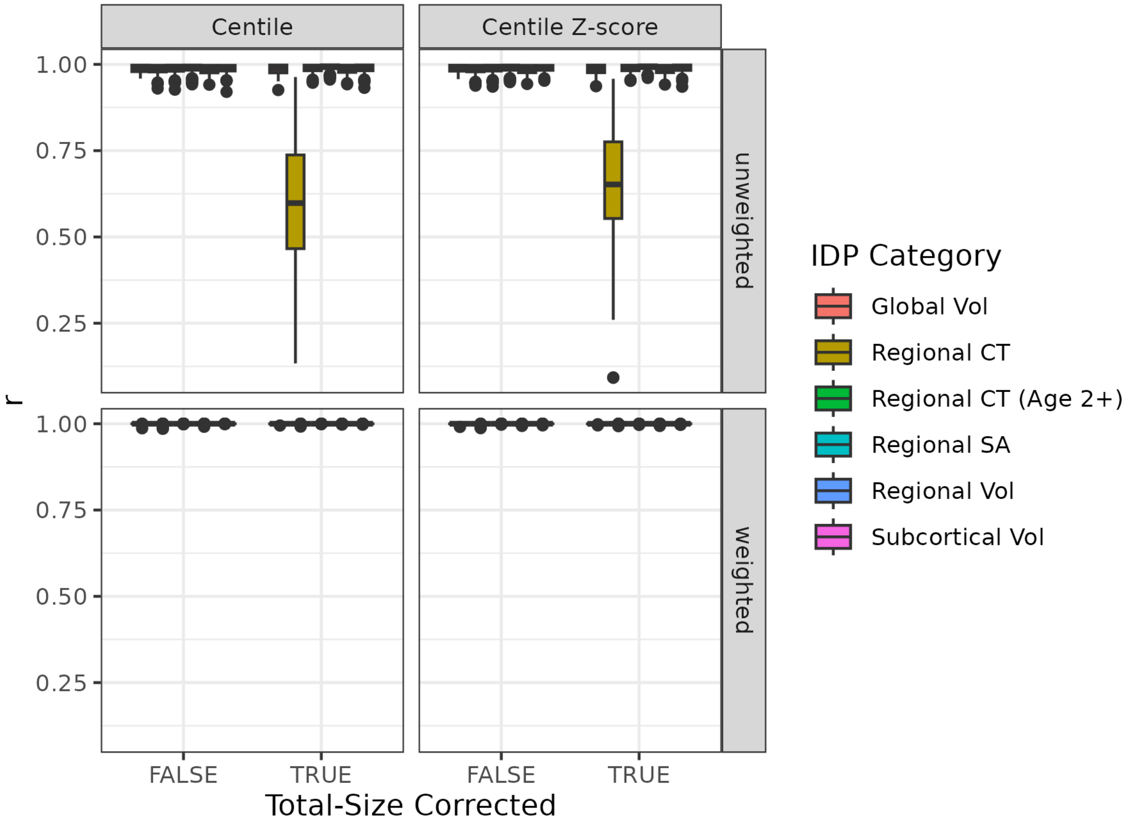


***Figure S4. Correlations between patients' reference scores derived from each split-half model show high reliability in most models and IDP types.*** *Centile (left panels) and centile z-scores (right panels) of held-out patients were derived from a series of models fit in each split-half of the data: correcting or not-correcting for total size (x-axis), and weighting (lower panels) or not weighting (upper panels) for scan quality. While most models and IDPs produce consistent scores for each individual (r close to 1) across each split-half, scores of total-size-corrected thickness IDPs (yellow boxes) were not well correlated. This is corrected by refitting models only with data from individuals aged two years or older (green boxes), which were used for all subsequent case-control analyses. Abbrv: CT, cortical thickness; SA, surface area; Vol, volume.*

*Analysis of mean case-control differences in reference score*

To assess brain charts' ability to uncover clinically meaningful deviation in IDP morphometry, we tested whether the average centile z-score of individuals with a given diagnosis differed from that of controls. Specifically, in each diagnosis and IDP, we used two-tailed *t*-tests with Welch's correction to compare z-scores in cases and site-matched controls, with FDR correction for testing multiple IDPs and diagnoses. Tests were conducted separately in centiles derived from the sex-moderated and sex-intercept-only models (see above).

*Extremeness analyses*

We also assessed whether clinical cases were over- or under-represented among those with extreme reference scores. As in prior literature(*62*), we defined extreme high centiles as those greater than 95% (centile z-score > 1.64) and extreme low centiles as those less than 5% (centile z-score < -1.64). For each diagnosis and IDP, we used Fisher's exact test of proportions (EnvStats' `twoSamplePermutationTestProportion` function) to compare the proportion of cases with extremely low or high centiles to the proportion of controls in the same group, using FDR correction to account for testing multiple diagnoses and IDPs. As above, we tested separately for differences in z-scores derived from the full and sex-null models.

### Supplementary Text

#### A. Sex Effect Replication Results

Here we show the results of our analyses probing the significance of age-varying and overall sex effects across both split-half replications. For the complete results underlying these figures, please see Supplemental Data S1.

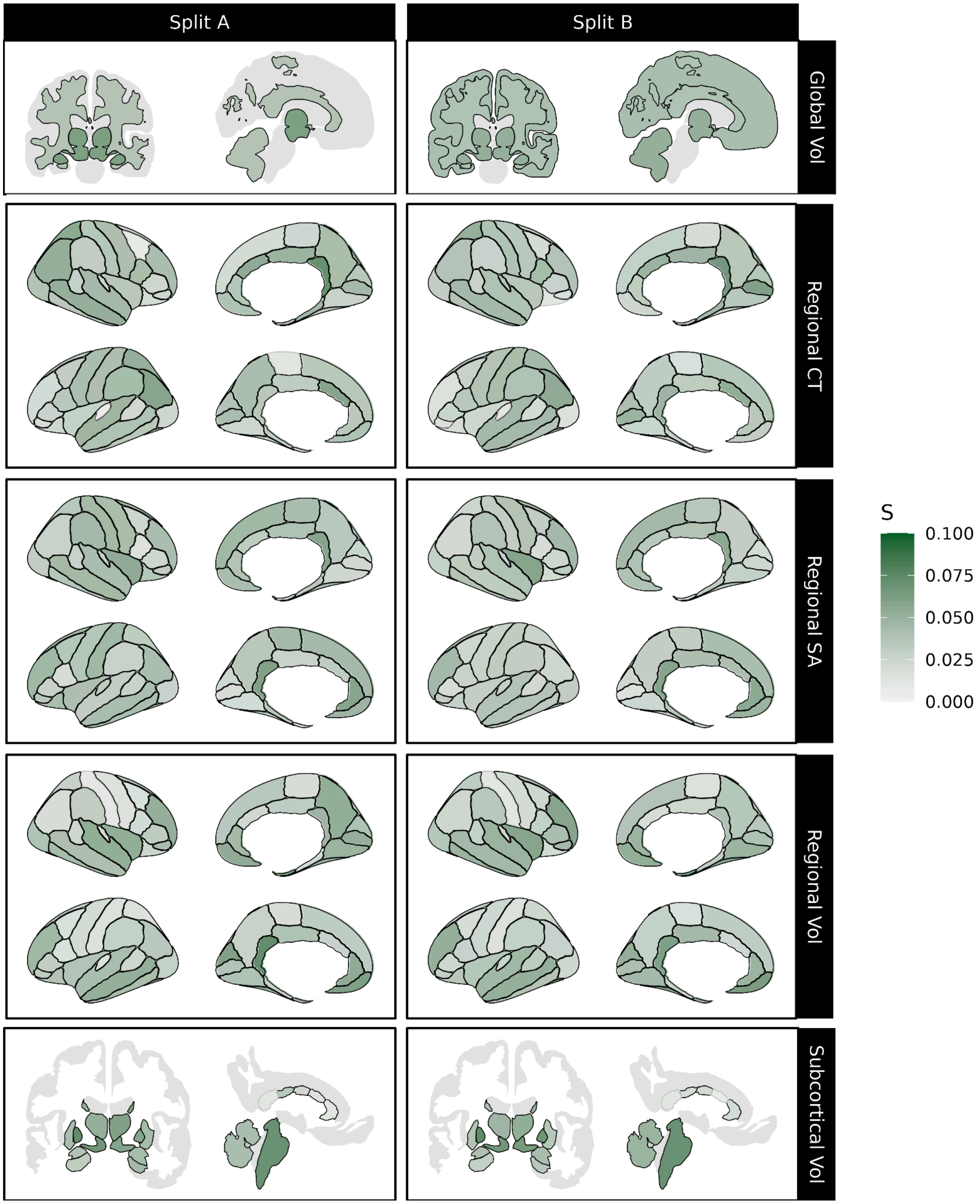


***Figure S5. Nearly all neuroanatomical features exhibit sex biases that vary across the lifespan.*** *Magnitude of the age-varying component of sex (age-by-sex term) on IDPs each test split. Effect sizes (S) are reported using the robust effect size index (RESI). Black outlines indicate significant effects at p <0.05, using false discovery rate (FDR) correction for multiple comparisons. Abbrv: CT, cortical thickness; SA, surface area; Vol, volume.*

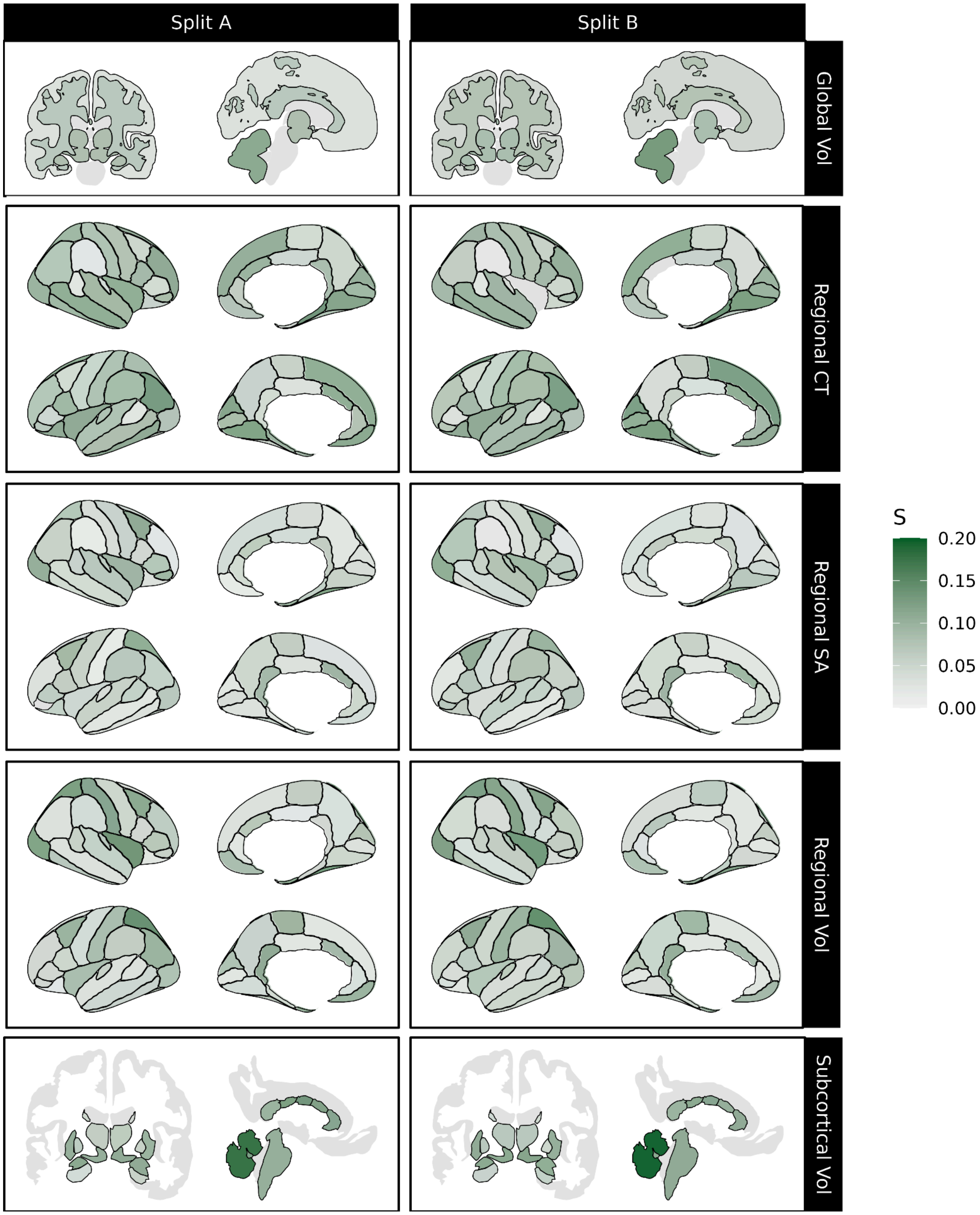


***Figure S6. Sex effects survive when controlling for total brain size.*** *Magnitude of sex effects (sex intercept and age-by-sex interaction) on IDPs each test split when controlling for total brain size. Effect sizes (S) are reported using the robust effect size index (RESI). Black outlines indicate significant effects at p <0.05, using false discovery rate (FDR) correction for multiple comparisons. Abbrv: CT, cortical thickness; SA, surface area; Vol, volume.*

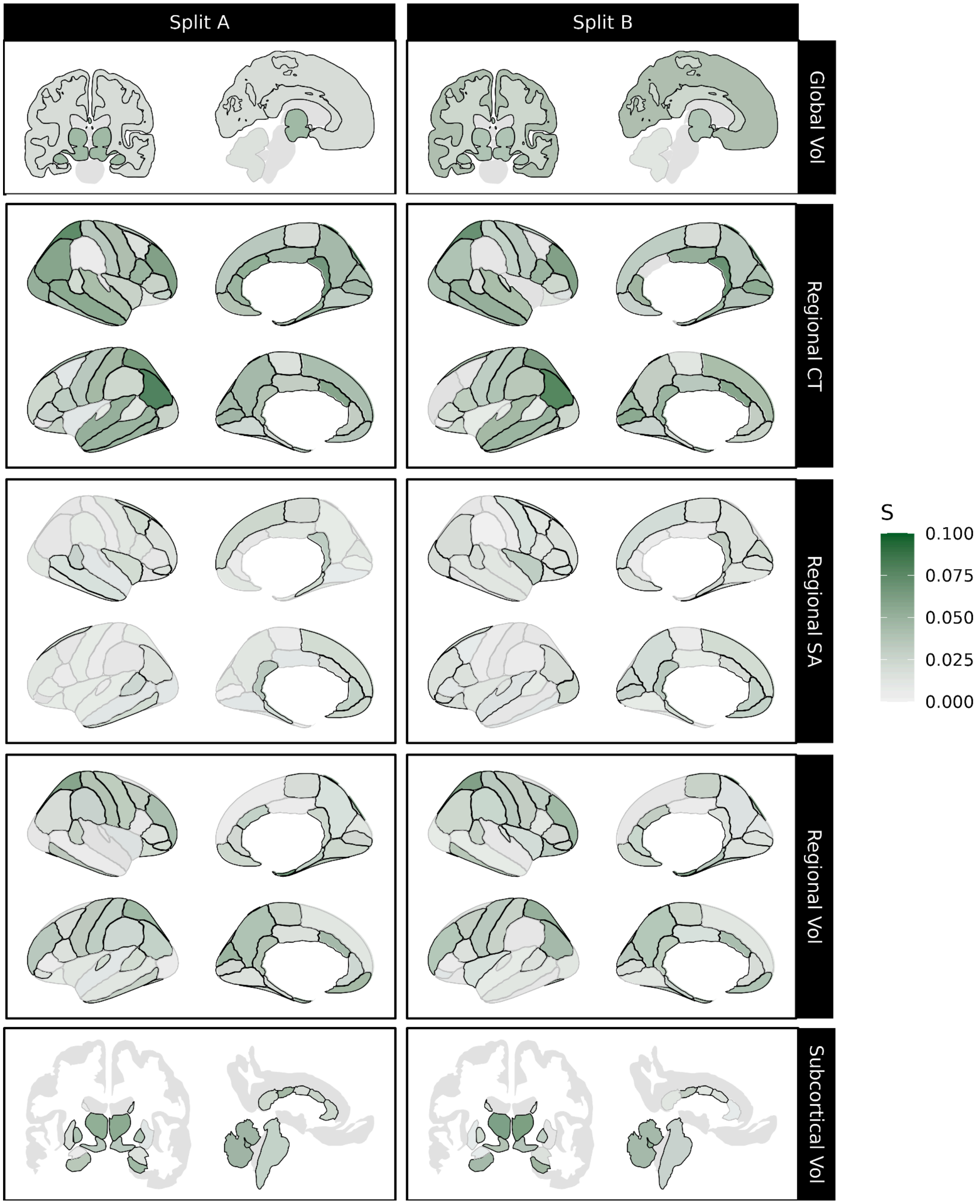


***Figure S7. Age-varying sex effect when controlling for total brain size.*** *Magnitude of age-by-sex interaction effect on IDPs each test split when controlling for total brain size. Effect sizes (S) are reported using the robust effect size index (RESI). Black outlines indicate significant effects at p <0.05, using false discovery rate (FDR) correction for multiple comparisons. Abbrv: CT, cortical thickness; SA, surface area; Vol, volume.*

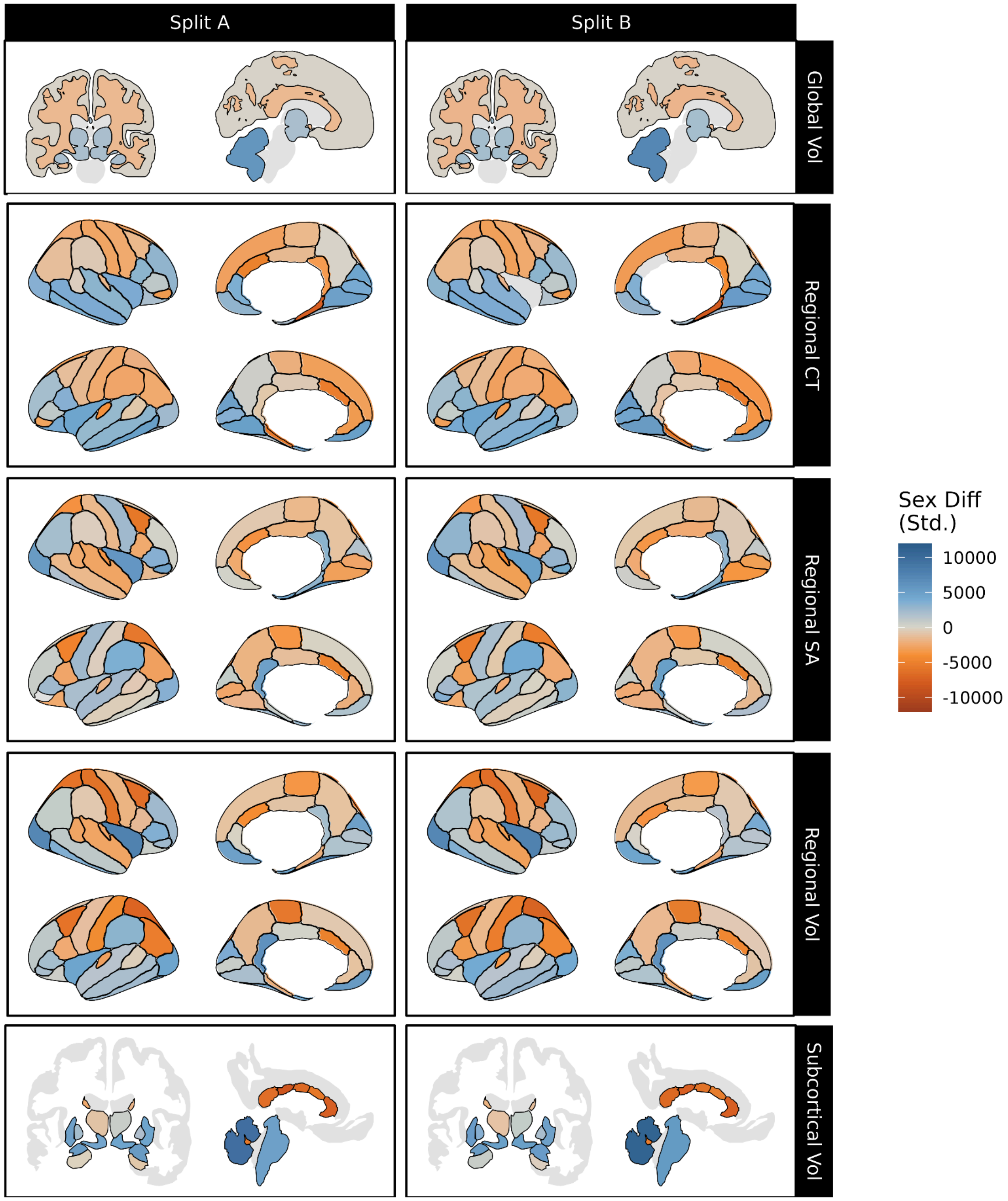


***Figure S8. Sex differences in IDPs' medians over the lifespan.*** *Sex bias in IDP median trajectory controlling for total size, with orange indicating female bias and blue indicating male bias. Predicted centiles for each sex were standardized by scaling relative to each IDP's standard deviation before subtracting predicted female median centile from the predicted male median centile at each age; overall sex differences were operationalized as the integral of the resulting sex difference trajectory, normalized to a 100-year lifespan (see Methods). Black outlines indicate significant sex effects at p <0.05, using false discovery rate (FDR) correction for multiple comparisons. Abbrv: CT, cortical thickness; SA, surface area; Vol, volume.*

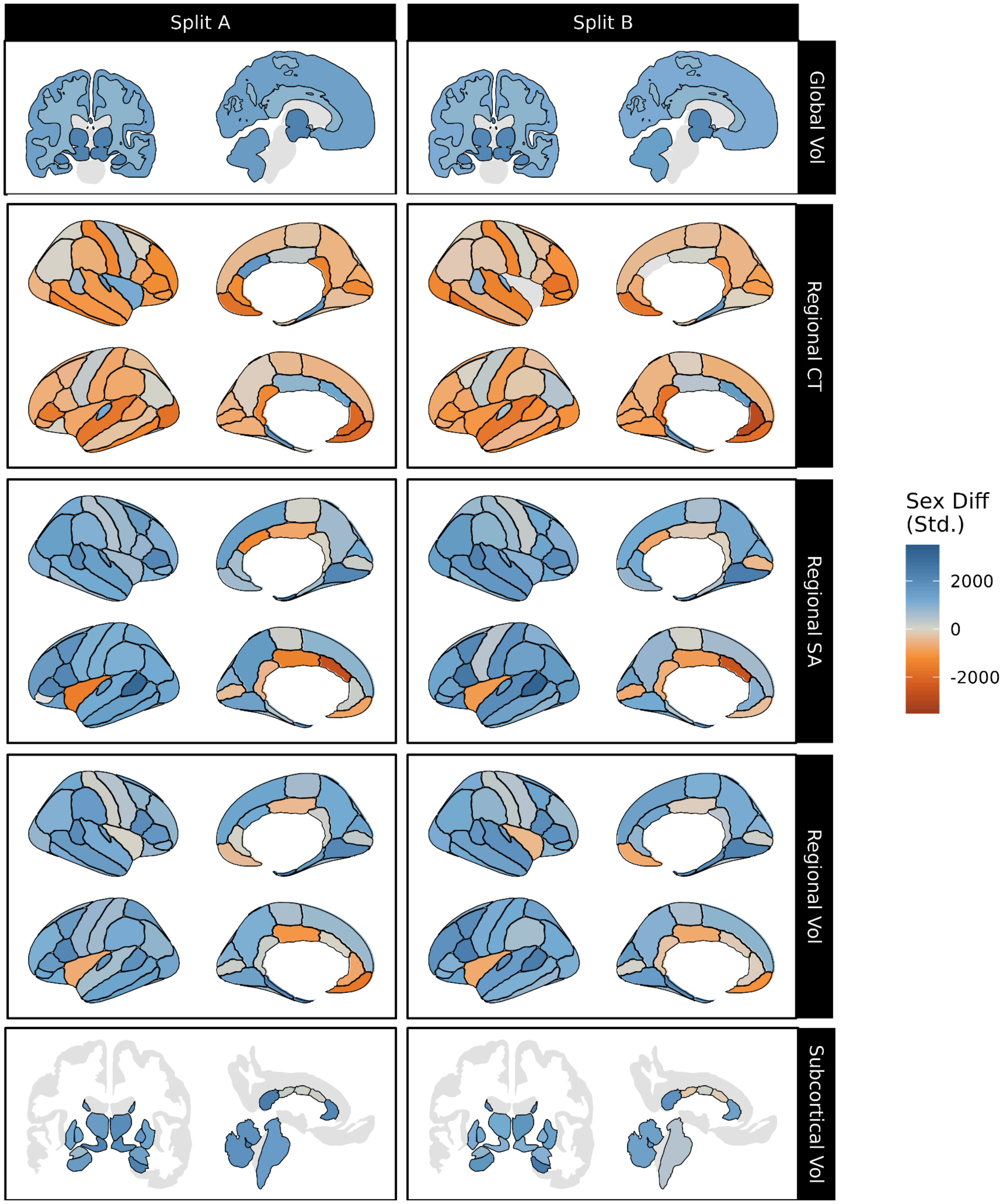


***Figure S9. Standardized sex differences in IDPs' variabilities over the lifespan.*** *Sex bias in IDP variability controlling for total size, with orange indicating female bias and blue indicating male bias. Predicted centiles for each sex were standardized by scaling relative to each IDP's standard deviation before calculating the natural log of the ratio of males' over females' coefficient of variation (i.e. log(male sigma/female sigma)); overall sex differences were operationalized as the integral of the resulting sex difference trajectory, normalized to a 100-year lifespan (see Methods). Black outlines indicate significant sex effects at p <0.05, using false discovery rate (FDR) correction for multiple comparisons. Abbrv: Std., standardized; CT, cortical thickness; SA, surface area; Vol, volume.*

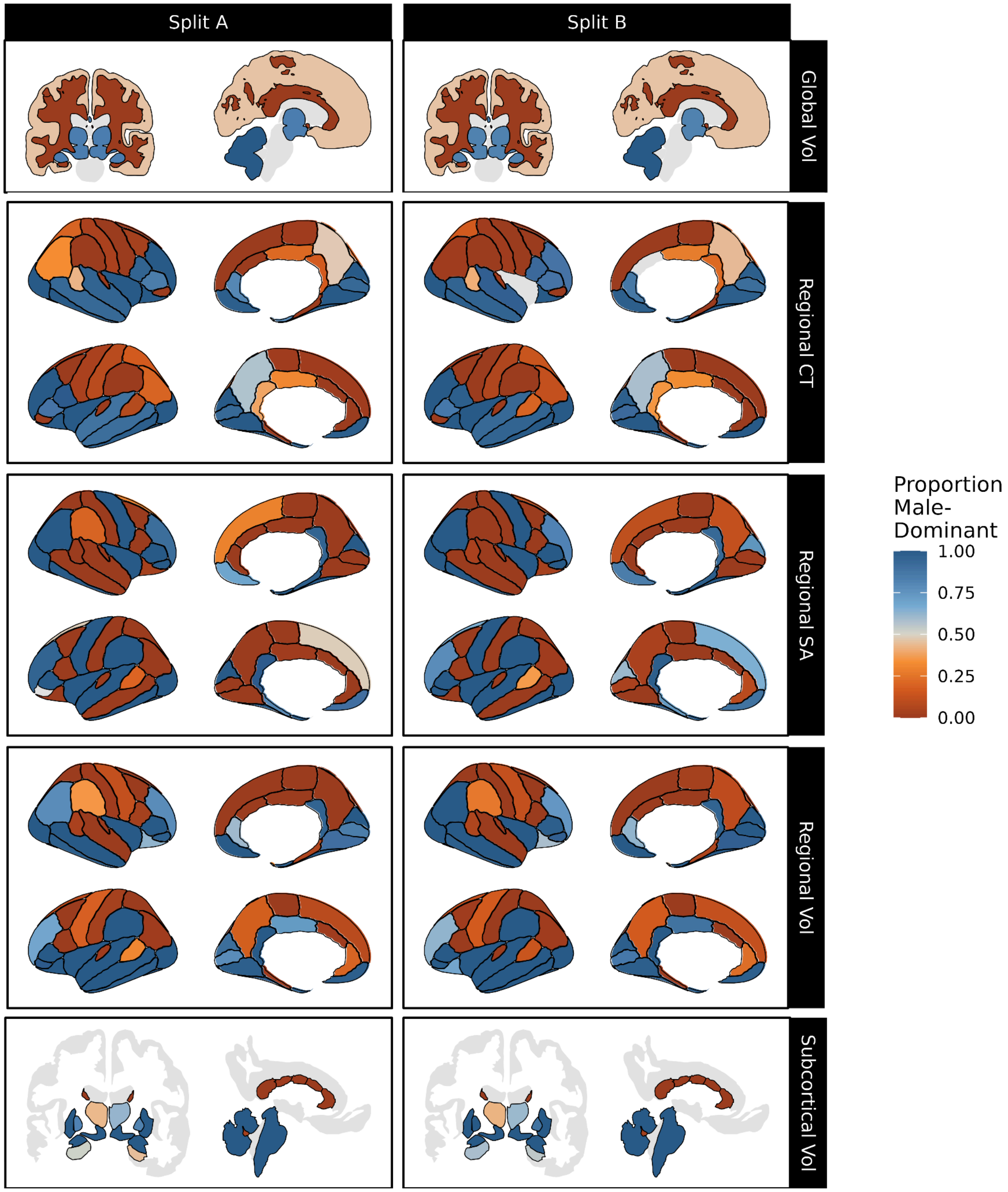


***Figure S10. Proportion of the lifespan an IDP's median is biased towards males.*** *Blue indicates more of the lifespan being male-biased (i.e. larger in males), while orange indicates a longer time spent female-biased. Black outlines indicate significant sex effects at p <0.05, using false discovery rate (FDR) correction for multiple comparisons. Abbrv: CT, cortical thickness; SA, surface area; Vol, volume.*

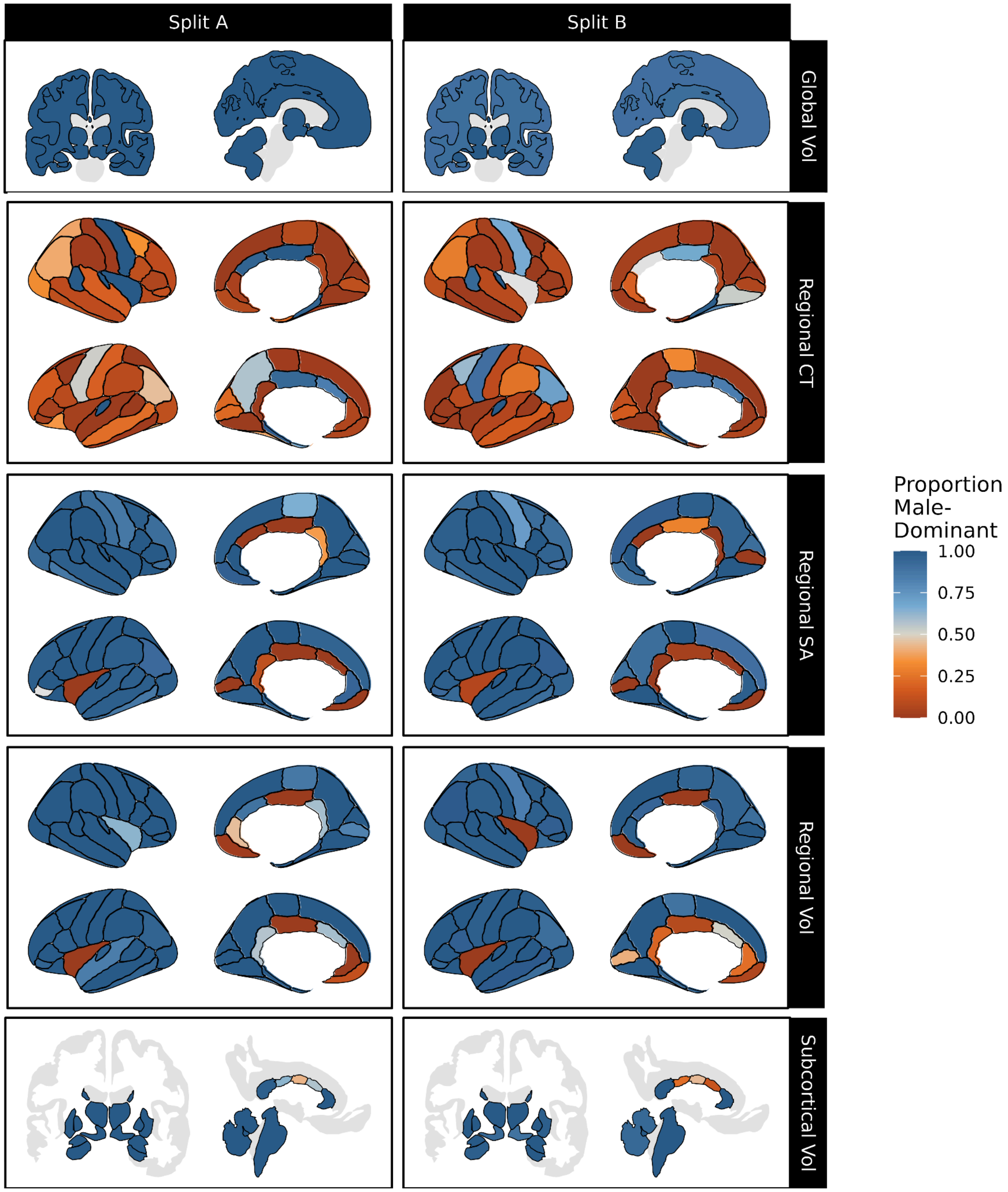


***Figure S11. Proportion of the lifespan an IDP's variability is biased towards males.*** *Blue indicates more of the lifespan being male-biased (i.e. more variable in males), while orange indicates a longer time spent female-biased. Black outlines indicate significant sex effects at p <0.05, using false discovery rate (FDR) correction for multiple comparisons. Abbrv: CT, cortical thickness; SA, surface area; Vol, volume.*

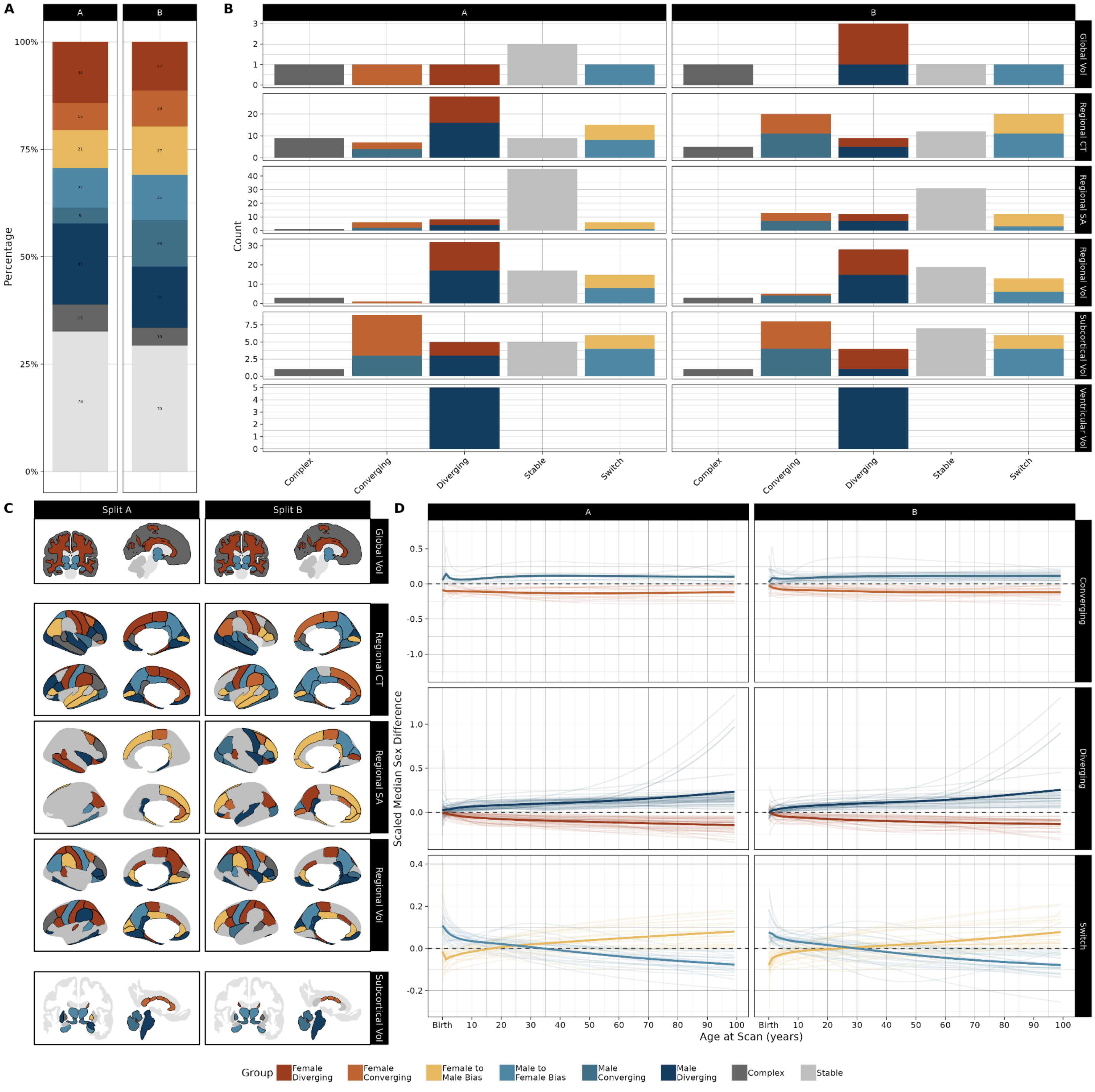


***Figure S12. IDP groupings by* a priori *classes of median sex difference trajectories. (A)*** *Percentage of IDPs belonging to each group, as determined by the trajectory of the IDPs' median sex difference over the lifespan, in each split-half. Groups are defined as follows: female diverging, where consistently female-biased features become more biased over the lifespan; female converging, where female bias decreases over time; female bias to male bias, where IDPs that are female-biased early in life become male-biased later; male bias to female bias, where bias changes from male to female with age; male converging, where male bias decreases with age; male diverging, where male bias increases with age; stable, where sex biases do not significantly vary with age (as per likelihood ratio tests); complex, for significantly age-varying sex biases that do not fit any previously defined groups.* ***(B)*** *Number of IDPs belonging to each group, subdivided by phenotype class.* ***(C)*** *IDPs colored by group.* ***(D)*** *Summarized sex bias trajectories (dark lines) within each*

*group. Summaries for each group were created by standardizing sex-difference trajectories within each IDP (translucent lines) and fitting a simple generalized additive model.*

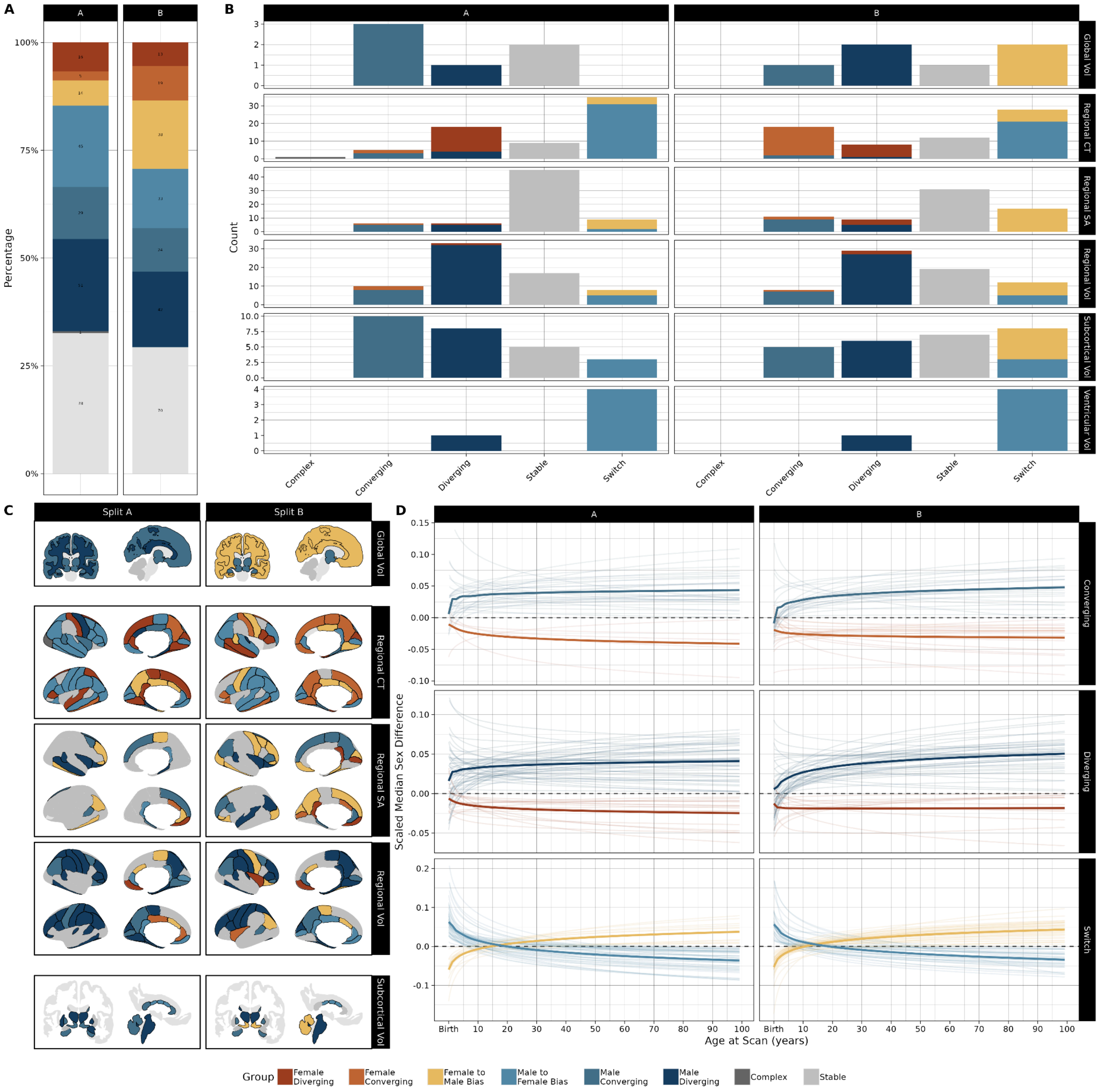


***Figure S13. IDP groupings by*** **a priori** ***classes of sex difference trajectories in variability. (A)*** *Percentage of IDPs belonging to each group, as determined by the trajectory of the IDPs' sex difference in variability over the lifespan, in each split-half. Groups are defined as follows: female diverging, where consistently female-biased features become more biased over the lifespan; female converging, where female bias decreases over time; female bias to male bias, where IDPs that are female-biased early in life become male-biased later; male bias to female bias, where bias changes from male to female with age; male converging, where male bias decreases with age; male diverging, where male bias increases with age; stable, where sex biases do not significantly vary with age (as per likelihood ratio tests); complex, for significantly*

*age-varying sex biases that do not fit any previously defined groups.* ***(B)*** *Number of IDPs belonging to each group, subdivided by phenotype class.* ***(C)*** *IDPs colored by group.* ***(D)*** *Summarized sex bias trajectories (dark lines) within each group. Summaries for each group were created by standardizing sex-difference trajectories within each IDP (translucent lines) and fitting a simple generalized additive model.*

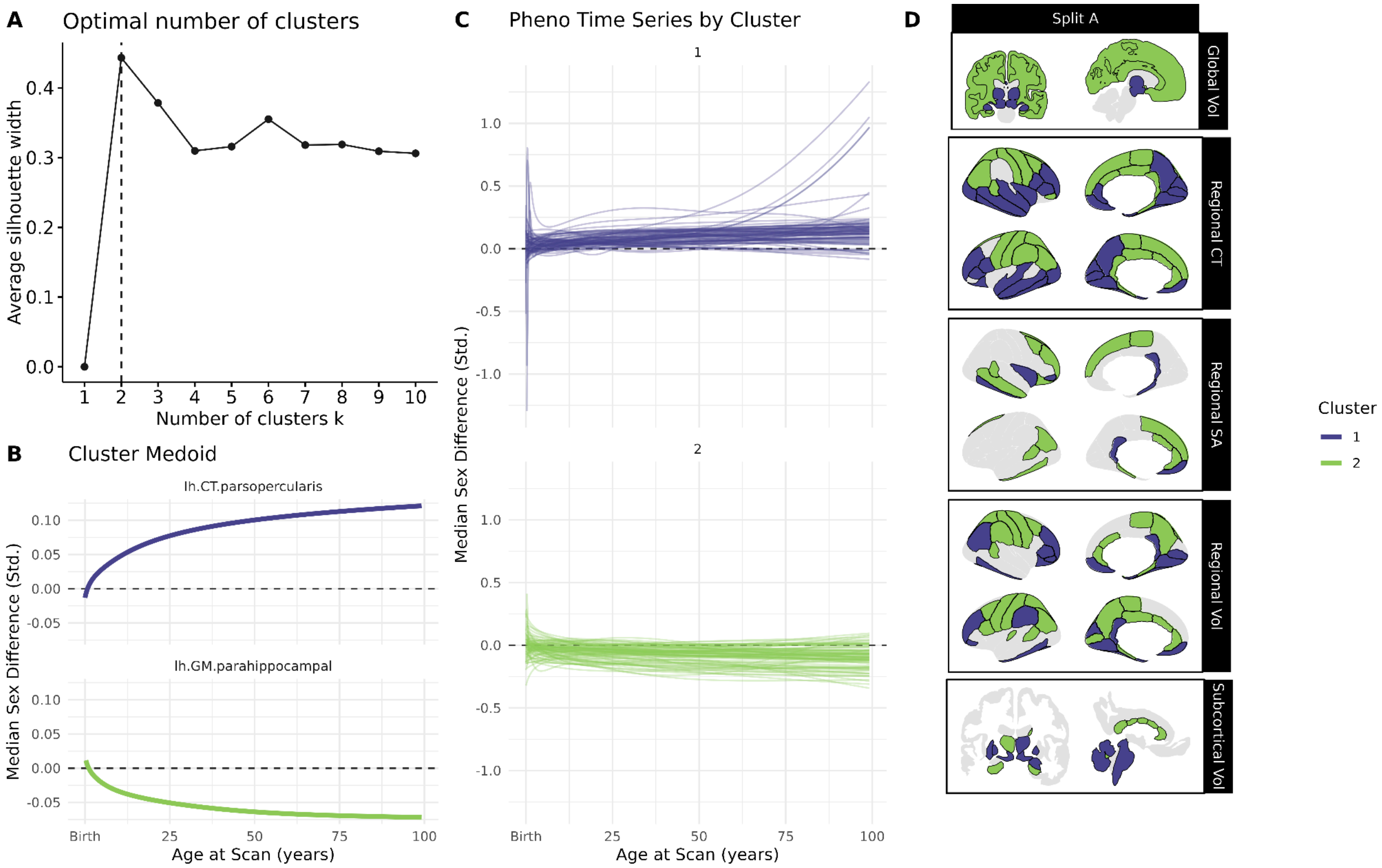


***Figure S14. Data-driven clustering of IDPs by median sex difference trajectories in test split A. (A)*** *Plot of average silhouette width by number of k-medoid clusters, indicating a two-cluster solution.* ***(B)*** *Scaled median sex bias trajectories of cluster medoids for 2-cluster solution, corresponding to increasing male bias (Cluster 1, purple) and increasing female bias (Cluster 2, green)* ***(C)*** *Scaled median sex bias trajectories of each IDP, grouped by cluster. (D) IDPs visualized in neuroanatomical context, colored by sex bias trajectory cluster. All IDPs included in k-medoid clustering protocol had significantly age-varying sex difference trajectories, as indicated by our FDR corrected likelihood ratio tests (black outlines). Abbrv: CT, cortical thickness; SA, surface area; Vol, volume; Std., standardized*

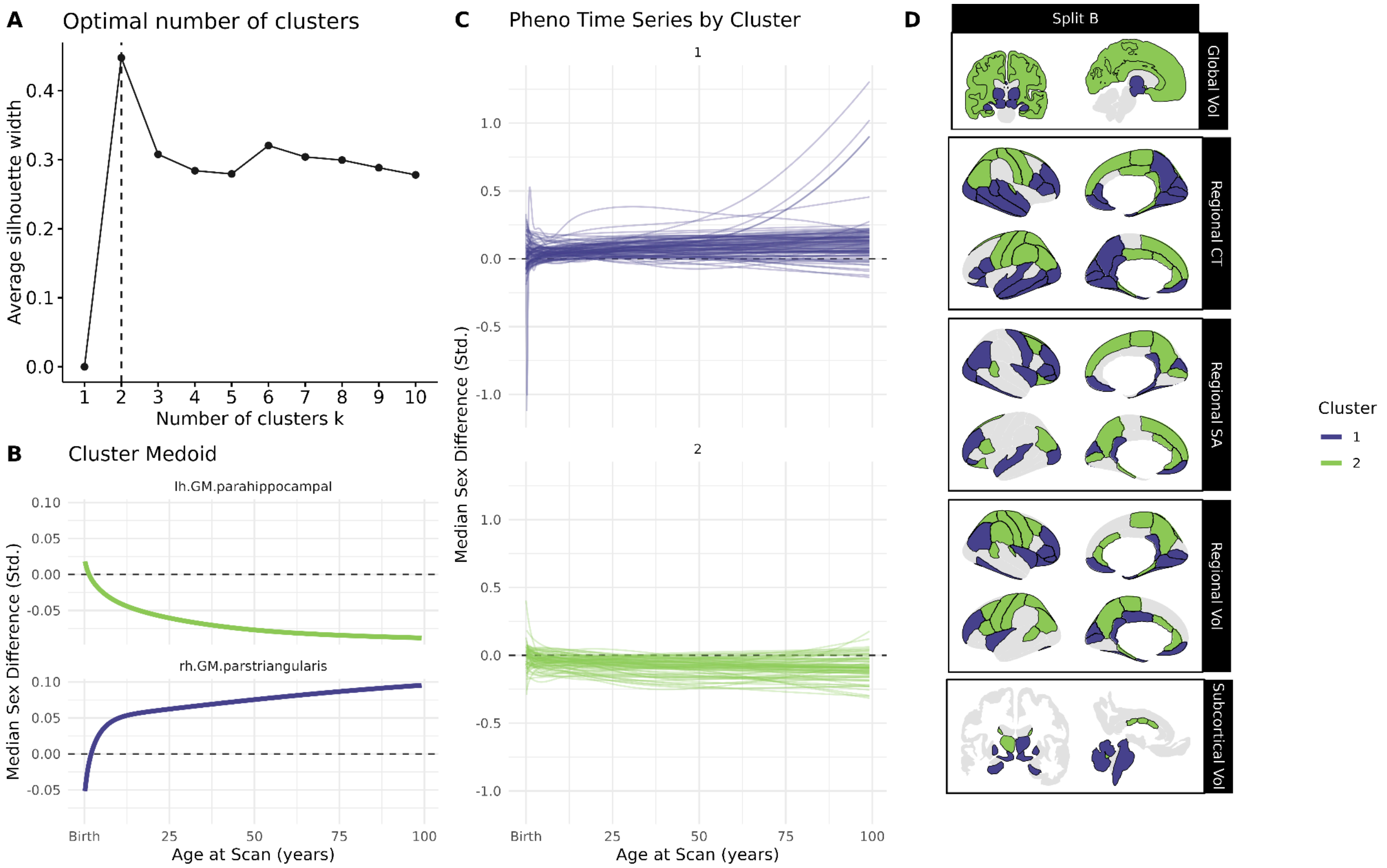


***Figure S15. Data-driven clustering of IDPs by median sex difference trajectories in test split B. (A)*** *Plot of average silhouette width by number of k-medoid clusters, indicating a two-cluster solution.* ***(B)*** *Scaled median sex bias trajectories of cluster medoids for 2-cluster solution, corresponding to increasing male bias (Cluster 1, purple) and increasing female bias (Cluster 2, green)* ***(C)*** *Scaled median sex bias trajectories of each IDP, grouped by cluster. (D) IDPs visualized in neuroanatomical context, colored by sex bias trajectory cluster. All IDPs included in k-medoid clustering protocol had significantly age-varying sex difference trajectories, as indicated by our FDR corrected likelihood ratio tests (black outlines). Abbrv: CT, cortical thickness; SA, surface area; Vol, volume; Std., standardized*

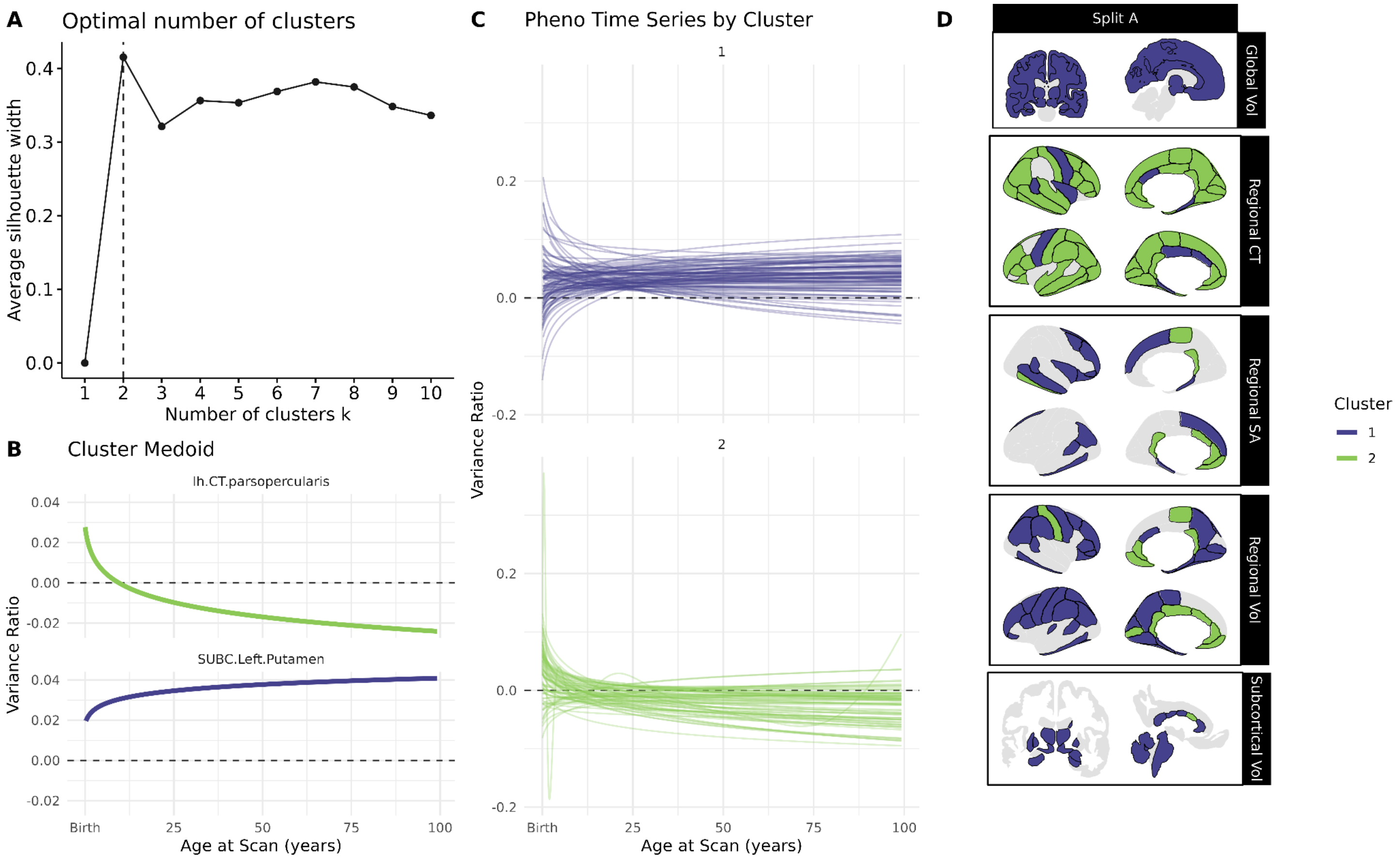


***Figure S16. Data-driven clustering of IDPs by sex difference trajectories in variability in test split A. (A)*** *Plot of average silhouette width by number of k-medoid clusters, indicating a two-cluster solution.* ***(B)*** *Scaled sex bias trajectories in variability of cluster medoids for 2-cluster solution, corresponding to increasing male bias (Cluster 1, purple) and increasing female bias (Cluster 2, green)* ***(C)*** *Scaled variability sex bias trajectories of each IDP, grouped by cluster. (D) IDPs visualized in neuroanatomical context, colored by sex bias trajectory cluster. All IDPs included in k-medoid clustering protocol had significantly age-varying sex difference trajectories, as indicated by our FDR corrected likelihood ratio tests (black outlines). Abbrv: CT, cortical thickness; SA, surface area; Vol, volume; Std., standardized*

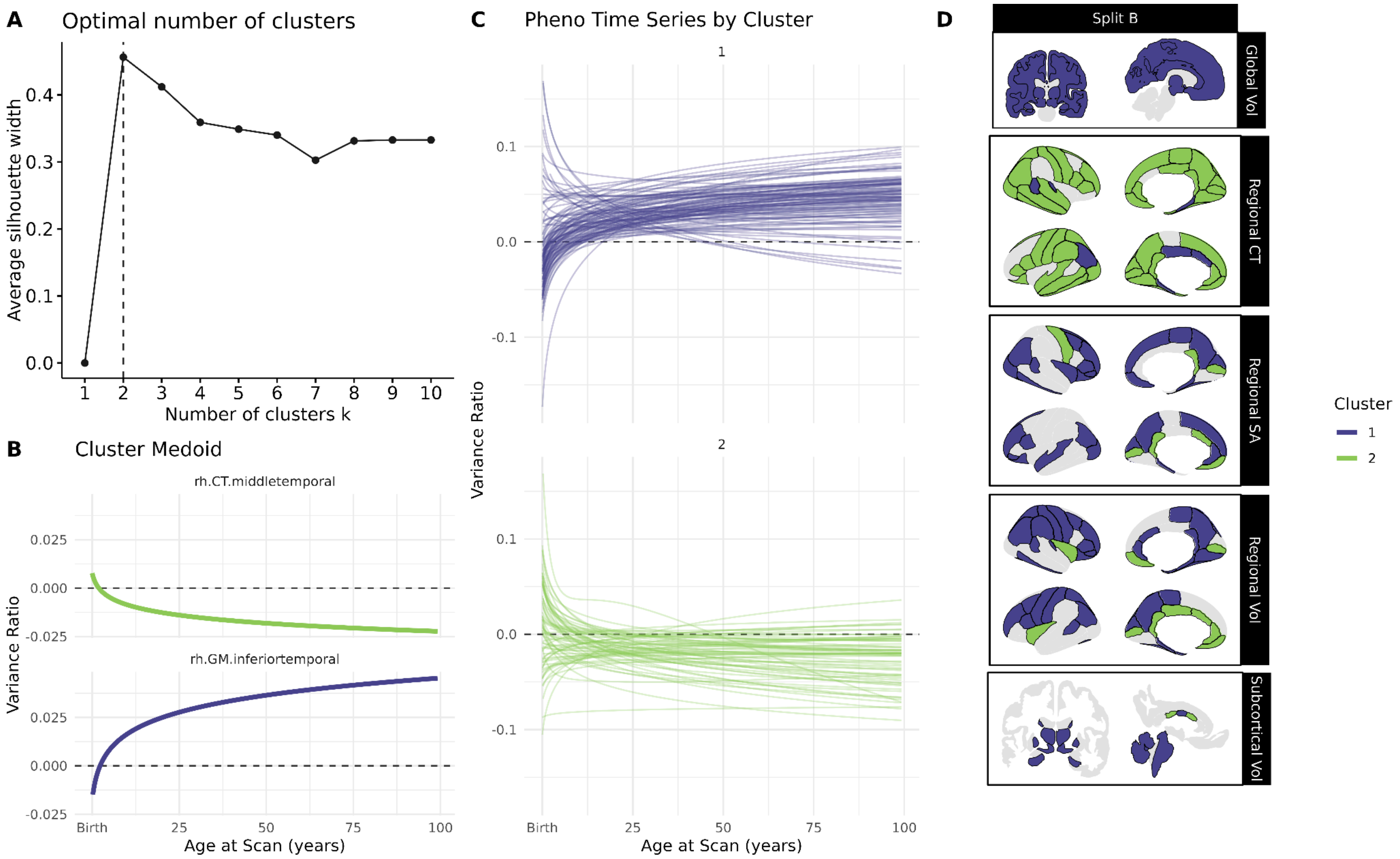


***Figure S17. Data-driven clustering of IDPs by sex difference trajectories in variability in test split B. (A)*** *Plot of average silhouette width by number of k-medoid clusters, indicating a two-cluster solution.* ***(B)*** *Scaled sex bias trajectories in variability of cluster medoids for 2-cluster solution, corresponding to increasing male bias (Cluster 1, purple) and increasing female bias (Cluster 2, green)* ***(C)*** *Scaled variability sex bias trajectories of each IDP, grouped by cluster. (D) IDPs visualized in neuroanatomical context, colored by sex bias trajectory cluster. All IDPs included in k-medoid clustering protocol had significantly age-varying sex difference trajectories, as indicated by our FDR corrected likelihood ratio tests (black outlines). Abbrv: CT, cortical thickness; SA, surface area; Vol, volume; Std., standardized*

#### B. Additional Case-Control Analysis Results

Here we present extended results from our case-control analyses of reference scores derived from sex-moderated brain charts, including tests of over- or under-representation in extreme centiles. We also compare disease effects uncovered by sex-moderated models (i.e. those including age-varying sex effects) to those uncovered by sex-intercept-only models, with side-by-side comparison in each neuropsychiatric condition. Finally, we test for case-control differences in reference scores derived from models controlling for total brain size in order to index regional disease atypicalities above and beyond total-brain-size effects (Figs S31 & S32). For full results underlying these figures, please see Supplemental Data 2 & 3.

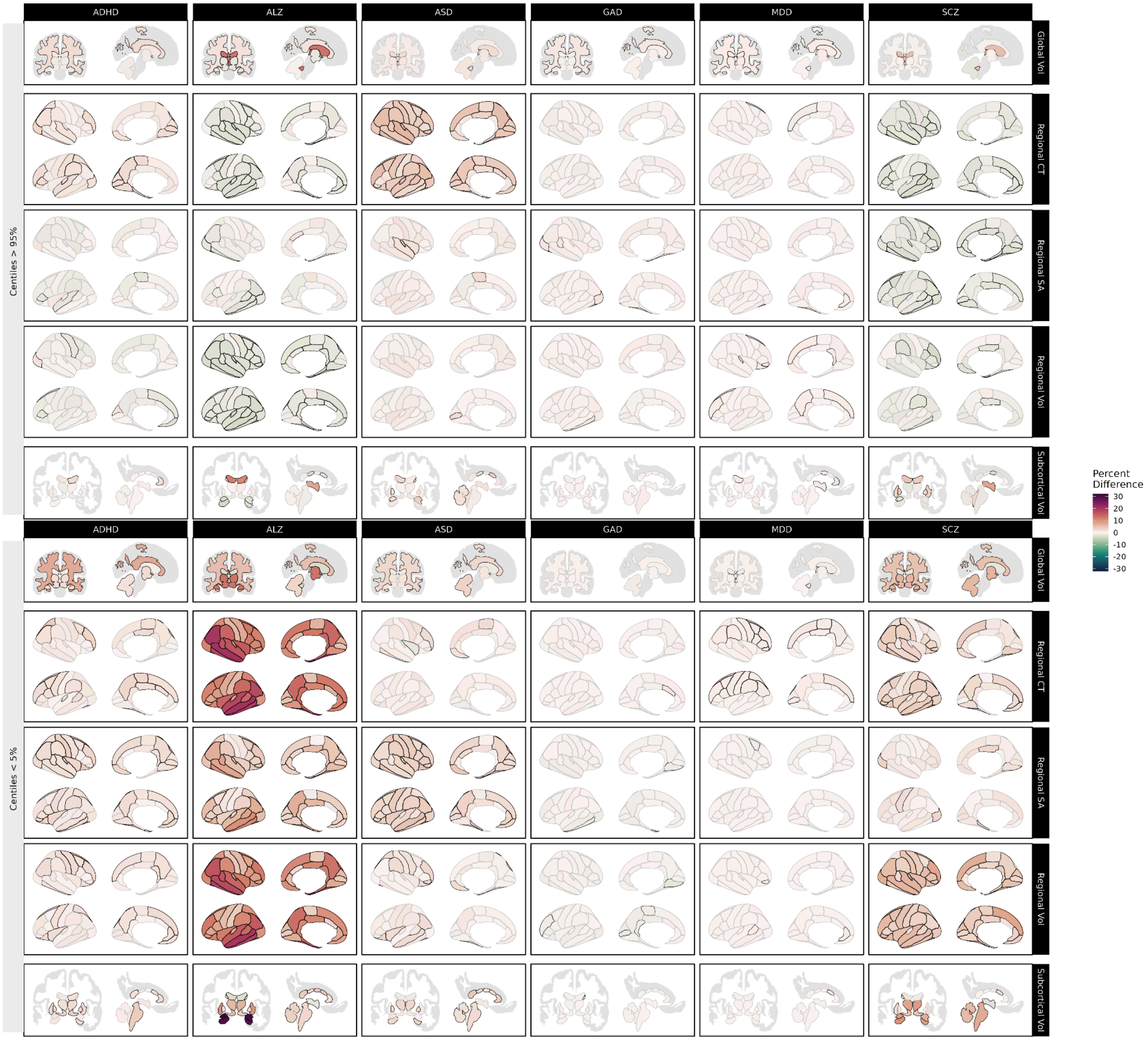


***Figure S18. Over- and under-representation of clinical cases in extreme centiles for each IDP.*** *Using centile scores derived from our sex-moderated brain charts, we compared the proportions of cases and site-matched controls with centiles above the 95th percentile (upper panel) and below the 5th percentile (lower panel). Reds indicate cases are over-represented relative to controls, while greens indicate that cases are under-represented relative to controls. Black outlines indicate significant effects at p <0.05, FDR-corrected. Abbrv: ADHD, attention-deficit/hyperactivity disorder; ALZ, Alzheimer's disease; ASD, autism spectrum disorder; GAD, generalized anxiety disorder; MDD, major depressive disorder; SCZ, schizophrenia; CT, cortical thickness; SA, surface area; Vol, volume.*

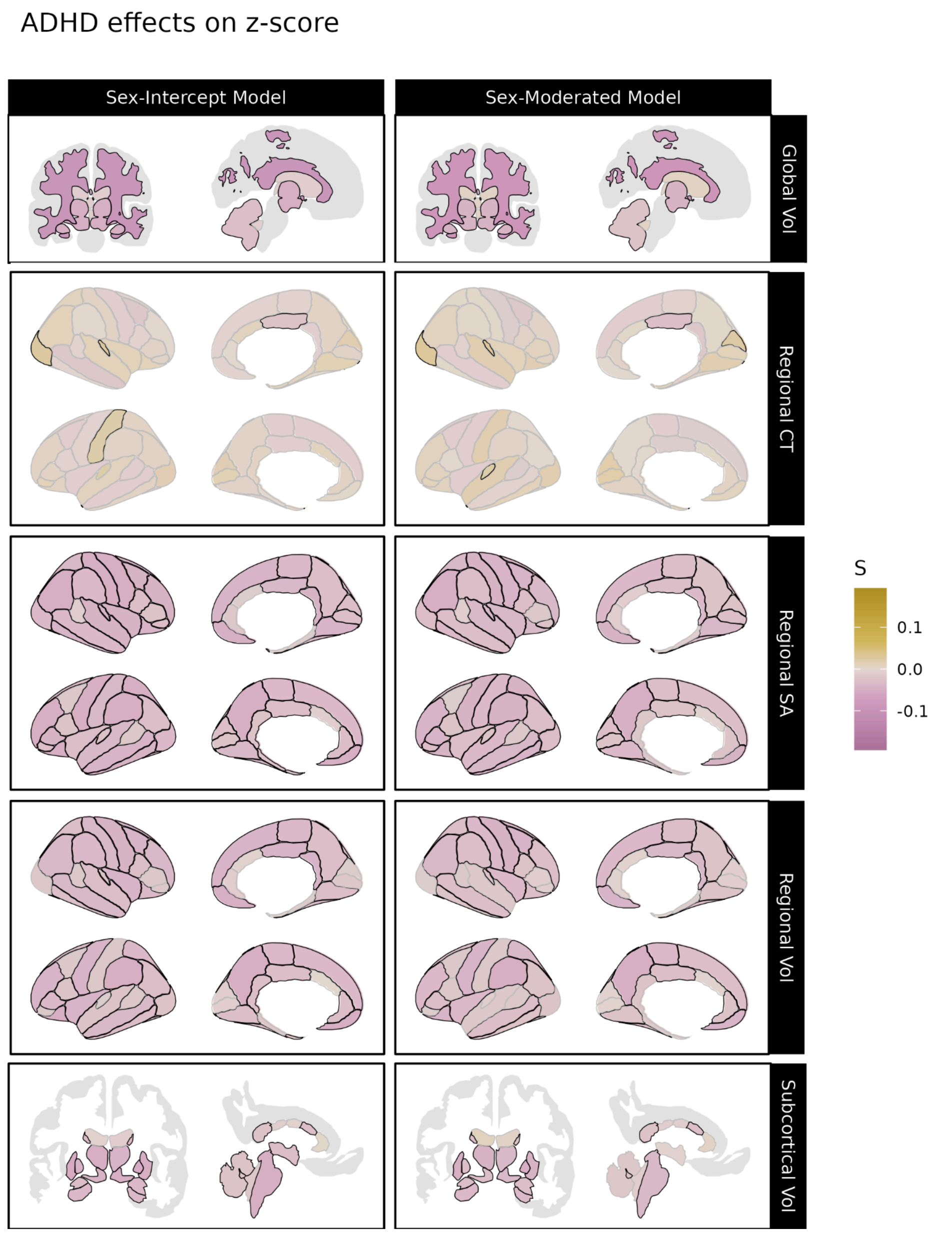


***Figure S19. Significant differences associated with ADHD in centile z-scores derived from sex-moderated and sex-intercept brain charts.*** *Differences in centile z-scores in individuals with attention-deficit/hyperactivity disorder (ADHD) relative to site-matched controls. The left panel displays differences in centile z-scores derived from sex-intercept models (i.e. models keeping sex differences in IDPs constant across the lifespan) while the right panel displays differences in centile z-scores derived from sex-moderated models, which allow age-by-sex effects. Gold indicates cases tend to have higher centile z-scores than controls for a given IDP, while purple indicates cases tend to have smaller centile z-scores. Black outlines indicate significant effects at p <0.05, FDR-corrected in each model. Abbrv: CT, cortical thickness; SA, surface area; Vol, volume.*

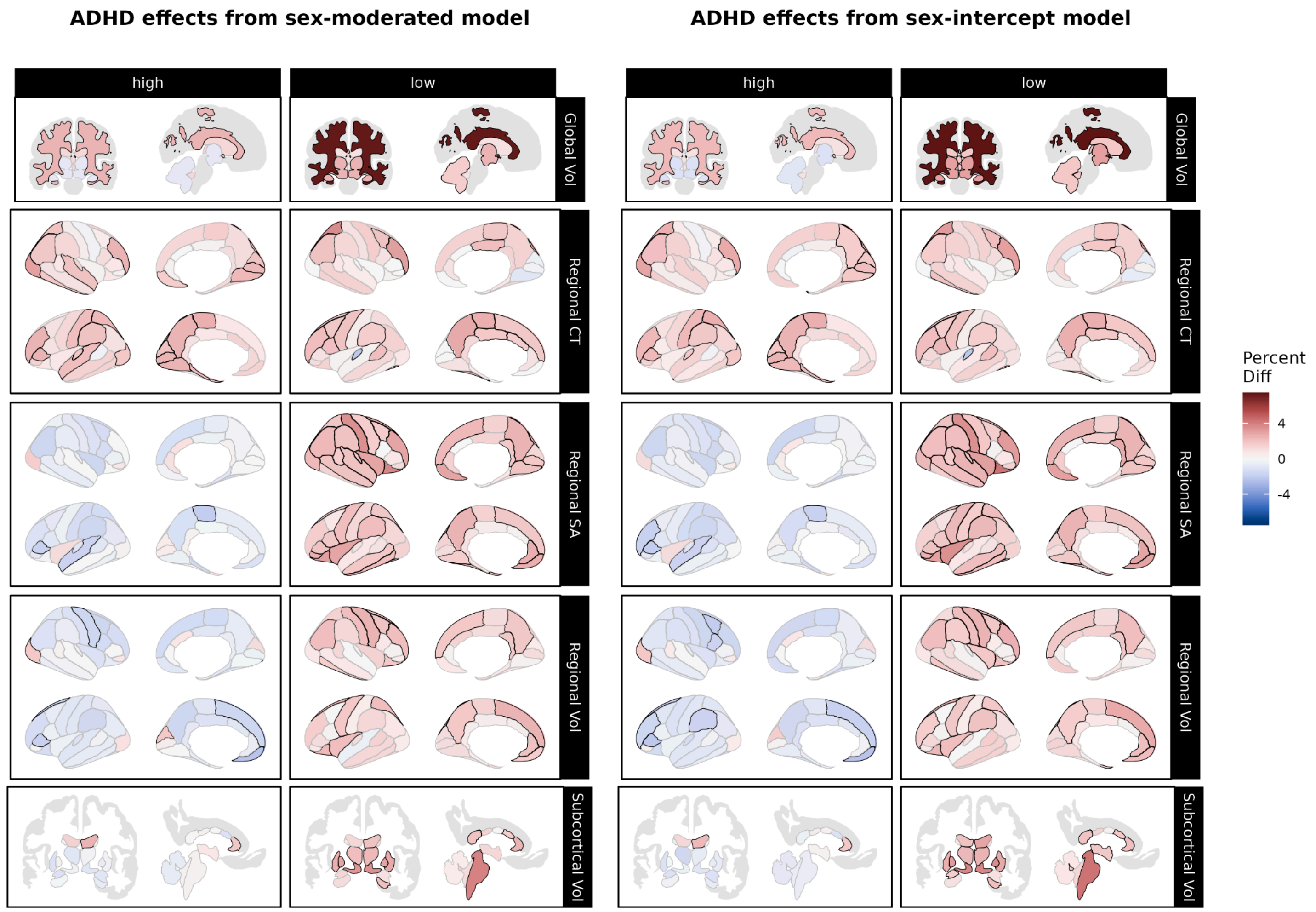


***Figure S20. Over- and under-representation of ADHD in extreme centiles according to sex-moderated and sex-intercept brain charts.*** *Using centile scores derived from our sex-moderated (left panel) and sex-intercept (right panel) brain charts, we compared the proportions of individuals with attention-deficit/hyperactivity disorder (ADHD) to the proportion of site-matched controls with centiles above the 95th percentile ("high", left column) and below the 5th percentile ("low", right column). Reds indicate cases are over-represented relative to controls, while greens indicate that cases are under-represented relative to controls. Black outlines indicate significant effects at p <0.05, FDR-corrected within each brain chart model. Abbrv: ADHD, attention-deficit/hyperactivity disorder; CT, cortical thickness; SA, surface area; Vol, volume.*

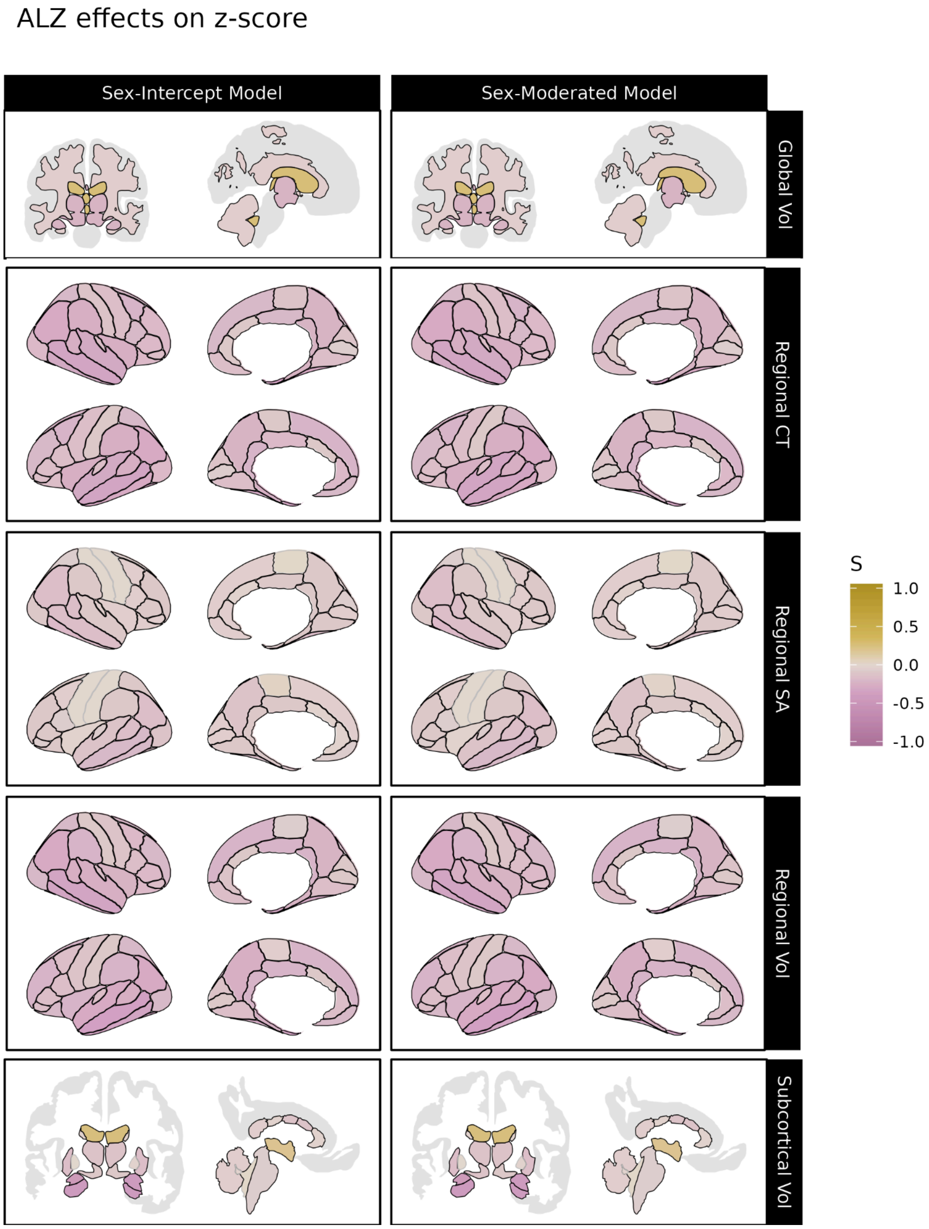


***Figure S21. Significant differences associated with Alzheimer's disease in centile z-scores derived from sex-moderated and sex-intercept brain charts.*** *Differences in centile z-scores in individuals with Alzheimer's disease relative to site-matched controls. The left panel displays differences in centile z-scores derived from sex-intercept models (i.e. models keeping sex differences in IDPs constant across the lifespan) while the right panel displays differences in centile z-scores derived from sex-moderated models, which allow age-by-sex effects. Gold indicates cases tend to have higher centile z-scores than controls for a given IDP, while purple indicates cases tend to have smaller centile z-scores. Black outlines indicate significant effects at p <0.05, FDR-corrected in each model. Abbrv: CT, cortical thickness; SA, surface area; Vol, volume.*

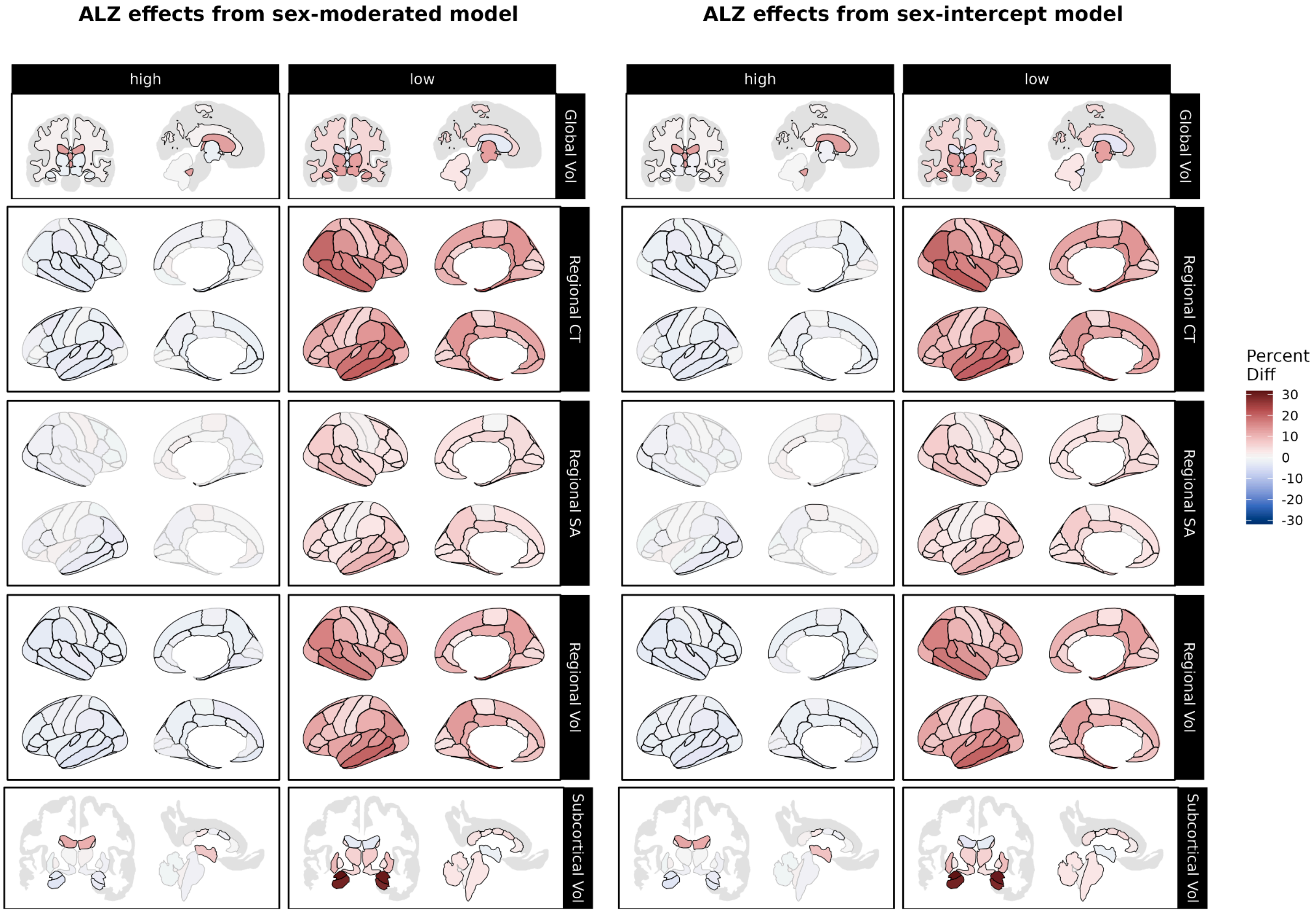


***Figure S22. Over- and under-representation of Alzheimer's disease in extreme centiles according to sex-moderated and sex-intercept brain charts.*** *Using centile scores derived from our sex-moderated (left panel) and sex-intercept (right panel) brain charts, we compared the proportions of individuals with Alzheimer's disease to the proportion of site-matched controls with centiles above the 95th percentile ("high", left column) and below the 5th percentile ("low", right column). Reds indicate cases are over-represented relative to controls, while greens indicate that cases are under-represented relative to controls. Black outlines indicate significant effects at p <0.05, FDR-corrected within each brain chart model. Abbrv: ALZ, Alzheimer's disease; CT, cortical thickness; SA, surface area; Vol, volume.*

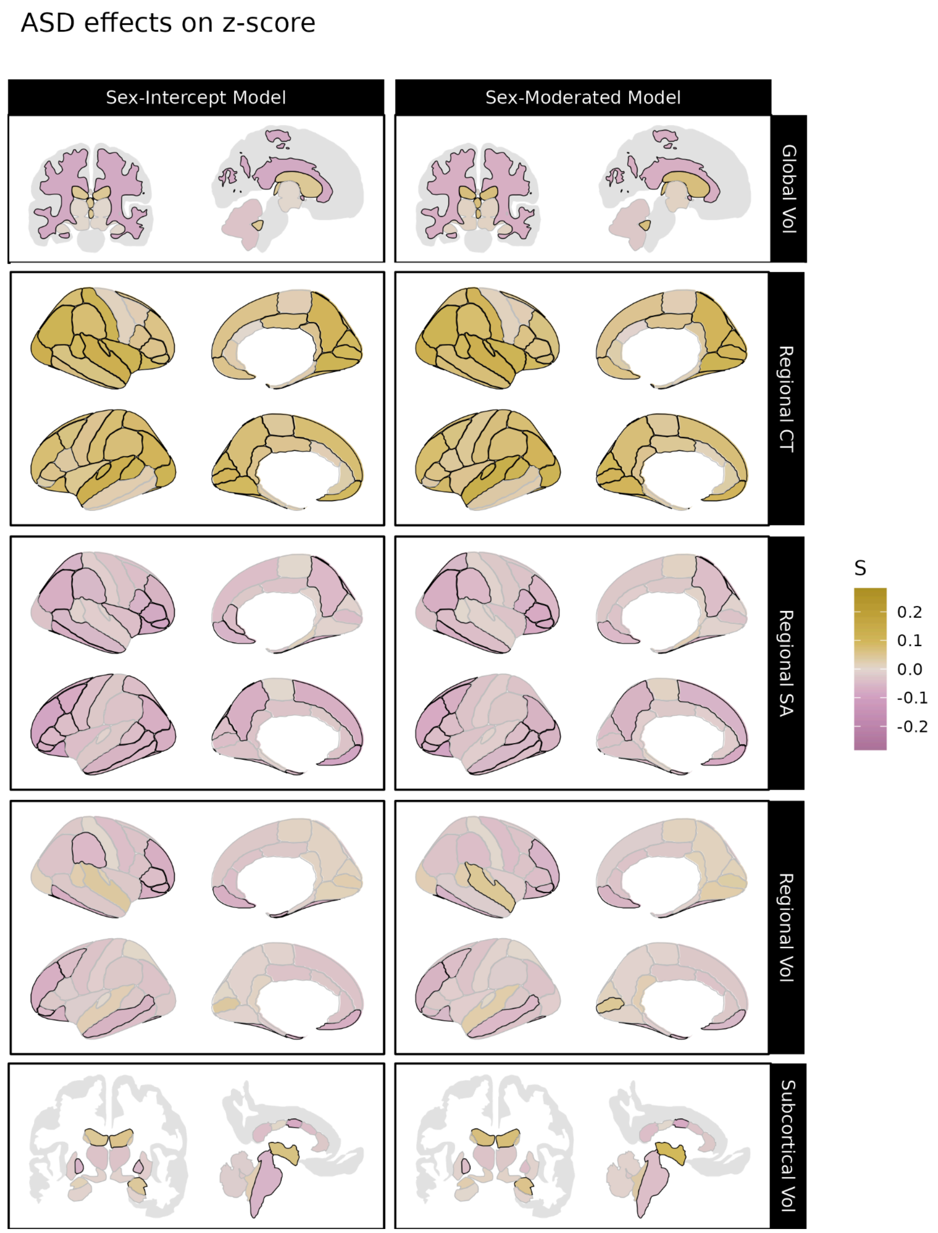


***Figure S23. Significant differences associated with autism spectrum disorder in centile z-scores derived from sex-moderated and sex-intercept brain charts.*** *Differences in centile z-scores in individuals with autism spectrum disorder relative to site-matched controls. The left panel displays differences in centile z-scores derived from sex-intercept models (i.e. models keeping sex differences in IDPs constant across the lifespan) while the right panel displays differences in centile z-scores derived from sex-moderated models, which allow age-by-sex effects. Gold indicates cases tend to have higher centile z-scores than controls for a given IDP, while purple indicates cases tend to have smaller centile z-scores. Black outlines indicate significant effects at p <0.05, FDR-corrected in each model. Abbrv: CT, cortical thickness; SA, surface area; Vol, volume.*

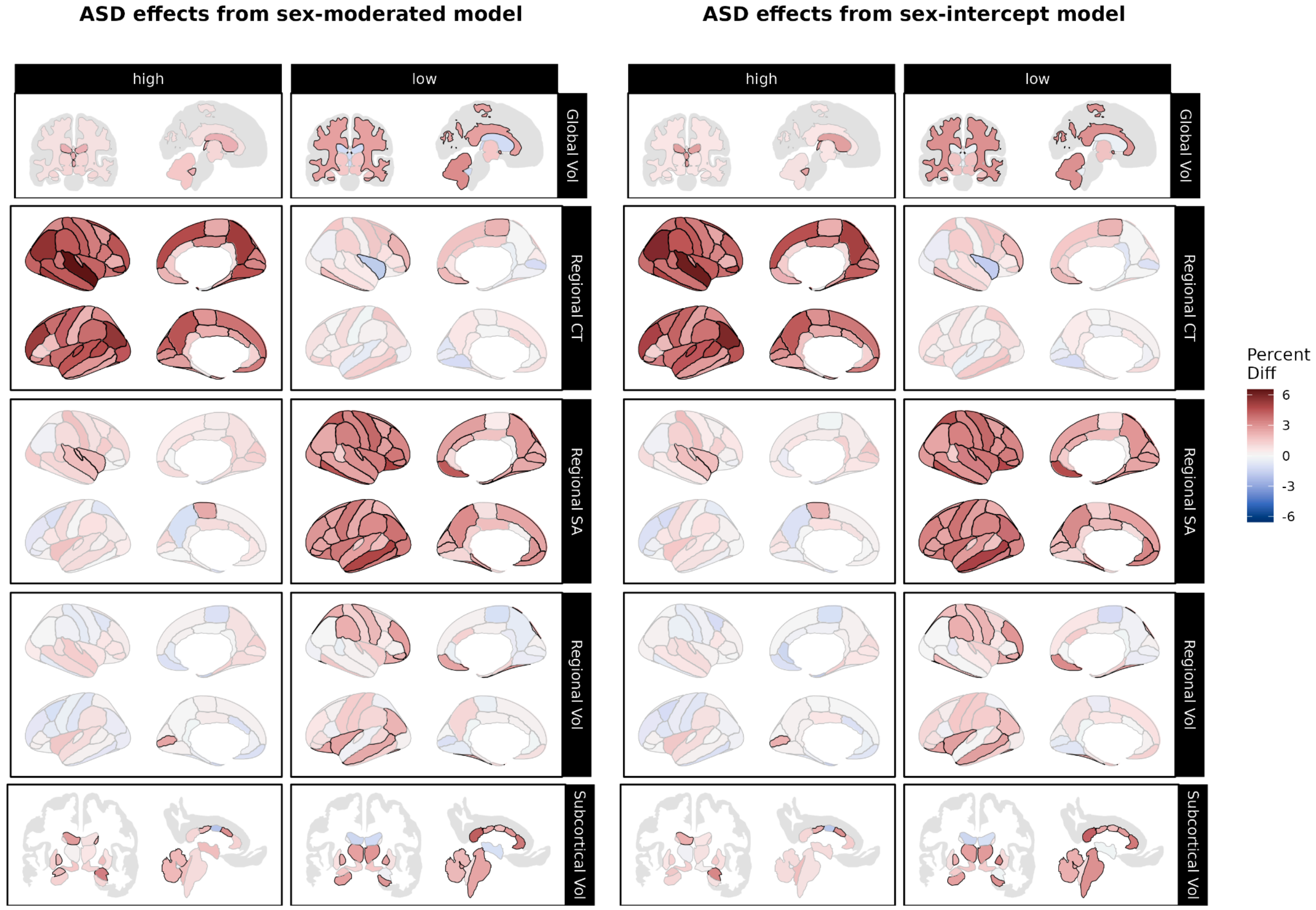


***Figure S24. Over- and under-representation of ASD in extreme centiles according to sex-moderated and sex-intercept brain charts.*** *Using centile scores derived from our sex-moderated (left panel) and sex-intercept (right panel) brain charts, we compared the proportions of individuals with autism spectrum disorder (ASD) to the proportion of site-matched controls with centiles above the 95th percentile ("high", left column) and below the 5th percentile ("low", right column). Reds indicate cases are over-represented relative to controls, while greens indicate that cases are under-represented relative to controls. Black outlines indicate significant effects at p <0.05, FDR-corrected within each brain chart model. Abbrv: ASD, autism spectrum disorder; CT, cortical thickness; SA, surface area; Vol, volume.*

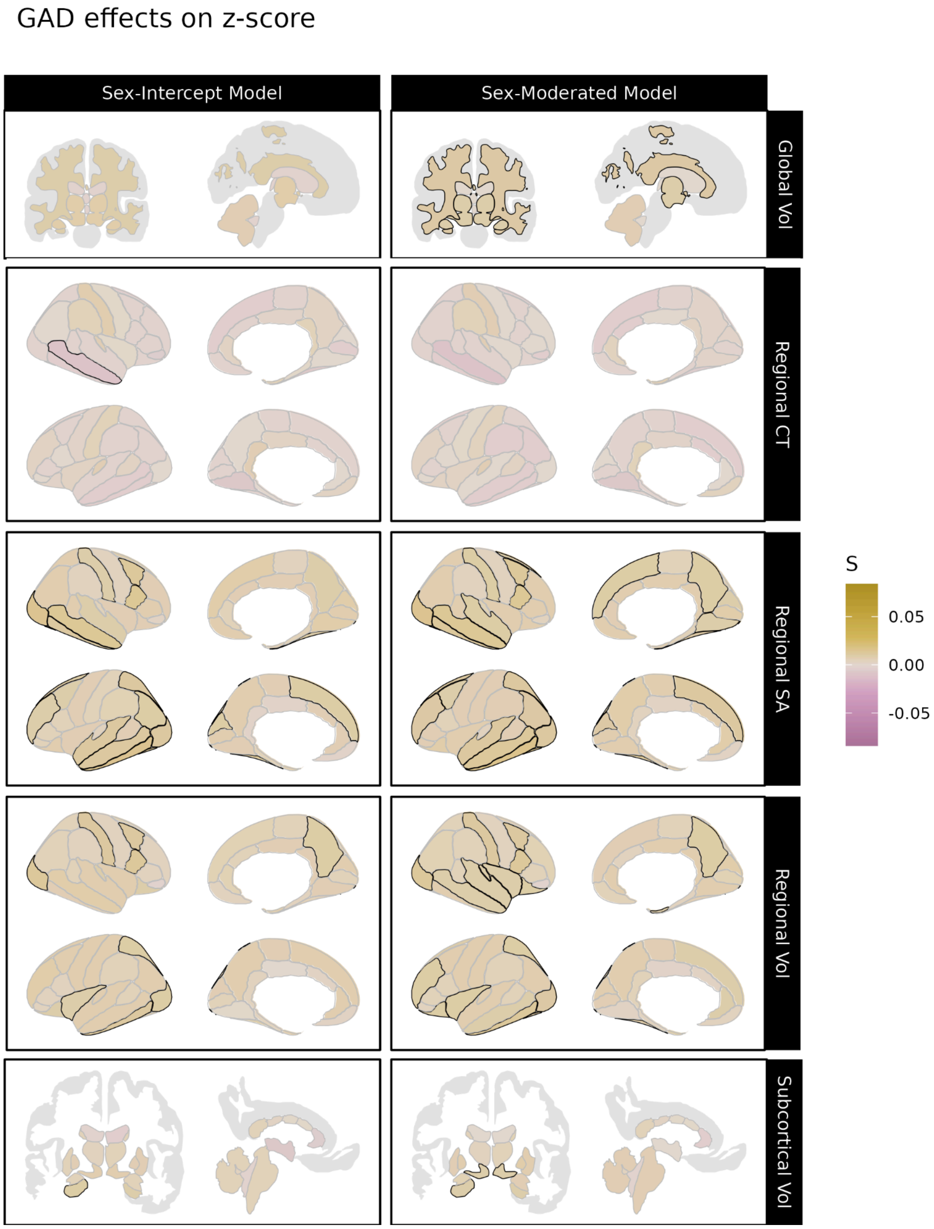


***Figure S25. Significant differences associated with generalized anxiety disorder in centile z-scores derived from sex-moderated and sex-intercept brain charts.*** *Differences in centile z-scores in individuals with generalized anxiety disorder relative to site-matched controls. The left panel displays differences in centile z-scores derived from sex-intercept models (i.e. models keeping sex differences in IDPs constant across the lifespan) while the right panel displays differences in centile z-scores derived from sex-moderated models, which allow age-by-sex effects. Gold indicates cases tend to have higher centile z-scores than controls for a given IDP, while purple indicates cases tend to have smaller centile z-scores. Black outlines indicate significant effects at p <0.05, FDR-corrected in each model. Abbrv: CT, cortical thickness; SA, surface area; Vol, volume.*

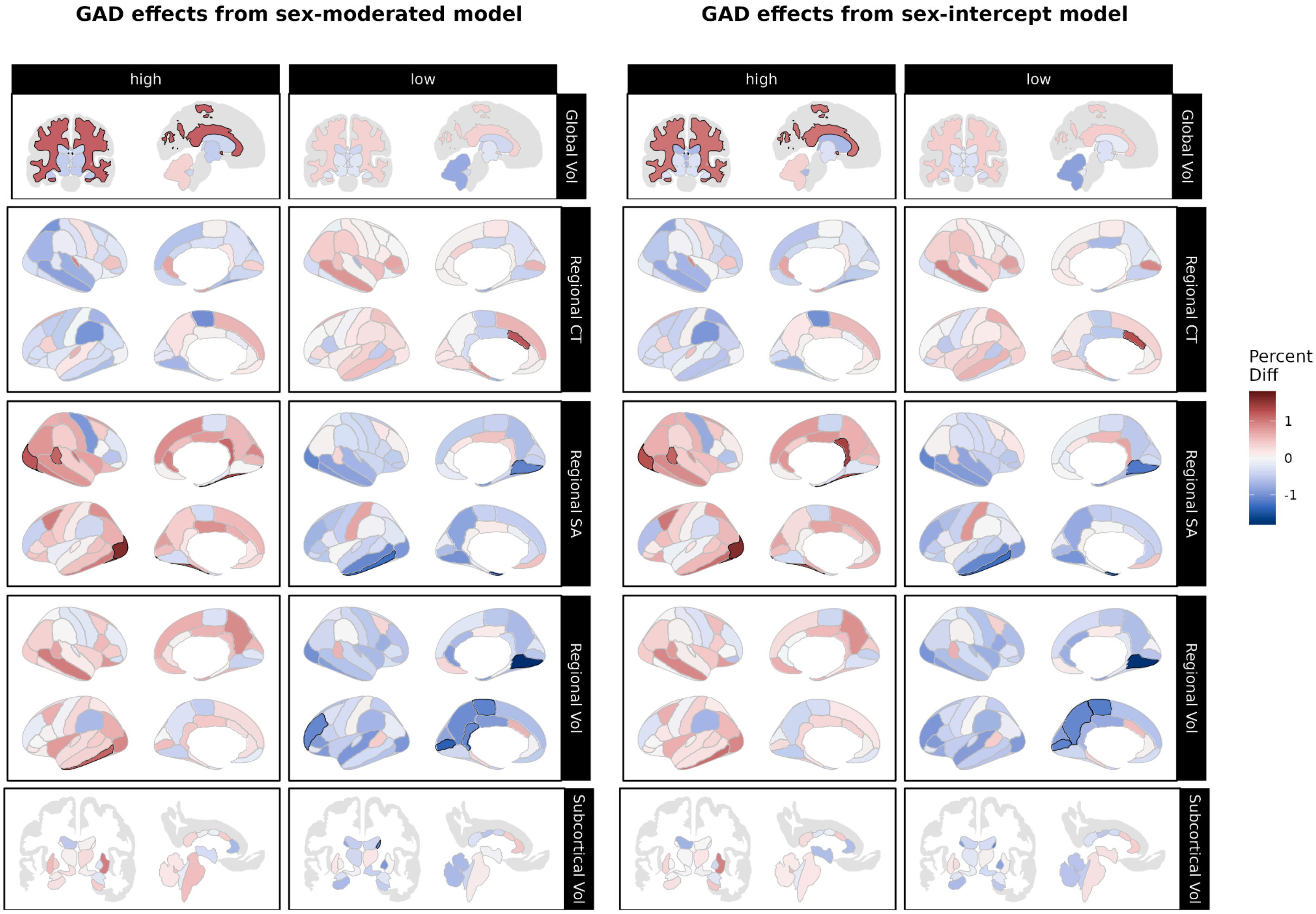


***Figure S26. Over- and under-representation of GAD in extreme centiles according to sex-moderated and sex-intercept brain charts.*** *Using centile scores derived from our sex-moderated (left panel) and sex-intercept (right panel) brain charts, we compared the proportions of individuals with generalized anxiety disorder (GAD) to the proportion of site-matched controls with centiles above the 95th percentile ("high", left column) and below the 5th percentile ("low", right column). Reds indicate cases are over-represented relative to controls, while greens indicate that cases are under-represented relative to controls. Black outlines indicate significant effects at p <0.05, FDR-corrected within each brain chart model. Abbrv: GAD, generalized anxiety disorder; CT, cortical thickness; SA, surface area; Vol, volume.*

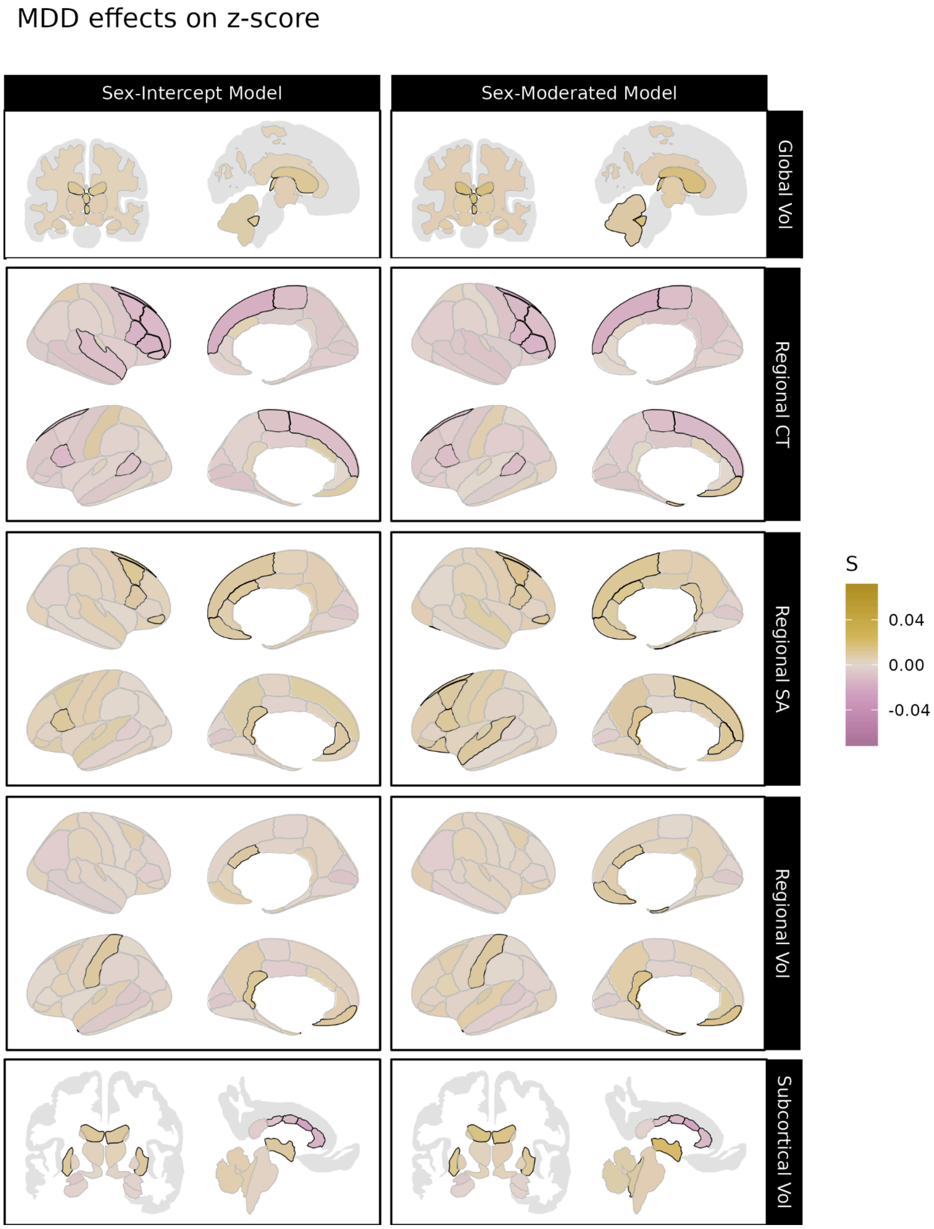


***Figure S27. Significant differences associated with major depressive disorder in centile z-scores derived from sex-moderated and sex-intercept brain charts.*** *Differences in centile z-scores in individuals with major depressive disorder relative to site-matched controls. The left panel displays differences in centile z-scores derived from sex-intercept models (i.e. models keeping sex differences in IDPs constant across the lifespan) while the right panel displays differences in centile z-scores derived from sex-moderated models, which allow age-by-sex effects. Gold indicates cases tend to have higher centile z-scores than controls for a given IDP, while purple indicates cases tend to have smaller centile z-scores. Black outlines indicate significant effects at p <0.05, FDR-corrected in each model. Abbrv: CT, cortical thickness; SA, surface area; Vol, volume.*

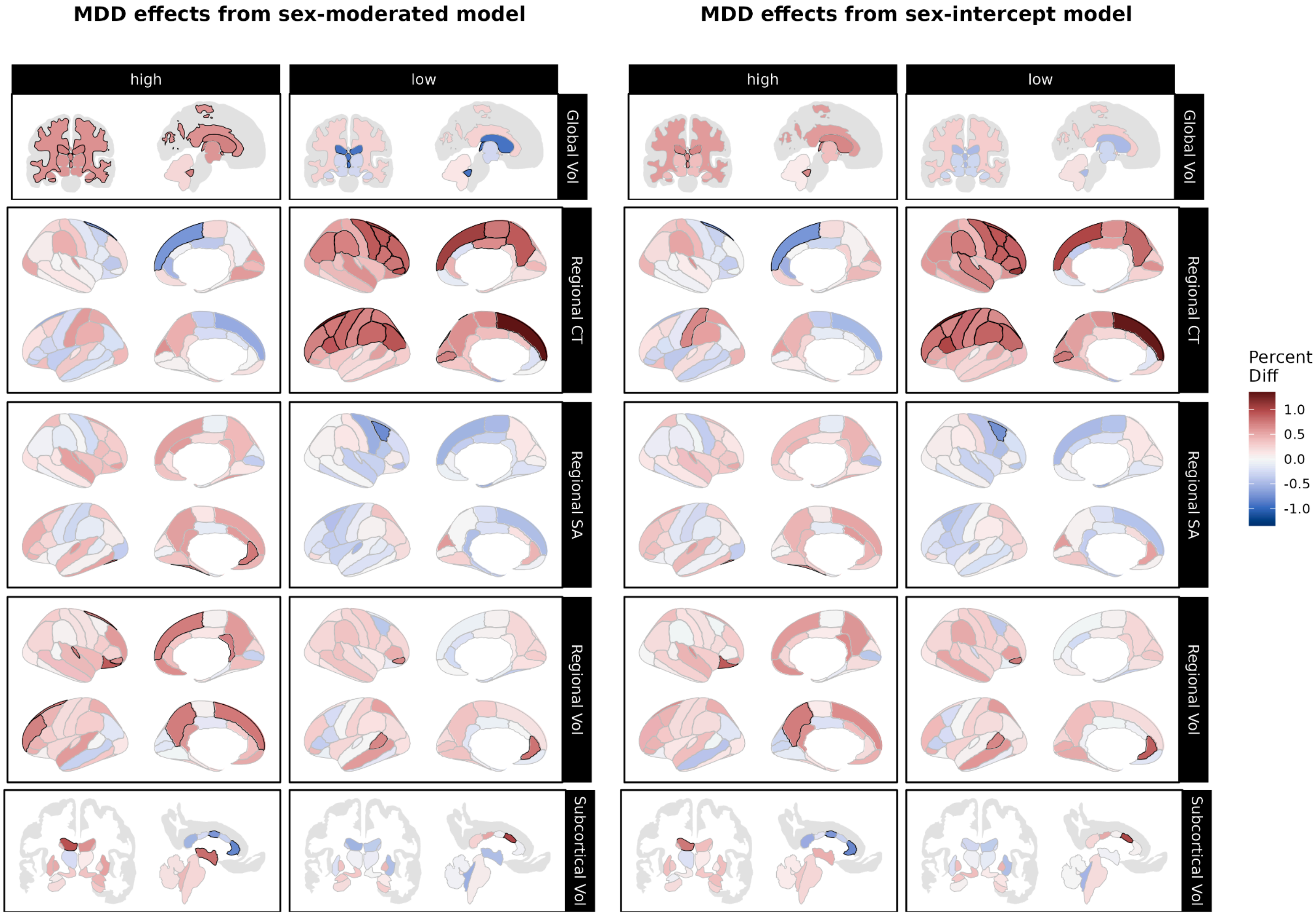


***Figure S28. Over- and under-representation of MDD in extreme centiles according to sex-moderated and sex-intercept brain charts.*** *Using centile scores derived from our sex-moderated (left panel) and sex-intercept (right panel) brain charts, we compared the proportions of individuals with major depressive disorder (MDD) to the proportion of site-matched controls with centiles above the 95th percentile ("high", left column) and below the 5th percentile ("low", right column). Reds indicate cases are over-represented relative to controls, while greens indicate that cases are under-represented relative to controls. Black outlines indicate significant effects at p <0.05, FDR-corrected within each brain chart model. Abbrv: MDD, major depressive disorder; CT, cortical thickness; SA, surface area; Vol, volume.*

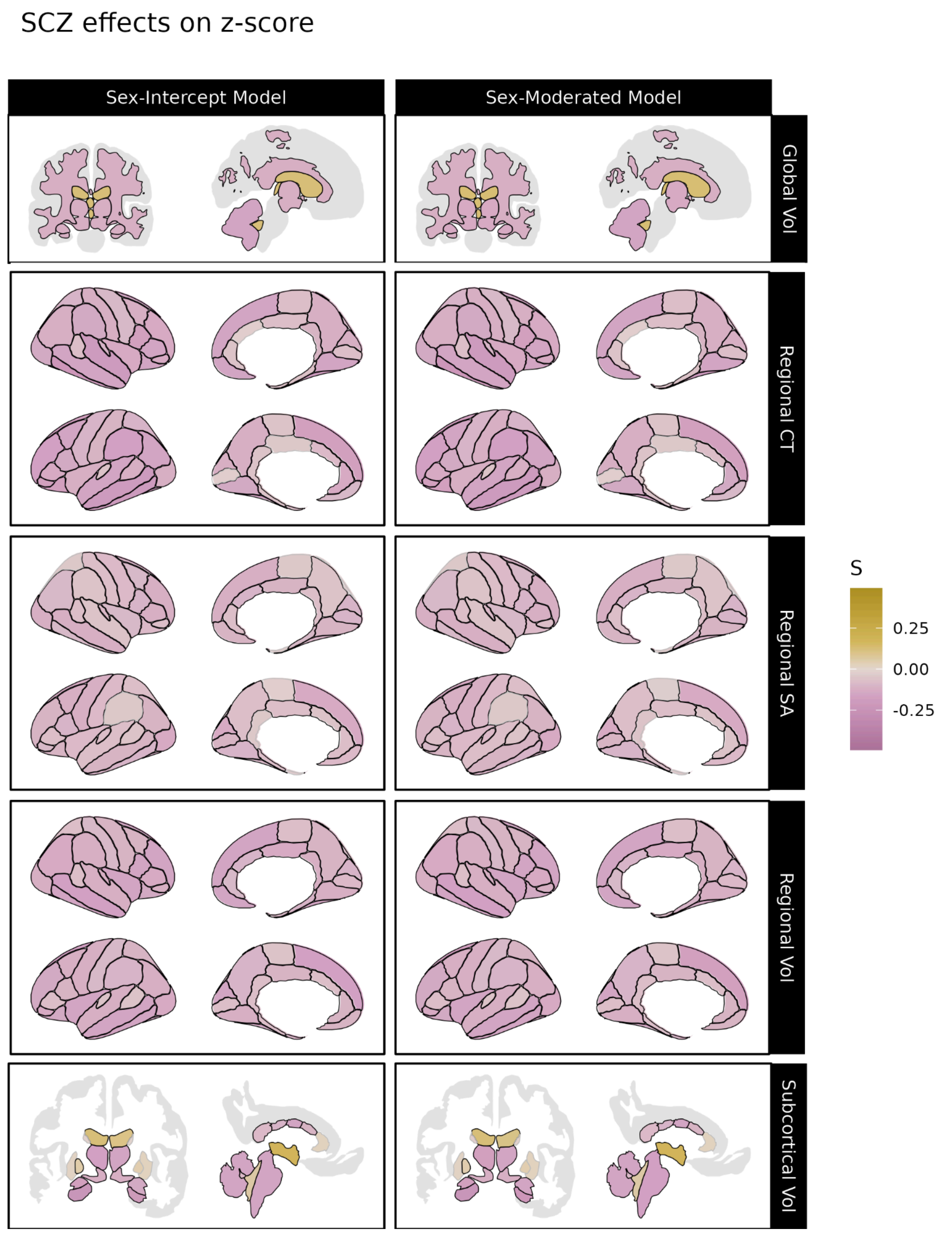


***Figure S29. Significant differences associated with schizophrenia in centile z-scores derived from sex-moderated and sex-intercept brain charts.*** *Differences in centile z-scores in individuals with schizophrenia relative to site-matched controls. The left panel displays differences in centile z-scores derived from sex-intercept models (i.e. models keeping sex differences in IDPs constant across the lifespan) while the right panel displays differences in centile z-scores derived from sex-moderated models, which allow age-by-sex effects. Gold indicates cases tend to have higher centile z-scores than controls for a given IDP, while purple indicates cases tend to have smaller centile z-scores. Black outlines indicate significant effects at p <0.05, FDR-corrected in each model. Abbrv: CT, cortical thickness; SA, surface area; Vol, volume.*

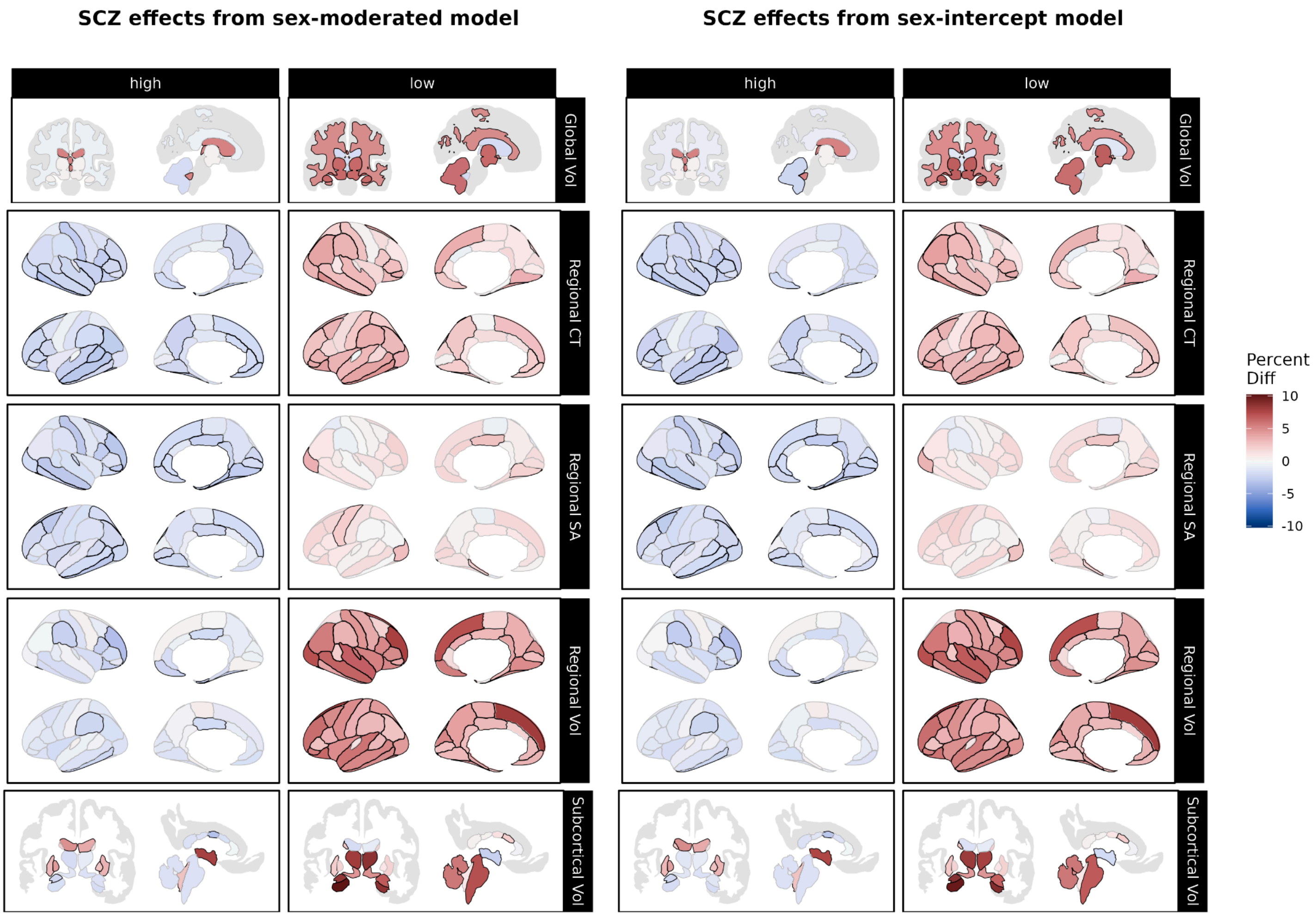


***Figure S30. Over- and under-representation of schizophrenia in extreme centiles according to sex-moderated and sex-intercept brain charts.*** *Using centile scores derived from our sex-moderated (left panel) and sex-intercept (right panel) brain charts, we compared the proportions of individuals with schizophrenia to the proportion of site-matched controls with centiles above the 95th percentile ("high", left column) and below the 5th percentile ("low", right column). Reds indicate cases are over-represented relative to controls, while greens indicate that cases are under-represented relative to controls. Black outlines indicate significant effects at p <0.05, FDR-corrected within each brain chart model. Abbrv: SCZ, schizophrenia; CT, cortical thickness; SA, surface area; Vol, volume.*

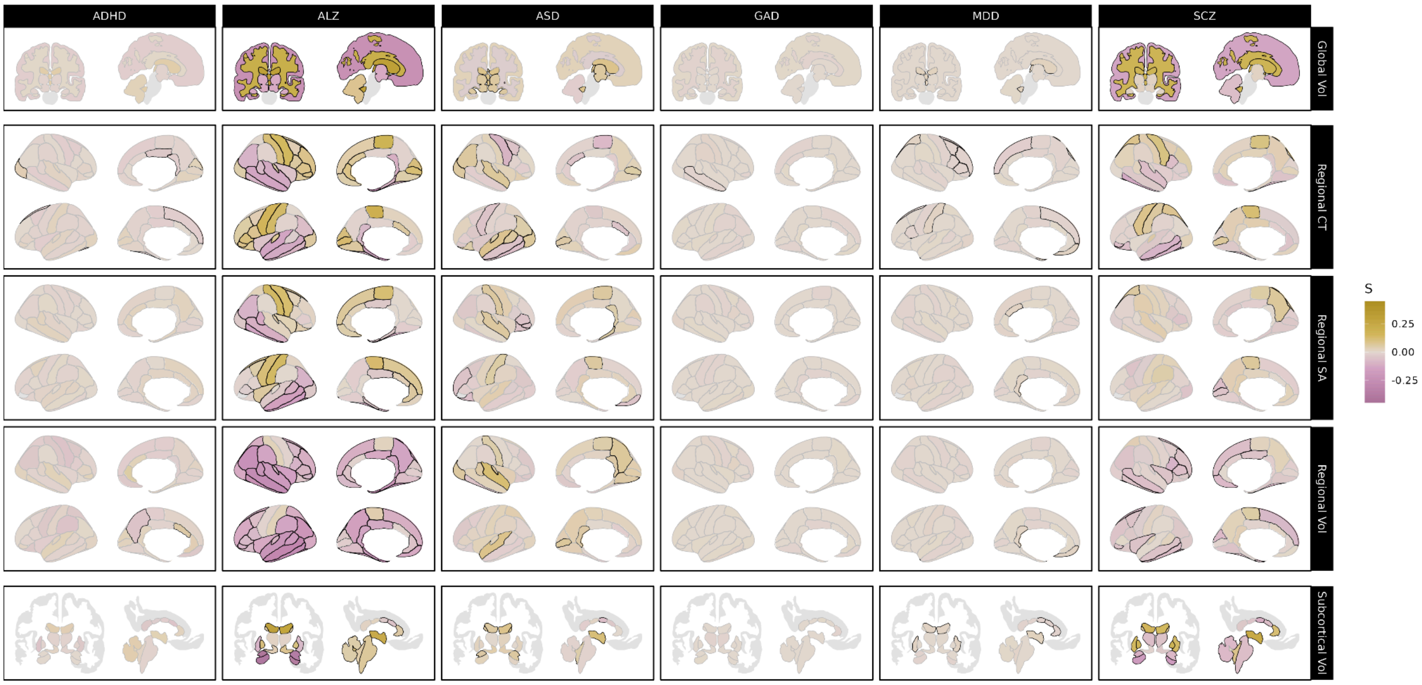


***Figure S31. Case-control differences in centile z-scores derived from models controlling for total brain size.*** *Case-control differences in centile z-score. Effect size (S) using the robust effect size index (RESI), with positive values indicating cases tend to be larger than controls and negative values indicating they tend to be smaller. Black outlines indicate significant effects at p <0.05, FDR-corrected. Abbrv: ADHD, attention-deficit/hyperactivity disorder; ALZ, Alzheimer's disease; ASD, autism spectrum disorder; GAD, generalized anxiety disorder; MDD, major depressive disorder; SCZ, schizophrenia; CT, cortical thickness; SA, surface area; Vol, volume.*

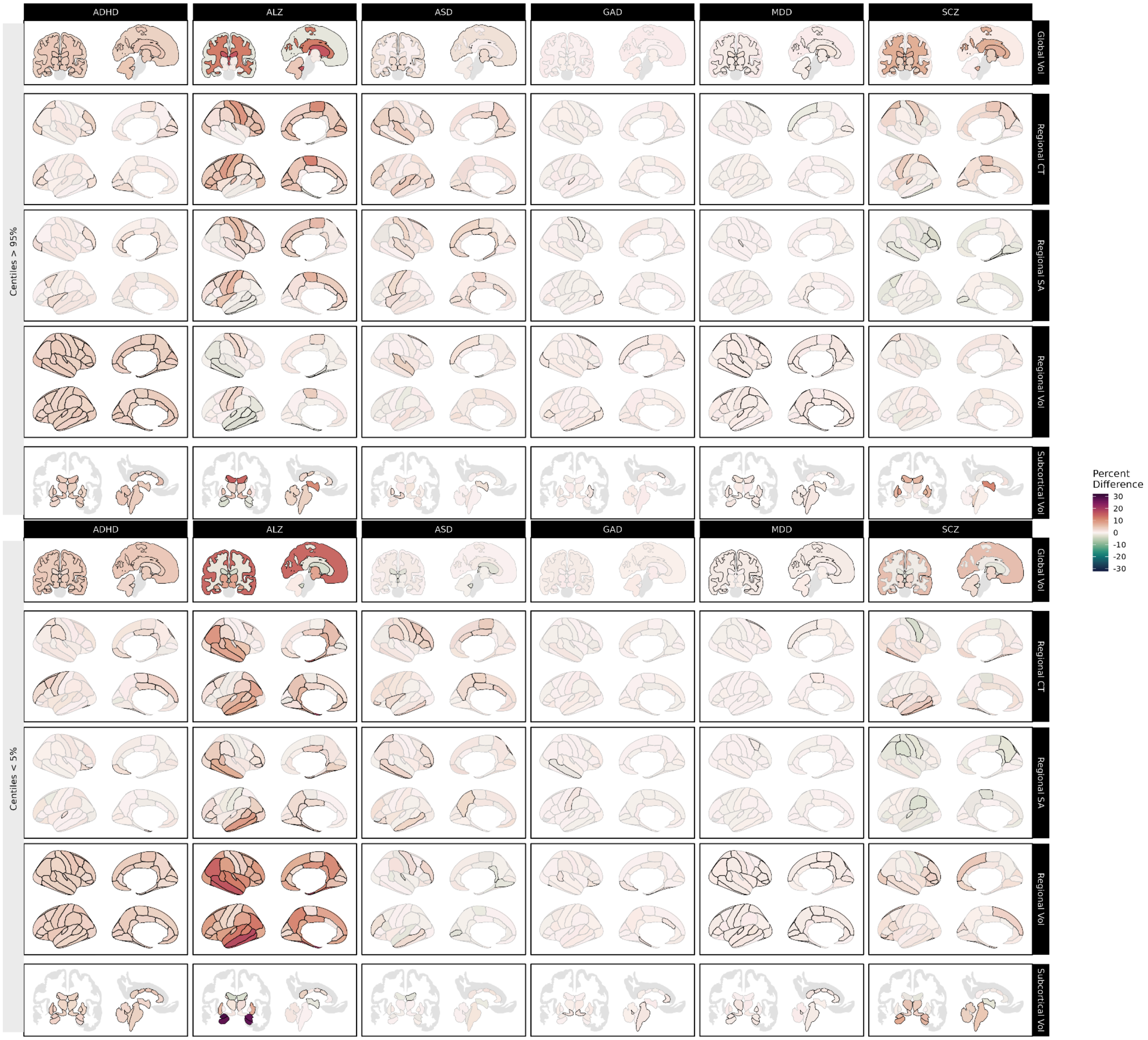


***Figure S32. Over- and under-representation of clinical cases in extreme centiles for each IDP, controlling for total brain size.*** *Using centile scores derived from our sex-moderated brain charts with a nonlinear control for total brain size, we compared the proportions of cases and site-matched controls with centiles above the 95th percentile (upper panel) and below the 5th percentile (lower panel). Reds indicate cases are over-represented relative to controls, while greens indicate that cases are under-represented relative to controls. Black outlines indicate significant effects at p <0.05, FDR-corrected. Abbrv: ADHD, attention-deficit/hyperactivity disorder; ALZ, Alzheimer's disease; ASD, autism spectrum disorder; GAD, generalized anxiety disorder; MDD, major depressive disorder; SCZ, schizophrenia; CT, cortical thickness; SA, surface area; Vol, volume.*

#### C. Sensitivity Analysis Results

Here we show the results of our sensitivity analyses. For the complete results underlying these figures, please see Supplemental Data S1-S3.

**C1. Weighting by scan quality**

To assess sensitivity to scan reconstruction quality above and beyond our scan quality inclusion criteria, we refit all train and test models weighting inversely by surface hole number. Despite a lack of surface hole data in scans younger than 1.5 years of age, results were largely consistent with our main analyses, including the presence and directionality of age-varying sex effects. It is worth noting that these *a priori* grouping of features by sex difference trajectories did show an increase in the proportion of converging phenotypes (A, Diverging = 21.2%, Converging = 33.1%; B, Diverging = 21.6%, Converging = 36.9%; Figs S40 and S41) relative to our main analyses. However, k-medoids clustering (Fig S42-S45) supported diverging male and female trajectories as the dominant mode of age-related change, consistent with our main analyses.

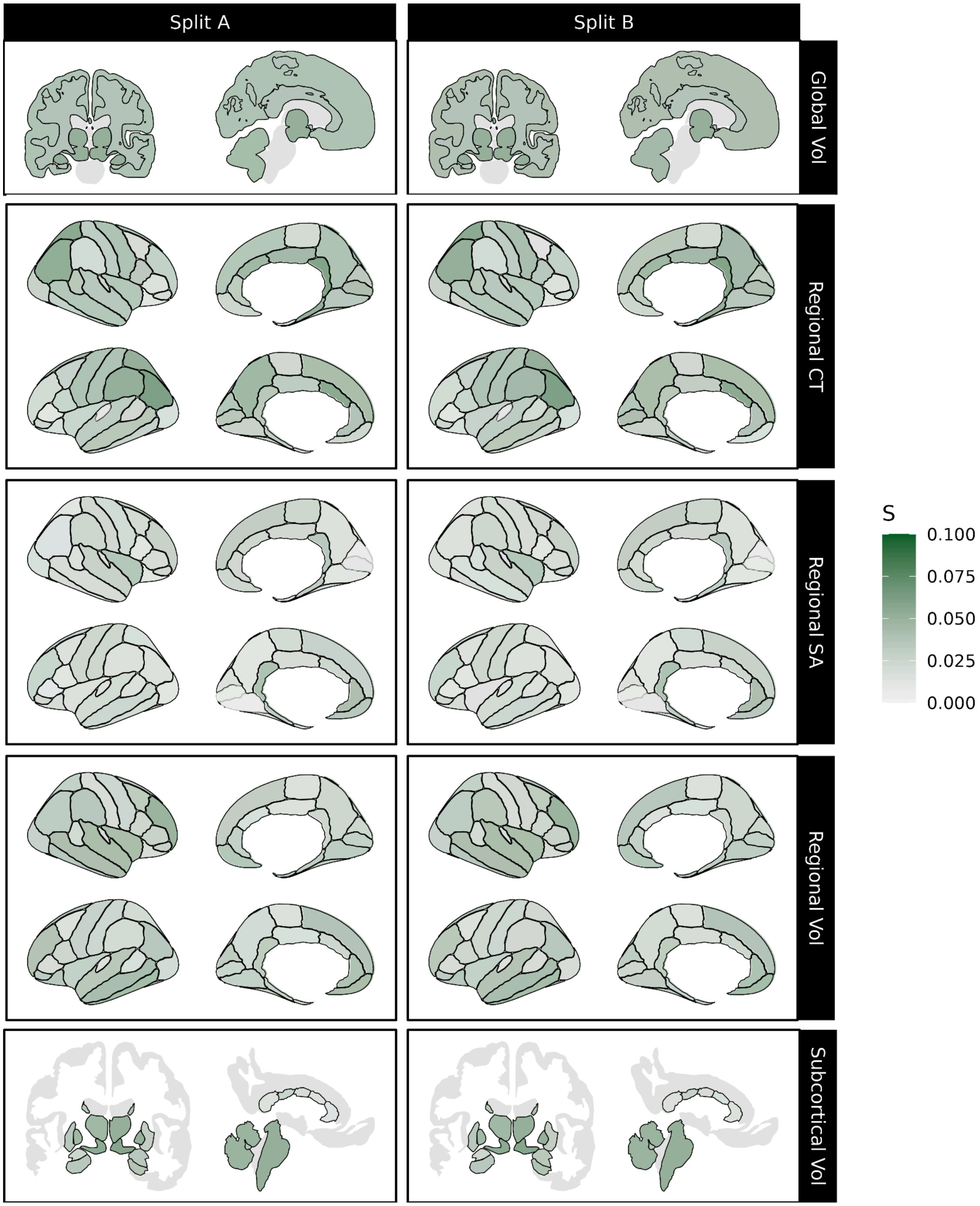


***Figure S33. Age-varying sex-biases persist when weighting by scan quality.*** *Magnitude of the age-varying component of sex (age-by-sex term) on IDPs each test split when weighted inversely by surface hole number. Effect sizes (S) are reported using the robust effect size index (RESI). Black outlines indicate significant effects at p <0.05, using false discovery rate (FDR) correction for multiple comparisons. Abbrv: CT, cortical thickness; SA, surface area; Vol, volume.*

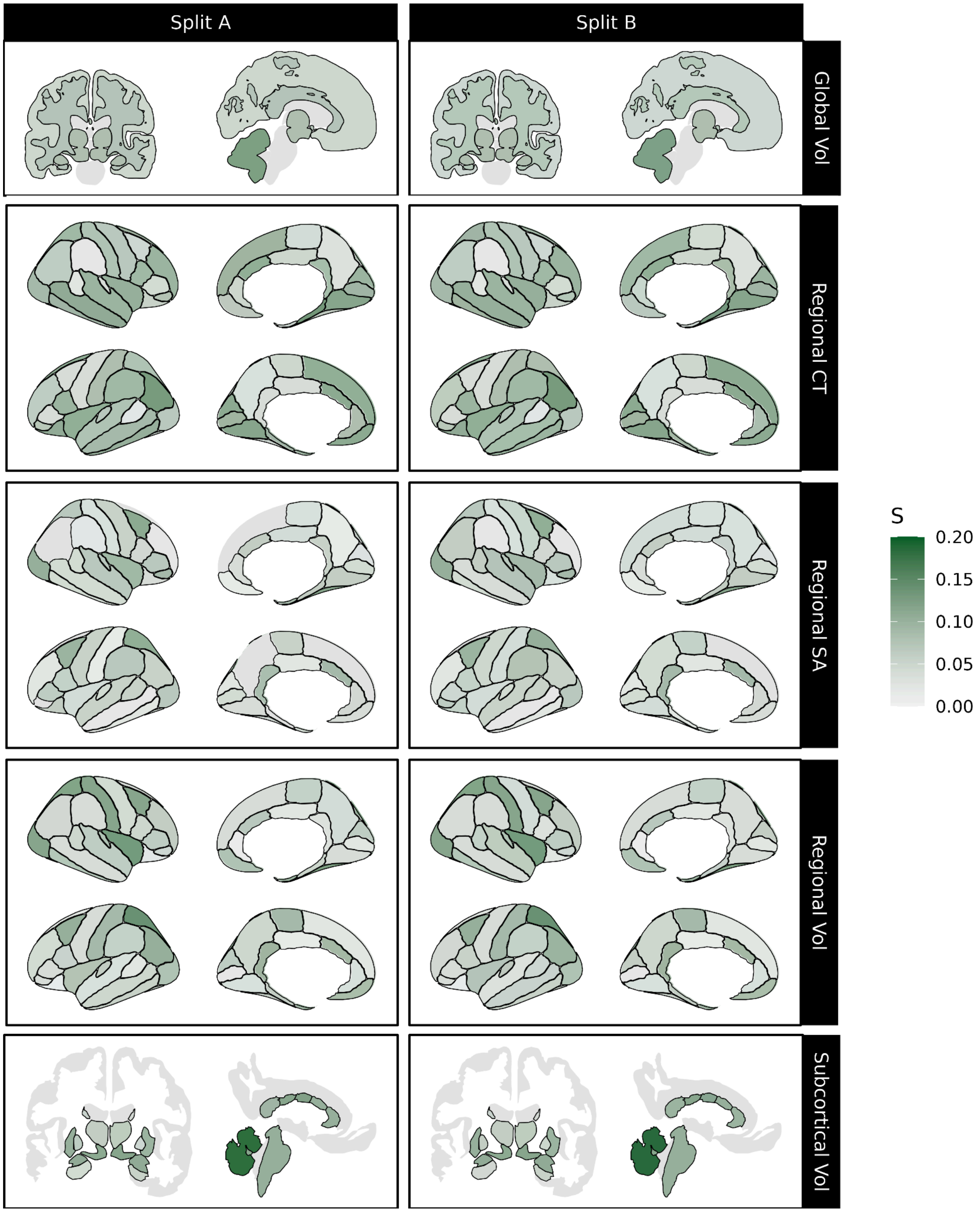


***Figure S34. Sex effects survive when controlling for total brain size and weighting by scan quality.*** *Magnitude of sex effects (sex intercept and age-by-sex interaction) on IDPs each test split when controlling for total brain size and weighting each scan inversely by surface hole number. Effect sizes (S) are reported using the robust effect size index (RESI). Black outlines indicate significant effects at p <0.05, using false discovery rate (FDR) correction for multiple comparisons. Abbrv: CT, cortical thickness; SA, surface area; Vol, volume.*

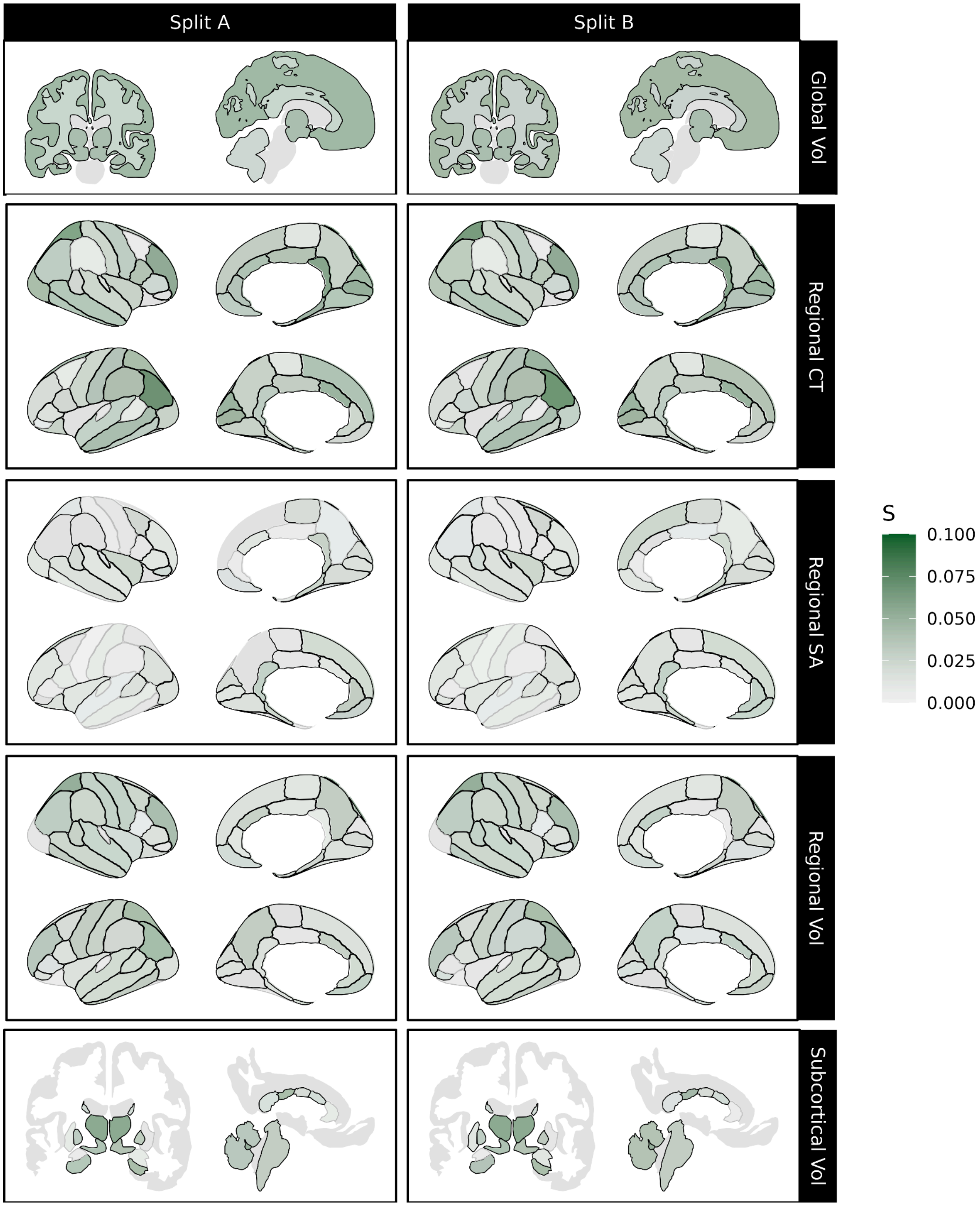


***Figure S35. Age-varying sex effect when controlling for total brain size and weighting by scan quality.*** *Magnitude of age-by-sex interaction effect on IDPs each test split when controlling for total brain size and weighting data inversely to surface hole number. Effect sizes (S) are reported using the robust effect size index (RESI). Black outlines indicate significant effects at p <0.05, using false discovery rate (FDR) correction for multiple comparisons. Abbrv: CT, cortical thickness; SA, surface area; Vol, volume.*

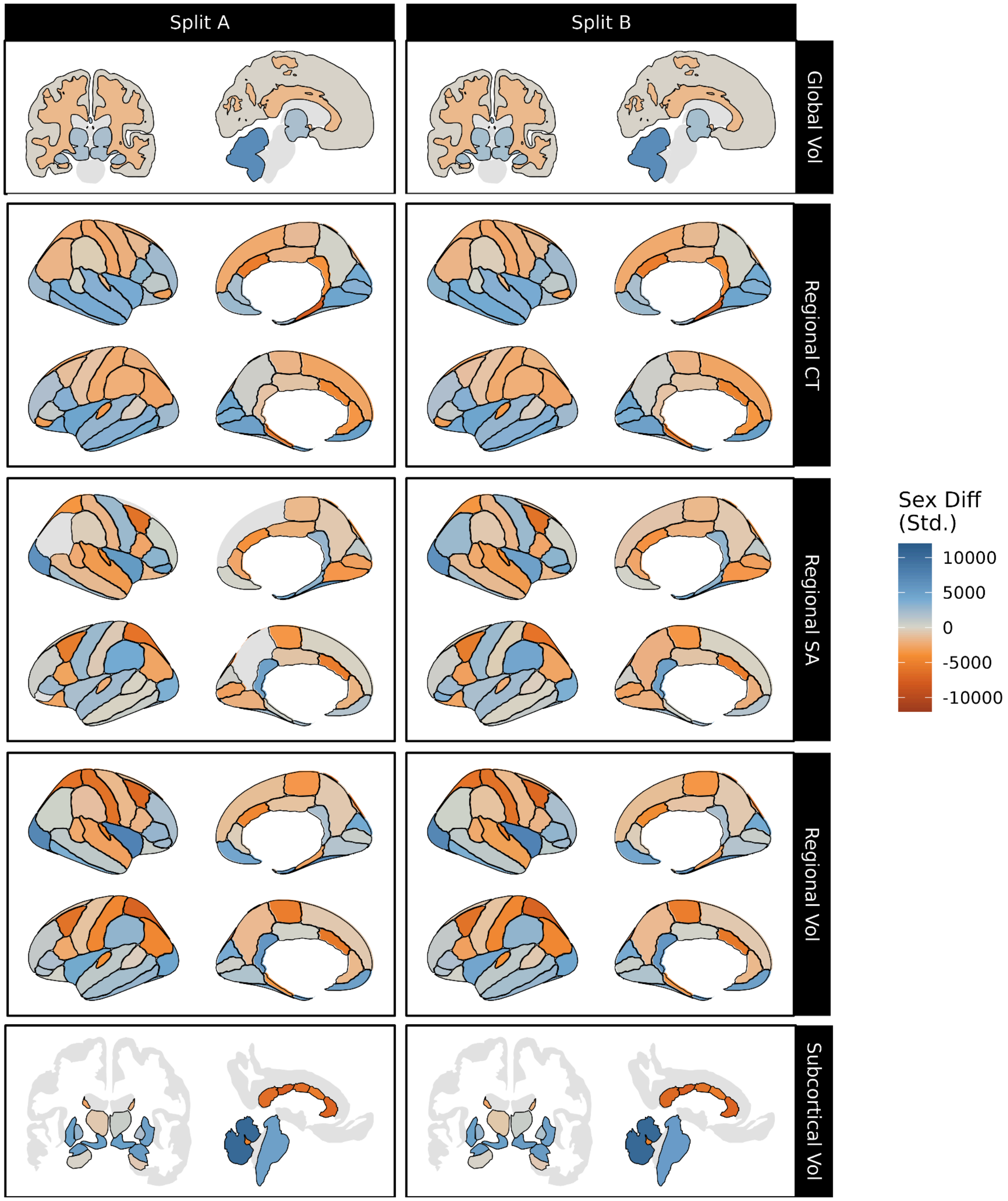


***Figure S36. Sex differences in IDPs' medians when weighting by scan quality.*** *Sex bias in IDP median trajectory derived from models controlling for total size and weighting by scan quality, with orange indicating female bias and blue indicating male bias. Predicted centiles for each sex were standardized by scaling relative to each IDP's standard deviation before subtracting predicted female median centile from the predicted male median centile at each age; overall sex differences were operationalized as the integral of the resulting sex difference trajectory, normalized to a 100-year lifespan (see Methods). Black outlines indicate significant sex effects at p <0.05, using false discovery rate (FDR) correction for multiple comparisons. Abbrv: CT, cortical thickness; SA, surface area; Vol, volume.*

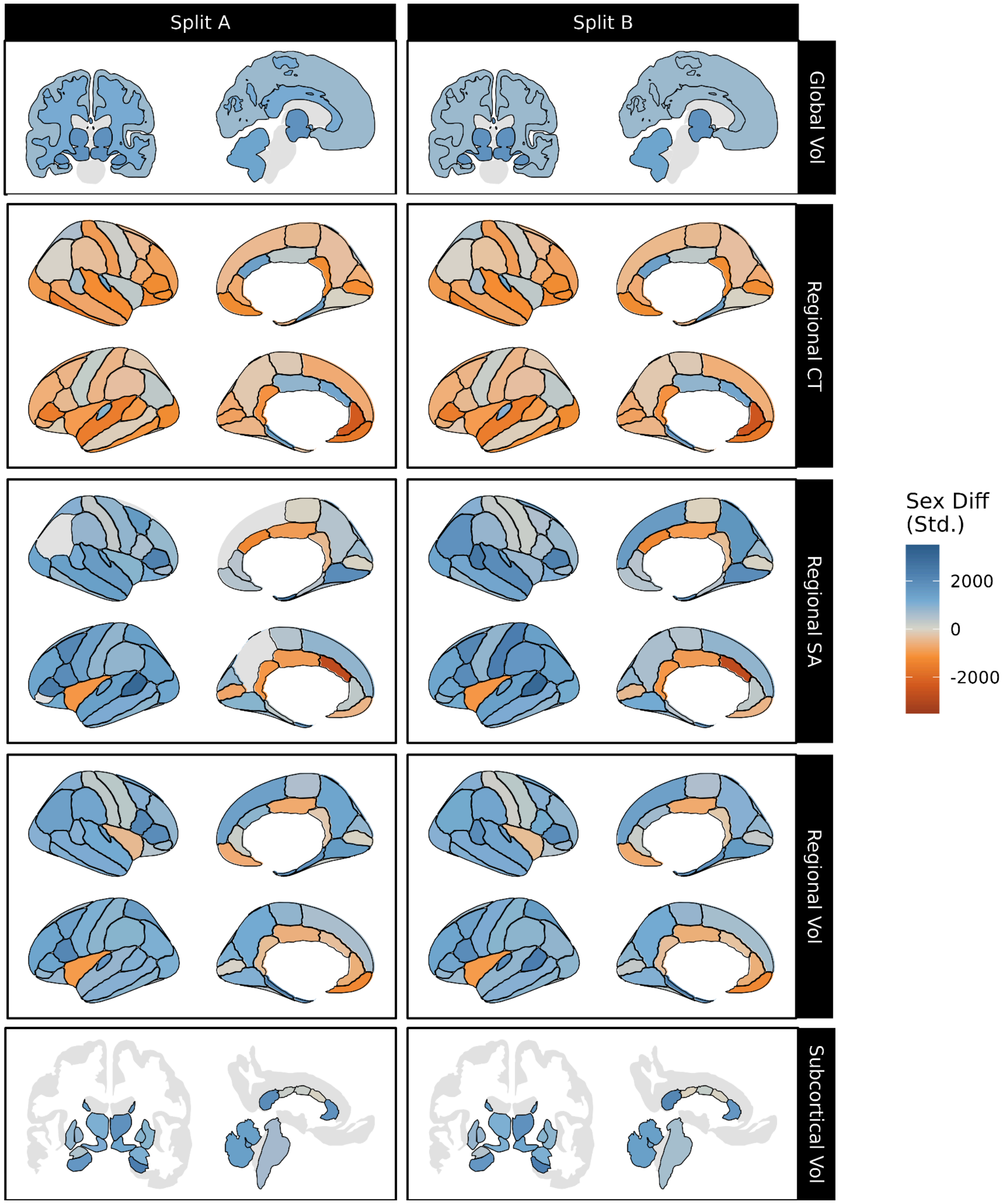


***Figure S37. Standardized sex differences in IDPs' variabilities when weighting by scan quality.*** *Sex bias in IDP variability controlling for total size and weighted by scan quality, with orange indicating female bias and blue indicating male bias. Predicted centiles for each sex were standardized by scaling relative to each IDP's standard deviation before calculating the natural log of the ratio of males' over females' coefficient of variation (i.e. log(male sigma/female sigma)); overall sex differences were operationalized as the integral of the resulting sex difference trajectory, normalized to a 100-year lifespan (see Methods). Black outlines indicate significant sex effects at p <0.05, using false discovery rate (FDR) correction for multiple comparisons. Abbrv: Std., standardized; CT, cortical thickness; SA, surface area; Vol, volume.*

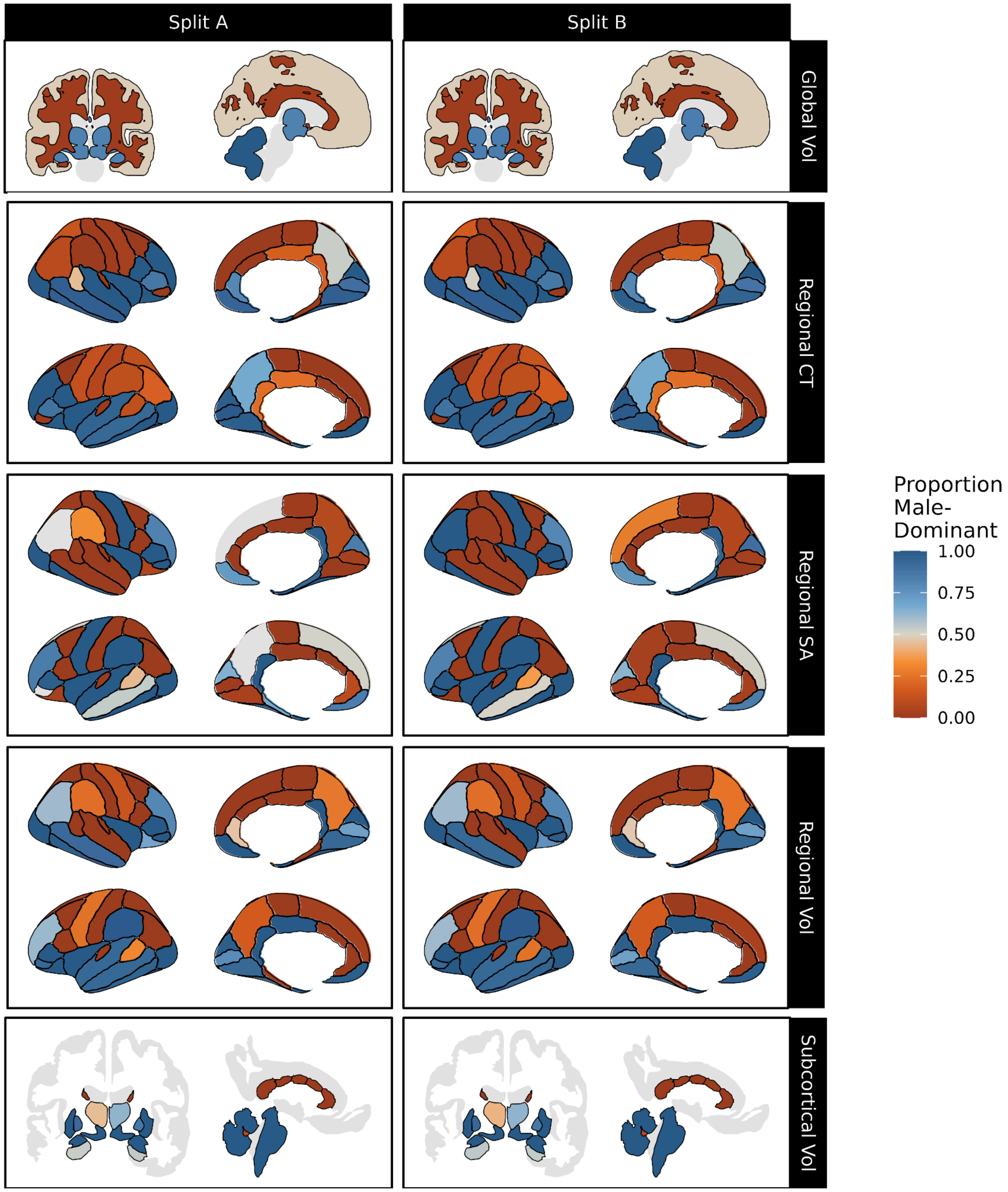


***Figure S38. Proportion of the lifespan an IDP's median is biased towards males when weighting by scan quality.*** *Blue indicates more of the lifespan being male-biased (i.e. larger in males), while orange indicates a longer time spent female-biased. Black outlines indicate significant sex effects at p <0.05, using false discovery rate (FDR) correction for multiple comparisons. Abbrv: CT, cortical thickness; SA, surface area; Vol, volume.*

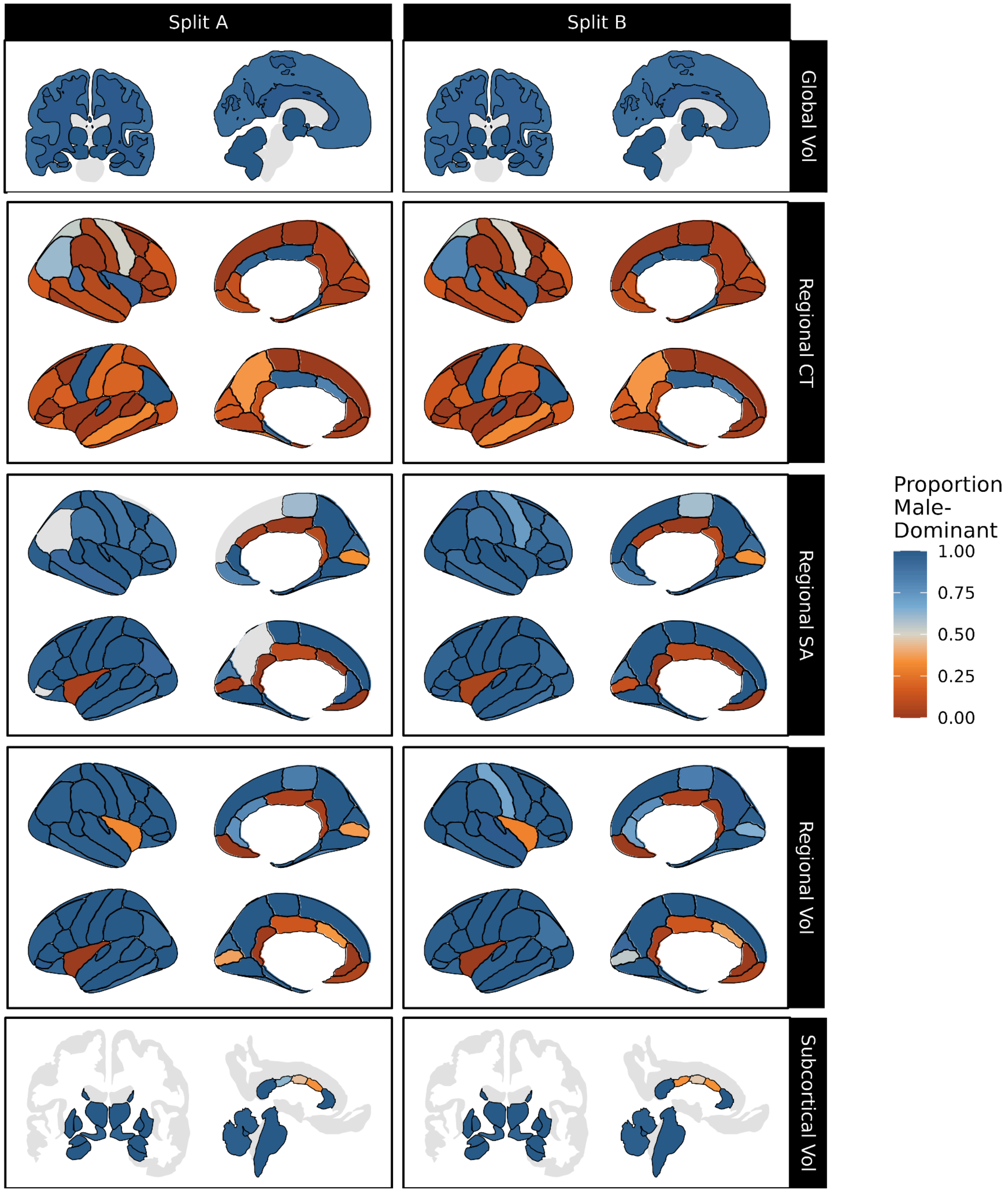


***Figure S39. Proportion of the lifespan an IDP's variability is biased towards males when weighting by scan quality.*** *Blue indicates more of the lifespan being male-biased (i.e. more variable in males), while orange indicates a longer time spent female-biased. Black outlines indicate significant sex effects at p <0.05, using false discovery rate (FDR) correction for multiple comparisons. Abbrv: CT, cortical thickness; SA, surface area; Vol, volume.*

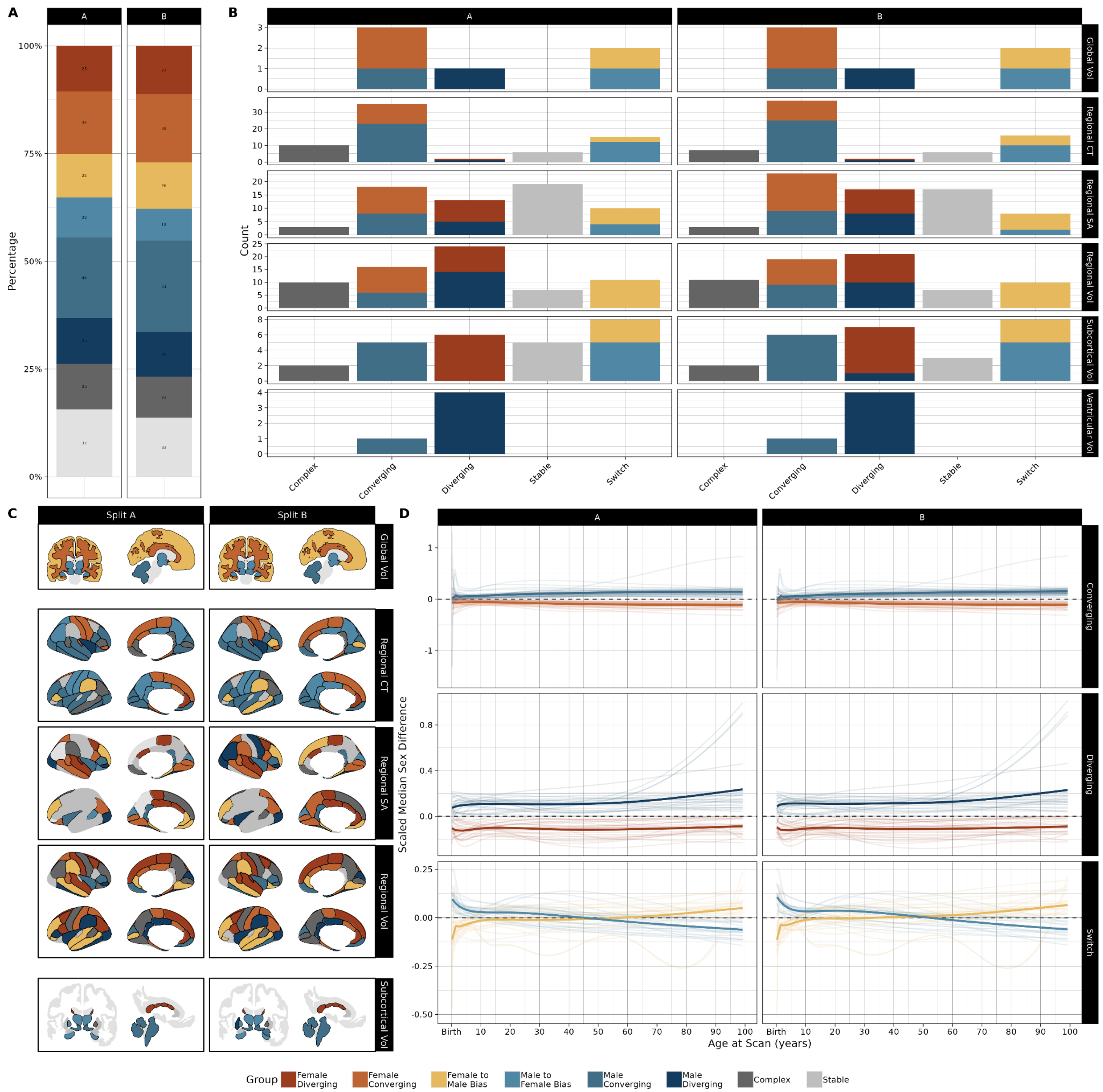


***Figure S40. IDP groupings by*** **a priori** ***classes of median sex difference trajectories when weighting by scan quality. (A)*** *Percentage of IDPs belonging to each group, as determined by the trajectory of the IDPs' median sex difference over the lifespan, in each split-half. Groups are defined as follows: female diverging, where consistently female-biased features become more biased over the lifespan; female converging, where female bias decreases over time; female bias to male bias, where IDPs that are female-biased early in life become male-biased later; male bias to female bias, where bias changes from male to female with age; male converging, where male bias decreases with age; male diverging, where male bias increases with age; stable, where sex biases do not significantly vary with age (as per likelihood ratio tests); complex, for significantly age-varying sex biases that do not fit any previously defined groups.* ***(B)*** *Number of IDPs belonging to each group, subdivided by phenotype class.* ***(C)*** *IDPs colored by group.* ***(D)*** *Summarized sex bias trajectories (dark*

*lines) within each group. Summaries for each group were created by standardizing sex-difference trajectories within each IDP (translucent lines) and fitting a simple generalized additive model.*

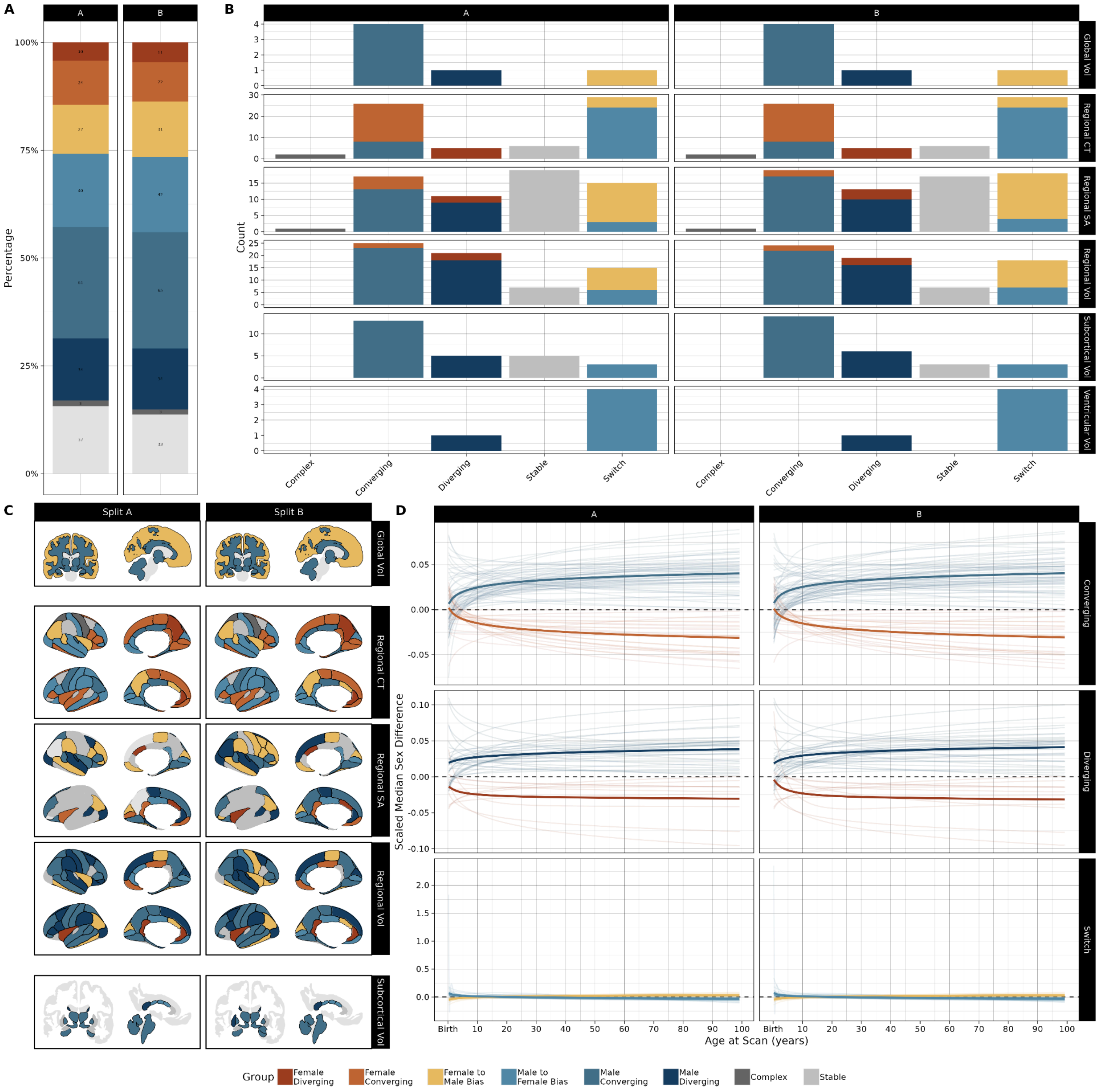


***Figure S41. IDP groupings by*** **a priori** ***classes of sex difference trajectories in variability when weighting by scan quality. (A)*** *Percentage of IDPs belonging to each group, as determined by the trajectory of the IDPs' sex difference in variability over the lifespan, in each split-half. Groups are defined as follows: female diverging, where consistently female-biased features become more biased over the lifespan; female converging, where female bias decreases over time; female bias to male bias, where IDPs that are female-biased early in life become male-biased later; male bias to female bias, where bias changes from male to female with age; male converging, where male bias decreases with age; male diverging, where male bias increases with age; stable, where sex biases do not significantly vary with age (as per likelihood ratio tests);*

*complex, for significantly age-varying sex biases that do not fit any previously defined groups.* ***(B)*** *Number of IDPs belonging to each group, subdivided by phenotype class.* ***(C)*** *IDPs colored by group.* ***(D)*** *Summarized sex bias trajectories (dark lines) within each group. Summaries for each group were created by standardizing sex-difference trajectories within each IDP (translucent lines) and fitting a simple generalized additive model.*

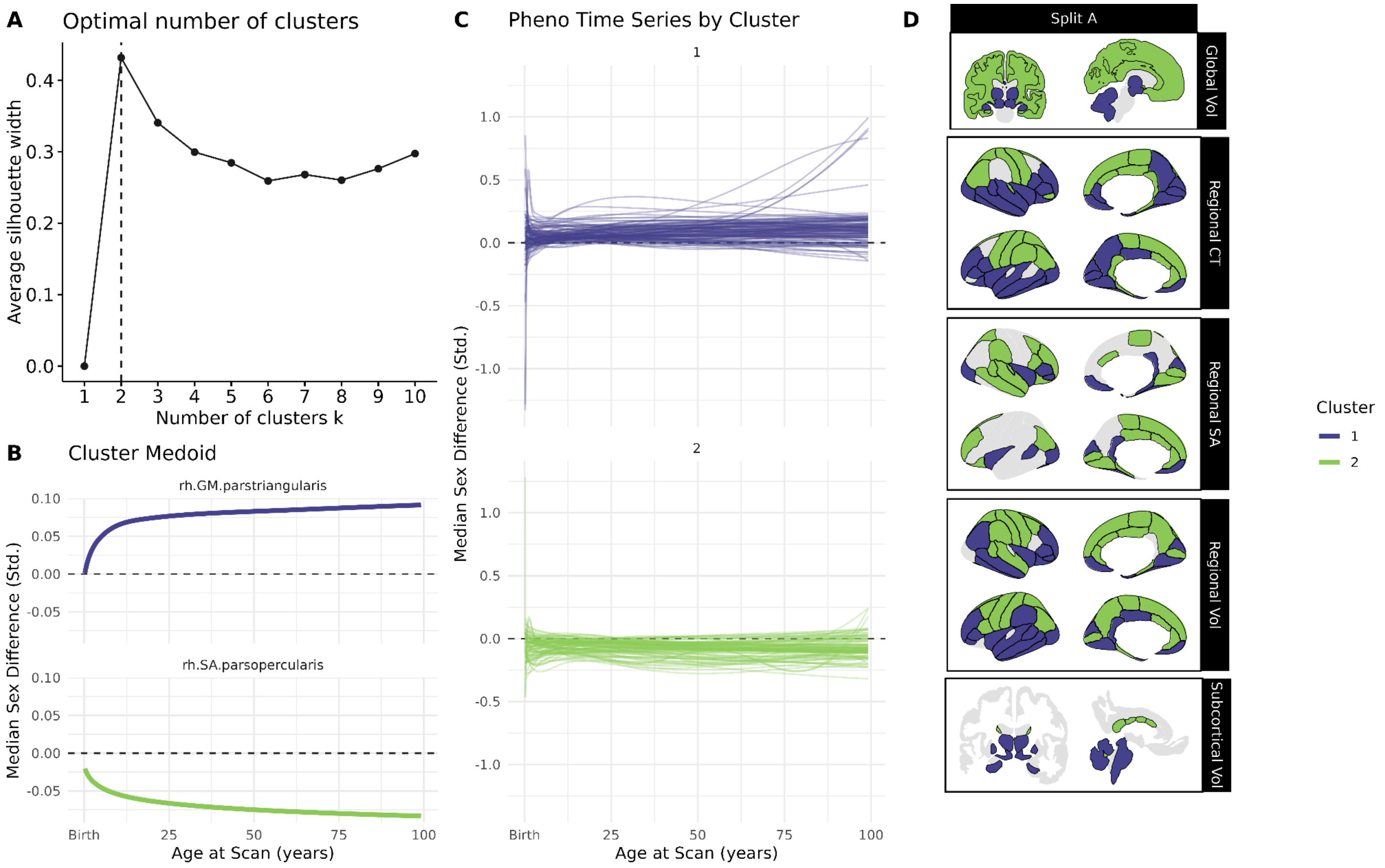


***Figure S42. Data-driven clustering of IDPs by median sex difference trajectories in test split A when weighted by scan quality. (A)*** *Plot of average silhouette width by number of k-medoid clusters, indicating a two-cluster solution.* ***(B)*** *Scaled median sex bias trajectories of cluster medoids for 2-cluster solution, corresponding to increasing male bias (Cluster 1, purple) and increasing female bias (Cluster 2, green)* ***(C)*** *Scaled median sex bias trajectories of each IDP, grouped by cluster. (D) IDPs visualized in neuroanatomical context, colored by sex bias trajectory cluster. All IDPs included in k-medoid clustering protocol had significantly age-varying sex difference trajectories, as indicated by our FDR corrected likelihood ratio tests (black outlines). Abbrv: CT, cortical thickness; SA, surface area; Vol, volume; Std., standardized*

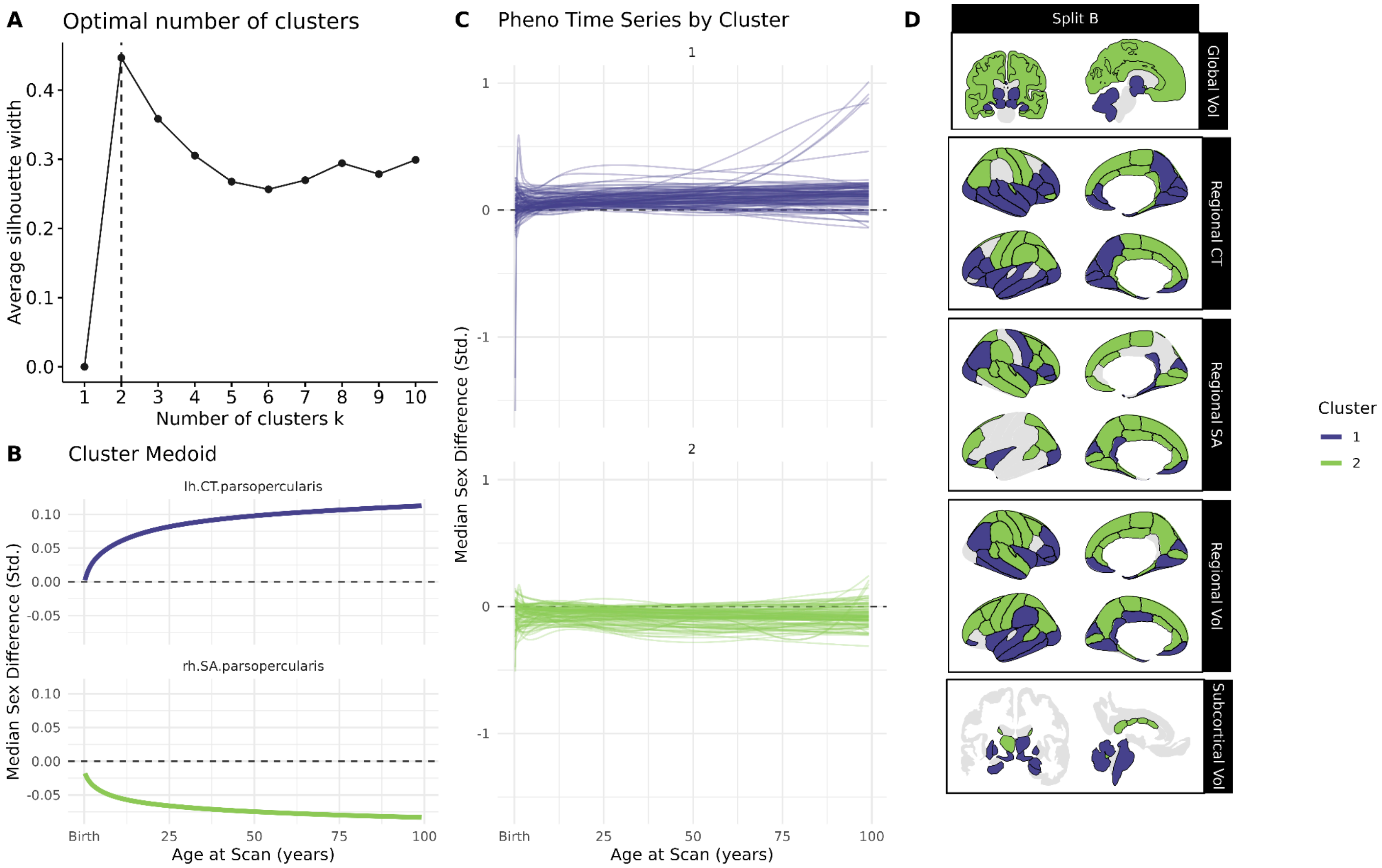


***Figure S43. Data-driven clustering of IDPs by median sex difference trajectories in test split B when weighted by scan quality. (A)*** *Plot of average silhouette width by number of k-medoid clusters, indicating a two-cluster solution.* ***(B)*** *Scaled median sex bias trajectories of cluster medoids for 2-cluster solution, corresponding to increasing male bias (Cluster 1, purple) and increasing female bias (Cluster 2, green)* ***(C)*** *Scaled median sex bias trajectories of each IDP, grouped by cluster. (D) IDPs visualized in neuroanatomical context, colored by sex bias trajectory cluster. All IDPs included in k-medoid clustering protocol had significantly age-varying sex difference trajectories, as indicated by our FDR corrected likelihood ratio tests (black outlines). Abbrv: CT, cortical thickness; SA, surface area; Vol, volume; Std., standardized*

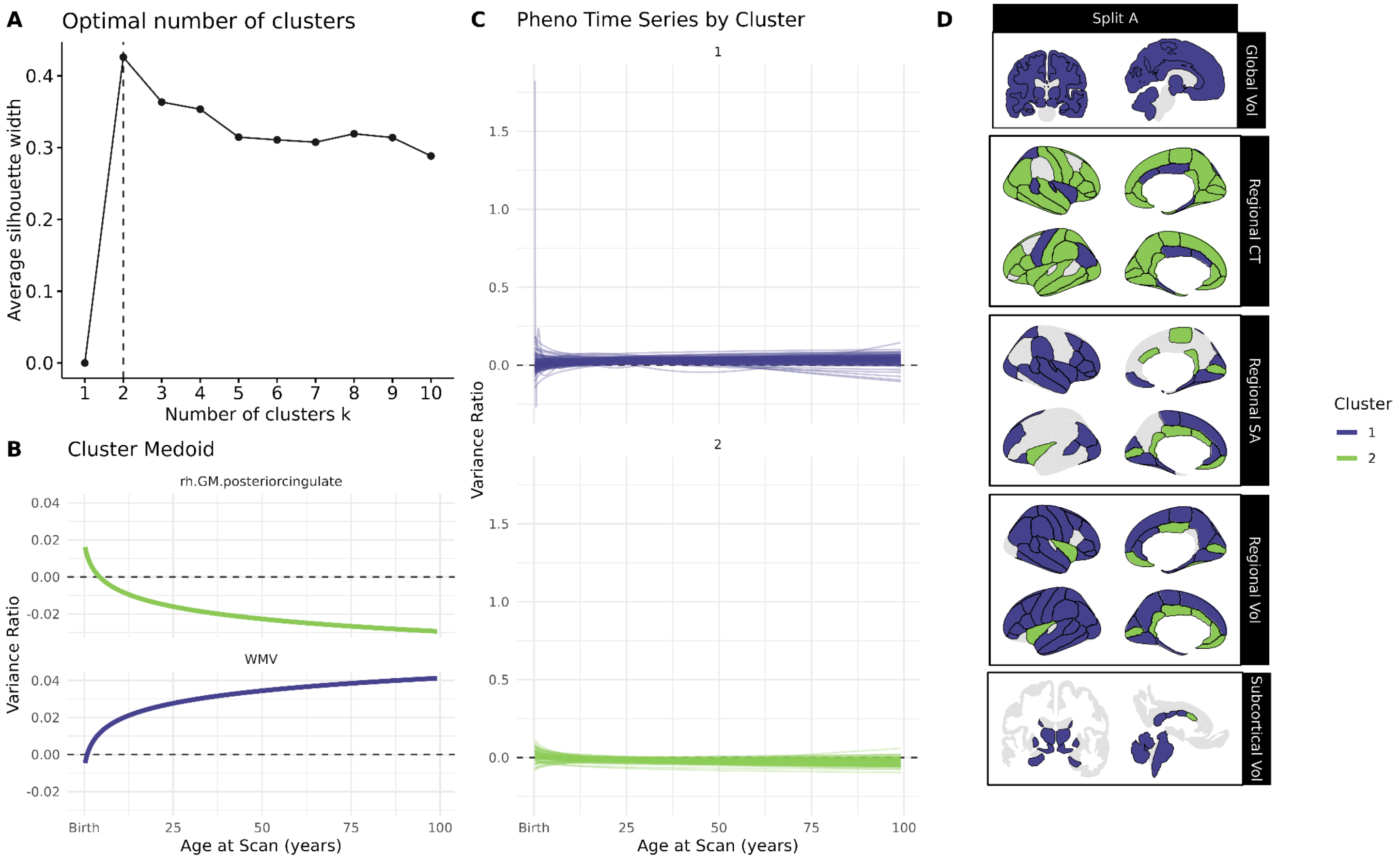


***Figure S44. Data-driven clustering of IDPs by sex difference trajectories in variability in test split A when weighting by scan quality. (A)*** *Plot of average silhouette width by number of k-medoid clusters, indicating a two-cluster solution.* ***(B)*** *Scaled sex bias trajectories in variability of cluster medoids for 2-cluster solution, corresponding to increasing male bias (Cluster 1, purple) and increasing female bias (Cluster 2, green)* ***(C)*** *Scaled variability sex bias trajectories of each IDP, grouped by cluster. (D) IDPs visualized in neuroanatomical context, colored by sex bias trajectory cluster. All IDPs included in k-medoid clustering protocol had significantly age-varying sex difference trajectories, as indicated by our FDR corrected likelihood ratio tests (black outlines). Abbrv: CT, cortical thickness; SA, surface area; Vol, volume; Std., standardized*

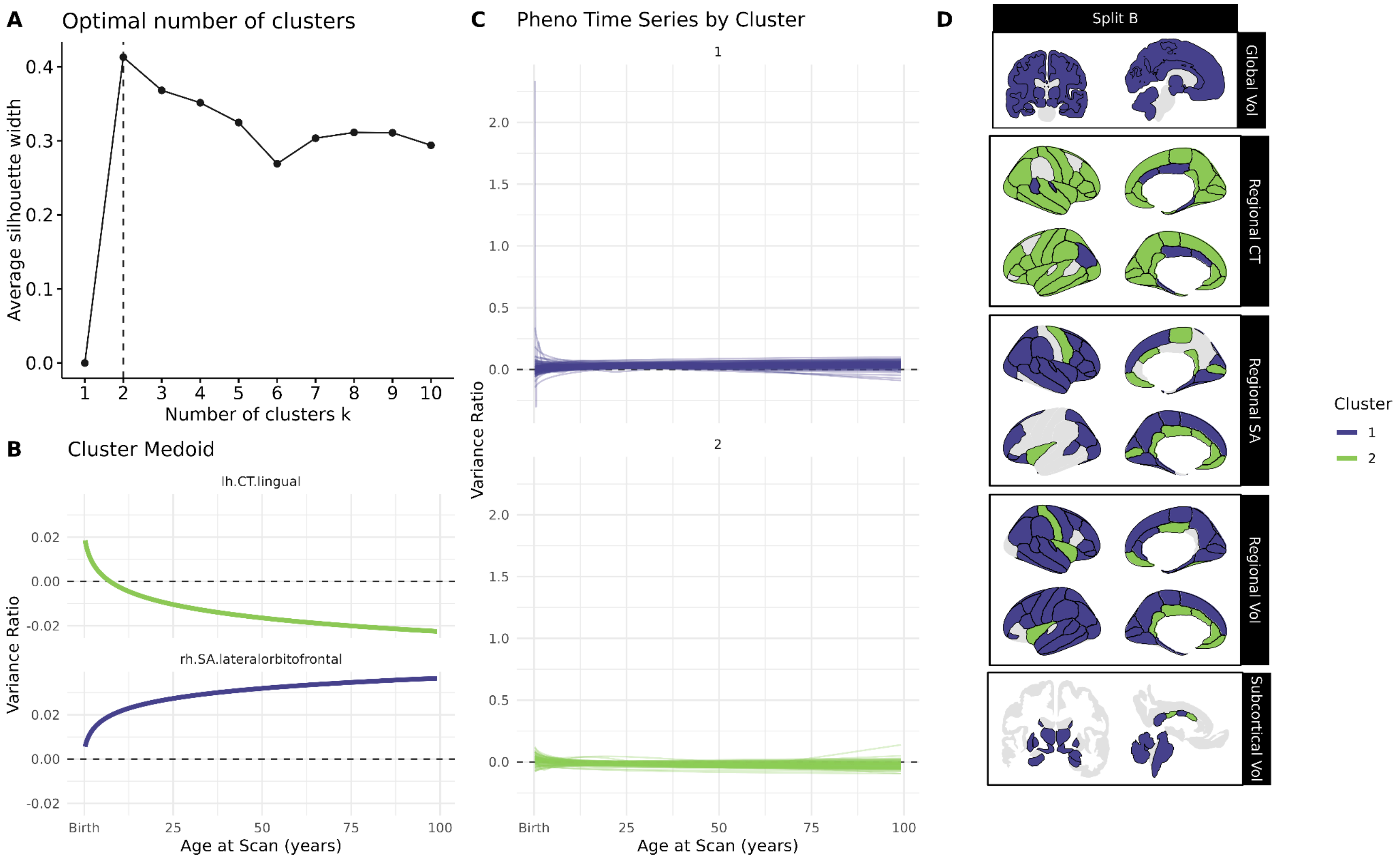


***Figure S45. Data-driven clustering of IDPs by sex difference trajectories in variability in test split B when weighted by scan quality. (A)*** *Plot of average silhouette width by number of k-medoid clusters, indicating a two-cluster solution.* ***(B)*** *Scaled sex bias trajectories in variability of cluster medoids for 2-cluster solution, corresponding to increasing male bias (Cluster 1, purple) and increasing female bias (Cluster 2, green)* ***(C)*** *Scaled variability sex bias trajectories of each IDP, grouped by cluster. (D) IDPs visualized in neuroanatomical context, colored by sex bias trajectory cluster. All IDPs included in k-medoid clustering protocol had significantly age-varying sex difference trajectories, as indicated by our FDR corrected likelihood ratio tests (black outlines). Abbrv: CT, cortical thickness; SA, surface area; Vol, volume; Std., standardized*

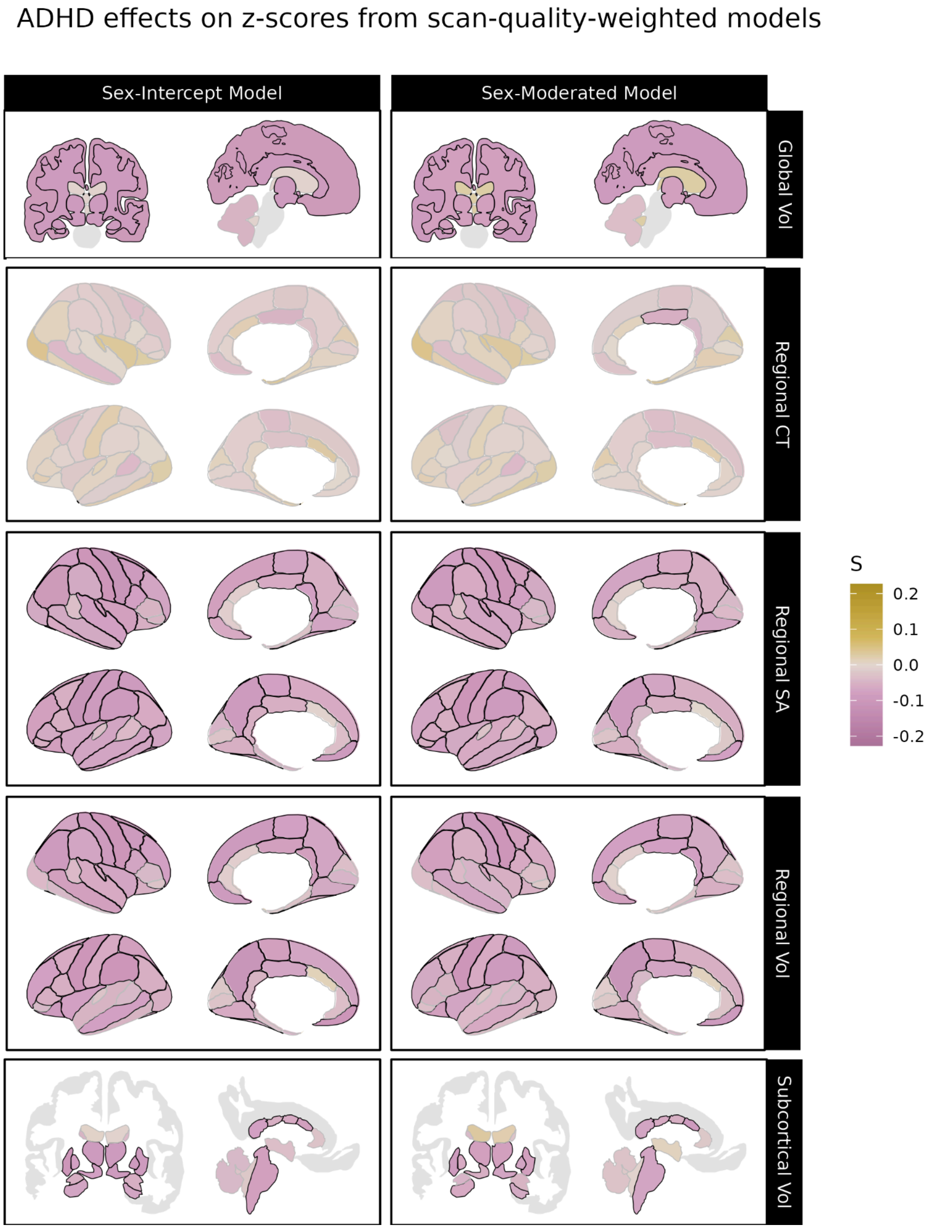


***Figure S46. Significant differences associated with ADHD in centile z-scores derived from sex-moderated and sex-intercept brain charts weighting by scan quality.*** *Differences in centile z-scores in individuals with attention-deficit/hyperactivity disorder (ADHD) relative to site-matched controls. The left panel displays differences in centile z-scores derived from sex-intercept models (i.e. models keeping sex differences in IDPs constant across the lifespan) while the right panel displays differences in centile z-scores derived from sex-moderated models, which allow age-by-sex effects. Gold indicates cases tend to have higher centile z-scores than controls for a given IDP, while purple indicates cases tend to have smaller centile z-scores. Black outlines indicate significant effects at p <0.05, FDR-corrected in each model. Abbrv: CT, cortical thickness; SA, surface area; Vol, volume.*

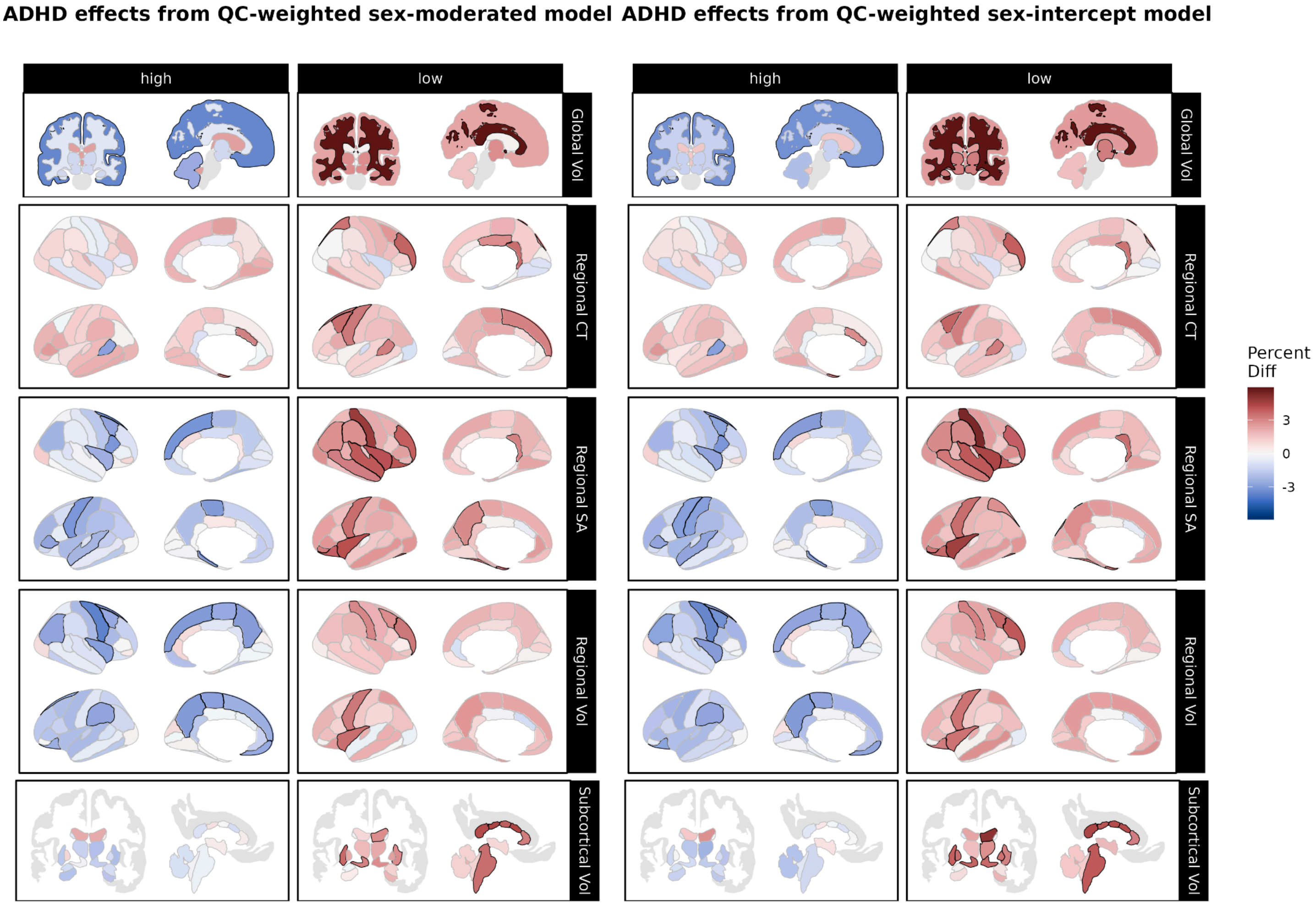


***Figure S47. Over- and under-representation of ADHD in extreme centiles according to sex-moderated and sex-intercept brain charts weighted by scan quality.*** *Using centile scores derived from our sex-moderated (left panel) and sex-intercept (right panel) brain charts, we compared the proportions of individuals with attention-deficit/hyperactivity disorder (ADHD) to the proportion of site-matched controls with centiles above the 95th percentile ("high", left column) and below the 5th percentile ("low", right column). Reds indicate cases are over-represented relative to controls, while greens indicate that cases are under-represented relative to controls. Black outlines indicate significant effects at p <0.05, FDR-corrected within each brain chart model. Abbrv: ADHD, attention-deficit/hyperactivity disorder; CT, cortical thickness; SA, surface area; Vol, volume.*

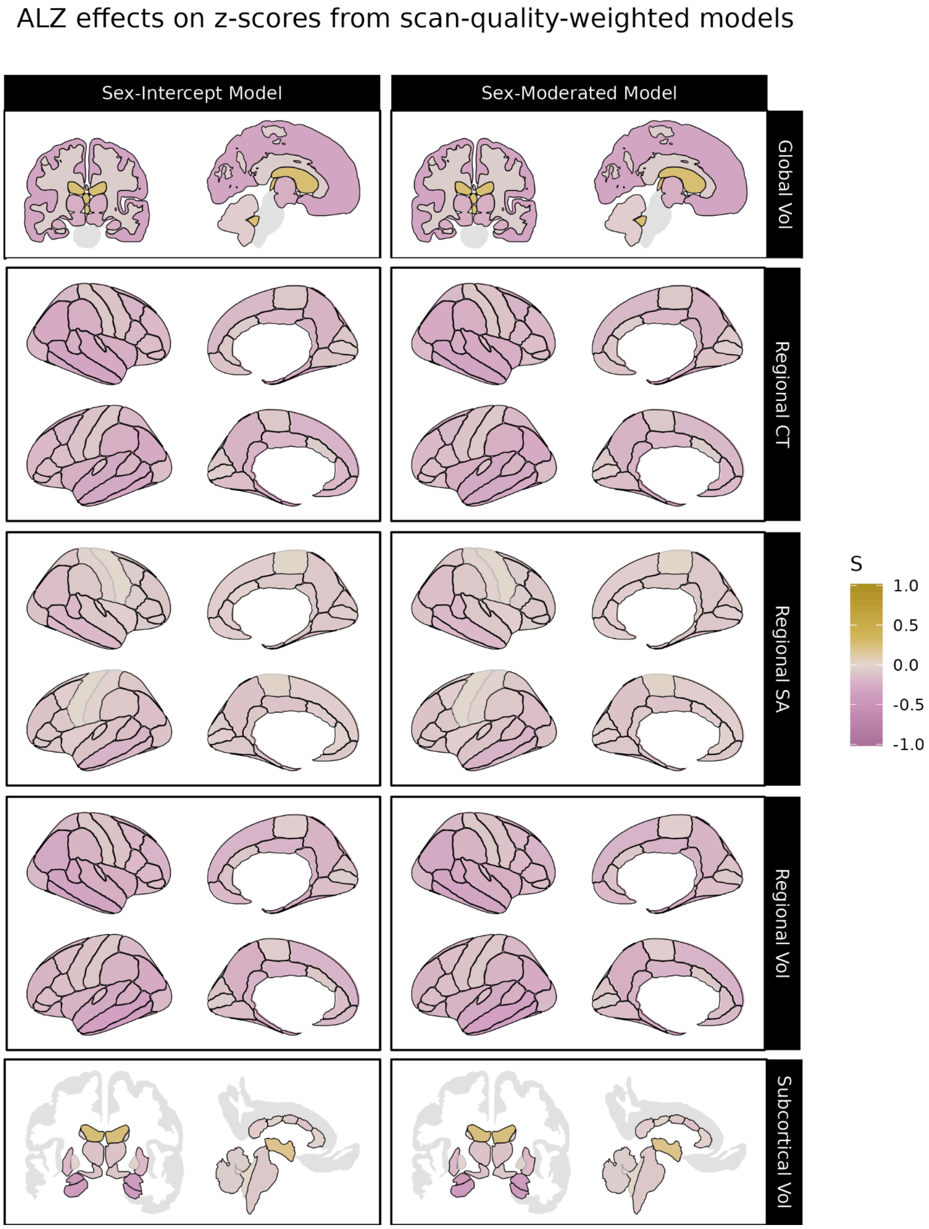


***Figure S48. Significant differences associated with Alzheimer's disease in centile z-scores derived from sex-moderated and sex-intercept brain charts weighted by scan quality.*** *Differences in centile z-scores in individuals with Alzheimer's disease relative to site-matched controls. The left panel displays differences in centile z-scores derived from sex-intercept models (i.e. models keeping sex differences in IDPs constant across the lifespan) while the right panel displays differences in centile z-scores derived from sex-moderated models, which allow age-by-sex effects. Gold indicates cases tend to have higher centile z-scores than controls for a given IDP, while purple indicates cases tend to have smaller centile z-scores. Black outlines indicate significant effects at p <0.05, FDR-corrected in each model. Abbrv: ALZ, Alzheimer's disease; CT, cortical thickness; SA, surface area; Vol, volume.*

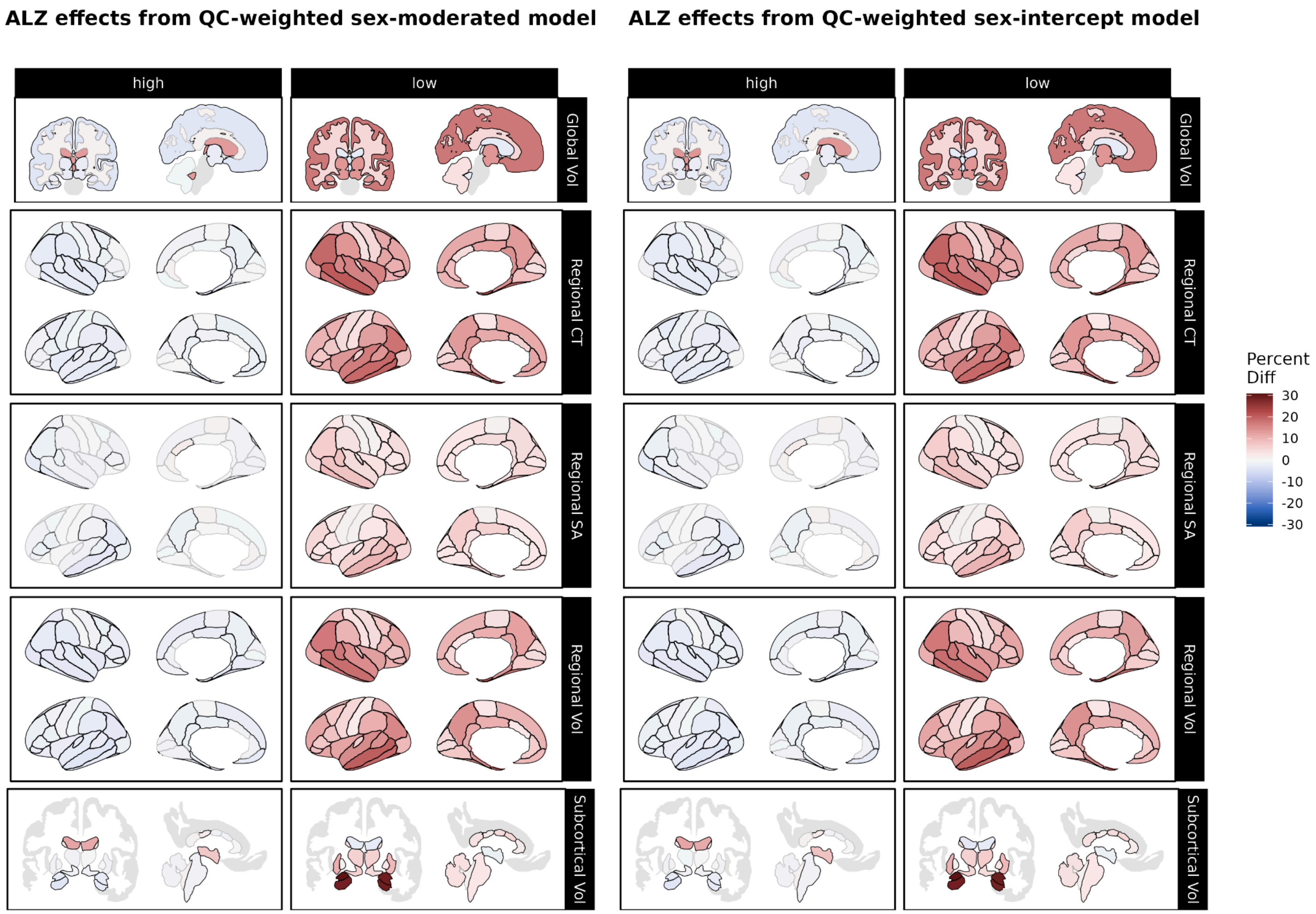


***Figure S49. Over- and under-representation of Alzheimer's disease in extreme centiles according to sex-moderated and sex-intercept brain charts weighted by scan quality.*** *Using centile scores derived from our sex-moderated (left panel) and sex-intercept (right panel) brain charts, we compared the proportions of individuals with Alzheimer's disease (ALZ) to the proportion of site-matched controls with centiles above the 95th percentile ("high", left column) and below the 5th percentile ("low", right column). Reds indicate cases are over-represented relative to controls, while greens indicate that cases are under-represented relative to controls. Black outlines indicate significant effects at p <0.05, FDR-corrected within each brain chart model. Abbrv: ALZ, Alzheimer's disease; CT, cortical thickness; SA, surface area; Vol, volume.*

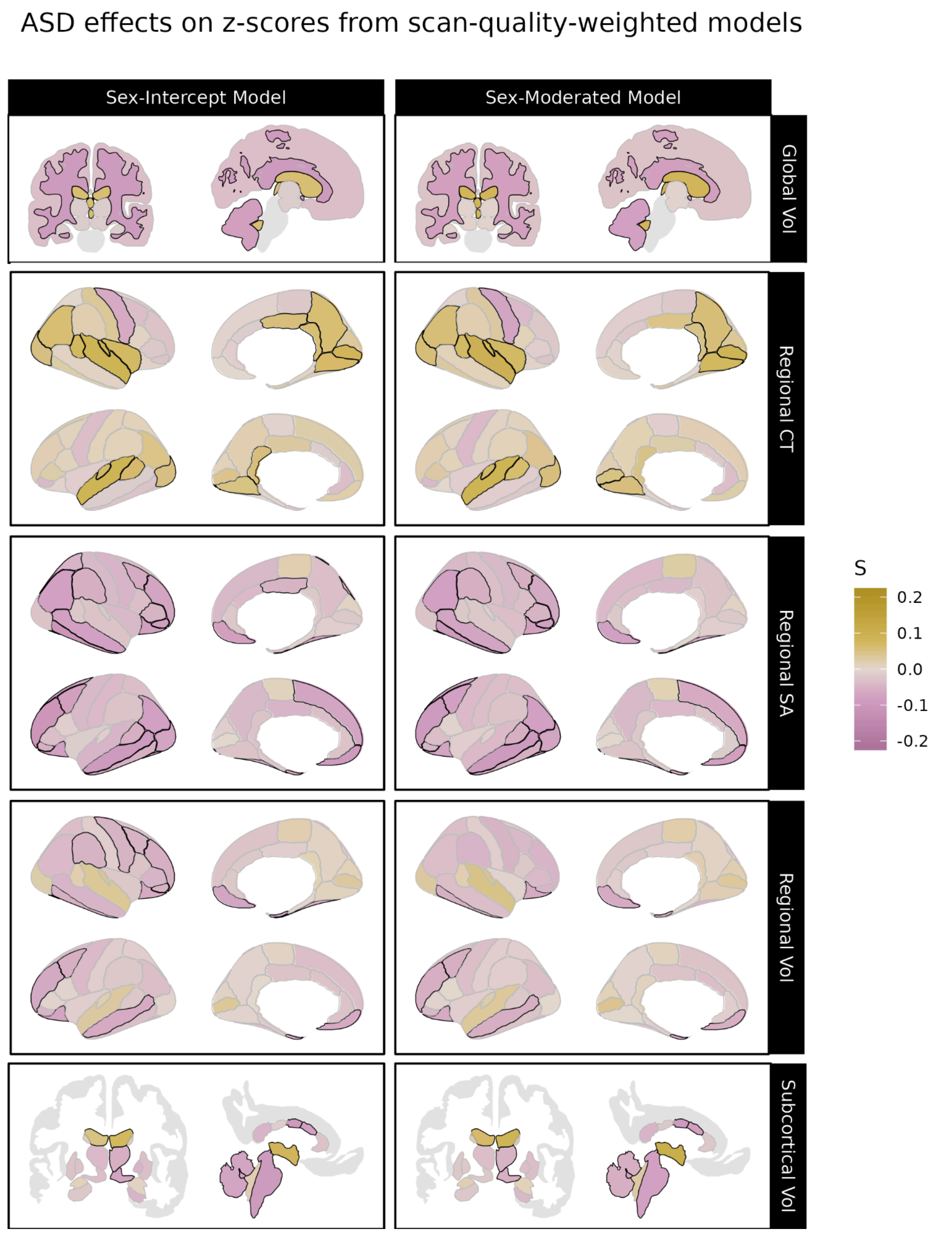


***Figure S50. Significant differences associated with autism spectrum disorder in centile z-scores derived from sex-moderated and sex-intercept brain charts weighted by scan quality.*** *Differences in centile z-scores in individuals with autism spectrum disorder (ASD) relative to site-matched controls. The left panel displays differences in centile z-scores derived from sex-intercept models (i.e. models keeping sex differences in IDPs constant across the lifespan) while the right panel displays differences in centile z-scores derived from sex-moderated models, which allow age-by-sex effects. Gold indicates cases tend to have higher centile z-scores than controls for a given IDP, while purple indicates cases tend to have smaller centile z-scores. Black outlines indicate significant effects at p <0.05, FDR-corrected in each model. Abbrv: ASD, autism spectrum disorder; CT, cortical thickness; SA, surface area; Vol, volume.*

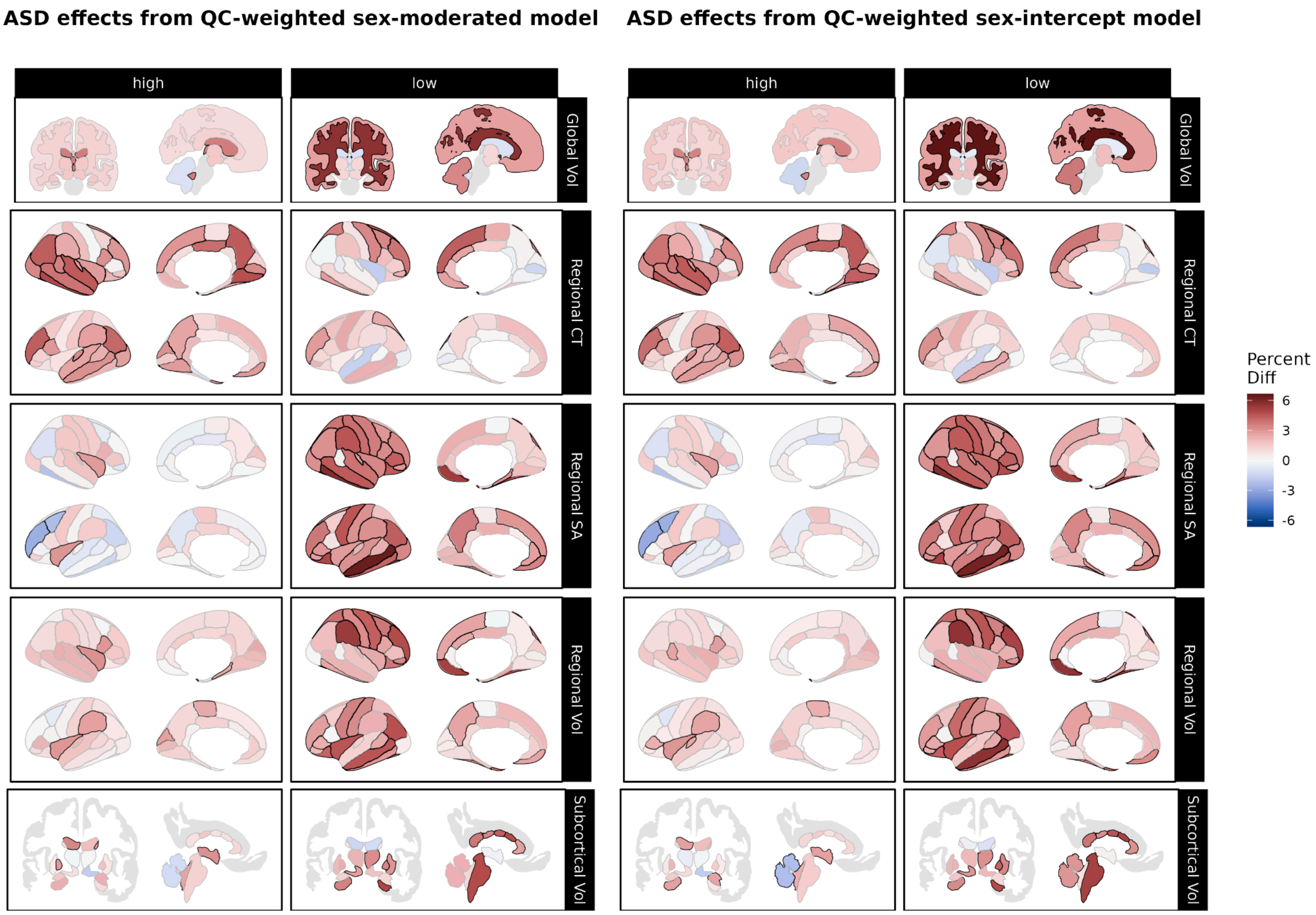


***Figure S51. Over- and under-representation of autism spectrum disorder in extreme centiles according to sex-moderated and sex-intercept brain charts weighted by scan quality.*** *Using centile scores derived from our sex-moderated (left panel) and sex-intercept (right panel) brain charts, we compared the proportions of individuals with autism spectrum disorder (ASD) to the proportion of site-matched controls with centiles above the 95th percentile ("high", left column) and below the 5th percentile ("low", right column). Reds indicate cases are over-represented relative to controls, while greens indicate that cases are under-represented relative to controls. Black outlines indicate significant effects at p <0.05, FDR-corrected within each brain chart model. Abbrv: ASD, autism spectrum disorder; CT, cortical thickness; SA, surface area; Vol, volume.*

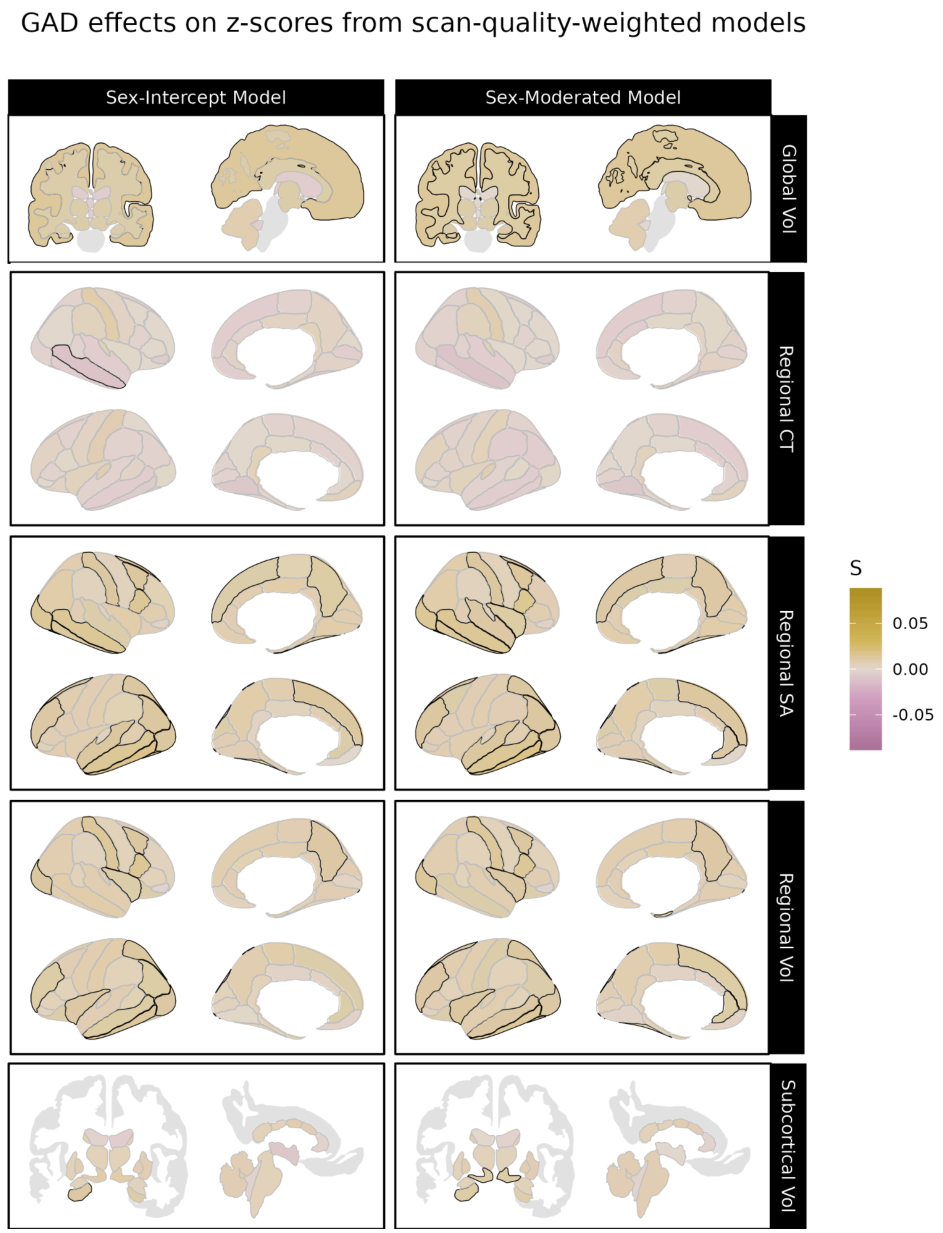


***Figure S52. Significant differences associated with generalized anxiety disorder in centile z-scores derived from sex-moderated and sex-intercept brain charts weighted by scan quality.*** *Differences in centile z-scores in individuals with generalized anxiety disorder (GAD) relative to site-matched controls. The left panel displays differences in centile z-scores derived from sex-intercept models (i.e. models keeping sex differences in IDPs constant across the lifespan) while the right panel displays differences in centile z-scores derived from sex-moderated models, which allow age-by-sex effects. Gold indicates cases tend to have higher centile z-scores than controls for a given IDP, while purple indicates cases tend to have smaller centile z-scores. Black outlines indicate significant effects at p <0.05, FDR-corrected in each model. Abbrv: GAD, generalized anxiety disorder; CT, cortical thickness; SA, surface area; Vol, volume.*

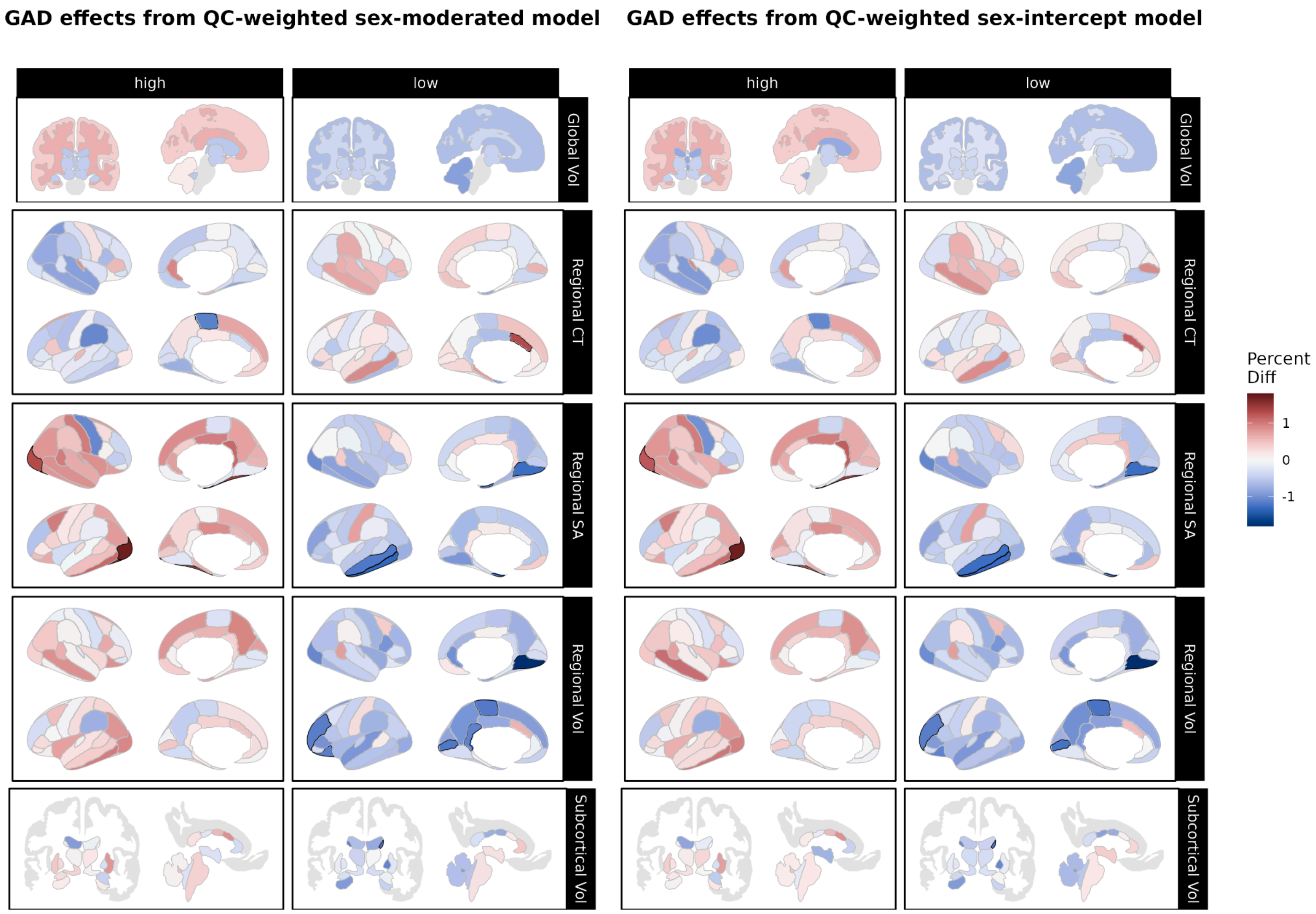


***Figure S53. Over- and under-representation of generalized anxiety disorder in extreme centiles according to sex-moderated and sex-intercept brain charts weighted by scan quality.*** *Using centile scores derived from our sex-moderated (left panel) and sex-intercept (right panel) brain charts, we compared the proportions of individuals with generalized anxiety disorder (GAD) to the proportion of site-matched controls with centiles above the 95th percentile ("high", left column) and below the 5th percentile ("low", right column). Reds indicate cases are over-represented relative to controls, while greens indicate that cases are under-represented relative to controls. Black outlines indicate significant effects at p <0.05, FDR-corrected within each brain chart model. Abbrv: GAD, generalized anxiety disorder; CT, cortical thickness; SA, surface area; Vol, volume.*

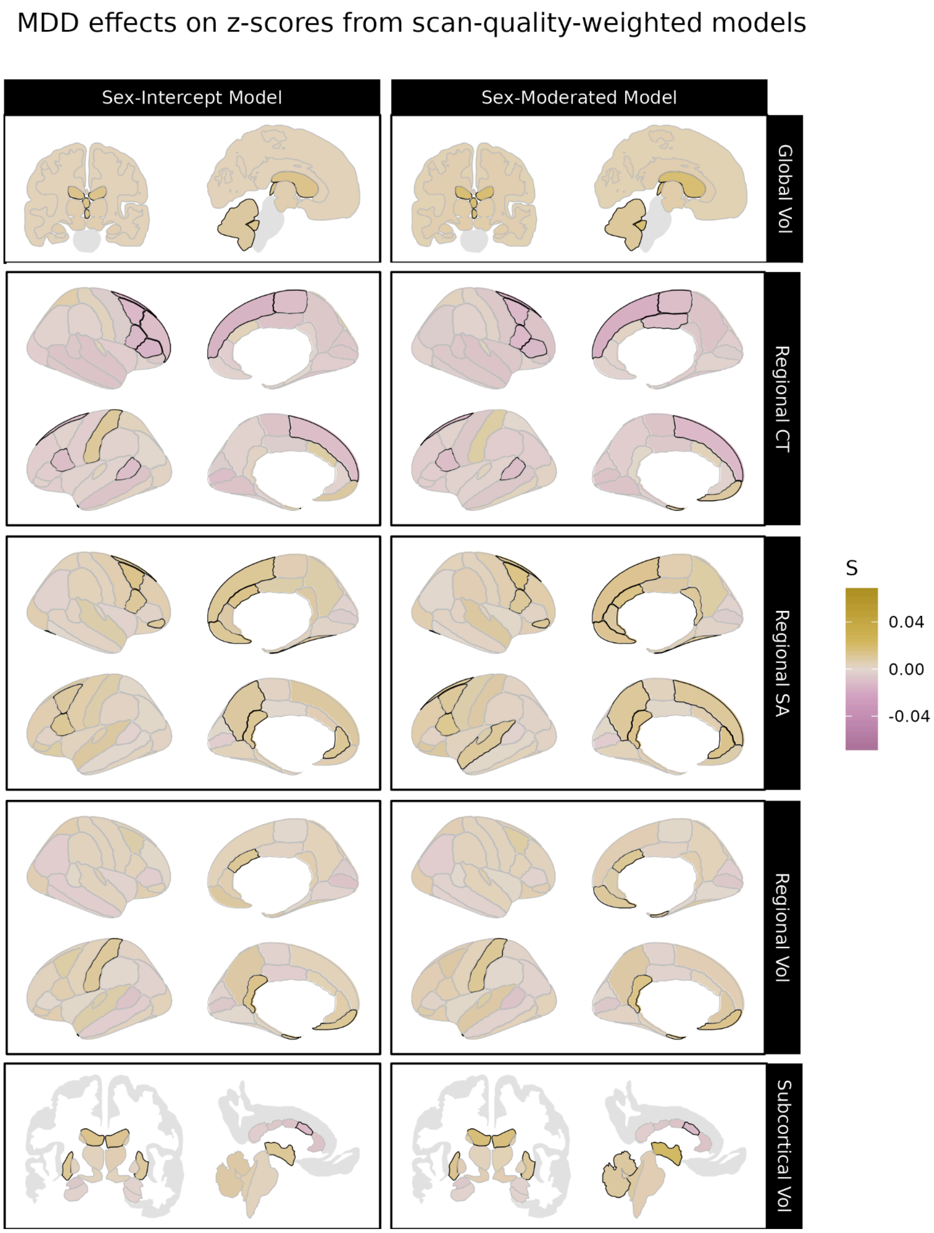


***Figure S54. Significant differences associated with major depressive disorder in centile z-scores derived from sex-moderated and sex-intercept brain charts weighted by scan quality.*** *Differences in centile z-scores in individuals with major depressive disorder (MDD) relative to site-matched controls. The left panel displays differences in centile z-scores derived from sex-intercept models (i.e. models keeping sex differences in IDPs constant across the lifespan) while the right panel displays differences in centile z-scores derived from sex-moderated models, which allow age-by-sex effects. Gold indicates cases tend to have higher centile z-scores than controls for a given IDP, while purple indicates cases tend to have smaller centile z-scores. Black outlines indicate significant effects at p <0.05, FDR-corrected in each model. Abbrv: MDD, major depressive disorder; CT, cortical thickness; SA, surface area; Vol, volume.*

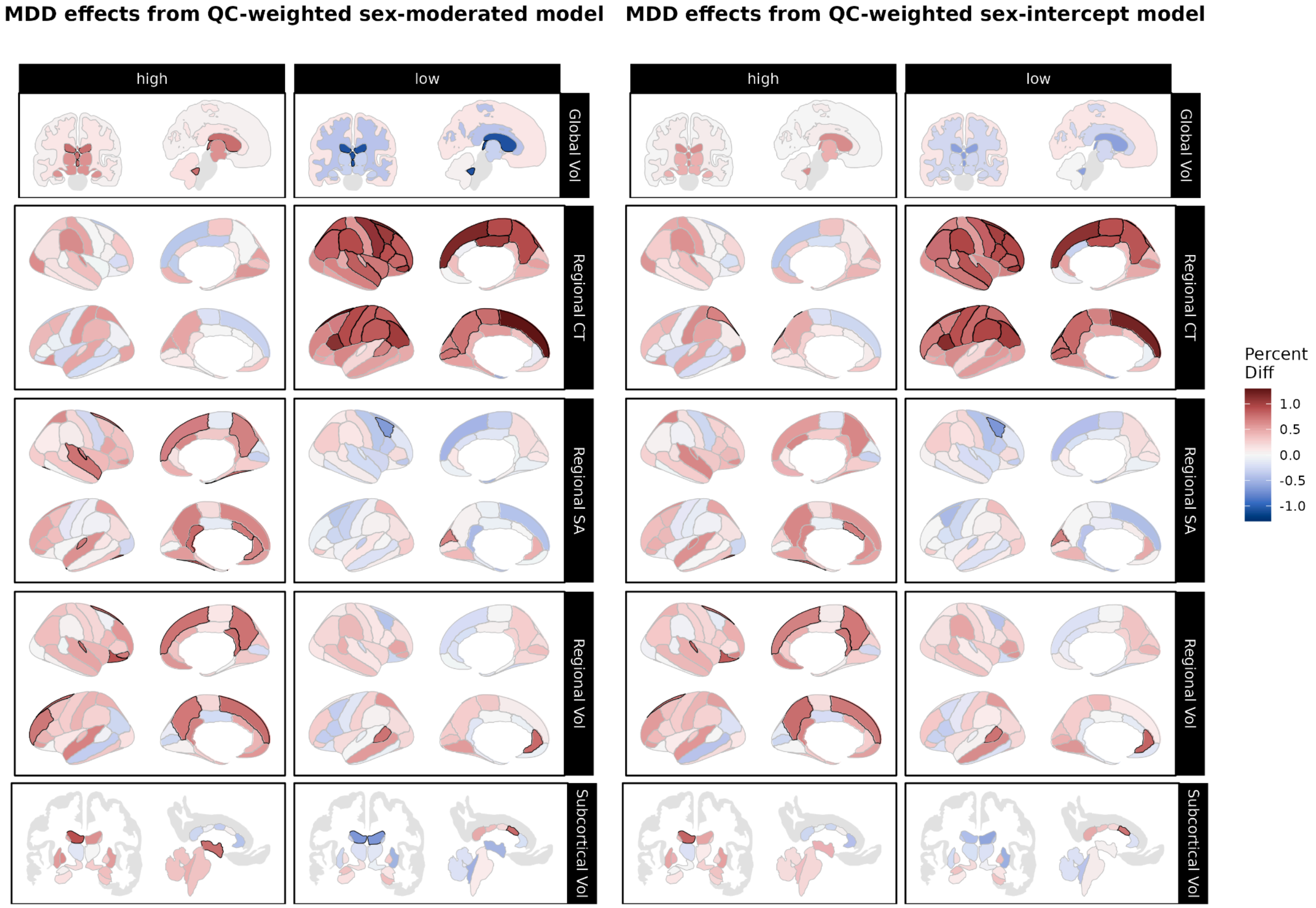


***Figure S55. Over- and under-representation of major depressive disorder in extreme centiles according to sex-moderated and sex-intercept brain charts weighted by scan quality.*** *Using centile scores derived from our sex-moderated (left panel) and sex-intercept (right panel) brain charts, we compared the proportions of individuals with major depressive disorder (MDD) to the proportion of site-matched controls with centiles above the 95th percentile ("high", left column) and below the 5th percentile ("low", right column). Reds indicate cases are over-represented relative to controls, while greens indicate that cases are under-represented relative to controls. Black outlines indicate significant effects at p <0.05, FDR-corrected within each brain chart model. Abbrv: MDD, major depressive disorder; CT, cortical thickness; SA, surface area; Vol, volume.*

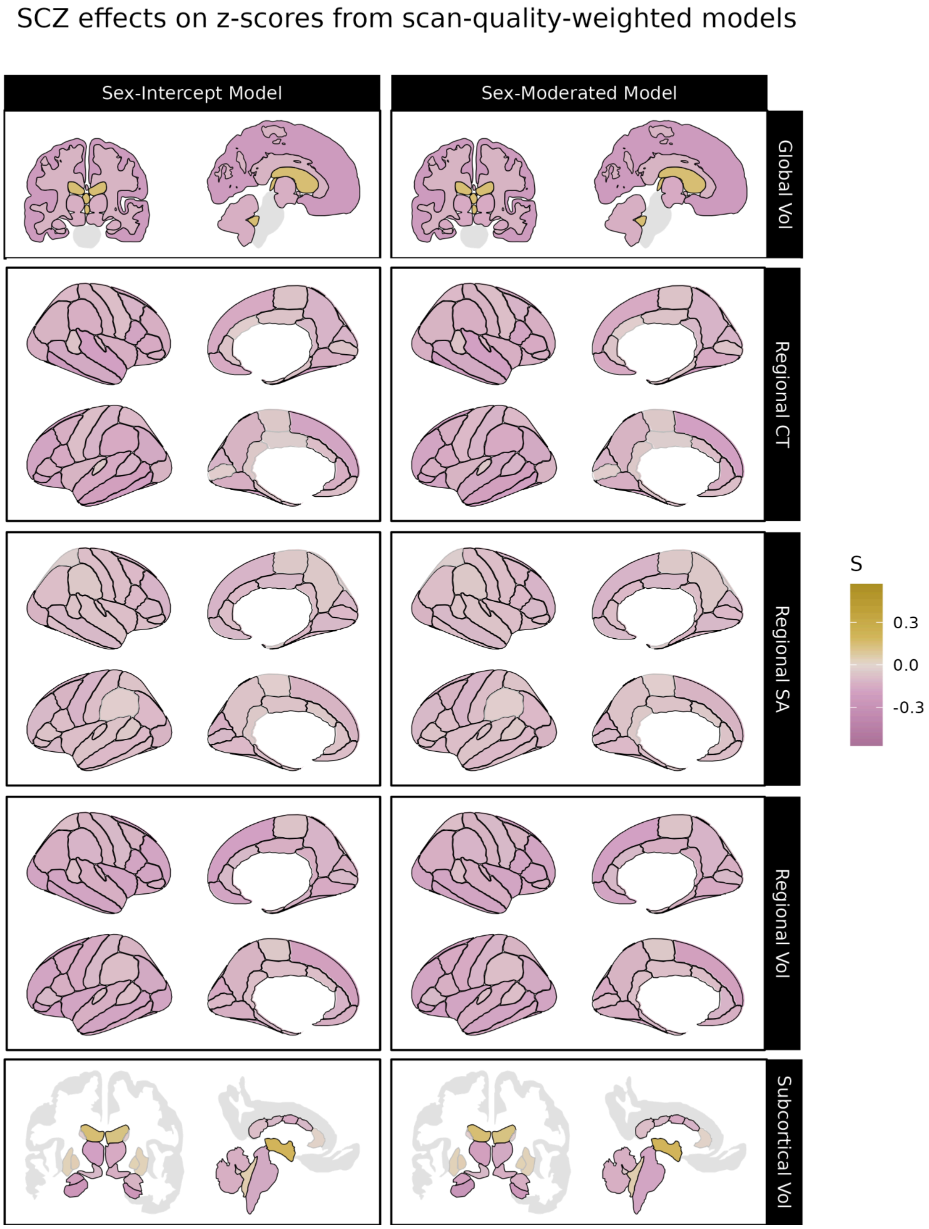


***Figure S56. Significant differences associated with schizophrenia in centile z-scores derived from sex-moderated and sex-intercept brain charts weighted by scan quality.*** *Differences in centile z-scores in individuals with schizophrenia (SCZ) relative to site-matched controls. The left panel displays differences in centile z-scores derived from sex-intercept models (i.e. models keeping sex differences in IDPs constant across the lifespan) while the right panel displays differences in centile z-scores derived from sex-moderated models, which allow age-by-sex effects. Gold indicates cases tend to have higher centile z-scores than controls for a given IDP, while purple indicates cases tend to have smaller centile z-scores. Black outlines indicate significant effects at p <0.05, FDR-corrected in each model. Abbrv: SCZ, schizophrenia; CT, cortical thickness; SA, surface area; Vol, volume.*

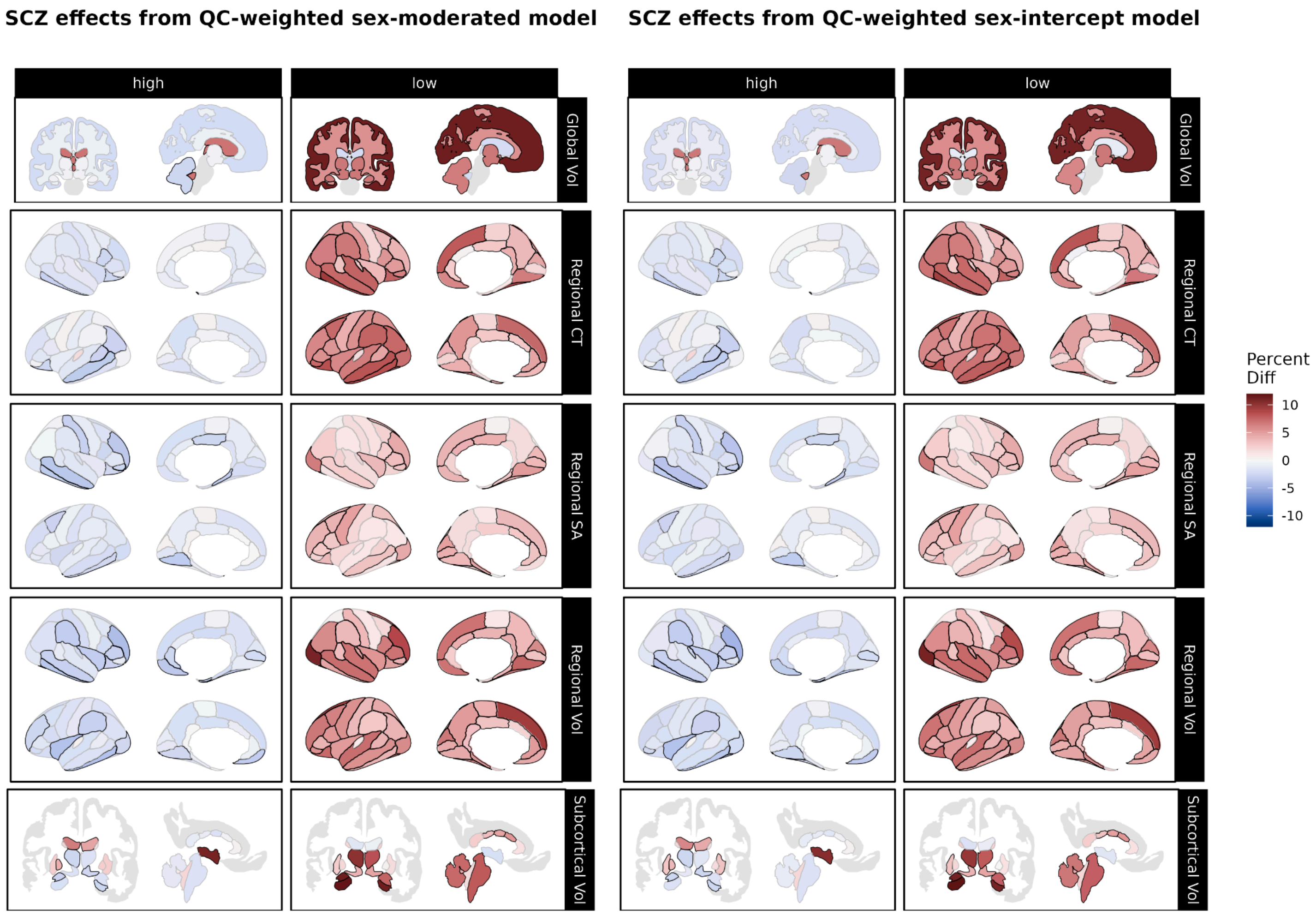


***Figure S57. Over- and under-representation of schizophrenia in extreme centiles according to sex-moderated and sex-intercept brain charts weighted by scan quality.*** *Using centile scores derived from our sex-moderated (left panel) and sex-intercept (right panel) brain charts, we compared the proportions of individuals with schizophrenia (SCZ) to the proportion of site-matched controls with centiles above the 95th percentile ("high", left column) and below the 5th percentile ("low", right column). Reds indicate cases are over-represented relative to controls, while greens indicate that cases are under-represented relative to controls. Black outlines indicate significant effects at p <0.05, FDR-corrected within each brain chart model. Abbrv: SCZ, schizophrenia; CT, cortical thickness; SA, surface area; Vol, volume.*

##### C2. Removing infant and neonatal data

We additionally assessed our cortical thickness analyses' sensitivity to the inclusion of early-life (i.e. fetal and neonatal) data. All results, including the presence and directionality of age-varying and size-corrected sex differences, were highly consistent with our main analyses.

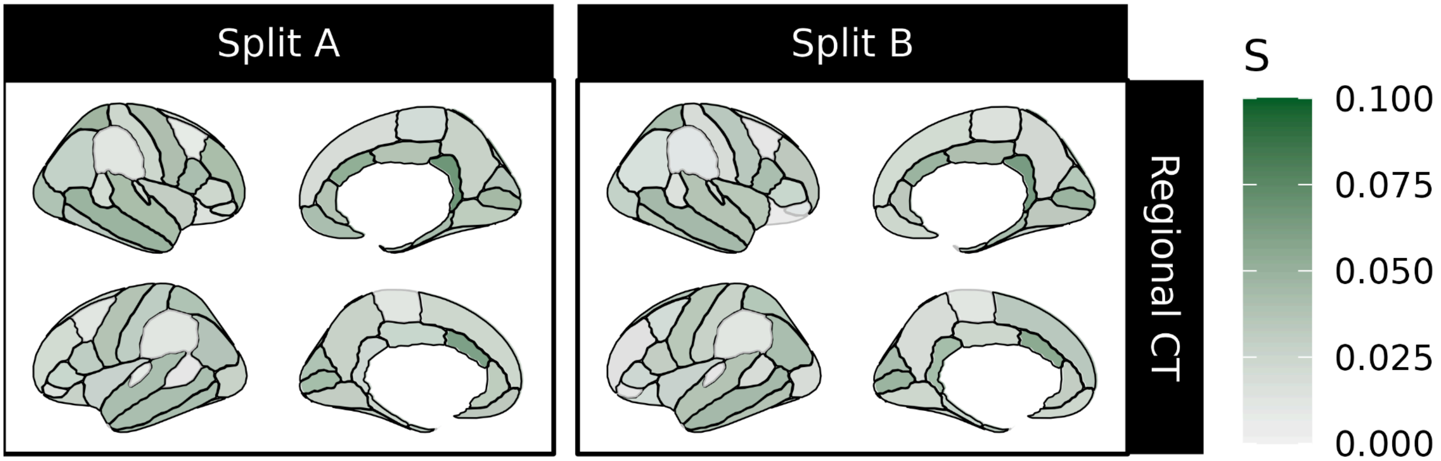


***Figure S58. Age-varying sex-biases excluding neonatal and infant data.*** *Magnitude of the age-varying component of sex (age-by-sex term) on cortical thickness (CT) IDPs after age two years. Effect sizes (S) are reported using the robust effect size index (RESI). Black outlines indicate significant effects at p <0.05, using false discovery rate (FDR) correction for multiple comparisons. Abbrv: CT, cortical thickness.*

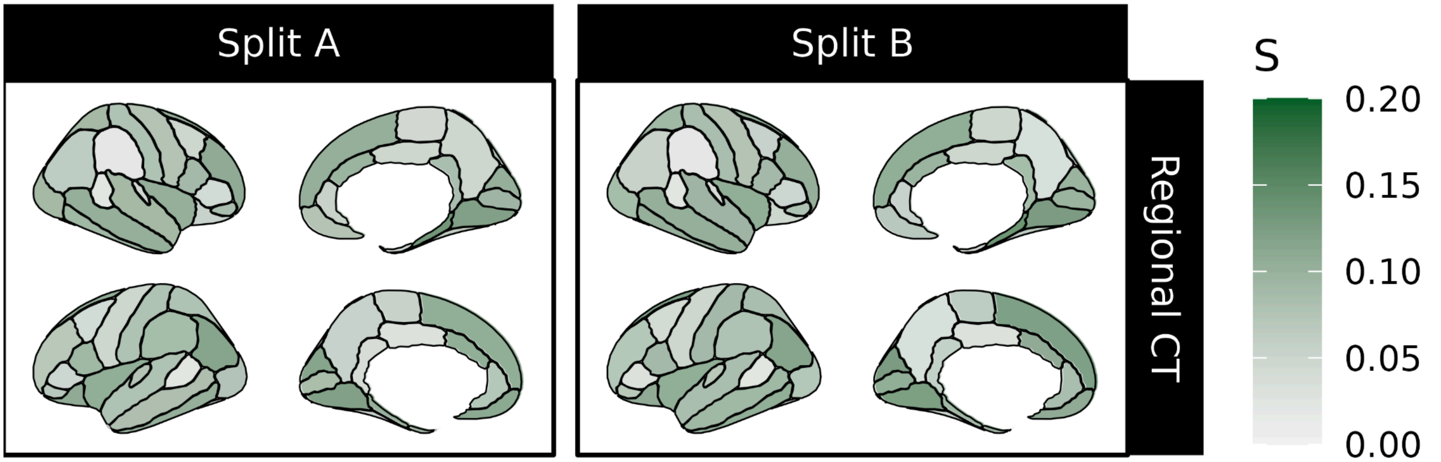


***Figure S59. Sex effects when controlling for total brain size, removing infant and neonatal data.*** *Magnitude of all sex effects on cortical thickness (CT) IDPs after age two years when controlling for mean cortical thickness. Effect sizes (S) are reported using the robust effect size index (RESI). Black outlines indicate significant effects at p <0.05, using false discovery rate (FDR) correction for multiple comparisons. Abbrv: CT, cortical thickness.*

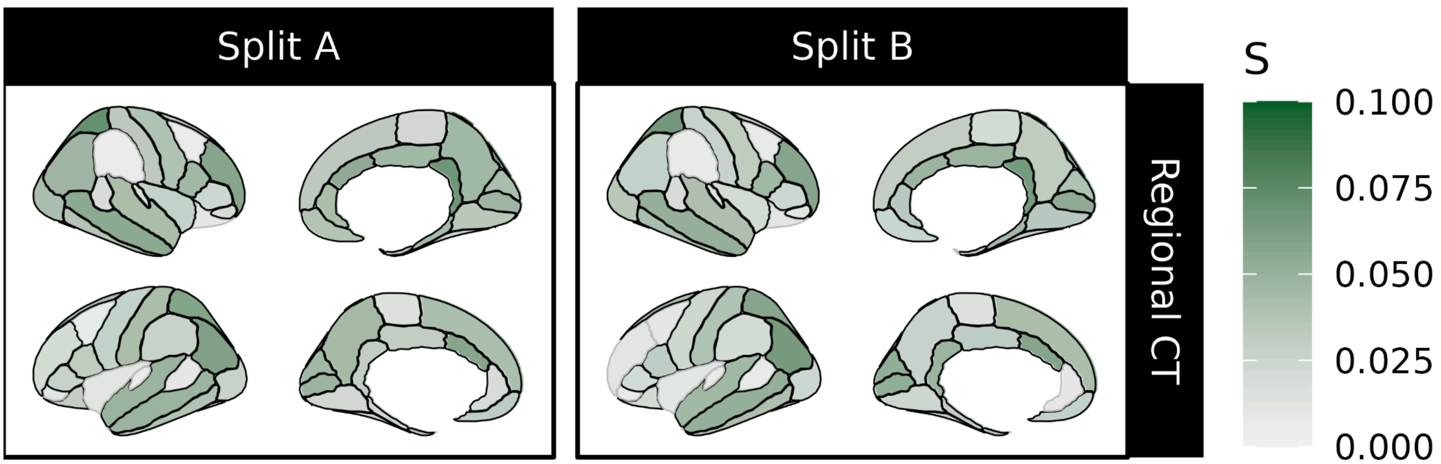


***Figure S60. Age-varying sex effects when controlling for total brain size, removing infant and neonatal data.*** *Magnitude of the age-varying component of sex (age-by-sex term) on cortical thickness (CT) IDPs after age two years when controlling for mean cortical thickness. Effect sizes (S) are reported using the robust effect size index (RESI). Black outlines indicate significant effects at p <0.05, using false discovery rate (FDR) correction for multiple comparisons. Abbrv: CT, cortical thickness.*

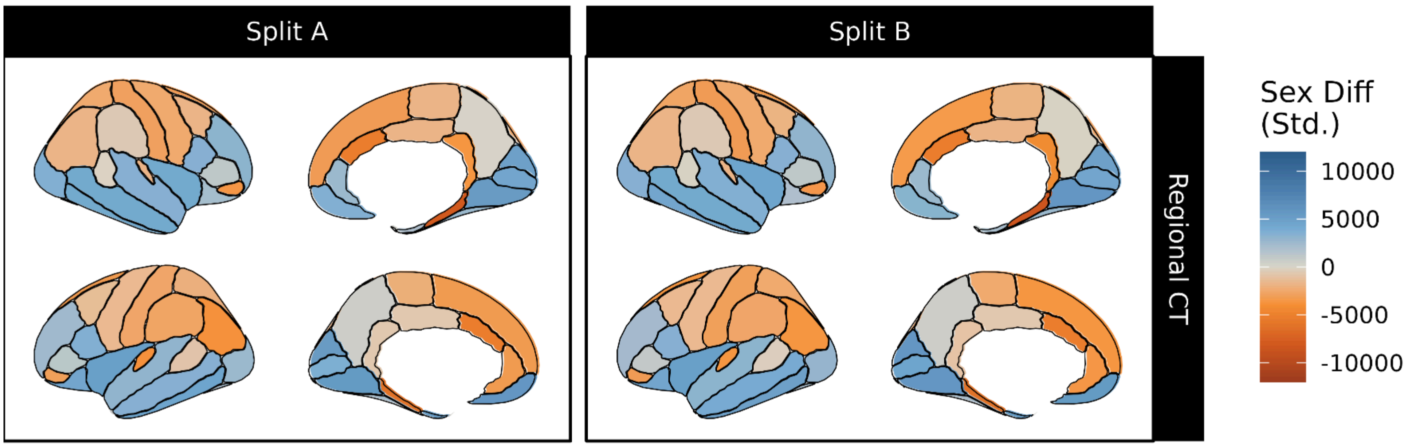


***Figure S61. Sex differences in IDPs' medians over the lifespan after age two.*** *Sex bias in IDP median trajectory after age two, controlling for mean cortical thickness. Orange indicates female bias and blue indicates male bias. Black outlines indicate significant effects at p <0.05, using false discovery rate (FDR) correction for multiple comparisons. Abbrv: CT, cortical thickness.*

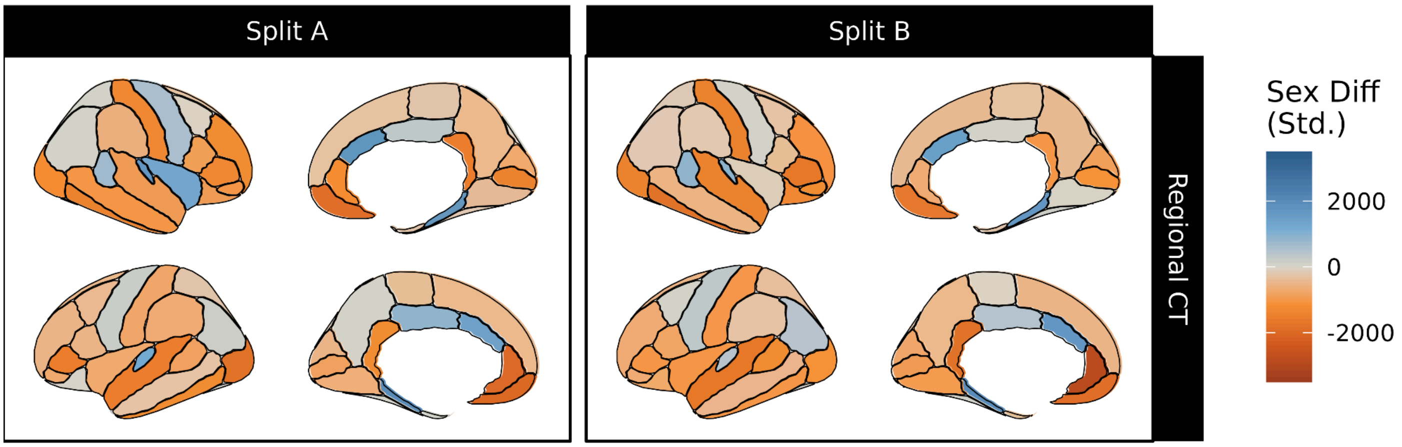


***Figure S62. Sex differences in IDPs' variabilities over the lifespan after age two.*** *Sex bias in IDP variability after age two, controlling for mean cortical thickness. Orange indicates female bias and blue indicates male bias. Black outlines indicate significant effects at p <0.05, using false discovery rate (FDR) correction for multiple comparisons. Abbrv: CT, cortical thickness.*

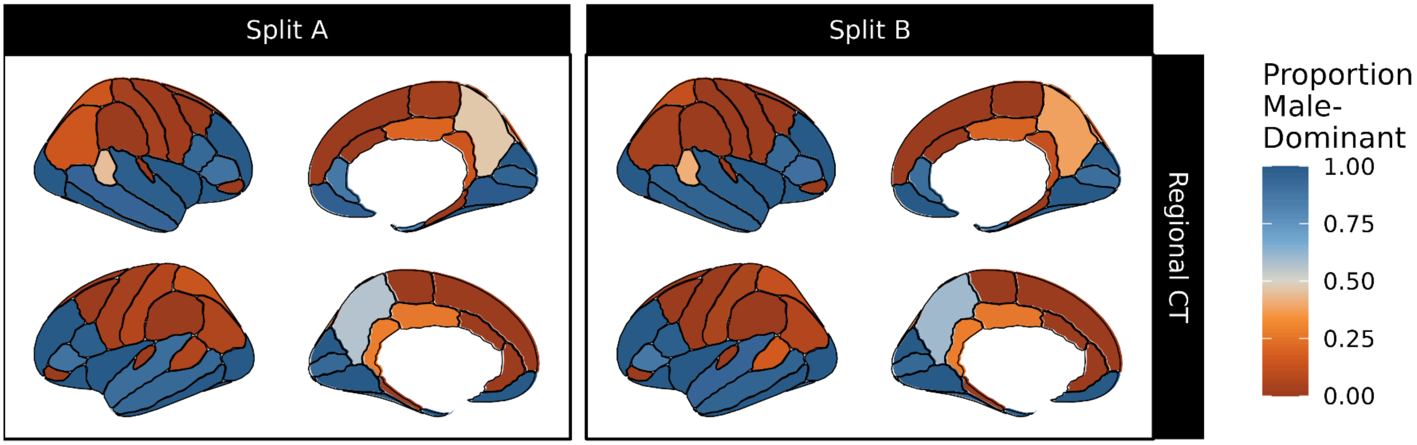


***Figure S63. Proportion of the lifespan an IDP's median is biased towards males after age two.*** *Blue indicates more of the lifespan being male-biased (i.e. larger in males), while orange indicates a longer time spent female-biased. Black outlines indicate significant sex effects at p <0.05, using false discovery rate (FDR) correction for multiple comparisons. Abbrv: CT, cortical thickness; SA, surface area; Vol, volume.*

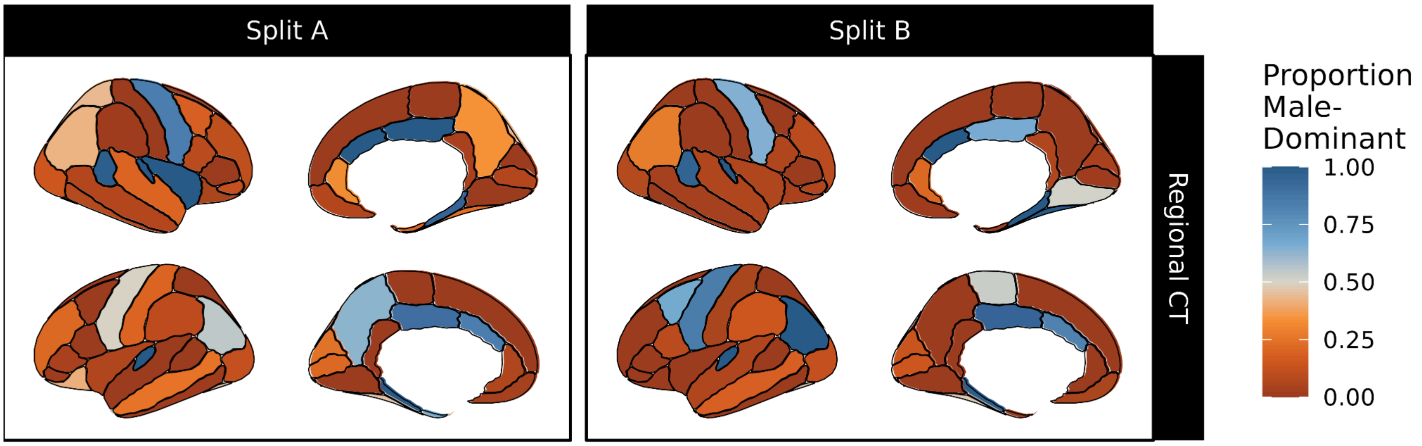


***Figure S64. Proportion of the lifespan an IDP's variability is biased towards males after age two.*** *Blue indicates more of the lifespan being male-biased (i.e. more variable in males), while orange indicates a longer time spent female-biased. Black outlines indicate significant sex effects at p <0.05, using false discovery rate (FDR) correction for multiple comparisons. Abbrv: CT, cortical thickness; SA, surface area; Vol, volume.*

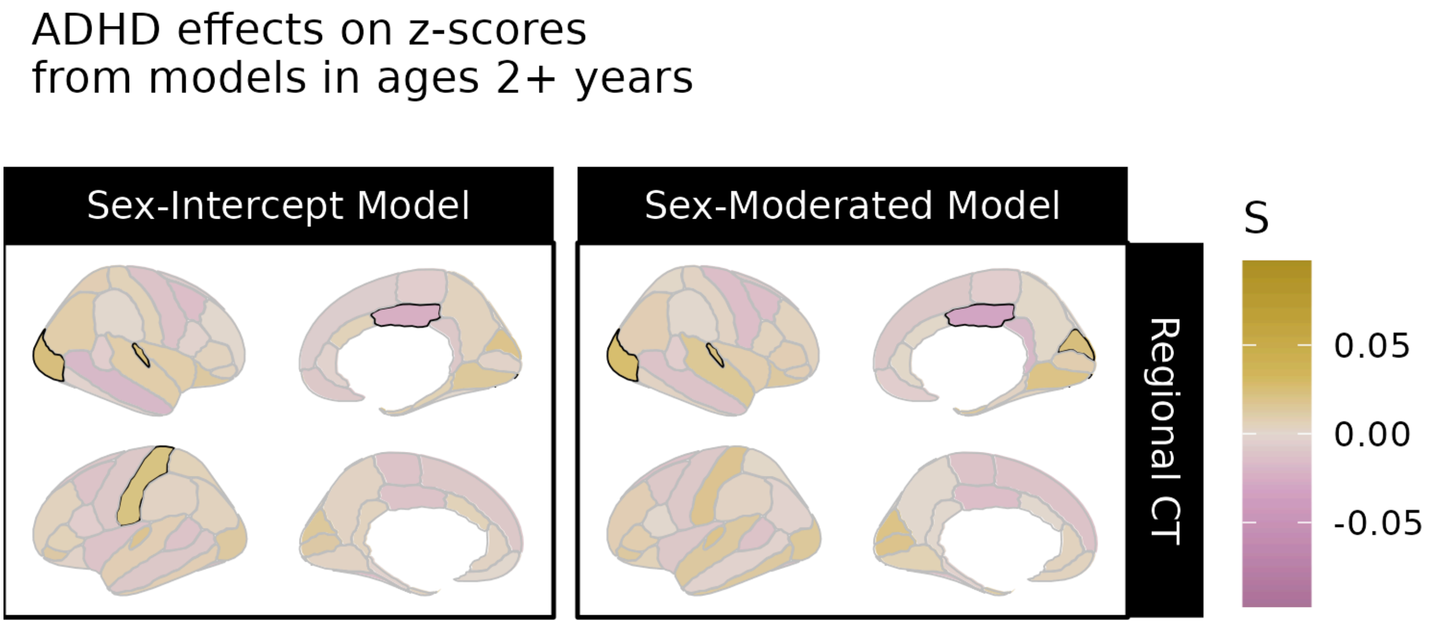


***Figure S65. Significant differences associated with ADHD in centile z-scores derived from sex-moderated and sex-intercept brain charts fit without fetal or infant data.*** *Differences in centile z-scores in individuals with attention-deficit/hyperactivity disorder (ADHD) relative to site-matched controls. The left panel displays differences in centile z-scores derived from sex-intercept models (i.e. models keeping sex differences in IDPs constant across the lifespan) while the right panel displays differences in centile z-scores derived from sex-moderated models, which allow age-by-sex effects. Gold indicates cases tend to have higher centile z-scores than controls for a given IDP, while purple indicates cases tend to have smaller centile z-scores. Black outlines indicate significant effects at p <0.05, FDR-corrected in each model. Abbrv: CT, cortical thickness.*

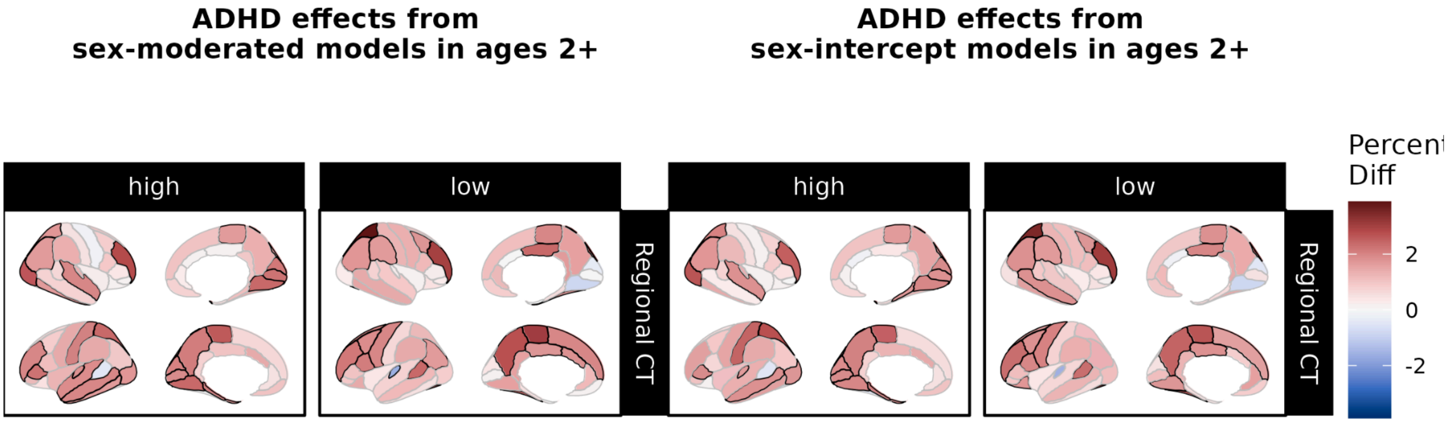


***Figure S66. Over- and under-representation of ADHD in extreme centiles according to sex-moderated and sex-intercept brain charts fit without fetal or infant data.*** *Using centile scores derived from our sex-moderated (left panel) and sex-intercept (right panel) brain charts, we compared the proportions of individuals with attention-deficit/hyperactivity disorder (ADHD) to the proportion of site-matched controls with centiles above the 95th percentile ("high", left column) and below the 5th percentile ("low", right column). Reds indicate cases are over-represented relative to controls, while greens indicate that cases are under-represented relative to controls. Black outlines indicate significant effects at p <0.05, FDR-corrected within each brain chart model. Abbrv: ADHD, attention-deficit/hyperactivity disorder; CT, cortical thickness.*

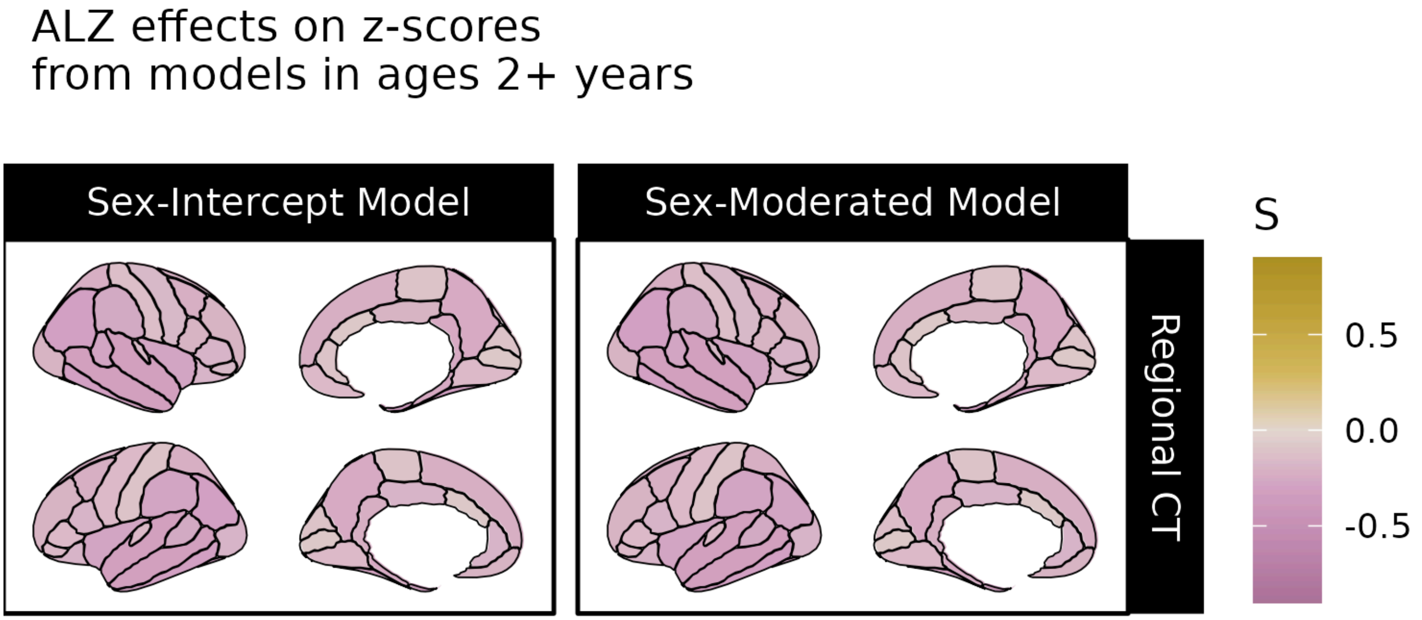


***Figure S67. Significant differences associated with Alzheimer's disease in centile z-scores derived from sex-moderated and sex-intercept brain charts without fetal and neonatal data.*** *Differences in centile z-scores in individuals with Alzheimer's disease (ALZ) relative to site-matched controls. The left panel displays differences in centile z-scores derived from sex-intercept models (i.e. models keeping sex differences in IDPs constant across the lifespan) while the right panel displays differences in centile z-scores derived from sex-moderated models, which allow age-by-sex effects. Gold indicates cases tend to have higher centile z-scores than controls for a given IDP, while purple indicates cases tend to have smaller centile z-scores. Black outlines indicate significant effects at p <0.05, FDR-corrected in each model. Abbrv: CT, cortical thickness.*

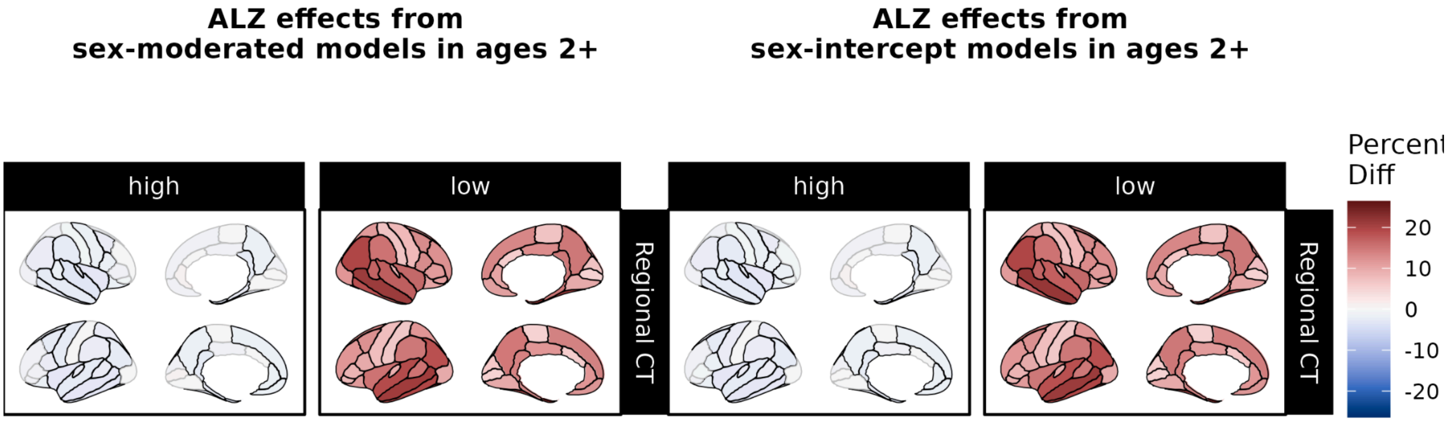


***Figure S68. Over- and under-representation of Alzheimer's disease in extreme centiles according to sex-moderated and sex-intercept brain charts fit without fetal or infant data.*** *Using centile scores derived from our sex-moderated (left panel) and sex-intercept (right panel) brain charts, we compared the proportions of individuals with Alzheimer's disease (ALZ) to the proportion of site-matched controls with centiles above the 95th percentile ("high", left column) and below the 5th percentile ("low", right column). Reds indicate cases are over-represented relative to controls, while greens indicate that cases are under-represented relative to controls. Black outlines indicate significant effects at p <0.05, FDR-corrected within each brain chart model. Abbrv: ALZ, Alzheimer's disease; CT, cortical thickness.*

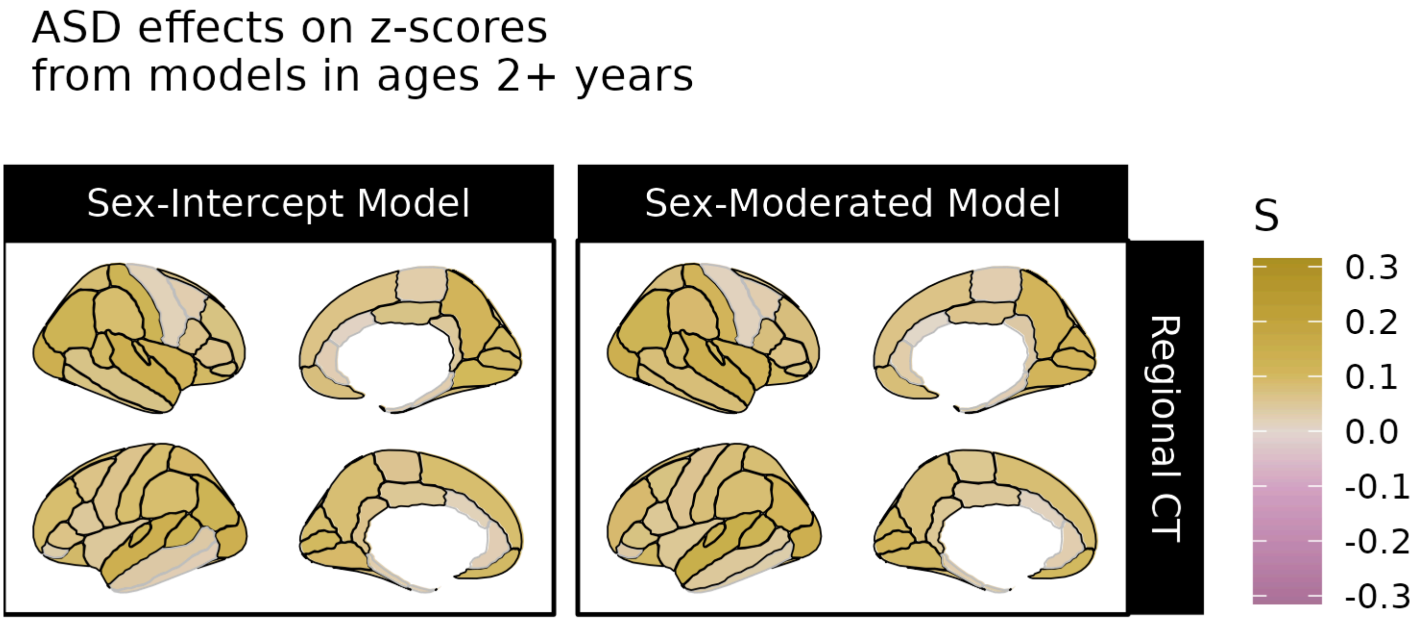


***Figure S69. Significant differences associated with autism spectrum disorder in centile z-scores derived from sex-moderated and sex-intercept brain charts without fetal or infant data.*** *Differences in centile z-scores in individuals with autism spectrum disorder (ASD) relative to site-matched controls. The left panel displays differences in centile z-scores derived from sex-intercept models (i.e. models keeping sex differences in IDPs constant across the lifespan) while the right panel displays differences in centile z-scores derived from sex-moderated models, which allow age-by-sex effects. Gold indicates cases tend to have higher centile z-scores than controls for a given IDP, while purple indicates cases tend to have smaller centile z-scores. Black outlines indicate significant effects at p <0.05, FDR-corrected in each model. Abbrv: CT, cortical thickness.*

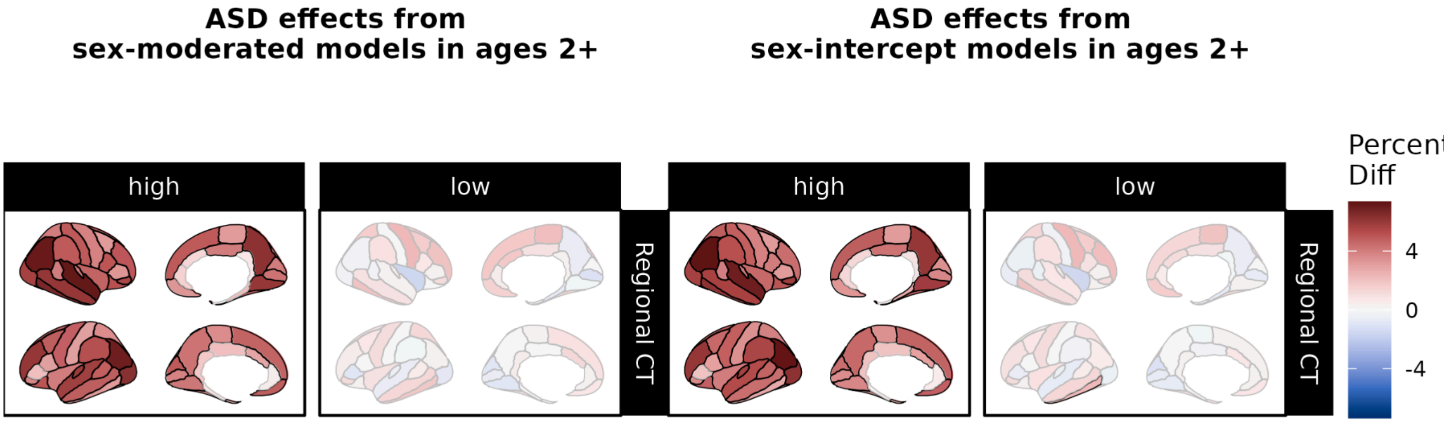


***Figure S70. Over- and under-representation of autism spectrum disorder in extreme centiles according to sex-moderated and sex-intercept brain charts fit without fetal or infant data.*** *Using centile scores derived from our sex-moderated (left panel) and sex-intercept (right panel) brain charts, we compared the proportions of individuals with autism spectrum disorder (ASD) to the proportion of site-matched controls with centiles above the 95th percentile ("high", left column) and below the 5th percentile ("low", right column). Reds indicate cases are over-represented relative to controls, while greens indicate that cases are under-represented relative to controls. Black outlines indicate significant effects at p <0.05, FDR-corrected within each brain chart model. Abbrv: ASD, autism spectrum disorder; CT, cortical thickness.*

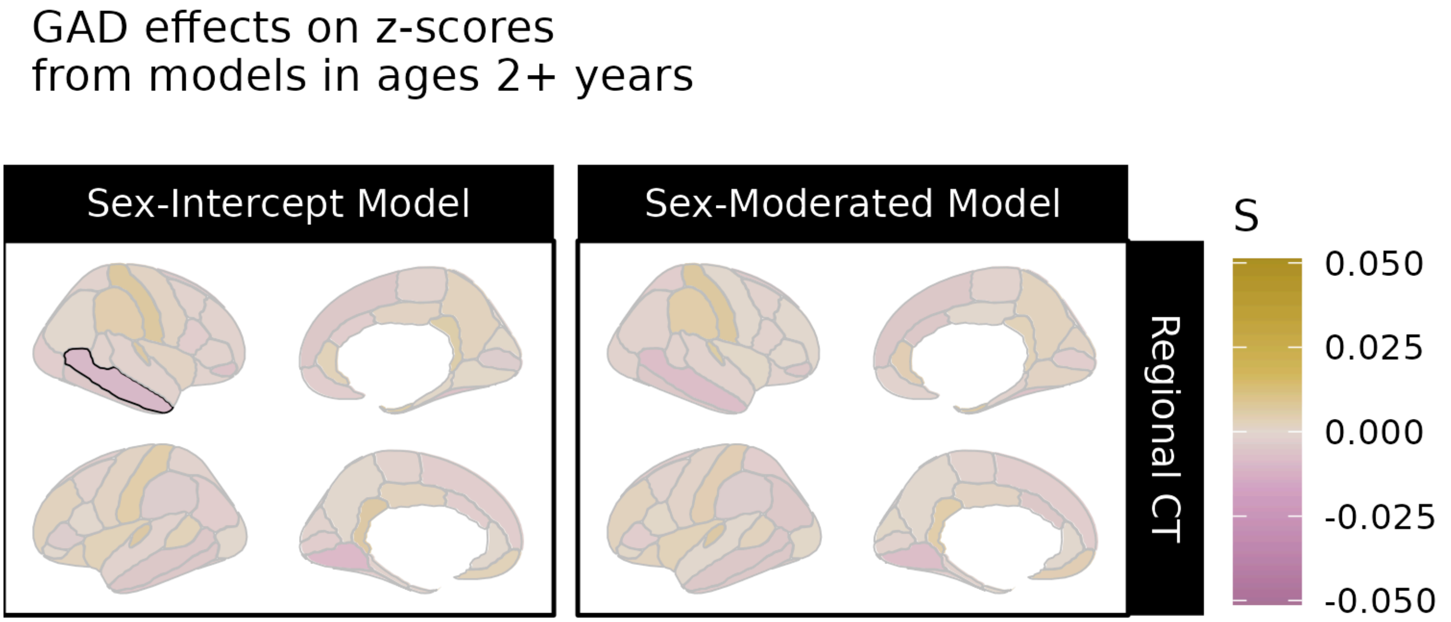


***Figure S71. Significant differences associated with generalized anxiety disorder in centile z-scores derived from sex-moderated and sex-intercept brain charts without fetal or infant data.*** *Differences in centile z-scores in individuals with generalized anxiety disorder (GAD) relative to site-matched controls. The left panel displays differences in centile z-scores derived from sex-intercept models (i.e. models keeping sex differences in IDPs constant across the lifespan) while the right panel displays differences in centile z-scores derived from sex-moderated models, which allow age-by-sex effects. Gold indicates cases tend to have higher centile z-scores than controls for a given IDP, while purple indicates cases tend to have smaller centile z-scores. Black outlines indicate significant effects at p <0.05, FDR-corrected in each model. Abbrv: CT, cortical thickness.*

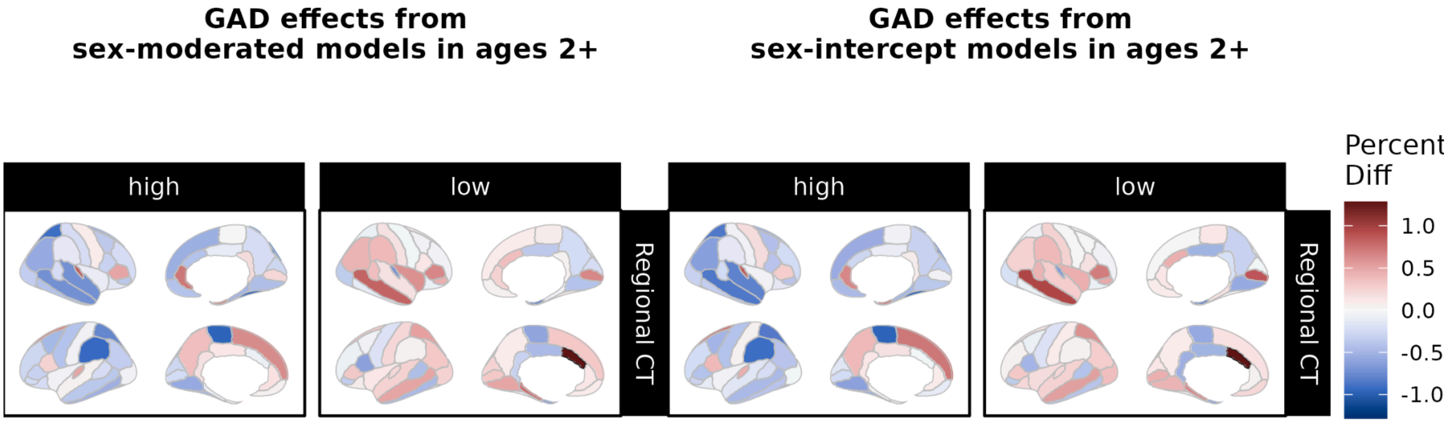


***Figure S72. Over- and under-representation of generalized anxiety disorder in extreme centiles according to sex-moderated and sex-intercept brain charts fit without fetal or infant data.*** *Using centile scores derived from our sex-moderated (left panel) and sex-intercept (right panel) brain charts, we compared the proportions of individuals with generalized anxiety disorder (GAD) to the proportion of site-matched controls with centiles above the 95th percentile ("high", left column) and below the 5th percentile ("low", right column). Reds indicate cases are over-represented relative to controls, while greens indicate that cases are under-represented relative to controls. Black outlines indicate significant effects at p <0.05, FDR-corrected within each brain chart model. Abbrv: GAD, generalized anxiety disorder; CT, cortical thickness.*

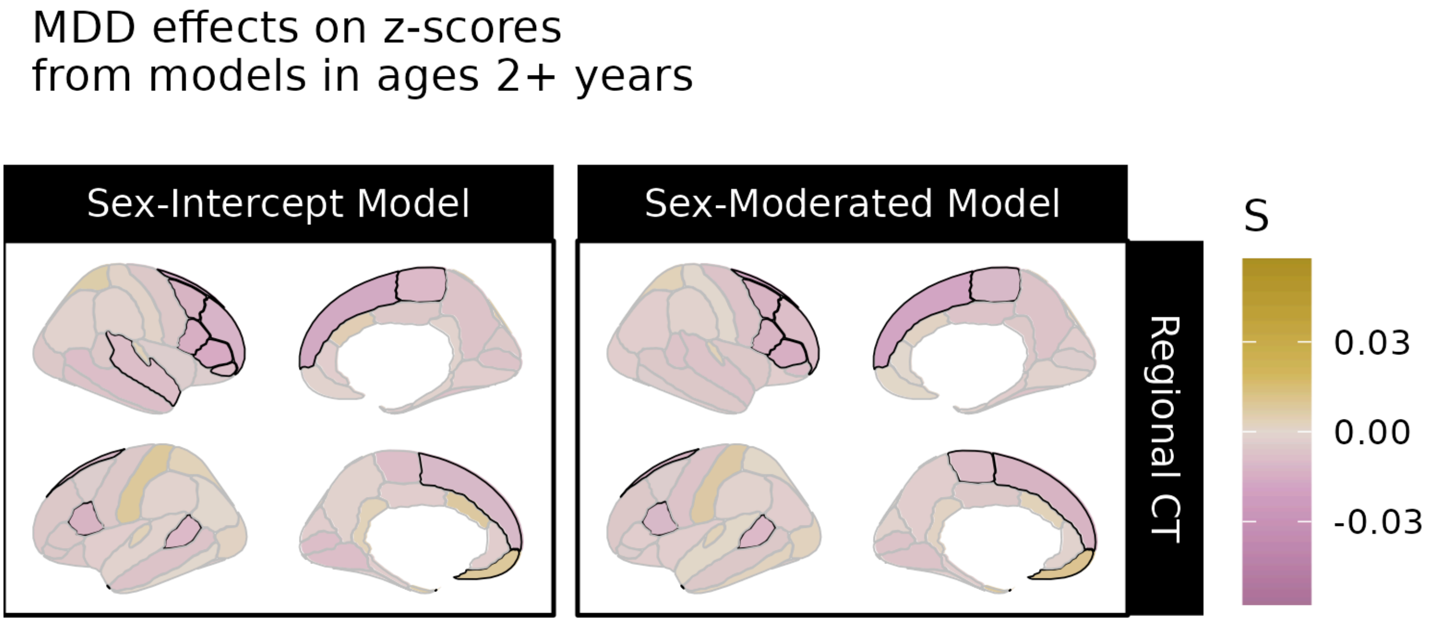


***Figure S73. Significant differences associated with major depressive disorder in centile z-scores derived from sex-moderated and sex-intercept brain charts.*** *Differences in centile z-scores in individuals with major depressive disorder (MDD) relative to site-matched controls. The left panel displays differences in centile z-scores derived from sex-intercept models (i.e. models keeping sex differences in IDPs constant across the lifespan) while the right panel displays differences in centile z-scores derived from sex-moderated models, which allow age-by-sex effects. Gold indicates cases tend to have higher centile z-scores than controls for a given IDP, while purple indicates cases tend to have smaller centile z-scores. Black outlines indicate significant effects at p <0.05, FDR-corrected in each model. Abbrv: CT, cortical thickness.*

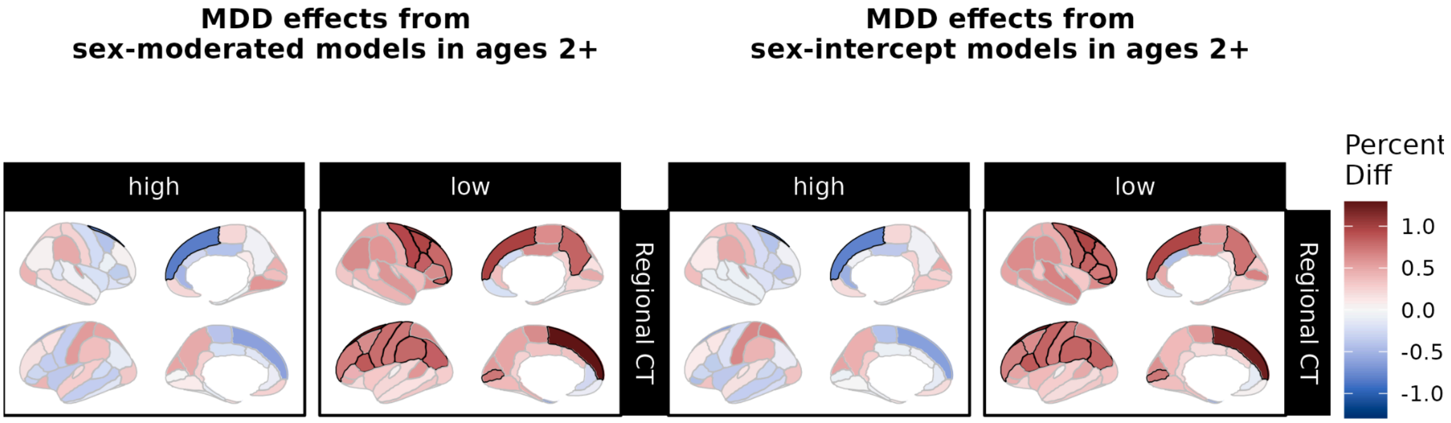


***Figure S74. Over- and under-representation of major depressive disorder in extreme centiles according to sex-moderated and sex-intercept brain charts fit without fetal or infant data.*** *Using centile scores derived from our sex-moderated (left panel) and sex-intercept (right panel) brain charts, we compared the proportions of individuals with major depressive disorder (MDD) to the proportion of site-matched controls with centiles above the 95th percentile ("high", left column) and below the 5th percentile ("low", right column). Reds indicate cases are over-represented relative to controls, while greens indicate that cases are under-represented relative to controls. Black outlines indicate significant effects at p <0.05, FDR-corrected within each brain chart model. Abbrv: MDD, major depressive disorder; CT, cortical thickness.*

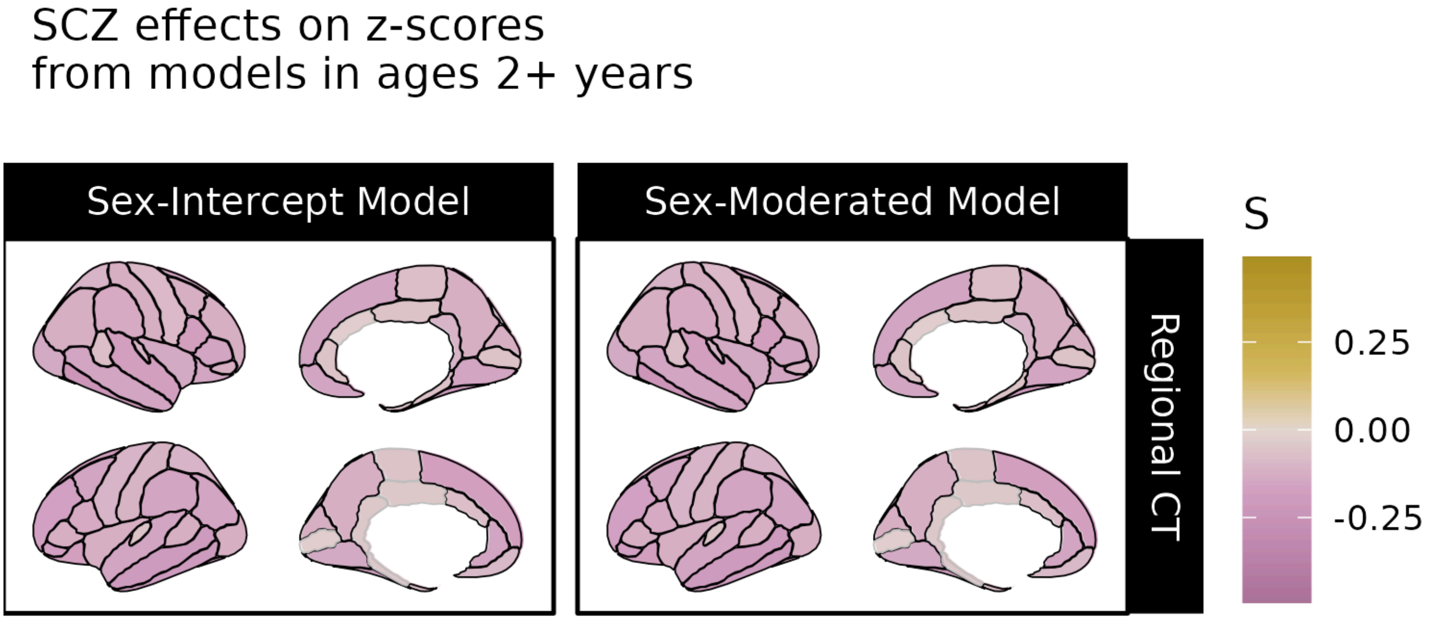


***Figure S75. Significant differences associated with schizophrenia in centile z-scores derived from sex-moderated and sex-intercept brain charts.*** *Differences in centile z-scores in individuals with schizophrenia (SCZ) relative to site-matched controls. The left panel displays differences in centile z-scores derived from sex-intercept models (i.e. models keeping sex differences in IDPs constant across the lifespan) while the right panel displays differences in centile z-scores derived from sex-moderated models, which allow age-by-sex effects. Gold indicates cases tend to have higher centile z-scores than controls for a given IDP, while purple indicates cases tend to have smaller centile z-scores. Black outlines indicate significant effects at p <0.05, FDR-corrected in each model. Abbrv: CT, cortical thickness.*

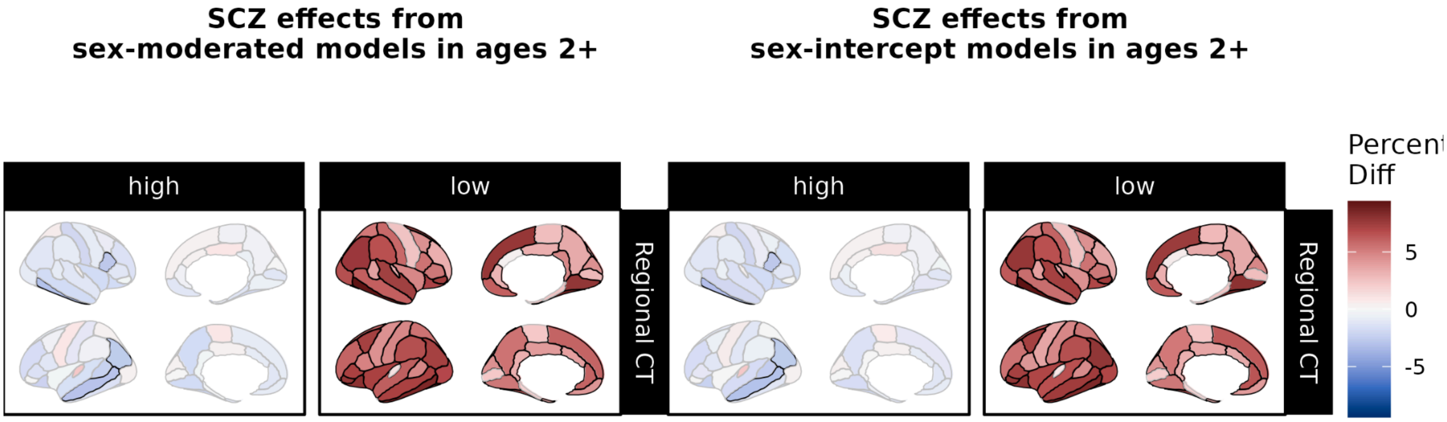


***Figure S76. Over- and under-representation of schizophrenia in extreme centiles according to sex-moderated and sex-intercept brain charts fit without fetal or infant data.*** *Using centile scores derived from our sex-moderated (left panel) and sex-intercept (right panel) brain charts, we compared the proportions of individuals with schizophrenia (SCZ) to the proportion of site-matched controls with centiles above the 95th percentile ("high", left column) and below the 5th percentile ("low", right column). Reds indicate cases are over-represented relative to controls, while greens indicate that cases are under-represented relative to controls. Black outlines indicate significant effects at p <0.05, FDR-corrected within each brain chart model. Abbrv: SCZ, schizophrenia; CT, cortical thickness.*

#### D. Consortium Information

The members of consortia whose data was used in this manuscript include the following:

**Lifespan Brain Chart Consortium**

Gardner M.[1,2,3], Adamson C.[4,5], Adler S.[6], Anagnostou E.[7,8], Anderson, K.M.[9], Apergis-Schoute, A.M.[10], Areces-Gonzalez A.[11,12], Astle, D.E.[13], Auyeung B.[14,15], Ayub M.[16], Bae J.[17], Ball G.[4,18], Banca P.[19,20], Baron-Cohen S.[15,21], Beare R.[4,5], Bedford, S.A.[15], Benegal V.[22], Berken J.[1,2], Bethlehem, R.A.I.[15,23], Beyer F.[24], Biria M.[25,26], Blangero J.[27], Blesa C√°bez M.[28,29], Boardman, R.P.[28,29], Borzage M.[30,31,32], Bosch-Bayard, J.F.[33,34], Bourke N.[35], Buczek M.[1,2], Bullmore, E.T.[36], Calhoun, V.D.[37], Chakravarty, M.M.[38,34], Chen C.[39], Chertavian C.[40], Chetelat G.[41], Chong, Y.S.[42,43], Corvin A.[44], Costantino M.[45,46], Courchesne E.[47,48], Crivello F.[49], Cropley, V.L.[50], Crosbie J.[51], Crossley N.[52,53], Cyr K.[1,2], David, A.S.[54], Delarue M.[41], Delorme R.[55,56], Desrivieres S.[57], Devenyi, G.A.[58,59], Di Biase, M.A.[50,60], Dolan R.[61,62,63], Donald, K.A.[64], Donohoe G.[65], Dorfschmidt L.[66,67,40], Dunlop K.[68,69], Edwards, A.D.[70,71,72], Elison, J.T.[73], Ellis, C.T.[74,75], Elman, J.A.[76], Eyler L.[77,78], Fair, D.A.[73], Feczko E.[79,80], Fletcher, P.C.[81,82], Fonagy P.[83,84], Franz, C.E.[85], Galan-Garcia L.[86], Gholipour A.[87], Giedd J.[88,89], Gilmore, J.H.[90], Gispert López J.[91,92,93], Glahn, D.C.[95,96], Goodyer, I.M.[36], Grant, P.E.[97], Groenewold, N.A.[98,64], Gunning, F.M.[99], Gur, R.E.[66,40], Gur, R.C.[66,100], Hammill, C.F.[51,101], Hansson O.[102,103], Hedden T.[104,105], Heinz A.[106], Henson, R.N.[13,107], Heuer K.[108], Hoare J.[109], Holla B.[110,111], Holmes, A.J.[112], Huang H.[113], Im K.[114,115], Ipser J.[116], Jack Jr, C.R.[117], Jackowski, A.P.[118,119], Jia T.[120,121,122], Jones, D.T.[123,124], Jones, P.B.[36,125], Jung B.[1,2], Kafadar E.[1,2,3], Kahn, R.S.[126,127], Karandikar S.[1,2], Karlsson H.[128,129], Karlsson L.[130,129], Kawashima R.[131], Kelley, E.A.[132], Kern S.[133,134], Kim K.[135,136,137], Kitzbichler, M.G.[36], Kremen, W.S.[76], Lalonde F.[139], Landeau B.[41], Lerch J.[140,141,142], Lewis, J.D.[143], Li J.[144], Liao W.[144], Liston C.[145], Lombardo, M.V.[146,15], Luque Laguna P.[147], Lv J.[148,50], Macedo B.[1,2,3], Mallard, T.T.[149], Mandal A.[1,2,3], Marcelis M.[150], Mathias, S.R.[95], Mazoyer B.[49,151], McGuire P.[152], Meaney, M.J.[153], Mechelli A.[152], Mercedes L.[1,2], Merisaari H.[130], Merritt K.[54], Misic B.[154], Montagnese M.[155], Morgan, S.E.[156,157,158], Mothersill D.[159,160,161], Ortinau C.[162], Ossenkoppele R.[163,164], Ouyang M.[113], Palacios Martínez E.[91], Palaniyappan L.[165,166], Paly L.[41], Pan, P.M.[167,168], Pantelis C.[169,170,171], Park, M.M.[172,173], Paus T.[174,175], Pausova Z.[51,176], Paz-Linares D.[11,177], Pichet Binette A.[178,179], Pierce K.[180], Prem S.[1,2,3], Pulli, E.P.[130], Qian X.[181], Qiu A.[182],

Raznahan A.[139], Rittman T.[183], Robbins, T.W.[184], Rodrigue A.[95], Rollins, C.K.[185,186], Romero-Garcia R.[36,187], Ronan L.[36], Rosenberg, M.D.[188], Rowe, J.B.[189], Rowitch, D.H.[190], Salum, G.A.[191,192,193], Satterthwaite, T.D.[194,66], Schaare H.[195], Schachar, R.J.[51], Schultz, A.P.[196,197,96], Schöll M.[198,199,200], Seidlitz J.[66,67,40], Sha Z.[1,2,3], Sharp D.[201,202], Shinohara, R.T.[39,203], Skoog I.[133,134], Smyser, C.D.[204], Sperling, R.A.[196,205,96], Stein, D.J.[206], Stolicyn A.[207], Suckling J.[36,208], Sullivan G.[209], Sun K.[1,2,3], Thyreau B.[131], Toro R.[108,210], Traut N.[211,212], Tsvetanov, K.A.[183,213], Turk E.[214,215], Turk-Browne, N.B.[9,216], Tuulari, J.J.[130,217,129], Tzourio C.[218], Vachon-Presseau É.[219,220,221], Vaghi, M.M.[222,223], Valdes-Sosa, M.J.[86], Valdes-Sosa, P.A.[224,225], Valk, S.L.[226], van Amelsvoort T.[227], Vandekar, S.N.[228,229], Vasung L.[230], Victoria, L.W.[99], Villeneuve S.[179,231,178], Villringer A.[24,232], Vogel, J.W.[194,66], Váša F.[233], Vértes, P.E.[36,234], Wagstyl K.[235], Wang, Y.S.[236,237,238], Warfield, S.K.[87], Warrier V.[240], Westman E.[241,242], Westwater, M.L.[36], Whalley, H.C.[207,243], White, S.R.[36,244], Williams R.[1,2], Witte A.[232,24,245], Yang N.[236,237,238], Yeo B.[246,247,248], Yun H.[250], Zalesky A.[251], Zar, H.J.[98,252], Zettergren A.[133], Zhou, J.H.[181,253,246], Ziauddeen H.[36,254,255], Zimmerman D.[66,67,40], Zugman A.[256,257,168], Zuo, X.N.[236,237,238], Alexander-Bloch, A.F.[66,67,40]

[1] Lifespan Brain Institute, The Children's Hospital of Philadelphia and Penn Medicine, Philadelphia PA 19139, USA
[2] Department of Child and Adolescent Psychiatry and Behavioral Science, The Children's Hospital of Philadelphia, Philadelphia, PA, USA
[3] Department of Psychiatry, University of Pennsylvania, Philadelphia, PA, USA
[4] Developmental Imaging, Murdoch Children's Research Institute, Melbourne, Victoria, Australia
[5] Department of Medicine, Monash University, Melbourne, Victoria, Australia
[6] UCL Great Ormond Street Institute for Child Health, 30 Guilford St, Holborn, London WC1N 1EH
[7] Department of Pediatrics University of Toronto
[8] Holland Bloorview Kids Rehabilitation Hospital, Toronto, Canada
[9] Department of Psychology, Yale University, New Haven, CT, USA
[10] School of Biological and Behavioural Sciences, Centre for Brain and Behaviour, Queen Mary University of London, London, United Kingdom
[11] The Clinical Hospital of Chengdu Brain Science Institute, MOE Key Lab for NeuroInformation, University of Electronic Science and Technology of China, No. 2006, Xiyuan Ave., West Hi-Tech Zone, Chengdu, 611731, China
[12] University of Pinar del Río "Hermanos Saiz Montes de Oca", Cuba
[13] MRC Cognition and Brain Sciences Unit, University of Cambridge, Cambridge UK
[14] Department of Psychology, School of Philosophy, Psychology and Language Sciences, University of Edinburgh, Edinburgh, United Kingdom
[15] Autism Research Centre, Department of Psychiatry, University of Cambridge, Cambridge, CB2 0SZ, UK.
[16] University College London, Mental Health Neuroscience Research Department, Division of Psychiatry, London UK
[17] Department of Neuropsychiatry, Seoul National University Bundang Hospital, Seongnam, Korea
[18] Department of Paediatrics, University of Melbourne, Melbourne, Victoria, Australia
[19] Department of Neuroscience, Faculty of Medicine and Nursing, University of the Basque Country, UPV/EHU, Spain
[20] IKERBASQUE, Basque Foundation for Science, Bilbao, Spain
[21] Cambridge Lifetime Asperger Syndrome Service (CLASS), Cambridgeshire and Peterborough NHS Foundation Trust, Cambridge, United Kingdom
[22] Centre for Addiction Medicine, National Institute of Mental Health and Neurosciences (NIMHANS), Bengaluru, India 560029
[23] Brain Mapping Unit, Department of Psychiatry, University of Cambridge, Cambridge, CB2 0SZ, UK
[24] Department of Neurology, Max Planck Institute for Human Cognitive and Brain Sciences, Leipzig, 04103, Germany

[25] The Institute of Psychiatry, Psychology & Neuroscience (IoPPN), King's College London, UK
[26] Department of Psychology King's College London South London & Maudsley NHS Foundation Trust, UK
[27] Department of Human Genetics, South Texas Diabetes and Obesity Institute, University of Texas Rio Grande Valley
[28] Centre for Reproductive Health, Institute for Regeneration and Repair, University of Edinburgh, UK
[29] Centre for Clinical Brain Sciences, University of Edinburgh, Edinburgh, UK
[30] Fetal and Neonatal Institute, Division of Neonatology, Children's Hospital Los Angeles, Department of Pediatrics, Keck School of Medicine, University of Southern California, Los Angeles, California USA
[31] Alfred E. Mann Department of Biomedical Engineering, Viterbi School of Engineering, University of Southern California, Los Angeles, USA
[32] Department of Regulatory and Quality Sciences, Alfred E. Mann School of Pharmacy and Pharmaceutical Sciences, University of Southern California, Los Angeles, USA
[33] McGill Centre for Integrative Neuroscience, Ludmer Centre for Neuroinformatics and Mental Health, Montreal Neurological Institute
[34] McGill University
[35] Department of Brain Sciences, Imperial College London, London UK & Care Research & Technology Centre, UK Dementia Research Institute
[36] Department of Psychiatry, University of Cambridge, Cambridge, CB2 0SZ, UK
[37] Tri-institutional Center for Translational Research in Neuroimaging and Data Science, Georgia State University, Georgia Institute of Technology, and Emory University, Atlanta, GA, USA
[38] Computational Brain Anatomy (CoBrA) Laboratory, Cerebral Imaging Centre, Douglas Mental Health University Institute
[39] Penn Statistics in Imaging and Visualization Endeavor, Department of Biostatistics, Epidemiology, and Informatics, Perelman School of Medicine, University of Pennsylvania, Philadelphia, PA, USA
[40] Lifespan Brain Institute, The Children's Hospital of Philadelphia, Philadelphia, PA 19104
[41] Normandie Univ, UNICAEN, INSERM, U1237, PhIND "Physiopathology and Imaging of Neurological Disorders", Institut Blood and Brain @ Caen-Normandie, Cyceron, 14000 Caen, France
[42] Singapore Institute for Clinical Sciences, Agency for Science, Technology and Research, Singapore
[43] Department of Obstetrics and Gynaecology, Yong Loo Lin School of Medicine, National University of Singapore, Singapore
[44] Department of Psychiatry, Trinity College, Dublin, Ireland
[45] Cerebral Imaging Centre, Douglas Mental Health University Institute, Verdun, Canada
[46] Undergraduate program in Neuroscience, McGill University, Montreal, Canada
[47] Department of Neuroscience, University of California, San Diego, San Diego, CA 92093, USA
[48] Autism Center of Excellence, University of California, San Diego, San Diego, CA 92037, USA
[49] Institute of Neurodegenerative Disorders, CNRS UMR5293, CEA, University of Bordeaux
[50] Melbourne Neuropsychiatry Centre, University of Melbourne, Melbourne, Australia
[51] The Hospital for Sick Children, Toronto, Canada
[52] Department of Psychiatry, School of Medicine, Pontificia Universidad Católica de Chile, Diagonal Paraguay 362, Santiago 8330077, Chile
[53] Department of Psychiatry, University of Oxford OX3 7JX
[54] Division of Psychiatry, University College London, London, UK
[55] Child and Adolescent Psychiatry Department, Robert Debré University Hospital, AP-HP, F-75019, Paris France
[56] Human Genetics and Cognitive Functions , Institut Pasteur, F-75015, Paris France
[57] Social, Genetic and Developmental Psychiatry Centre, Institute of Psychiatry, Psychology & Neuroscience, King's College London, London, United Kingdom

[58] Cerebral Imaging Centre, Douglas Mental Health University Institute, Montreal, QC, Canada, McGill Department of Psychiatry, Montreal, QC, Canada
[59] Department of Psychiatry, McGill University, Montreal, QC, Canada
[60] Department of Psychiatry, Brigham and Women's Hospital, Harvard Medical School, Boston, Massachusetts, United States
[61] Max Planck UCL Centre for Computational Psychiatry and Ageing Research, University College London, London, UK.
[62] Wellcome Centre for Human Neuroimaging, University College London, London, UK
[63] Wellcome Centre for Human Neuroimaging, 12 Queen Square, London WC1N 3AR
[64] Neuroscience Institute, University of Cape Town, Cape Town, South Africa
[65] Center for Neuroimaging, Cognition & Genomics (NICOG), School of Psychology, National University of Ireland Galway, Galway, Ireland
[66] Department of Psychiatry, University of Pennsylvania, Philadelphia, PA 19104
[67] Department of Child and Adolescent Psychiatry and Behavioral Science, The Children's Hospital of Philadelphia, Philadelphia, PA 19104
[68] Department of Psychiatry, University of Toronto, Toronto, Ontario, Canada
[69] Centre for Depression and Suicide Studies, Unity Health Network, Toronto, Ontario, Canada
[70] Centre for the Developing Brain, King's College London, London, UK
[71] Evelina London Children's Hospital
[72] MRC Centre for Neurodevelopmental Disorders, London.
[73] Institute of Child Development, Department of Pediatrics, Masonic Institute for the Developing Brain, University of Minnesota, Minneapolis, MN, United States
[74] Department of Psychology, Stanford University, Stanford, CA, USA
[75] Haskins Laboratories, New Haven, CT, USA
[76] Department of Psychiatry, Center for Behavior Genetics of Aging, University of California, San Diego, La Jolla, CA
[77] Desert-Pacific Mental Illness Research Education and Clinical Center, VA San Diego Healthcare, San Diego, CA, USA
[78] Department of Psychiatry, University of California San Diego, Los Angeles, CA, USA
[79] Masonic Institute for the Developing Brain, University of Minnesota, Minneapolis, Minnesota
[80] Department of Pediatrics, University of Minnesota, Minneapolis, Minnesota
[81] Department of Psychiatry, University of Cambridge, and Wellcome Trust MRC Institute of Metabolic Science, Cambridge Biomedical Campus, Cambridge, United Kingdom
[82] Cambridgeshire and Peterborough NHS Foundation Trust
[83] Department of Clinical, Educational and Health Psychology, University College London, London, UK
[84] Anna Freud National Centre for Children and Families, London UK
[85] Department of Psychiatry, Center for Behavior Genetics of Aging, University of California, San Diego, La Jolla, CA 92093
[86] Cuban Center for Neuroscience, La Habana, Cuba
[87] Computational Radiology Laboratory, Boston Children's Hospital, Boston, MA 02115
[88] Department of Child and Adolescent Psychiatry, University of California, San Diego, San Diego, CA 92093, USA
[89] Department of Psychiatry, University of California San Diego, San Diego, CA, USA
[90] Department of Psychiatry, University of North Carolina, Chapel Hill, NC, USA
[91] Barcelonaβeta Brain Research Center (BBRC), Pasqual Maragall Foundation, Barcelona, Spain
[92] Hospital del Mar Research Institute, Barcelona, Spain
[93] Centro de Investigación Biomédica en Red Bioingeniería, Biomateriales y Nanomedicina, Instituto de Salud Carlos III, Madrid, Spain

[94] Centro Nacional de Investigaciones Cardiovasculares (CNIC), Madrid, Spain
[95] Department of Psychiatry, Boston Children's Hospital and Harvard Medical School, Boston, MA 02115
[96] Harvard Medical School, Boston, MA 02115
[97] Division of Newborn Medicine and Neuroradiology, Fetal Neonatal Neuroimaging and Developmental Science Center, Boston Children's Hospital, Harvard Medical School, Boston, MA 02115, USA
[98] Department of Paediatrics and Child Health, Red Cross War Memorial Children's Hospital, SA-MRC Unit on Child & Adolescent Health, University of Cape Town, South Africa
[99] Weill Cornell Institute of Geriatric Psychiatry, Department of Psychiatry, Weill Cornell Medicine
[100] Lifespan Brain Institute, The Children's Hospital of Philadelphia, Philadelphia, PA 19105
[101] Mouse Imaging Centre, Toronto, Canada
[102] Clinical Memory Research Unit, Department of Clinical Sciences Malmö, Lund University, Malmö, Sweden
[103] Memory Clinic, Skåne University Hospital, Malmö, Sweden
[104] Department of Neurology, Icahn School of Medicine at Mount Sinai, New York, NY 10029, USA
[105] Athinoula A. Martinos Center for Biomedical Imaging, Department of Radiology, Massachusetts General Hospital, Harvard Medical School, Boston, MA 02129, USA
[106] Department of Psychiatry and Psychotherapy, Charite University Hospital Berlin, Berlin, Germany
[107] Department of Psychiatry, University of Cambridge, Cambridge, UK
[108] Institut Pasteur, Université Paris Cité, Unité de Neuroanatomie Appliquée et Théorique, F-75015 Paris, France
[109] Department of Psychiatry, University of Cape Town, Cape Town, South Africa.
[110] Department of Integrative Medicine, NIMHANS, Bengaluru-560029, India
[111] Accelerator Program for Discovery in Brain disorders using Stem cells (ADBS), Department of Psychiatry, NIMHANS, Bengaluru-560029, India
[112] Department of Psychiatry, Brain Health Institute, Rutgers University, Piscataway, NJ, USA
[113] Department of Radiology, Children's Hospital of Philadelphia and University of Pennsylvania, Philadelphia, PA 19104.
[114] Division of Newborn Medicine, Fetal Neonatal Neuroimaging and Developmental Science Center, Boston Children's Hospital, Harvard Medical School, Boston, MA 02115, USA
[115] Boston Children's Hospital, Boston, MA 02115
[116] Department of Psychiatry and Mental Health, Clinical Neuroscience Institute, University of Cape Town
[117] Department of Radiology, Mayo Clinic, Rochester, MN 55905, USA
[118] Department of Psychiatry, Universidade Federal de S√£o Paulo
[119] National Institute of Developmental Psychiatry, CNPq
[120] Institute of Science and Technology for Brain-Inspired Intelligence, Fudan University, Shanghai, 200433, China
[121] Key Laboratory of Computational Neuroscience and BrainInspired Intelligence (Fudan University), Ministry of Education, Shanghai, China
[122] Centre for Population Neuroscience and Precision Medicine (PONS), Institute of Psychiatry, Psychology and Neuroscience, SGDP Centre, King's College London, London SE5 8AF, UK
[123] Department of Neurology, Mayo Clinic, Rochester, MN, USA
[124] Department of Radiology, Mayo Clinic, Rochester, MN, USA
[125] Cambridgeshire and Peterborough NHS Foundation Trust, Huntingdon, United Kingdom
[126] Department of Psychiatry, Icahn School of Medicine at Mount Sinai, New York, NY, USA
[127] Department of Psychiatry, Icahn School of Medicine, Mount Sinai, New York, USA
[128] Department of Clinical Medicine, Department of Psychiatry and Turku Brain and Mind Center, FinnBrain Birth Cohort Study, University of Turku and Turku University Hospital, Turku, Finland
[129] Centre for Population Health Research, Turku University Hospital and University of Turku, Turku, Finland

[130] FinnBrain Birth Cohort Study, Turku Brain and Mind Center, Department of Clinical Medicine, University of Turku and Turku University Hospital, Turku, Finland
[131] Institute of Development, Aging and Cancer, Tohoku University, Seiryocho, Aobaku, Sendai 980-8575, Japan
[132] Queen's University, Departments of Psychology and Psychiatry, Centre for Neuroscience Studies, Kingston, Ontario, Canada
[133] Neuropsychiatric Epidemiology Unit, Department of Psychiatry and Neurochemistry, Institute of Neuroscience and Physiology, the Sahlgrenska Academy, Centre for Ageing and Health (AGECAP) at the University of Gothenburg, Sweden
[134] Region Västra Götaland, Sahlgrenska University Hospital, Psychiatry, Cognition and Old Age Psychiatry Clinic, Gothenburg, Sweden
[135] Department of Brain and Cognitive Sciences, Seoul National University College of Natural Sciences, Seoul, Republic of Korea
[136] Department of Neuropsychiatry, Seoul National University Bundang Hospital, Seongnam, Republic of Korea
[137] Department of Psychiatry, Seoul National University College of Medicine, Seoul, Republic of Korea
[138] Department of Brain and Cognitive Science, Seoul National University College of Natural Sciences
[139] Section on Developmental Neurogenomics, Human Genetics Branch, National Institute of Mental Health, Bethesda, MD, USA
[140] Department of Medical Biophysics, University of Toronto, Toronto, ON, Canada
[141] Mouse Imaging Centre, The Hospital for Sick Children, Toronto, ON, Canada
[142] Wellcome Centre for Integrative Neuroimaging, FMRIB, Nuffield Department of Clinical Neuroscience, University of Oxford, Oxford, UK
[143] Montreal Neurological Institute, McGill University, Montreal, Canada
[144] The Clinical Hospital of Chengdu Brain Science Institute, University of Electronic Science and Technology of China, Chengdu 611731, China
[145] Department of Psychiatry and Brain and Mind Research Institute, Weill Cornell Medicine
[146] Laboratory for Autism and Neurodevelopmental Disorders, Center for Neuroscience and Cognitive Systems @UniTn, Istituto Italiano di Tecnologia, Rovereto, Italy
[147] The Cardiff University Brain Research Imaging Centre (CUBRIC), Cardiff University, Cardiff, UK
[148] School of Biomedical Engineering & Brain and Mind Centre, The University of Sydney, Sydney, NSW, Australia
[149] Department of Psychology, University of Texas, Austin, Texas 78712, USA
[150] Department of Psychiatry and Neuropsychology, School of Mental Health and Neuroscience, EURON, Maastricht University Medical Centre, PO Box 616, 6200 MD, Maastricht, the Netherlands; Institute for Mental Health Care Eindhoven (GGzE), Eindhoven, the Netherlands.
[151] Bordeaux University Hospital
[152] Professor, Department of Psychosis Studies, Institute of Psychiatry, Psychology and Neuroscience, King's College London, UK
[153] Ludmer Centre for Neuroinformatics and Mental Health, Douglas Mental Health University Institute, McGill University, Montreal, Quebec, Canada; Singapore Institute for Clinical Sciences, Singapore
[154] McConnell Brain Imaging Centre, Montreal Neurological Institute, McGill University, Montreal, QC H3A 2B4, Canada
[155] Psychology Department, University of Cambridge, Downing Pl, Cambridge, UK
[156] School of Biomedical Engineering and Imaging Sciences, King's College London, London UK
[157] Department of Computer Science and Technology, University of Cambridge, Cambridge CB3 0FD, United Kingdom
[158] Department of Psychiatry, University of Cambridge, Cambridge CB2 0SZ, United Kingdom
[159] Department of Psychology, School of Business, National College of Ireland, Dublin, Ireland

[160] School of Psychology & Center for Neuroimaging and Cognitive Genomics, National University of Ireland Galway, Galway, Ireland
[161] Department of Psychiatry, Trinity College Dublin, Dublin, Ireland
[162] Department of Pediatrics, Washington University in St. Louis, St. Louis, Missouri, United States
[163] Alzheimer Center Amsterdam, Department of Neurology, Amsterdam Neuroscience, Vrije Universiteit Amsterdam, Amsterdam UMC, Amsterdam, The Netherlands
[164] Lund University, Clinical Memory Research Unit, Lund, Sweden
[165] Douglas Mental Health University Institute, Department of Psychiatry, McGill University, Quebec, Canada
[166] Robarts Research Institute, University of Western Ontario, London, Ontario, Canada.
[167] Department of Psychiatry, Universidade Federal de S√£o Paulo, Brazil
[168] National Institute of Developmental Psychiatry for Children and Adolescents (INPD), Brazil.
[169] Melbourne Neuropsychiatry Centre, Department of Psychiatry, The University of Melbourne and Melbourne Health, Carlton South, Victoria, Australia
[170] Melbourne School of Engineering, The University of Melbourne, Parkville, Victoria, Australia
[171] Florey Institute of Neuroscience and Mental Health, Parkville, VIC, Australia
[172] Cerebral Imaging Centre, Douglas Mental Health University Institute, Montreal, Canada
[173] Integrated Program in Neuroscience, McGill University, Montreal, Canada
[174] Department of Psychiatry, Faculty of Medicine and Centre Hospitalier Universitaire Sainte-Justine, University of Montreal, Montreal, Quebec, Canada
[175] Departments of Psychiatry and Psychology, University of Toronto, Toronto, ON, Canada.
[176] Departments of Physiology and Nutritional Sciences, University of Toronto, Toronto, Canada
[177] Cuban Neuroscience Center, Havana, Cuba
[178] Department of Psychiatry, Faculty of Medicine, McGill University, Montreal, Qc, H3A 1Y2, Canada
[179] Douglas Mental Health University Institute, Montreal, Qc, H4H 1R3, Canada.
[180] Department of Neurosciences, University of California, San Diego La Jolla, CA, USA
[181] Center for Sleep and Cognition, Yong Loo Lin School of Medicine, National University of Singapore, Singapore
[182] Department of Health Technology and Informatics, Mental Health Research Centre, The Hong Kong Polytechnic University
[183] Department of Clinical Neurosciences, University of Cambridge, Cambridge UK
[184] Department of Psychology and the Behavioural and Clinical Neuroscience Institute, University of Cambridge, Cambridge, CB2 3EB, UK
[185] Department of Neurology, Harvard Medical School
[186] Department of Neurology, Boston Children's Hospital, Boston, MA 02115
[187] Instituto de Biomedicina de Sevilla (IBiS) HUVR/CSIC/Universidad de Sevilla, Dpto. de Fisiología Médica y Biofísica, Spain
[188] Department of Psychology, Neuroscience Institute, University of Chicago
[189] Department of Clinical Neurosciences, University of Cambridge, Cambridge, UK
[190] Department of Paediatrics and Wellcome-MRC Cambridge Stem Cell Institute, University of Cambridge, Hills Road, Cambridge, UK
[191] Department of Psychiatry, Universidade Federal do Rio Grande do Sul (UFRGS)
[192] National Institute of Developmental Psychiatry (INPD)
[193] Child Mind Institute, New York, USA
[194] Lifespan Informatics & Neuroimaging Center, University of Pennsylvania, Philadelphia, PA 19104
[195] U Bremen Research Alliance, Bremen, Germany
[196] Harvard Aging Brain Study, Department of Neurology, Massachusetts General Hospital, Boston, MA 02114
[197] Athinoula A. Martinos Center for Biomedical Imaging, Department of Radiology, Massachusetts General Hospital, Charlestown, MA 02129, USA

[198] Wallenberg Centre for Molecular and Translational Medicine, University of Gothenburg, Gothenburg, Sweden
[199] Department of Psychiatry and Neurochemistry, University of Gothenburg, Sweden
[200] Dementia Research Centre, Queen's Square Institute of Neurology, University College London, UK
[201] Department of Brain Sciences, Imperial College London, London UK
[202] Care Research & Technology Centre, UK Dementia Research Institute
[203] Center For AI And Data Science For Integrated Diagnostics, Department of Radiology, Perelman School of Medicine, University of Pennsylvania, Philadelphia, PA, USA
[204] Departments of Neurology, Pediatrics, and Radiology, Washington University School of Medicine, St. Louis, United States
[205] Center for Alzheimer Research and Treatment, Department of Neurology, Brigham and Women's Hospital, Boston, MA 02115
[206] SA MRC Unit on Risk & Resilience in Mental Disorders, Dept of Psychiatry and Neuroscience Institute, University of Cape Town, Cape Town, South Africa
[207] Division of Psychiatry, Centre for Clinical Brain Sciences, University of Edinburgh, UK
[208] Cambridge and Peterborough Foundation NHS Trust
[209] Centre for Clinical Brain Sciences, University of Edinburgh, Edinburgh, UK.
[210] Université de Paris, Paris, France
[211] Department of Neuroscience, Institut Pasteur, Paris, France
[212] Center for Research and Interdisciplinarity (CRI), Université Paris Descartes, Paris, France
[213] Department of Psychology, University of Cambridge, Cambridge, UK
[214] Department of Cognitive Neuropsychology, Tilburg University, Warandelaan 2, 5000 LE, Tilburg, the Netherlands
[215] Department of Neonatology, University Medical Center Utrecht, Utrecht University Heidelberglaan 100, 3584 CX, Utrecht, the Netherlands
[216] Wu Tsai Institute, Yale University, New Haven, CT, USA
[217] Neurocenter, Turku University Hospital, Turku, Finland
[218] Univ. Bordeaux, Inserm, Bordeaux Population Health Research Center, U1219, CHU Bordeaux, F-33000 Bordeaux, France
[219] Faculty of Dental Medicine and Oral Health Sciences, McGill University, Montreal, Qc, H3A 1G1, Canada
[220] Faculty of Dentistry, McGill University, Montreal, Qc, H3A 1G1, Canada
[221] Alan Edwards Centre for Research on Pain (AECRP), McGill University, Montreal, Qc, H3A 1G1, Canada
[222] School of Psychological Sciences, Birkbeck University of London, London, UK
[223] Centre for Brain and Cognitive Development, Birkbeck College, University of London, London, UK
[224] Brain-Computer Interface & Brain-Inspired Intelligence Key Laboratory of Sichuan Province, Chengdu, Sichuan, China
[225] University of Electronic Science and Technology of China/Cuban Center for Neuroscience
[226] Institute for Neuroscience and Medicine 7, Forschungszentrum Juelich; Max Planck Institute for Human Cognitive and Brain Sciences
[227] Department of Psychiatry & Neurosychology, Maastricht University, Maastricht, The Netherlands
[228] Department of Biostatistics, Vanderbilt University, Nashville, Tennessee, USA
[229] Department of Biostatistics, Vanderbilt University Medical Center, Nashville, Tennessee, USA
[230] Division of Newborn Medicine, Fetal Neonatal Neuroimaging and Developmental Science Center, Department of Pediatrics, Boston Children's Hospital, Boston, MA 02115
[231] McConnell Brain Imaging Center, Montreal Neurological Institute, McGill University, Montreal, Quebec, Canada
[232] Clinic for Cognitive Neurology, University of Leipzig Medical Center, Leipzig, 04103, Germany
[233] Department of Neuroimaging, King's College London, London, UK

[234] The Alan Turing Institute, London NW1 2DB, UK
[235] Wellcome Centre for Human Neuroimaging, Institute of Neurology, University College London, WC1N 3AR
[236] State Key Laboratory of Cognitive Neuroscience and Learning, Beijing Normal University, Beijing 100875, China
[237] Developmental Population Neuroscience Research Center, IDG/McGovern Institute for Brain Research, Beijing Normal University, Beijing 100875, China
[238] National Basic Science Data Center, Beijing 100190, China
[239] Research Center for Lifespan Development of Brain and Mind, Institute of Psychology, Chinese Academy of Sciences, Beijing 100101, China
[240] Department of Psychiatry, University of Cambridge, Cambridge, CB2 0SZ, UK.
[241] Division of Clinical Geriatrics, Center for Alzheimer Research, Department of Neurobiology, Care Sciences and Society, Karolinska Institutet, Stockholm, Sweden
[242] Ageing Epidemiology Research Unit, School of Public Health, Imperial College London, UK
[243] Generation Scotland, University of Edinburgh
[244] MRC Biostatistics Unit, University of Cambridge, Cambridge, England
[245] Faculty of Medicine, CRC 1052 'Obesity Mechanisms', University of Leipzig, Leipzig, 04103, Germany
[246] Department of Electrical and Computer Engineering, National University of Singapore, Singapore
[247] Centre for Sleep & Cognition and Centre for Translational MR Research, Yong Loo Lin School of Medicine, National University of Singapore, Singapore
[248] N.1 Institute for Health & Institute for Digital Medicine, National University of Singapore, Singapore
[249] Integrative Sciences and Engineering Programme (ISEP), National University of Singapore, Singapore
[250] Fetal Neonatal Neuroimaging and Developmental Science Center, Division of Newborn Medicine, Boston Children's Hospital, Harvard Medical School, Boston, MA 02115, USA
[251] Melbourne Neuropsychiatry Centre, University of Melbourne, Melbourne, Australia; Department of Biomedical Engineering, University of Melbourne, Melbourne, Australia.
[252] SAMRC Unit on Child & Adolescent Health, University of Cape Town, South Africa
[253] Center for Translational Magnetic Resonance Research, Yong Loo Lin School of Medicine, National University of Singapore, Singapore
[254] Wellcome Trust-MRC Institute of Metabolic Science, University of Cambridge, Cambridge, CB2 0SZ
[255] Cambridgeshire and Peterborough Foundation Trust, Cambridge, CB21 5EF
[256] National Institute of Mental Health (NIMH), National Institutes of Health (NIH), Bethesda, Maryland, USA
[257] Department of Psychiatry, Escola Paulista de Medicina, S√£o Paulo, Brazil.
[258] Research Center for Lifespan Development of Brain and Mind, Institute of Psychology, Chinese Academy of Sciences, Beijing 100101, China
[259] Key Laboratory of Brain and Education, School of Education Science, Nanning Normal University, Nanning 530001, China

**Aging Brain: Vasculature, Ischemia, and Behavior Study (ABVIB)**

Helena C. Chui M.D.[1,2] (Principal Investigator), Charles C. DeCarli, M.D.[3], William G. Ellis, M.D.[3], William J. Jagust, M.D.[4,5], Joel H. Kramer, Ph.D.[6], Meng Law, M.D.[7,8], Dan Mungas Ph,D.[3], Bruce R. Reed, Ph.D.[9], Nerses Sanossian, M.D.[2,10], Michael W. Weiner, M.D.[11,12,13,14,15,16], Wendy J. Mack, Ph.D.[17], Harry V. Vinters, M.D.[18,19], Chris Zarow, Ph.D.[2], Ling Zheng, Ph.D.[2]

[1] Alzheimer's Disease Research Center, Keck School of Medicine, University of Southern California, Los Angeles, CA, USA
[2]Department of Neurology, Keck School of Medicine, University of Southern California, Los Angeles, CA, USA
[3] Department of Neurology, University of California Davis School of Medicine, Sacramento, California, USA
[4] Department of Neuroscience, University of California Berkeley, Berkeley, CA, USA

[5] Lawrence Berkeley National Laboratory, Berkeley, CA, USA
[6] Memory and Aging Center, Department of Neurology, University of California, San Francisco, San Francisco, CA, USA
[7] Department of Radiology, The Alfred, Melbourne, VIC, Australia
[8] Department of Neuroscience, School of Translational Medicine, Monash University, Clayton, VIC, Australia
[9] National Institutes of Health Center for Scientific Review, Bethesda, MD, USA
[10] Roxanna Todd Hodges Stroke Program, University of Southern California, Los Angeles, CA, USA
[11] Department of Veterans Affairs Medical Center, Center for Imaging of Neurodegenerative Diseases, San Francisco, California, USA
[12] Department of Radiology and Biomedical Imaging, University of California San Francisco, San Francisco, California, USA
[13] Department of Medicine, University of California San Francisco, San Francisco, California, USA
[14] Department of Psychiatry and Behavioral Sciences, University of California San Francisco, San Francisco, California, USA
[15] Department of Neurology, University of California San Francisco, San Francisco, California, USA
[16] Northern California Institute for Research and Education (NCIRE), San Francisco, California, USA
[17] Population and Public Health Sciences, Keck School of Medicine of the University of Southern California, Los Angeles, California, USA
[18] Department of Pathology and Laboratory Medicine, David Geffen School of Medicine, University of California Los Angeles, Los Angeles, California, USA
[19] Department of Neurology, David Geffen School of Medicine, University of California Los Angeles, Los Angeles, California, USA

**Alzheimer's Disease Neuroimaging Initiative (ADNI)**
A complete listing of ADNI investigators can be found at:
http://adni.loni.usc.edu/wp-content/uploads/how_to_apply/ADNI_Acknowledgement_List.pdf

**Australian Imaging, Biomarkers and Lifestyle (AIBL)**
AIBL researchers are listed at www.aibl.csiro.au.

**Alzheimer's Disease Repository Without Borders (ARWiBo)**
The Principal Investigator of ARWIBO is Giovanni B. Frisoni, MD, University Hospitals and University of Geneva, Geneva, Switzerland, and IRCCS Fatebenefratelli, The National Centre for Alzheimer's and Mental Diseases, Brescia, Italy. ARWIBO is the result of effort of many researchers of IRCCS Fatebenefratelli: G. Binetti, MD, Neurobiology; L. Bocchio-Chiavetto, PhD, Neuropharmacology; M. Cotelli, PhD, Neuropsychology Unit; C. Minussi, PhD, Neurophysiology; M. Gennarelli, PhD, Genetic Unit; R. Ghidoni, PhD, Proteomics Unit; D. Moretti, MD, and O. Zanetti, MD, Alzheimer's Unit. A complete listing of ARWiBo researchers can be found at: https://www.arwibo.it/acknowledgments.pdf

**Biomarkers of Cognitive Decline among Normal Individuals: the BIOCARD Study**
A listing of BIOCARD investigators can be found on the BIOCARD website, https://biocard.pathology.jhu.edu/our-team/.

**Centre for Attention Learning and Memory (CALM)**
Duncan E. Astle[1,2]. More information on CALM team members can be found at: https://calm.mrc-cbu.cam.ac.uk/team/

[1] MRC Cognition and Brain Sciences Unit, University of Cambridge, Cambridge, UK

[2] Department of Psychiatry, University of Cambridge, Cambridge, UK

**Cambridge Center for Ageing and Neuroscience (Cam-CAN) study**
A complete list of Cam-CAN reasearchers can be found at: https://cam-can.mrc-cbu.cam.ac.uk/people/

**Chinese Color Nest Project (CCNP)**
Yin-Shan Wang[1,2], Ning Yang[1] and Xi-Nian Zuo[1,2,3,4]

[1] State Key Laboratory of Cognitive Neuroscience and Learning, Beijing Normal University, Beijing 100875, China
[2] Developmental Population Neuroscience Research Center, International Data Group/McGovern Institute for Brain Research, Beijing Normal University, Beijing 100875, China
[3] Research Center for Lifespan Development of Mind and Brain, Institute of Psychology, Chinese Academy of Sciences, Beijing 100101, China
[4] Department of Psychology, University of Chinese Academy of Sciences, Beijing 100049, China

**Centers of Biomedical Research Excellence (COBRE)**
Vince D. Calhoun[1]

[1] Tri-Institutional Center for Translational Research in Neuroimaging and Data Science (TReNDS), Georgia State University, Georgia Institute of Technology, and Emory University, Atlanta, Georgia, USA

**Developing Human Connectome Project (dHCP)**
Stephen M. Smith[1], Daniel Rueckert[2,3], Joseph V. Hajnald[4,5] and A. David Edwards[6,7]

[1] Nuffield Department of Clinical Neurosciences, Wellcome Centre for Integrative Neuroimaging, FMRIB, University of Oxford, Oxford, UK
[2] Chair for AI in Healthcare and Medicine, Technical University of Munich (TUM) and TUM University Hospital, Munich, Germany
[3] Department of Computing, Imperial College London, London, UK
[4] Early Life Imaging Department, School of Biomedical Engineering and Imaging Sciences, King's College London, London, UK
[5] Imaging Physics and Engineering Department, School of Biomedical Engineering and Imaging Sciences, King's College London, London, UK
[6] Centre for the Developing Brain, Research Department of Early Life Imaging, School of Biomedical Engineering and Imaging Sciences, King’s College London, London SE1 7EH, United Kingdom
[7] Medical Research Council Centre for Neurodevelopmental Disorders, King’s College London, London SE1 1UL, United Kingdom

**Harvard Aging Brain Study (HABS)**
Reisa A. Sperling[1,2] and Keith A. Johnson[1,2,3,4,5]

[1] Department of Neurology, Massachusetts General Hospital, Harvard Medical School, Boston, MA, USA
[2] Department of Neurology, Brigham and Women's Hospital, Harvard Medical School, Boston, MA, USA
[3] Molecular Neuroimaging, Massachusetts General Hospital, Boston, Massachusetts, USA
[4] Harvard Medical School, Boston, Massachusetts, USA
[5] Department of Radiology, Massachusetts General Hospital, Boston, Massachusetts, USA

**Human Connectome Project (HCP)**
The investigators for each HCP substudy can be found at: https://www.humanconnectome.org/

**International Consortium for Brain Mapping (ICBM)**
John C Mazziotta[1]

[1] UCLA School of Medicine, Los Angeles, USA

**IMAGEN**
Sylvane Desrivieres[1], Andreas Heinz[2], Tianye Jia[1,3,4], and Gunter Schumann[5,6]

[1] Social, Genetic and Developmental Psychiatry Centre, Institute of Psychiatry, Psychology and Neuroscience, King's College, London, UK
[2] Department of Psychiatry and Psychotherapy CCM, Charité-Universitätsmedizin Berlin, corporate member of Freie Universität Berlin, Humboldt-Universität zu Berlin, and Berlin Institute of Health, Berlin, Germany
[3] Institute of Science and Technology for Brain-Inspired Intelligence (ISTBI), Fudan University, Shanghai 200433, China
[4] Key Laboratory of Computational Neuroscience and Brain-Inspired Intelligence, Fudan University, Ministry of Education, Shanghai 200433, China.
[5] Centre for Population Neuroscience and Stratified Medicine (PONS), Department of Psychiatry and Neuroscience, Charité Universitätsmedizin Berlin, Berlin, Germany
[6] Centre for Population Neuroscience and Precision Medicine (PONS), Institute for Science and Technology of Brain-inspired Intelligence (ISTBI), Fudan University, Shanghai, China

**Neuroscience in Psychiatry Network (NSPN)**
A full list of NSPN consortium members can be found at: https://www.nspn.org.uk/nspn-team/

**Province of Ontario Neurodevelopmental Disorders (POND) Network**
Evdokia Anagnostou[1,2], Muhammad Ayub[3,4], Jennifer Crosbie[5,6,7,8], Christopher F. Hammill[9], Elizabeth A. Kelley[3,10,11], Jason Lerch[7,9,12], and Russell J. Schachar[5,6,7,8]

[1] Autism Research Centre, Bloorview Research Institute, Holland Bloorview Kids Rehabilitation Hospital, Toronto, ON, Canada
[2] Department of Pediatrics, Temerty Faculty of Medicine, University of Toronto, Toronto, Ontario, Canada
[3] Department of Psychiatry, Queen's University, Kingston, ON, Canada
[4] Division of Psychiatry, University College London, London, UK
[5] Department of Psychiatry, The Hospital for Sick Children, Toronto, ON, Canada
[6] Department of Psychiatry, Temerty Faculty of Medicine, University of Toronto, Toronto, ON, Canada
[7] Program in Neurosciences and Mental Health, Research Institute, The Hospital for Sick Children, Toronto, ON, Canada
[8] Genetics and Genome Biology, The Hospital for Sick Children, Toronto, ON, Canada
[9] Mouse Imaging Centre, Hospital for Sick Children, Toronto, ON, Canada
[10] Department of Psychology, Queen’s University, Kingston, ON, Canada
[11] Centre for Neuroscience Studies, Queen’s University, Kingston, ON, Canada
[12] Wellcome Centre for Integrative Neuroimaging, FMRIB, Nuffield Department of Clinical Neurosciences, University of Oxford, Oxford, UK

**Pre-symptomatic Evaluation of Experimental or Novel Treatments for Alzheimer Disease (PREVENT-AD)**
Alexa Pichet Binette[1] and Sylvia Villeneuve[2,3]

[1] Clinical Memory Research Unit, Department of Clinical Sciences Malmö, Lund University, Lund, Sweden
[2] Douglas Mental Health University Institute Research Centre, McGill University, Montréal H4H 1R3, Canada
[3] Department of Psychiatry, McGill University, Montréal H3A 1A1, Canada

**Vietnam Era Twin Study of Aging (VETSA)**
A complete list of VETSA researchers can be found at: https://www.vetsatwins.org/people/

#### E. Extended Acknowledgements

Data used in the preparation of this article were obtained from the Alzheimer's Disease Neuroimaging Initiative (ADNI) database (adni.loni.usc.edu). The ADNI was launched in 2003 as a public-private partnership, led by Principal Investigator Michael W. Weiner, MD. The primary goal of ADNI has been to test whether serial magnetic resonance imaging (MRI), positron emission tomography (PET), other biological markers, and clinical and neuropsychological assessment can be combined to measure the progression of mild cognitive impairment (MCI) and early Alzheimer's disease (AD). Data collection and sharing for this project was funded by the Alzheimer's Disease Neuroimaging Initiative (ADNI) (National Institutes of Health Grant U01 AG024904) and DOD ADNI (Department of Defense award number W81XWH-12-2-0012). ADNI is funded by the National Institute on Aging, the National Institute of Biomedical Imaging and Bioengineering, and through generous contributions from the following: AbbVie, Alzheimer's Association; Alzheimer's Drug Discovery Foundation; Araclon Biotech; BioClinica, Inc.; Biogen; Bristol-Myers Squibb Company; CereSpir, Inc.; Cogstate; Eisai Inc.; Elan Pharmaceuticals, Inc.; Eli Lilly and Company; EuroImmun; F. Hoffmann-La Roche Ltd and its affiliated company Genentech, Inc.; Fujirebio; GE Healthcare; IXICO Ltd.; Janssen Alzheimer Immunotherapy Research & Development, LLC.; Johnson  Johnson Pharmaceutical Research  Development LLC.; Lumosity; Lundbeck; Merck  Co., Inc.; Meso Scale Diagnostics, LLC.;  NeuroRx Research; Neurotrack Technologies; Novartis Pharmaceuticals Corporation; Pfizer Inc.; Piramal Imaging; Servier; Takeda Pharmaceutical Company; and Transition Therapeutics. The Canadian Institutes of Health Research is providing funds to support ADNI clinical sites in Canada. Private sector contributions are facilitated by the Foundation for the National Institutes of Health (www.fnih.org). The grantee organization is the Northern California Institute for Research and Education, and the study is coordinated by the Alzheimer's Therapeutic Research Institute at the University of Southern California. ADNI data are disseminated by the Laboratory for Neuro Imaging at the University of Southern California.

ICBM data are disseminated by the Laboratory of Neuro Imaging at the University of Southern California.

Data used in the preparation of this work were obtained from the Mind Clinical Imaging Consortium database through the Mind Research Network (www.mrn.org).

YOUth is funded through the Gravitation program of the Dutch Ministry of Education, Culture, and Science and the Netherlands Organization for Scientific Research (NWO grant number 024.001.003). A complete listing of the study investigators and study management can be found at https://www.uu.nl/en/research/youth-cohort-study/about-us/who-is-involved. YOUth investigators and management designed and implemented the study and/or provided data but did not necessarily participate in the analysis or writing of this report. This manuscript reflects the views of the authors and may not reflect the opinions or views of the YOUth study investigators or YOUth management.

We would like to acknowledge the individuals and organizations that have made Data used for this research available including CAM-BIND, the Ontario Brain Institute, the Brain-CODE platform, and the Government of Ontario.

The SRPS1600 dataset was provided by the DecNef Department at the Advanced Telecommunication Research Institute International, Kyoto, Japan.

Data used in the preparation of this article were obtained from the Adolescent Brain Cognitive Development (ABCD) Study (https://abcdstudy.org), held in the NIMH Data Archive (NDA). This is a multisite, longitudinal study designed to recruit more than 10,000 children age 9-10 and follow them over 10 years into early adulthood. The ABCD Study® is supported by the National Institutes of Health and additional federal partners under award numbers U01DA041048, U01DA050989, U01DA051016, U01DA041022, U01DA051018, U01DA051037, U01DA050987, U01DA041174, U01DA041106, U01DA041117, U01DA041028, U01DA041134, U01DA050988, U01DA051039, U01DA041156, U01DA041025, U01DA041120, U01DA051038, U01DA041148, U01DA041093, U01DA041089, U24DA041123, U24DA041147. A full list of supporters is available at https://abcdstudy.org/federal-partners.html. A listing of participating sites and a complete listing of the study investigators can be found at https://abcdstudy.org/consortium_members/.
The ABCD data repository grows and changes over time. The ABCD data used in this report came from 10.15154/z563-zd24.

Data used in the preparation of this article were obtained from the Aging Brain: Vasculature, Ischemia, and Behavior Study database (https://ida.loni.usc.edu/login.jsp?project=ABVIB). The ABVIB study was launched in 1994 as a NIA-funded program project led by Principal Investigator Helena C. Chui MD. Data from a second study cohort was started in 2008-2013 and is included in the current database. Data collection and sharing for ABVIB was funded by the National Institutes on Aging (NIA) P01 AG12435.

We would like to acknowledge the primary funding source for the ABIDE II dataset (NIMH 5R21MH107045).

Data used in the preparation of this article was obtained from the Australian Imaging Biomarkers and Lifestyle flagship study of ageing (AIBL) funded by the Commonwealth Scientific and Industrial Research Organisation (CSIRO) which was made available at the ADNI database (www.loni.usc.edu/ADNI).

ARWiBo data collection and sharing for this project was supported by the Italian Ministry of Health, under the following grant agreements: Ricerca Corrente IRCCS Fatebenefratelli, Linea di Ricerca 2; Progetto Finalizzato Strategico 2000-2001 “Archivio normativo italiano di morfometria cerebrale con risonanza magnetica (età 40+)”; Progetto Finalizzato Strategico 2000-2001 “Decadimento cognitivo lieve non dementigeno: stadio preclinico di malattia di Alzheimer e demenza vascolare. Caratterizzazione clinica, strumentale, genetica e neurobiologica e sviluppo di criteri diagnostici utilizzabili nella realtà nazionale,”; Progetto Finalizzata 2002 “Sviluppo di indicatori di danno cerebrovascolare clinicamente significativo alla risonanza magnetica strutturale”; Progetto Fondazione CARIPLO 2005-2007 “Geni di suscettibilità per gli endofenotipi associati a malattie psichiatriche e dementigene”; “Fitness and Solidarietà”; and anonymous donors.

This paper uses data collected in the Accelerating Medicines Partnership in Schizophrenia (AMP SCZ) project. AMP SCZ is supported by NIMH grants U24MH124629, U01MH124631, U01MH124639. The research was also funded in part by the Welcome Trust (220664/Z/20/Z).

The BCP was supported by grants: U01MH110274, R01MH104324.

Data were provided in part by the Brain Genomics Superstruct Project of Harvard University and the Massachusetts General Hospital, (Principal Investigators: Randy Buckner, Joshua Roffman, and Jordan Smoller), with support from the Center for Brain Science Neuroinformatics Research Group, the Athinoula A. Martinos Center for Biomedical Imaging, and the Center for Human Genetic Research. 20 individual investigators at Harvard and MGH generously contributed data to GSP.

BHRCS was supported with grants from the National Institute of Development Psychiatric for Children and Adolescent (INPD). Grant: Fapesp 2014/50917-0 - CNPq 465550/2014-2

The BIODEP study was sponsored by the Cambridgeshire and Peterborough NHS Foundation Trust and the University of Cambridge, and funded by a strategic award from the Wellcome Trust (104025) in partnership with Janssen, GlaxoSmithKline, Lundbeck and Pfizer.

Data collection and sharing for this project was provided by the Centre for Attention, Learning and Memory (CALM). CALM funding was provided by the UK Medical Research Council and University of Cambridge, UK. Data used in the preparation of this work were obtained from CALM resource: https://calm.mrc-cbu.cam.ac.uk/. The study protocol is reported in Holmes et al, 2019, *BMC Pediatr*, doi:10.1186/s12887-018-1385-3.

We would like to acknowledge the individuals and organizations that have made Data used for this research available including the Ontario Brain Institute, the Brain-CODE platform, and the Government of Ontario.

Data collection and sharing for this project was provided by the Cambridge Centre for Ageing and Neuroscience (CamCAN). CamCAN funding was provided by the UK Biotechnology and Biological Sciences Research Council (grant number BB/H008217/1), together with support from the UK Medical Research Council and University of Cambridge, UK.

The CHILD study was funded by the Autism Research Trust.

The Drakenstein Child Health Study is funded by the Bill and Melinda Gates Foundation (OPP 1017641).

The Developing Human Connectome Project was supported by the European Research Council under the European Union Seventh Framework Programme (FP/2007-2013)/ERC Grant Agreement No. 319456.

Data were provided in part by the Human Connectome Project, WU-Minn Consortium (Principal Investigators: David Van Essen and Kamil Ugurbil; 1U54MH091657) funded by the 16 NIH Institutes and Centers that support the NIH Blueprint for Neuroscience Research; and by the McDonnell Center for Systems Neuroscience at Washington University.

Research using Human Connectome Project for Early Psychosis (HCP-EP) data reported in this publication was supported by the National Institute of Mental Health of the National Institutes of Health under Award Number U01MH109977. The HCP-EP 1.1 Release data used in this report came from DOI: 10.15154/1522899.

FinnBrain was funded by Jane and Aatos Erkko Foundation, Signe and Ane Gyllenberg Foundation

GUSTO study is supported by the Singapore National Research Foundation under its Translational and Clinical Research (TCR) Flagship Programme and administered by the Singapore Ministry of Health's National Medical Research Council (NMRC), Singapore - NMRC/TCR/004-NUS/2008; NMRC/TCR/012-NUHS/2014. Additional funding is provided by the Singapore Institute for Clinical Sciences, Agency for Science Technology and Research (A*STAR), Singapore; E.C. is supported by grants: NIMH P50163 MH081755, NIMH R01-MH036840, NIMH R01-MH110558, NIMH U01-MH108898, NIDCD R01DC016385.

Data used in the preparation of this work were obtained from the International Consortium for Brain Mapping (ICBM) database (www.loni.usc.edu/ICBM). The ICBM project (Principal Investigator John Mazziotta, M.D., University of California, Los Angeles) is supported by the National Institute of Biomedical Imaging and BioEngineering. ICBM is the result of efforts of co-investigators from UCLA, Montreal Neurologic Institute, University of Texas at San Antonio, and the Institute of Medicine, Juelich/Heinrich Heine University - Germany.

IMAP study (PI (scientific): G Chetelat; PI (MD) V de La Sayette)) was funded by Programme Hospitalier de Recherche Clinique (PHRCN 2011-A01493-38 and PHRCN 2012 12-006-0347) and Agence Nationale de la Recherche (LONGVIE 2007). Dr Chetelat's research including IMAP was also funded by Institut National de la Santé et de la Recherche Médicale (Inserm), Fondation Plan Alzheimer (Alzheimer Plan 2008-2012).

LIFE is funded by means of the European Union, by the European Regional Development Fund (ERDF) and by funds of the Free State of Saxony within the framework of the excellence initiative.

Data used in the preparation of this work were obtained in part from the Mind Clinical Imaging Consortium database through the Mind Research Network (www.mrn.org). The MCIC project was supported by the Department of Energy under Award Number DE-FG02-08ER64581. MCIC is the result of efforts of co-investigators from the University of Iowa, University of Minnesota, University of New Mexico, and Massachusetts General Hospital, who collected and shared the imaging data and demographic information.

The NACC database is funded by NIA/NIH Grant U24 AG072122. NACC data are contributed by the NIA-funded ADRCs: P30 AG062429 (PI James Brewer, MD, PhD), P30 AG066468 (PI Oscar Lopez, MD), P30 AG062421 (PI Teresa Gomez-Isla, MD), P30 AG066509 (PI Thomas Grabowski, MD), P30 AG066514 (PI Mary Sano, PhD), P30 AG066530 (PI Helena Chui, MD, Arthur Toga, PhD), P30 AG066507 (PI Marilyn Albert, PhD), P30 AG066444 (PI David Holtzman, MD), P30 AG066518 (PIs Lisa Silbert, MD, Kevin Duff, PhD), P30 AG066512 (PI Thomas Wisniewski, MD), P30 AG066462 (PI Scott Small, MD), P30 AG072979 (PI David Wolk, MD), P30 AG072972 (PIs Charles DeCarli, MD, Rachel Whitmer, PhD), P30 AG072976 (PI Andrew Saykin, PsyD), P30 AG072975 (PI Julie Schneider, MD, MS), P30 AG072978 (PI Ann McKee, MD), P30 AG072977 (PI Robert Vassar, PhD), P30 AG066519 (PI Joshua Grill, PhD), P30 AG062677 (PIs Brad Boeve, MD, Ronald Petersen, MD, PhD), P30 AG079280 (PI Jessica Langbaum, PhD), P30 AG062422 (PI Gil Rabinovici, MD), P30 AG066511 (PI Allan Levey, MD, PhD), P30 AG072946 (PI Linda Van Eldik, PhD), P30 AG062715 (PI Sanjay Asthana, MD, FRCP), P30 AG072973 (PI Russell Swerdlow, MD), P30 AG066506 (PIs Glenn Smith, PhD, ABPP, David Lowenstein, PhD, Ranjan Duara, MD), P30 AG066508 (PIs Stephen Strittmatter, MD, PhD, Christopher Van Dyck, MD), P30 AG066515 (PI Victor Henderson, MD, MS), P30 AG072947 (PI Suzanne Craft, PhD), P30 AG072931 (PI Henry Paulson, MD, PhD), P30 AG066546 (PIs Sudha Seshadri, MD, Gladys Maestre, MD, PhD), P30 AG086401 (PI Erik Roberson, MD, PhD), P30 AG086404 (PI Gary Rosenberg, MD), P30 AG086403 (PI Angela Jefferson, PhD), P30 AG072958 (PIs Heather Whitson, MD, Gwenn Garden, MD, PhD), P30 AG072959 (PI Jagan Pillai, MD, PhD), P30 AG092752 (Ihab Hajjar, MD, MS). Data collected by SCAN and shared by NACC are contributed by the NIA-funded ADRCs as follows: Arizona Alzheimer's Center - P30 AG072980 (PI: Eric Reiman, MD); R01 AG069453 (PI: Eric Reiman (contact), MD); P30 AG019610 (PI: Eric Reiman, MD); and the State of Arizona which provided additional funding supporting our center; Boston University - P30 AG013846 (PI Neil Kowall MD); Cleveland ADRC - P30 AG062428 (James Leverenz, MD); Cleveland Clinic, Las Vegas – P20AG068053; Columbia - P50 AG008702 (PI Scott Small MD); Duke/UNC ADRC – P30 AG072958; Emory University - P30AG066511 (PI Levey Allan, MD, PhD); Indiana University - R01 AG19771 (PI Andrew Saykin, PsyD); P30 AG10133 (PI Andrew Saykin, PsyD); P30 AG072976 (PI Andrew Saykin, PsyD); R01 AG061788 (PI Shannon Risacher, PhD); R01 AG053993 (PI Yu-Chien Wu, MD, PhD); U01 AG057195 (PI Liana Apostolova, MD); U19 AG063911 (PI Bradley Boeve, MD); and the Indiana University Department of Radiology and Imaging Sciences; Johns Hopkins - P30 AG066507 (PI Marilyn Albert, Phd.); Mayo Clinic - P50 AG016574 (PI Ronald Petersen MD PhD); Mount Sinai - P30 AG066514 (PI Mary Sano, PhD); R01 AG054110 (PI Trey Hedden, PhD); R01 AG053509 (PI Trey Hedden, PhD); New York University - P30AG066512-01S2 (PI Thomas Wisniewski, MD); R01AG056031 (PI Ricardo Osorio, MD); R01AG056531 (PIs Ricardo Osorio, MD; Girardin Jean-Louis, PhD); Northwestern University - P30 AG013854 (PI Robert Vassar PhD); R01 AG045571 (PI Emily Rogalski, PhD); R56 AG045571, (PI Emily Rogalski, PhD); R01 AG067781, (PI Emily Rogalski, PhD); U19 AG073153, (PI Emily Rogalski, PhD); R01 DC008552, (M.-Marsel Mesulam, MD); R01 AG077444, (PIs M.-Marsel Mesulam, MD, Emily Rogalski, PhD); R01 NS075075 (PI Emily Rogalski, PhD); R01 AG056258 (PI Emily Rogalski, PhD); Oregon Health and Science University - P30 AG008017 (PI Jeffrey Kaye MD); R56 AG074321 (PI Jeffrey Kaye, MD); Rush University - P30 AG010161 (PI David Bennett MD); Stanford – P30AG066515; P50 AG047366 (PI Victor Henderson MD MS); University of Alabama, Birmingham – P20; University of California, Davis - P30 AG10129 (PI Charles DeCarli, MD); P30 AG072972 (PI Charles DeCarli, MD); University of California, Irvine - P50 AG016573 (PI Frank LaFerla PhD); University of California, San Diego - P30AG062429 (PI James Brewer, MD, PhD); University of California, San Francisco - P30 AG062422 (Rabinovici, Gil D., MD); University of Kansas - P30 AG035982 (Russell Swerdlow, MD); University of Kentucky - P30 AG028283-15S1 (PIs Linda Van Eldik, PhD and Brian Gold, PhD); University of Michigan ADRC - P30AG053760 (PI Henry Paulson, MD, PhD) P30AG072931 (PI Henry Paulson, MD, PhD) Cure Alzheimer's Fund 200775 - (PI Henry Paulson, MD, PhD) U19 NS120384 (PI Charles DeCarli, MD, University of Michigan Site PI Henry Paulson, MD, PhD) R01 AG068338 (MPI Bruno Giordani, PhD, Carol Persad, PhD, Yi Murphey, PhD) S10OD026738-01 (PI Douglas Noll, PhD) R01 AG058724 (PI Benjamin Hampstead, PhD) R35 AG072262 (PI Benjamin Hampstead, PhD) W81XWH2110743 (PI Benjamin Hampstead, PhD) R01 AG073235 (PI Nancy Chiaravalloti, University of Michigan Site PI Benjamin Hampstead, PhD) 1I01RX001534 (PI Benjamin Hampstead, PhD) IRX001381 (PI Benjamin Hampstead, PhD); University of New Mexico - P20 AG068077 (Gary Rosenberg, MD); University of Pennsylvania - State of PA project 2019NF4100087335 (PI David Wolk, MD); Rooney Family Research Fund (PI David Wolk, MD); R01 AG055005 (PI David Wolk, MD); University of Pittsburgh - P50 AG005133 (PI Oscar Lopez MD); University of Southern California - P50 AG005142 (PI Helena Chui MD); University of Washington - P50 AG005136 (PI Thomas Grabowski MD); University of Wisconsin - P50 AG033514 (PI Sanjay Asthana MD FRCP); Vanderbilt University – P20 AG068082; Wake Forest - P30AG072947 (PI Suzanne Craft, PhD); Washington University, St. Louis - P01 AG03991 (PI John Morris MD); P01 AG026276 (PI John Morris MD); P20 MH071616 (PI Dan Marcus); P30 AG066444 (PI John Morris MD); P30 NS098577 (PI Dan Marcus); R01 AG021910 (PI Randy Buckner); R01 AG043434 (PI Catherine Roe); R01 EB009352 (PI Dan Marcus); UL1 TR000448 (PI Brad Evanoff); U24 RR021382 (PI Bruce Rosen); Avid Radiopharmaceuticals / Eli Lilly; Yale - P50 AG047270 (PI Stephen Strittmatter MD PhD); R01AG052560 (MPI: Christopher van Dyck, MD; Richard Carson, PhD); R01AG062276 (PI: Christopher van Dyck, MD); 1Florida - P30AG066506-03 (PI Glenn Smith, PhD); P50 AG047266 (PI Todd Golde MD PhD)

Data and/or research tools used in the preparation of this manuscript were obtained from the National Institute of Mental Health (NIMH) Data Archive (NDA). NDA is a collaborative informatics system created by the National Institutes of Health to provide a national resource to support and accelerate research in mental health. Dataset identifier(s): 2400, 2847, 2846, 2275, 2274, 2199, 2021, 1890, 19, 2936, 3142, 2803, 2607. This manuscript reflects the views of the authors and may not reflect the opinions or views of the NIH or of the Submitters submitting original data to NDA.

Data were provided in part by OASIS-3: Longitudinal Multimodal Neuroimaging: Principal Investigators: T. Benzinger, D. Marcus, J. Morris; NIH P30 AG066444, P50 AG00561, P30 NS09857781, P01 AG026276, P01 AG003991, R01 AG043434, UL1 TR000448, R01 EB009352. AV-45 doses were provided by Avid Radiopharmaceuticals, a wholly owned subsidiary of Eli Lilly.

Data used in the preparation of this article was obtained on 2023-09-23 from the Parkinson's Progression Markers Initiative (PPMI) database (www.ppmi-info.org/access-dataspecimens/download-data; Tier 1), RRID:SCR_006431. For up-to-date information on the study, visit www.ppmi-info.org. PPMI – a public-private partnership – is funded by the Michael J. Fox Foundation for Parkinson's Research, and funding partners; including  including 4D Pharma, Abbvie, AcureX, Allergan, Amathus Therapeutics, Aligning Science Across Parkinson's, AskBio, Avid Radiopharmaceuticals, BIAL, BioArctic, Biogen, Biohaven, BioLegend, BlueRock Therapeutics, Bristol-Myers Squibb, Calico Labs, Capsida Biotherapeutics, Celgene, Cerevel Therapeutics, Coave Therapeutics, DaCapo Brainscience, Denali, Edmond J. Safra Foundation, Eli Lilly, Gain Therapeutics, GE HealthCare, Genentech, GSK, Golub Capital, Handl Therapeutics, Insitro, Jazz Pharmaceuticals, Johnson & Johnson Innovative Medicine, Lundbeck, Merck, Meso Scale Discovery, Mission Therapeutics, Neurocrine Biosciences, Neuron23, Neuropore, Pfizer, Piramal, Prevail Therapeutics, Roche, Sanofi, Servier, Sun Pharma Advanced Research Company, Takeda, Teva, UCB, Vanqua Bio, Verily, Voyager Therapeutics, the Weston Family Foundation and Yumanity Therapeutics.

The POND study was supported by the Ontario Brain Institute (grant number IDS-I 1-02).

The results published here are in whole or in part based on data obtained from the AD Knowledge Portal (https://adknowledgeportal.org). Study data were provided by the Rush Alzheimer's Disease Center, Rush University Medical Center, Chicago. Data collection was supported through funding by NIA grants P30AG10161 (ROS), R01AG15819 (ROSMAP; genomics and RNAseq), R01AG17917 (MAP), R01AG30146, R01AG36042 (5hC methylation, ATACseq), RC2AG036547 (H3K9Ac), R01AG36836 (RNAseq), R01AG48015 (monocyte RNAseq) RF1AG57473 (single nucleus RNAseq), U01AG32984 (genomic and whole exome sequencing), U01AG46152 (ROSMAP AMP-AD, targeted proteomics), U01AG46161(TMT proteomics), U01AG61356 (whole genome sequencing, targeted proteomics, ROSMAP AMP-AD), the Illinois Department of Public Health (ROSMAP), and the Translational Genomics Research Institute (genomic). Additional phenotypic data can be requested at www.radc.rush.edu.

The content is the sole responsibility of the authors and does not necessarily represent official views of the NIA, NIH, or VA. The U.S. Department of Veterans Affairs, Department of Defense; National Personnel Records Center, National Archives and Records Administration; National Opinion Research Center; National Research Council, National Academy of Sciences; and the Institute for Survey Research, Temple University provided invaluable assistance in the creation of the VET Registry. The Cooperative Studies Program of the U.S. Department of Veterans Affairs provided financial support for development and maintenance of the Vietnam Era Twin Registry. We would also like to acknowledge the continued cooperation and participation of the members of the VET Registry and their families.

Data collection for the Vaghi datasets were supported by a Wellcome Trust Senior Investigator Award Grant No. 104631/Z/14/Z.

#### F. Captions for Other Supplementary Materials

**Supplemental Demographics Table: Study-level demographics and references**
This attached CSV contains detailed demographic and reference information for each of the studies included in the Lifespan Brain Chart Consortium.

**Supplemental Data S1: Results of sex effects significance testing**
This tab of the Supplemental Data contains the results of all likelihood ratio tests assessing the significance of age-by-sex and/or sex terms. Specifically, these data underlie Fig 1B, Fig 2D, and Figs S5 to S17, with each row corresponding to one likelihood ratio test.

The first column, “total_size_covar”, indicates whether the models tested were controlling for total brain size: TRUE indicates the model is size-corrected (e.g. Fig 2D) while FALSE indicates it is not (e.g. Fig 1B). The “effect_tested” column indicates what term(s) significance was being tested; a value of “age-varying sex component” indicates the age-by-sex interaction was being tested, while “any sex effect” indicates both the age-by-sex interaction and sex intercept. Note that “any sex effect” was tested only in models correcting for total size (i.e. “total_size_covar” is TRUE). The column “surfhole_weighted” is TRUE when the test was part of our sensitivity analyses in which data points were weighted inversely to their normalized surface hole count (Figs S5 & S8). “idp_category” broadly groups imaging derived phenotypes tested (noted in “idp”) as global measures (“Global Vols”), ventricular volumes (“Ventricular Vol”), subcortical volumes (“Subcortical Vol”), or cortical regions’ volumes (“Regional Vol”), thicknesses (“Regional CT”) or surface areas (“Regional SA”). “cv_sample” indicates which split-half, A or B, the test model was fitted and the likelihood ratio test(s) were performed in. Finally, the results of the likelihood ratio tests are reported in the remaining columns: chi statistic (“chi”); degrees of freedom (“df”); uncorrected p-value (“p.val”); Cohen’s F-squared effect size (“fsq”); the RESI S effect size (“resi”); p-values FDR-corrected within each level of ““total_size_covar”, “cv_sample” and “surfhole_weighted” (“p.val_fdr”); and an indicator of whether the test is significant at $q < 0.05$ (“significant_fdr”)

**Supplemental Data S2: Results of case-control t-tests**
This tab of the Supplemental Data contains the results of all t-tests comparing centile z-scores in individuals with psychiatric diagnoses to site-matched controls.

The first column, “total_size_covar”, indicates whether the centile scores tested were derived from models controlling for total brain size. The “model” column indicates whether centile scores were derived from sex-moderated models with an age-by-sex interaction (“sex_with_age_moderator”) or traditional models with only a sex intercept (“sex_intercept_only”). “dx_tested” indicates which neuropsychiatric conditions’ structural correlates are being tested. “centile_group” indicates whether the test is on the proportion of extremely low (“below_5th”) or high (“above_95th”) percentiles. “idp_category” broadly groups imaging derived phenotypes tested (noted in “idp”) as global measures (“Global Vols”), ventricular volumes (“Ventricular Vol”), subcortical volumes (“Subcortical Vol”), or cortical regions’ volumes (“Regional Vol”), thicknesses (“Regional CT”) or surface areas (“Regional SA”). The columns "percent_control", “percent_case”, and “percent_diff” index the percentage of cases and controls in the given “centile_group”, as well as their difference. The remaining columns show the results of Fisher’s exact test of proportions, specifically: uncorrected p-value (“p.val”); p-values FDR-corrected within each level of ““total_size_covar” and “model” (“p.val_fdr”); and an indicator of whether the test is significant at $q < 0.05$ (“significant_fdr”).

**Supplemental Data S3: Results of case-control extremeness analyses**
This tab of the Supplemental Data contains the results of all t-tests comparing the proportions of extremely high centiles (i.e. above the 95th percentile) and extremely low centiles (below the 5th percentile) in individuals with psychiatric diagnoses to site-matched controls.
